# Far-from-equilibrium assembly of multimers through DNA-based catalytic templating

**DOI:** 10.64898/2026.04.13.718267

**Authors:** Rakesh Mukherjee, Merry Mitra, Križan Jurinović, Jordan Juritz, Thomas E. Ouldridge

## Abstract

On-demand assembly of arbitrary, sequence-defined polymers from a pool of monomers is a major challenge in modern chemistry, towards which limited progress has been made. By contrast, biological systems routinely use information-bearing DNA and RNA templates to catalytically synthesize a precise, far-from-equilibrium ensemble of nucleic acid and protein sequences from the available pools of nucleoside triphosphates or aminoacyl-tRNAs. Inspired by these biological examples, we introduce an enzyme-free DNA strand displacement network in which single-stranded DNA sequences template the assembly of specific non-covalent DNA multimers of up to length five, under isothermal and autonomous conditions. The templates overcome product inhibition and can thereby catalyse the formation of a far-from-equilibrium ensemble of long-lived metastable products that are not otherwise addressable.

## Main

The ability to assemble on-demand, sequence-defined polymeric products from a mixture of building blocks has been described as a ‘‘Holy Grail’’ of chemistry (*1–3*). If such a feat were possible with arbitrary molecular building blocks, it would open the door to a new generation of designer complexes and materials (*4–9*) with a combinatorially large and easily accessible design space (*10–15*). In isolation, a small set of distinct building blocks can only reliably assemble a small set of structures (*16*). Biology circumvents this problem when constructing proteins and RNA by using the information in molecular templates to direct ribosome or polymerase-mediated assembly of amino acid or nucleotide sequences, respectively. Crucially, the template itself must act as a catalyst (*17*), accelerating the production of a specific sequence but not itself being consumed by the reaction. Without catalysis, a new template would be needed for each product, recreating the original challenge mitigated by the template. Since catalysts cannot affect equilibrium, highly specific catalytic templating necessarily drives systems extremely far from equilibrium by enriching certain product sequences relative to a pool of similarly stable alternatives (*17, 18*). Additionally, the nonequilibrium templating and folding process can force the product sequence into a metastable state that is far from the most stable one, even given the set of building blocks involved. For example, the functional states of molecules like serpins (*19*), cellular prion-related proteins (*20*), and viral fusion proteins (*21*) are metastable (*22*).

Ribosomes and polymerases are tuned to their natural building blocks; expanding the genetic alphabet to repurpose these biological mechanisms (*23*) is inherently limited in scope. Synthetic organic molecules (*24–26*), biopolymers such as peptides and nucleic acids (*27–29*), and bespoke materials (*30, 31*) have been used to template the assembly of a range of synthetic building blocks into target products. However, in these synthetic contexts, the specific attractive interactions between the monomers and the template, which make a template a good assembler, also tend to make it a poor catalyst. These attractive interactions typically act cooperatively to prevent the product spontaneously dissociating from the template, thereby inhibiting its participation in subsequent reaction cycles.

This effect, which is commonly known as ‘‘product inhibition’’, typically becomes exponentially worse with product length and is usually crippling for synthetic molecular templating systems that are not supported by highly-evolved enzymes or externally driven by non-chemical means (*32*). Recently, we (*33–35*) and others (*36–39*) have proposed strategies for overcoming product inhibition in which the product’s binding to the template is disrupted as the bond between its constituent monomer units are formed. However, effective, sequence-specific catalytic templating has not previously been demonstrated for products larger than dimers. Here we show that this approach can be applied to template the assembly of sequence-specific trimeric, tetrameric and pentameric products of base-paired DNA strands with catalytic turnover.

### A candidate mechanism for catalytic templating of multimers

The principle we employ is illustrated in Figure 1A. The monomers have a recognition interface and ‘‘right’’ and ‘‘left’’ polymerisation interfaces. Polymerisation is, however, inhibited by ‘‘blockers’’, molecules that are initially bound to the right polymerisation interfaces. Transient binding to the template can occur via the recognition interfaces, triggering full binding via the right polymerisation interface and release of the blockers. Once adjacent monomers are present on the template, their left and right polymerisation interfaces can bind to each other, because the right interfaces interact more strongly with neighbouring left interfaces than with the template. When this polymerisation reaction happens, the interaction of the left-hand monomer with the template is weakened. End sites of the template can be designed to recognise one-sided monomers, allowing a product to eventually form on the template with all its polymerisation interfaces incorporated into the product. At this point, the product is bound by only the weak recognition interfaces and can detach, releasing the template for further rounds of catalysis. Overall, the proposed mechanism uses recognition interactions with a template to assemble a specific product sequence that is not held together by these recognition interactions – just as the RNA and protein sequences assembled by transcription and translation are not held together by the base-pair recognition interactions that direct their synthesis. Product inhibition is overcome by diverting some of the free energy of product formation into disrupting interactions with the template.

**Figure 1:**
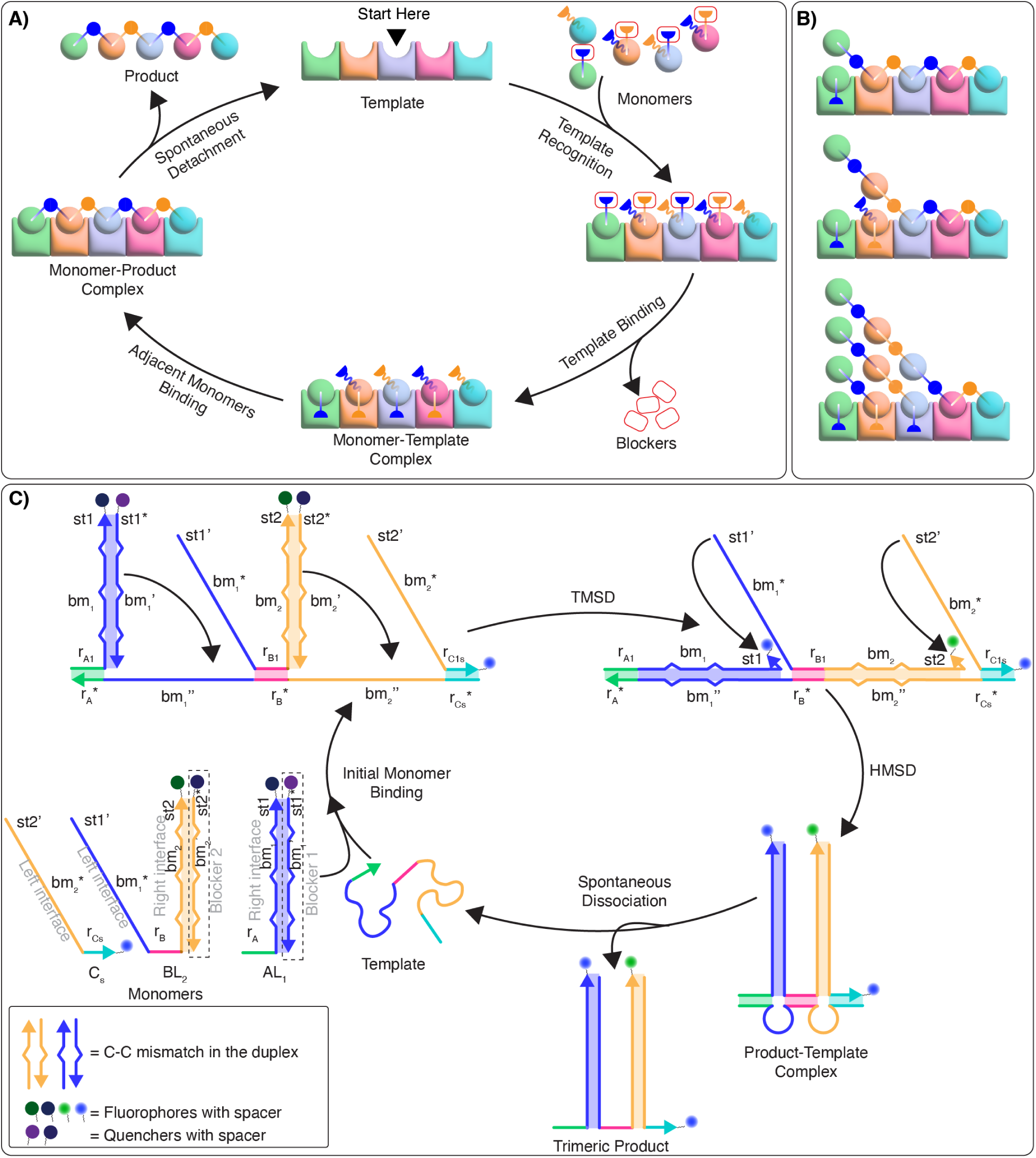
Design of catalytic templating networks. **A)** A schematic diagram of the intended templating mechanism. The spheres with protruding lines represent monomers with polymerization interfaces. The green and cyan spheres contain only one polymerization interface, and are the initial and terminal monomers. The other monomers with two polymerization interfaces are the propagating monomers. Initially, right-hand interfaces are blocked. The monomers are first recognised by the same-coloured grooves of the template, binding via their recognition interface and subsequently by their right-hand polymerization interfaces, releasing the blockers. Note that it is not expected that all monomers bind via their recognition interfaces prior to any blocker release, as illustrated here for simplicity. Polymerization occurs when right-hand interfaces are transferred from the template to an adjacent monomer. The final product then detaches spontaneously from the template, enabling further reaction cycles. **B)** Putative ‘‘brush’’ configurations where the templates are attached to multiple different intermediates. **C)** A domain-level representation of a DNA-based design of such a templating network for trimerization. The strands *A* and *B* bind to the template *via* recognition domain-triggered TMSD, releasing blocker strands *L*_1_ and *L*_2_. The *bm_1_*^j^ domain of *B* can then bind to the complementary template-bound domain *bm*_1_ of *A via* HMSD. Similarly, *C_s_* can perform HMSD after using its recognition domain as a handhold, forming a bond between *bm*_2_ and *bm*^j^_2_. These HMSD reactions lead to the formation of product-template complex, where the trimer and the template are bound by only three short recognition domains that can spontaneously dissociate, releasing the template for further reaction cycles.

The proposed mechanism faces challenges when extended beyond dimerisation to longer templates (*40*). First, the reactions may not happen in the order pictured in Figure 1A. It is possible, for example, that a subset of monomers bind to the right-hand end of the template, polymerise, and detach before the left-hand monomers arrive, resulting in left-hand truncated products. Second, although the interactions of the recognition interfaces are relatively weak, they will still be cooperative. Sufficiently long products will then still adhere strongly to the template, resulting in significant product inhibition and rendering the network non-catalytic.

Recently, we discovered a theoretical mechanism, illustrated in Figure 1B (see also figure S7, S8), with the potential to simultaneously overcome both of these challenges (*40*). If the recognition interactions are of moderate strength, products will tend to bind to the template cooperatively. However, transient unbinding of the relatively weak individual recognition interactions on the left-hand side allows new monomers to bind from solution and effectively displace the product from the template, due to the fact that these invading monomers can form stronger bonds with the template via both their recognition and right hand polymerisation interfaces. Eventually, given the right balance of binding strengths and reaction rates, the template can reach a steady state with a ‘‘brush’’ of partially formed products that advance to the right, pushing the product ahead of them off the template and thereby overcoming cooperative product inhibition. This dense conveyor belt of products also serves to limit the formation of partially-formed products that are truncated on their left hand side, since monomers that bind to the middle of the template are swept up by partially assembled products on their left.

### Implementing catalytic templating of multimers in an enzyme-free, DNA-based system

To implement the mechanism of Figure 1 in a synthetic molecular system, we apply two DNA strand displacement motifs in tandem (Figure 1C): toehold-mediated strand displacement (TMSD) (*41*) and handhold-mediated strand displacement (HMSD) (*42*). TMSD occurs when a single-stranded DNA (ssDNA) molecule (the *Invader*) binds to an extended complementary single-stranded domain (the toehold) of the *Target* strand in a *Target-Incumbent* duplex, and gradually displaces the *Incumbent* strand to form the *Target-Invader* duplex. In HMSD, the initial *Invader* binding occurs with the handhold, which is a single-stranded overhang on the *Incumbent* strand. Subsequently, the *Invader* binds to a very small (typically 1-2 nucleotide) secondary toehold on the *Target* strand and competes with the *Incumbent* for binding to the *Target*, eventually forming a *Target-Invader* duplex.

In the context of the mechanism of Figure 1A and B, TMSD has the appropriate chemical logic for recognition and blocker release. HMSD has the appropriate logic for the binding of left and right polymerisation domains between two adjacent, template-bound monomers. For DNA strands, domains – contiguous sections of bases that act as a unit – play the role of the interfaces in Figure 1A.

A domain-level illustration of a proposed DNA-based trimer-forming system is given in Figure 1C. The system comprises the initial monomer *A* (in a blocked complex *AL*_1_), the propagating monomer *B* (in a blocked complex *BL*_2_), the terminating monomer *C_s_*, and the trimeric ssDNA template ^3^*T_ABC_*. *A* has only a right polymerisation interface (*bm*_1_ + *st*_1_), *C_s_* only a left 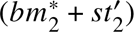, and *B* has both (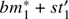 and *bm*_2_ + *st*_2_). All possess a unique recognition interface (*r _A_, r_B_, r_Cs_*) allowing them to bind to the template at the appropriate site. *A* and *B* can bind to the template via TMSD, releasing their blocking strands. *B* can then bind to *A* via HMSD. The rightmost monomer *C_s_* has only a recognition interaction with the template, but can use that to transiently bind to the template and then to *B* via HMSD. The network is then in principle capable of performing the reactions required by the proposed mechanism of Figure 1.

A number of design features are necessary to ensure that the system has the appropriate kinetic and thermodynamic properties. First, two different types of polymerisation interfaces are required to prevent monomer *B* from forming a highly stable intramolecular hairpin (figure S3) due to the flexibility of DNA oligomers. For the longer sequences considered later, these two interfaces will alternate – it is not necessary for each interface in a longer product to be unique, and they are not recognition elements. Second, domains *bm_i_*^j^ are complementary to *bm_i_* with the exception of three C-C mismatches. These mismatches, buried deep inside the duplexes formed by the blocking strands and the monomers, are strategically eliminated over multiple reaction steps to provide a ‘‘hidden’’ thermodynamic drive to the system (*43*) that ensures that polymerisation is thermodynamically favourable, but kinetically slow in the absence of a template. Third, the domains *st_i_* are secondary toeholds that are revealed after TMSD; they serve to accelerate HMSD. However, *st*’*_i_\** are truncated versions of the complement; an extra base pair in the initial monomer-blocker complex acts as a ‘‘clamp’’ to reduce unintended leak reactions (figure S2) (*44*).

To determine the domain lengths, we took heed of simulations of ODE models of trimerisation (*45*), parameterised using the the results of our earlier TMSD-HMSD systems (*33, 42*), which sought to identify regimes in which the mechanism of Figure 1B would allow a high yield of trimer products. We tested a few different designs around the optimum suggested by these simulations. For the systems reported, the recognition domains are 6-nucleotide (nt) long, except for the terminal monomers which have recognition domains of 8 nt. The *st* and *bm* domains are formed of 2 and 33 nt respectively. All sequences are given in Table S1.

### Binding of the blocked monomers to trimer templates

To test the viability of the design, we first checked the ability of the template to bind with the monomers by displacing the blocker strands. To monitor the reactions, we labelled the 3’-ends of the monomers with different fluorophores and the 5’-ends of the blocker strands with corresponding quencher species. When the template binds to the monomers, displacing the quencher-labelled blocker strands, fluorescence intensity increases.

To demonstrate the ability of the templates to direct the assembly of different monomers, we considered two variants for each of the sites on the template: *A* & *Ā* possessing only a right polymerisation interface, *B* & *D* possessing both, and *C_s_* & *E* possessing only a left polymerisation interface. The monomers within each of these pairs only differ by their recognition domains and fluorescence labels; the polymerisation interfaces are equivalent. The nomenclature reflects the relationship of the monomers to those we will subsequently use to demonstrate pentamer templating.

In separate experiments containing fixed concentrations (intended to be 10 nM) of either *ĀL*_1_ or *DL*_2_, we added varying concentrations of trimer template ^3^*T_ĀDE_*. Corresponding fluorescence signals (and inferred concentrations of reacted monomer as described in Methods) increased to levels proportional to the concentrations of added ^3^*T_ĀDE_* within 3 hours, as shown in Figure 2A and figure S9, indicating successful TMSD with mean estimated rate constants of 9.63 × 10^4^ ± 9.62 × 10^2^ M^−1^ s^−1^ for *Ā*, and 5.49 × 10^4^ ± 5.57 × 10^2^ M^−1^ s^−1^ for *D* (Table S28, S29). Similar results were obtained when different concentrations of ^3^*T_ABC_* were added to a fixed concentration of an orthogonal set of blocked monomers *AL*_1_ and *BL*_2_ (figure S9, table S26, S27).

**Figure 2:**
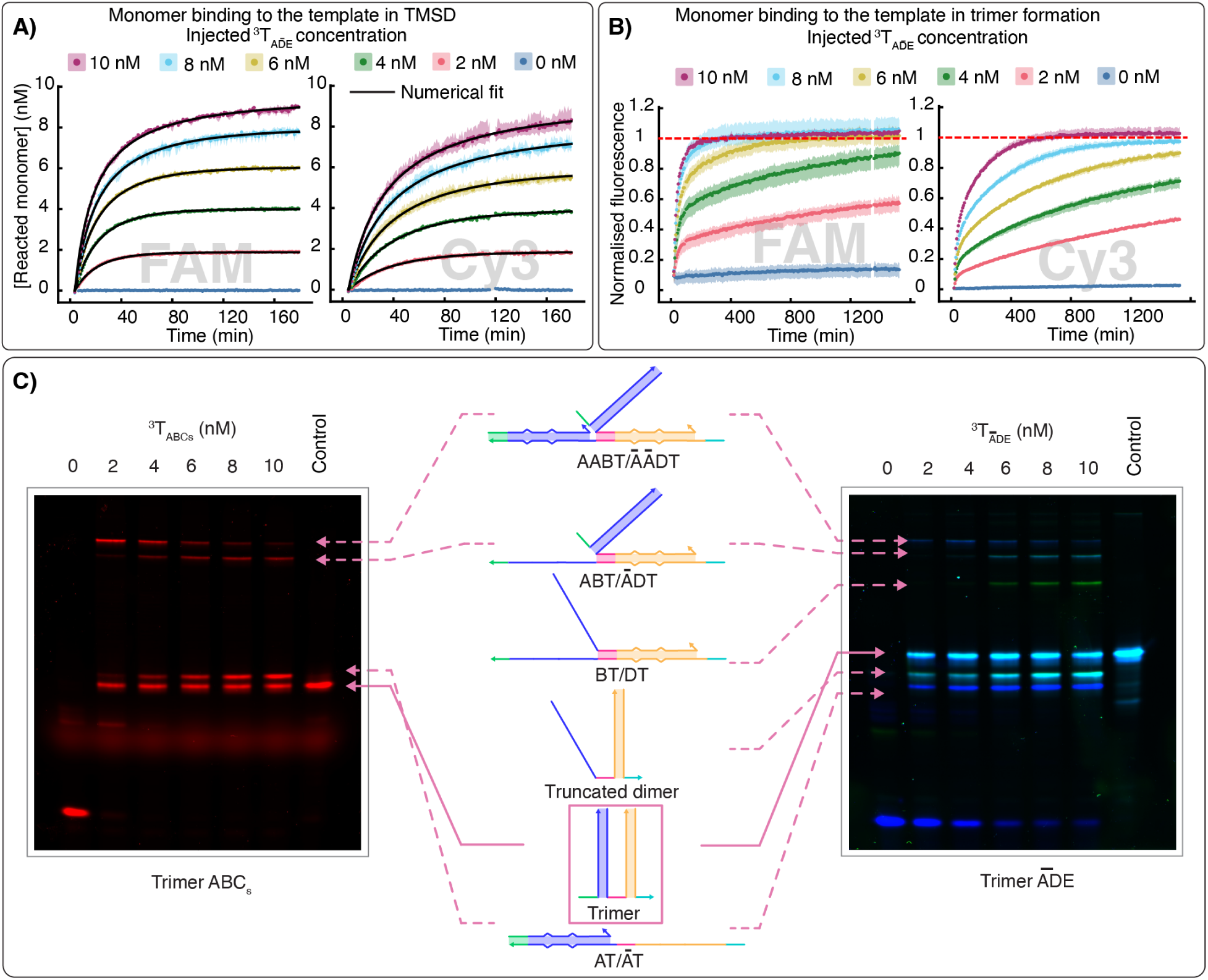
Trimer formation via templated catalysis. **A)** Individual monomer binding to the template ^3^*T_ĀDE_ via* TMSD. 2-10 nM of ^3^*T_ĀDE_* was added to 10 nM partially blocked monomers *ĀL*_1_ or *DL*_2_, with concentrations of reacted monomers inferred as described in methods. Lines are fits to a second order kinetic model. **B)** Fluorescence arising from monomer turnover during catalytic trimer formation, normalised relative to a 10 nM positive control (*ABC_s_* or *ĀDE*). 2-10 nM of ^3^*T_ĀDE_* was added to a mixture of 10 nM partially blocked monomers *ĀL*_1_, *DL*_2_, and free monomer *E*. The reported fluorescence signals correspond to reacted *Ā* and *D*. For both A) and B), data was recorded in triplicate, with the mean and min-max range shown. **C)** Fluorescence images of PAGE gels loaded with 10 pL aliquots from experimental wells of the experiments in B), along with pre-annealed products as positive controls for two trimer formation networks. Bands are identified as discussed in methods section.

### Complete catalytic cycle of trimerisation

We then tested the ability of the templates to catalyse the assembly of trimeric products from the available monomers. We prepared an equimolar mixture containing 10 nM blocked monomers *ĀL*_1_, *DL*_2_, and a free terminal monomer *E*, and added different concentrations (0 – 10 nM) of ^3^*T_ĀDE_* to the monomer mixtures. Fluorescence relative to a 10 nM positive control is reported in Figures 2B; later time points and equivalent data for ^3^*T_ABCs_* are given in figure S11. Unlike the TMSD reactions, which saturate at a level reflecting reaction stoichiometry, the concentration of reacted monomers continues to grow towards the positive control in all experiments, at a rate dependent on the amount of template added to the mixture. This behaviour is indicative of effective catalytic turnover, and hence limited product inhibition.

After ∼24 hours, 10 µL aliquots of the reaction mixtures were examined by polyacrylamide gel electrophoresis (PAGE), along with a positive control of 10 nM pre-annealed trimeric product (Figure 2C, raw data in figure S25, S26).

We observed strong product bands when the templates were present, and very low or no product formation in absence of the templates. From the strength of the band, the free product yield relative to control is estimated to be 50-60% for *ABC_s_* and 35-55% for *ĀDE* (Table S25). We observed apparent brush-like configurations with two template-bound partial products as slow-migrating bands at the top of the gels at higher monomer:template ratios. We also saw truncated dimers *DE* that were particularly prevalent at low monomer:template ratios. This behaviour is exactly as predicted by the mechanism in Figure 1B (*40*); when there are many monomers for every template, dense brushes can form and the production of truncated dimers is expected to be suppressed as the templates reach a brush-like steady state. When monomers do not outnumber templates, this dense brush-like state is not generally reached and truncated products are easier to form. Note that equivalent *BC_s_* band is not visible in the gel on left due to the choice of fluoropohores, and does not necessarily mean that they are not formed.

### Extended catalytic turnover of templates and product inhibition

We next tested the capacity of the templates to perform many catalytic cycles by exposing 100 nM of monomers to 0 – 5 nM of the corresponding trimer template. For all template concentrations, we observed 40% or more turnover of the monomers for both variants in 2 days (Figure 3A), rising to 80% or more after 9 days (figure S12). This successful turnover, and the formation of the intended products, is confirmed by PAGE experiments performed after 2 days from the template injection. With 5 nM concentrations of templates, all monomer bands at the bottom of the gel have disappeared due to their conversion into corresponding products and intermediates (Figure 3B, raw data in figure S27). Corresponding pixel density profiles of the gels can be used to estimate free product yield, which is 40% even with a 50 fold excess of monomers.

**Figure 3:**
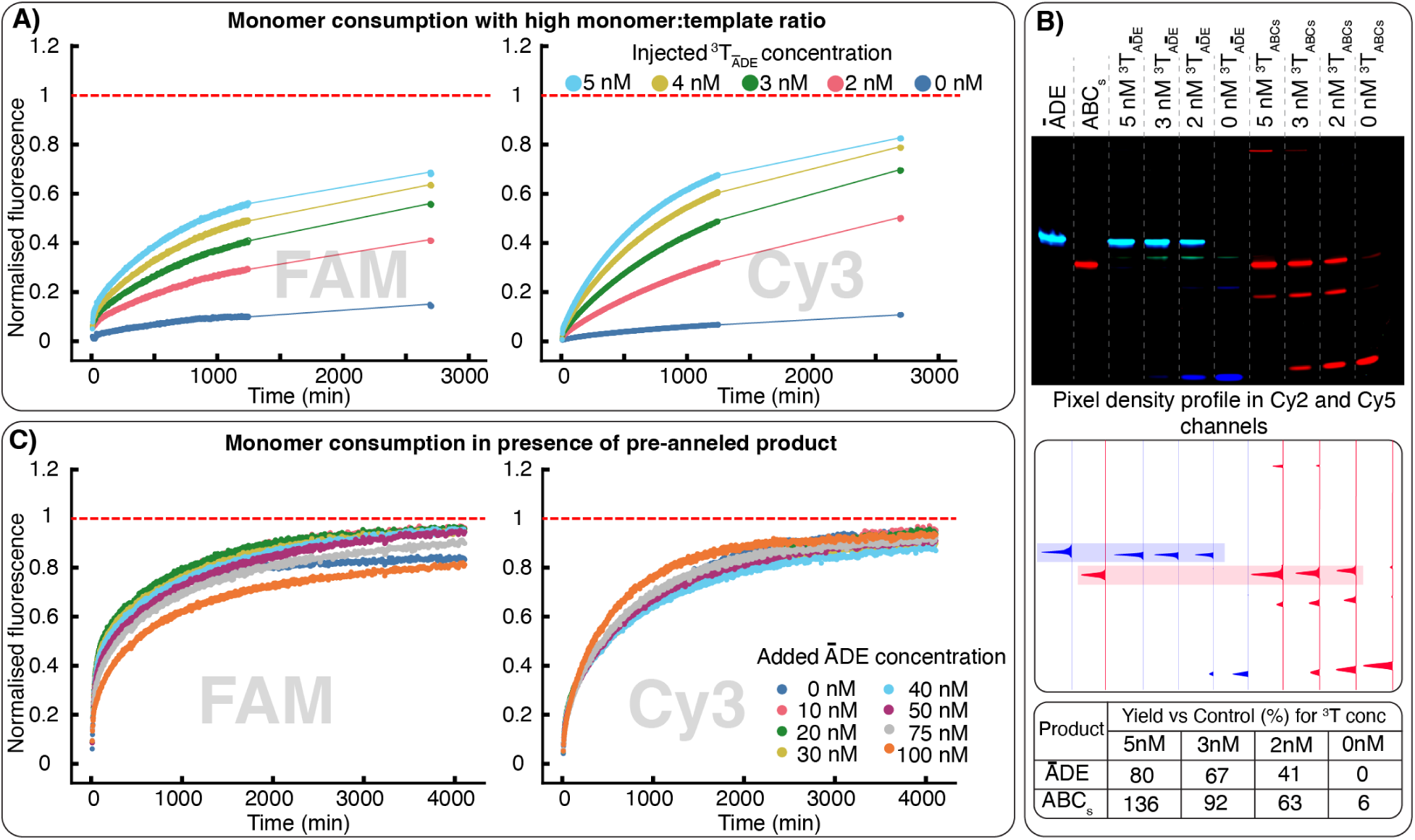
Effective catalytic trimerisation in the presence of large excesses of monomer and product. **A)** Fluorescence relative to 100 nM positive control (*ĀDE*) in the channels corresponding to unblocked *Ā* and *D* in response to addition of low (0 – 5 nM) concentrations of template ^3^*T_ĀDE_* to 100 nM each of *ĀL*_1_ and *DL*_2_ and *E*. **B)** PAGE image of 10 pL aliquots from the experiments in A), and equivalents for *ABC_s_* along with the pre-annealed controls. The pixel density plot of each lane of the gel is shown below. Shaded area in light blue shows the *ĀDE* peaks, and light red shows *ABC_s_*. Analysis of the same image provides estimated yields relative to positive control. **C)** The limited effect of product concentration on the template’s catalytic ability. A trimerisation reaction containing 10 nM of the monomers and 4 nM of ^3^*T_ĀDE_* was initiated with an additional 0 – 100 nM of pre-annealed, unlabelled *ĀDE*, with fluoresence reported relative to 10 nM of fluorescent positive control (*ĀDE*), in the FAM (corresponding to *Ā*) and Cy3 (corresponding to *D*) channels.

Separately, we checked the effect of pre-formed products in the reaction mixture, to explore the effect of product inhibition on catalytic turnover. We pre-annealed unlabelled *ĀDE* and added 10 –100 nM of this competitor product into different reaction wells containing 10 nM of monomers and 4 nM ^3^*T_ĀDE_*. The fluorescence signal of reaction kinetics showed little variation with excess product, suggesting very limited product inhibition (Figure 3C). Further experiments to test the propensity of the templates to bind to the fully formed products confirmed the instability of template-product complexes in a similar system with the same domain lengths (figure S37).

### Catalytic production of tetrameric and pentameric complexes

#### Differences from the trimer-forming network

To test the generalisability of the templating mechanism for making longer products, we attempted to make tetrameric *ABCD_s_*(^4^ *P*) and pentameric *ABCDE* (^5^ *P*) products (design and mechanism schematics in Figure 4A, figure S1 and S6). ^4^ *P* is formed by initial monomer *A*, two propagating monomers *B* and *C*, and a short terminating monomer *D_s_* with no right-hand polymerisation interface. *A* and *B* are the same monomers used in trimer formation, and *C* is an extended version of *C_s_*. Similarly, ^5^ *P* is formed by initial monomer *A*, three propagating monomers *B*, *C* and *D*, and the terminating monomer *E*. *A*, *B* and *C* are as above, and *D* and *E* are the same monomers used in the trimer formation experiments. Initially, *A*, *B*, *C* and *D* are present in their blocked form *AL*_1_, *BL*_2_, *CL*_1_, and *DL*_2_.

**Figure 4:**
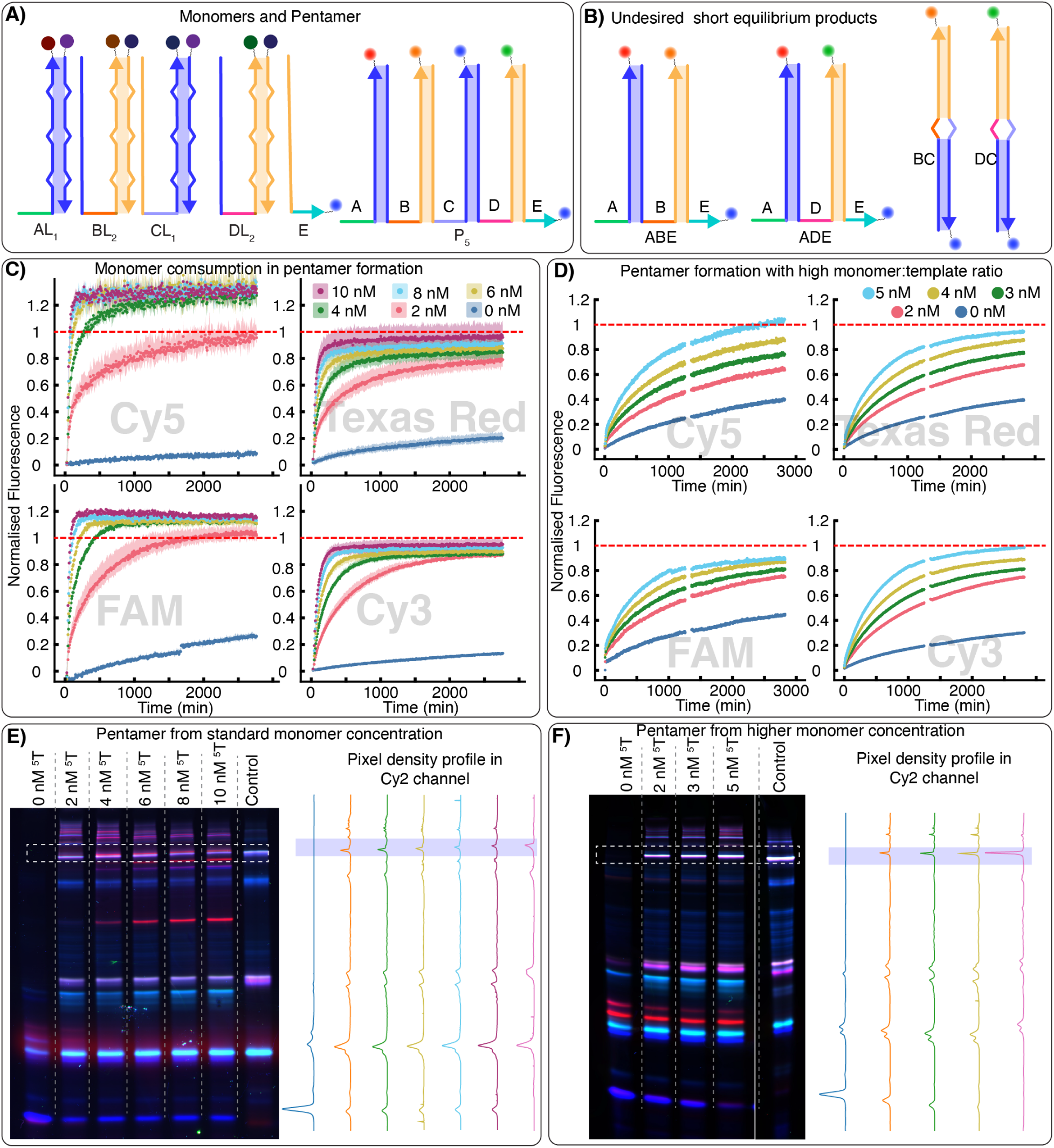
Catalytic formation of pentamers. **A)** domain-level diagram of intended pentameric products. **B)** domain-level illustration of thermodynamically stable dimeric and trimeric side-products. **C)** Fluorescence relative to 10 nM positive control (a 1:1:1 mixture of *AB*, *DE*, and *C*), arising from the unblocking of the monomers. Data shows average of triplicate with min-max range in the channels corresponding to unquenched *A*, *B*, *C*, and *D* in pentamerisation reactions containing 10 nM of all monomers and 0 – 10 nM of ^5^*T*. **D)** Fluorescence relative to 100 nM of same positive control for templated pentamer formation with a high concentration (100 nM) of monomers and 0 – 5 nM of ^5^*T*. **E-F**) Fluorescence PAGE images of 10 pL aliquots of the experiments in C) and D) with their corresponding pixel density profile. The product band/peak is shown in the boxed areas.

Multimer products are kept together by the long duplexes formed by alternating *bm*_1_ and *bm*_2_ domains and their complements. This repetitive design for the polymerisation interfaces ensures that a finite set of monomers can, in principle, be combined into an arbitrarily long product – as in transcription and translation. However, it also presents another challenge. In systems containing only *AL*_1_, *BL*_2_ and *C_s_*, or only *ĀL*_1_, *DL*_2_ and *E*, the intended trimer is the most thermodynamically favoured structure. Unless these sets of monomers are mixed, to create sequence variation that must be selected against in a non-equilibrium fashion, the equilibrium state is enriched in the intended product. In ^4^ *P*- and ^5^ *P*-forming systems, however, the longer products are metastable even without sequence variation. When multiple monomers with left and right polymerisation interfaces are present, the most thermodynamically favoured assemblies are a set of dimers and trimers as depicted in Figure 4B and figure S6C. These unwanted products saturate the polymerization domains while forming as many separate complexes as possible; they are therefore entropically favoured (*46*), as confirmed by NUPACK (*47*) analysis (figure S4, S5). A far-from-equilibrium templating mechanism is therefore essential if these longer products are to form, even without sequence specificity.

We first confirmed the binding of each monomer to the template ^5^*T via* TMSD (figure S10). All TMSD reactions were completed within four hours with estimated rate constants between 4.61 × 10^4^ ± 1.86 × 10^3^ M^−1^ s^−1^ and 1.21 × 10^5^ ± 8.1 × 10^2^ M^−1^ s^−1^ (Table S30 - S33). Next, we tested the ability of the right-hand (*n* +1) monomer to bind to the left-hand (*n*) monomer *via* HMSD and open up site *n* on the template for further TMSD. We exposed the template to large excesses of each monomer *n* in a sequential manner, and observed the TMSD reaction for each monomer saturate (figures S21, S22). Addition of monomer *n* + 1 then triggered further reactions of monomer *n*, consistent with successful HMSD. Secondary effects are observed when further monomers are added, as expected.

To template formation of ^4^ *P* and ^5^ *P*, we injected 0 – 10 nM of ^4^*T* and ^5^*T* into a mixture of the required monomers. Addition of the templates into the monomer mixture showed far faster increase of all fluorescence signals compared to the non-templated monomer mixtures (Figure 4C, figure S13). PAGE experiments showed clear product bands at the upper section of the gels, aligned with hierarchically assembled positive controls (Figures 4E, raw data in figure S28, S30) with a yield of up to ∼10% and 9% for ^4^ *P* and ^5^ *P* respectively, as estimated by comparing the band brightness to fluorescence in the positive controls (Table S25).

Higher molecular weight complexes, consistent with brushes (figure S7, S8) were also observed in the upper section of the gels. We also successfully tested a similar pentamer (^5^ *P_Ē_*) formation system in which the final monomer (*Ē*) and the template (^5^*T_Ē_*) are distinct (figure S15 and S32).

Multiple overlapping bands containing several labelled strands can be a source of uncertainty. Our procedure for identifying species, discussed in the methods, is based on migration distance and a binary classification of the presence or otherwise of the band in each fluorescent channel. To provide simpler results, we also performed experiments in which only the initial and terminal monomers were labelled, using two distinct fluorophores, with the results reported in figure S36. For both tetramers and pentamers the product band is clearly visible, and no product forms in the absence of a template. Slower migrating bands typically do not include the terminal monomer, as expected from brush-like conformations. There is some yield of truncated trimers containing both initial and terminal monomers in the pentamer case, but essentially no sign of the dimers *AD_s_* for the tetramer. A number of partially formed products containing the terminal monomer (but not the initial monomer) are visible, as is expected if the mechanism of (*40*) is only partially successful and lef-hand truncated products form.

The production of ^4^ *P* and ^5^ *P* reported in Figure 4E and figure S28 is catalytic as the products are not template-bound (see also figure S37); however, the yields are not sufficiently high to conclude that any given template is engaged in multiple rounds of successful catalysis. To verify that multiple rounds of turnover are possible, we added low concentrations of template (0 – 5 nM) to a reaction mixture containing 100 nM of each monomer (Figure 4D, figure S14). In all cases, fluorescence exceeded 50% of a positive control within a single day. Corroborative PAGE experiments (Figure 4F, figure S29) showed strong product bands and complete consumption of traceable monomers at 5 nM template concentration. Quantitative analysis of a gel run on a separate experiment (figure S41, S42, table S11, S14) with more template concentrations indicates up to 5x and 3x average catalytic turnover for 4T and 5T at the lowest template concentrations (S25), with negligible formation of full products without a template.

Increased monomer concentrations also increased the extent of undesired leak reactions in these experiments, where ∼40% of monomers *B* and *C*, and relatively lower amounts of other monomers, were consumed without any template. These leaks formed the short, thermodynamically stable by-products of Figure 4B; the templates are essential in directing assembly of long-lived metastable products. Importantly, however, the intended products do not readily disproportionate into smaller, thermodynamically favoured complexes once assembled (figure S17).

### Specificity of the templating processes

Having demonstrated catalytic formation of products of up to five units in length, we proceeded to test the specificity with which templates can selectively incorporate monomers from a pool of competing alternatives. For the trimer formation network, we used the two previously described sets of monomers for this purpose. For these sets, the pairs of strands ( *A, Ā*), (*B, D*) and (*C_s_, E*) are identical apart from their recognition interfaces, and thus any trimeric product could be formed by selecting one strand from each set. We initially tested the specificity of TMSD; in separate experiments for each blocked monomer (*AL*_1_, *ĀL*_1_, *BL*_2_ or *DL*_2_), we added either matching or non-matching templates and monitored the change in fluorescence. In all cases, matching templates reacted at a much faster rate with the monomer compared to the non-matching ones (figure S18). We then added the matching or non-matching templates to either the first (*AL*_1_, *BL*_2_, *C_s_*) or the second (*ĀL*_1_, *DL*_2_, *E*) set of monomers, monitoring fluorescence increases (S19, S20) and analysing the final products by PAGE (Figure 5B, raw images in figure S38).

**Figure 5:**
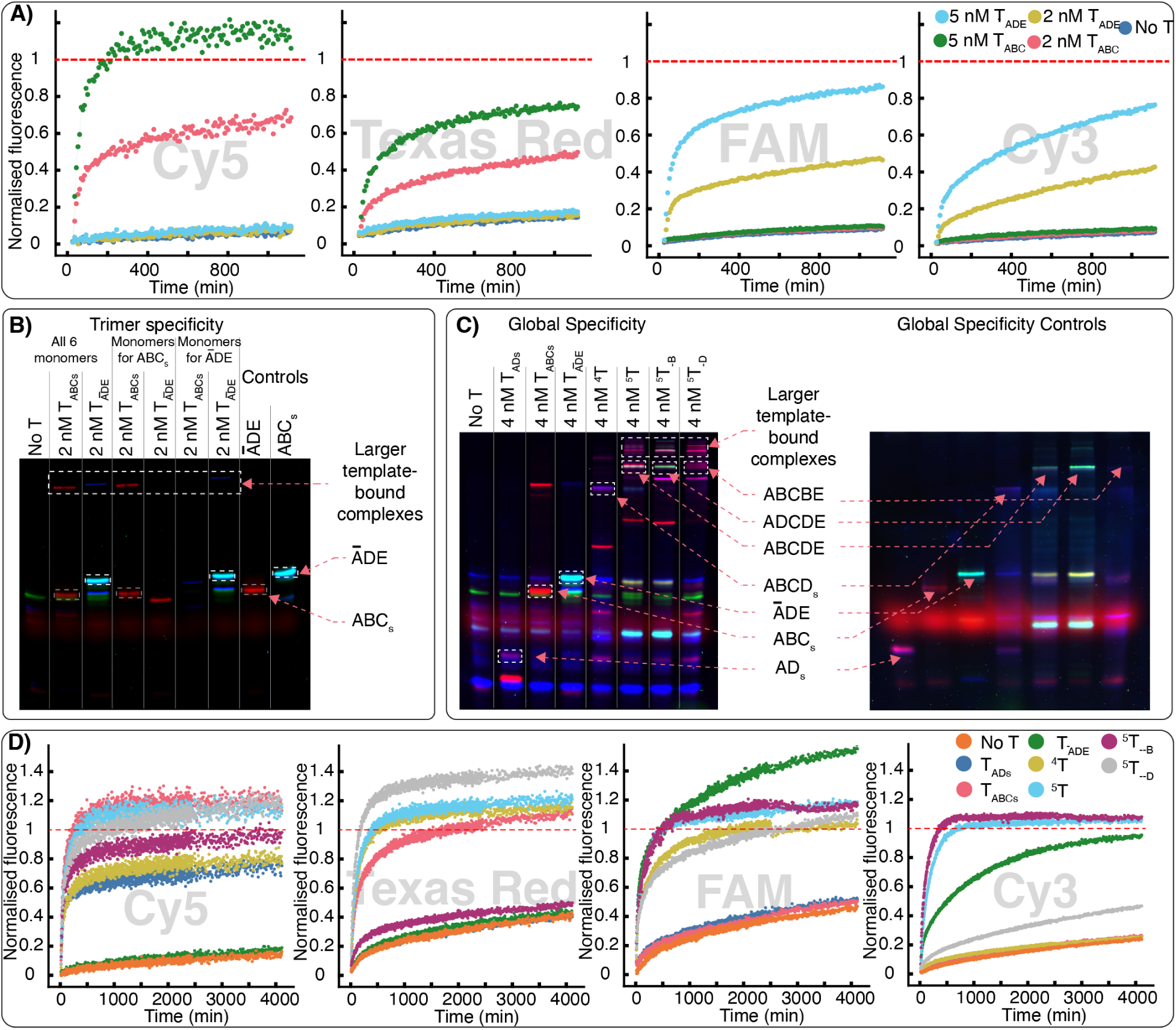
Sequence-specific templating. **A)** Fluorescence normalised relative to 10 nM positive controls (*ABC_s_* or *ĀDE*) in the channels of unquenched *A*, *B*, *Ā*, and *D* (from left to right) when mixtures of 10 nM each of 6 monomers (*AL*_1_, *ĀL*_1_, *BL*_2_, *DL*_2_, *C_s_*, and *E*) were triggered with ^3^*T_ABCs_*, ^3^*T_ĀDE_*, or nothing. Fluorescence PAGE images showing **B)** specificity of the trimeric templates alongside pre-formed products, and (**C)**) global specificity with pre-formed products (left) and controls (right). Bands are identified as discussed in materials and methods section. **D)** Fluorescence normalised relative to 10 nM positive controls in the channels of unquenched *A*, *B*, *Ā* + *C*, and *D* (left to right) after adding 4 nM of 7 different templates (see legends) to separate experimental wells containing a mixture of 8 monomers (*AL*_1_, *ĀL*_1_, *BL*_2_, *CL*_1_, *DL*_2_, *C_s_*, *D_s_*, *E*), each at 10 nM.

In both cases, the trimeric products formed efficiently when the right template was added to the system, with little-to-no product in presence of the wrong template. This behaviour was also observed when all six monomers were mixed in the reaction and triggered with either template (Figure 5A, B and figure S38)). In these PAGE tests, conducted after ∼20 hours from template injection, some undesired leak reactions are visible – particularly the apparent formation of *AT_ĀDE_* in lane 5, consistent with the fact that *T_ĀDE_* can slowly react with *A* when the template is not occupied with other monomers (figure S19). However, we see no evidence of the full assembly of either trimeric product in the absence of the matchnig template.

As a final test for the global specificity of the templating process, we mixed 8 monomers (*AL*_1_, *ĀL*_1_, *BL*_2_, *CL*_1_, *DL*_2_, *E*, *C_s_*, and *D_s_*) together and injected into separate wells one dimer template (for *AD_s_*), two trimer templates (for *ABC_s_*, and *ĀDE*), one tetramer template (for *ABCD_s_*), and three pentamer templates (for *ABCDE*, *ABCBE*, and *ADCDE*). PAGE gels (Figure 5C, figure S39) confirm that the intended product forms in each case, but not if the specific template is absent. Although crosstalk and leak reactions are exacerbated when all monomers are present in the same reaction, kinetic traces in Figure 5D clearly show that monomers still react far faster when a complementary template is present.

### Discussion and Outlook

DNA has been widely used to assemble complex structures, including crystals (*48, 49*), finite-size objects (*50,51*), and algorithmic assemblies (*52,53*). However, these assemblies are designed to out-compete alternatives due to stronger base pairing, either throughout the structure (*48–51,53*) or due to interactions with a nucleating seed (*52*). The intended products are thus favoured in equilibrium. Assembly by catalytic templating is qualitatively different in nature, since the use of catalytic templates to influence experimental outcome is an inherently far-from-equilibrium phenomenon. In this paper we have used information-bearing catalytic templates to selectively assemble long-lived, metastable multimers containing up to five monomers. Our catalytic mechanism overcomes the central challenge of templated assembly – product inhibition – allowing our system to operate enzyme-free under constant external conditions, even for assemblies containing five units.

Recently, interest has grown in demonstrating non-equilibrium nanotechnological systems that are driven out of their equilibrium state by the continuous consumption of molecular fuel, often facilitated by enzymatic action (*54, 55*). As with templating, these works are heavily inspired by natural systems; in this case, enzyme cascades (*56*), kinetic proofreading (*33*), and microtubule growth (*57*). The results presented here do not use such a motif – products are intended to assemble and then persist in a metastable non-equilibrium state, rather than being driven round a cycle. An intriguing extension would be to develop networks in which products are driven round cycles of templating and destruction, as RNA and proteins are in cells. Here, the design of our system holds some promise: one possibility would be to degrade longer products by catalysing their conversion into the more stable dimers and trimers of Figure 4B.

The system as it stands is a proof of principle that enzyme-free, sequence-specific catalytic templating of multimers is possible. Although the mechanism is a lot less efficient than natural templating systems, its advantage is that the objects templated do not have to conform to the requirements of a polymerase or ribosome. A natural extension, would, therefore, be to use our templating mechanism to direct the co-localisation of functional moieties beyond the fluorophores attached to the strands here – for example, functional groups that drive phase separation (*58, 59*) or perform cooperative molecular recognition (*60*). Sequence-specific templating, as in nature, would allow a relatively small library of functionalised monomers to be brought together in a combinatorially large number of ways, allowing the efficient exploration of an enormous design space.

While our experiments show side reactions that lead to undesired products, the results – particularly the apparent brush-like configurations observed in PAGE (Figure 4E, F), and the behaviour of the system when monomers are omitted (figure S24) or only added later on (figure S23) – are consistent with the correct products being produced exclusively by catalytic templating, with at least a partially successful implementation of the specific mechanism proposed in Ref. (*40*). Nonetheless, potential applications likely require improved accuracy and speed of templating. In particular, it is not trivial to design the template to avoid folding onto itself, which we suspect is a potential kinetic bottleneck. Stretching the ssDNA using, for example, an origami structure may be beneficial in that regard, and it would also limit the tendency of monomers to interact with other monomers that are not their neighbours. We also expect that future applications may exploit the *principle* demonstrated here, while using a modified detailed mechanism. One particularly important question is whether a similar principle can be used to demonstrate sequence-specific catalytic templating in which monomers form a covalent bond, rather than one based on non-covalent base pairing.

## Author contributions

Conceptualization: RM and TEO; Investigation and Methodology: RM and MM; Formal analysis, Data curation, and Visualization: RM, JJ, KJ, and TEO; Software: KJ; Supervision: RM, and TEO; Funding acquisition: TEO; Writing– original draft: All authors.

## Data and materials availability

All original fluorescence data for reaction kinetics, and original fluorescence images are freely available in Zenodo at the following URL: https://doi.org/10.5281/zenodo.20822987

Source code and user instructions for the Clearissa version 1.71, which was used to analyse the kinetic fluorescence data, is available at the following URL: https://github.com/PrinciplesKJ/ Clearissa_1.71

## Supplementary materials

Methods

Figures S1 to S42

Tables S1 to S33

References (*61–63*)

## Methods

### Materials

All the DNA strands were purchased from Integrated DNA Technologies as 100 µM solutions in pH8 IDTE buffer. NaCl and lyophilised bovine serum albumin (BSA) were purchased from Sigma. The flat-bottomed 96-well plates were obtained from Greiner Bio-One. Precast polyacrylamide gels and 10x Tris-Borate-EDTA (TBE) buffer were purchased from Invitrogen. Gel loading dyes were obtained from New England Biolabs.

### Details of strand designs

The strand design process started with a hand-drawn scheme and mechanism of reactions. Then we assigned specific domains to each strand where each domain is responsible for some specific task. We assigned the domain length based on previous studies on TMSD (*61*) and our own work on HMSD (*33–35, 42, 45*). These complex structures and design parameters are then fed into the NUPACK strand design programme to obtain sequence-level designs of the required strands (*62*). The design process is particularly challenging because Nupack bases strand designs on equilibria, whereas our system is intended to remain out of equilibrium even at the end of the process. Moreover, the nature of templating is that two monomers need to bind to both each other and the template. This prohibits the design of strands that are either predominantly ATC or ATG to minimise secondary structure, a common strategy in DNA nanotechnology. It was thus necessary to proceed through several iterations of trial and error to obtain sequences that functioned well, we present those sequences here. However, The rates of the strand displacement reactions, particularly of HMSD are also sensitive to sequences and secondary structures. We tested, and tried to incrementally optimise our design with several iterations. But this method is taxing on resources and time. A more robust way of generating sequence-level designs for different reaction topologies with predictable rate constants would help to further enhance the speed and yield of the systems.

### Preparation of complexes

Solutions of ∼2 pM ssDNA strands were prepared by diluting the purchased strands with pH7 IDTE buffers and concentrations were re-estimated by measuring the absorbance at 260 nm using Implen N50 nanophotometer. The blocked monomers were prepared by first mixing 200 nM of a monomer with 300 nM of its corresponding blocker strand and 1X TAE buffer containing 1 M NaCl to yield a 100 μL solution.The mixture was then heated to 95^◦^C and held there for 5 minutes before gradually cooling it down by 1^◦^C every 2 minutes to 20^◦^C, and was kept at that temperature until used for experiments. Other complexes used as positive controls were prepared by the same method, but varying the concentrations of the components as required. Complexes older than a week were not used in experiments.

### General method for strand displacement experiments

DNA strand displacement experiments were performed using fluorescently labelled strands and the progress of the reactions was monitored by BMG Labtech ClarioSTAR plate reader. The gain and filter settings used for the corresponding fluorophores at standard and high concentrations are listed in the Table S3. All the experiments, unless otherwise stated, were conducted at 25 ^◦^C in 200 µL reaction volume in Greiner pClear flat-bottom 96-well plates.

### Gel electrophoresis and fluorescence imaging

Native Polyacrylamide Gel Electrophoresis (PAGE) was performed to investigate the identities of the formed complexes. The samples were prepared by mixing 10 µL aliquot of the reaction mixtures or 10 nM pre-annealed positive controls with 2 µL 6x purple gel loading dye. The entire samples were loaded in the 10% TBE gel pockets. The electrophoresis was performed at a constant 90 V for 120 or 180 minutes depending on the samples, using 1x TBE as running buffer. After the electrophoresis was completed, the plastic cassettes of the gels were removed, and the gels were placed in 1X TBE solution in a staining box on a rocker for 20 minutes to remove the loading dyes. Then the gels were removed from the TBE solution and placed directly on the imaging stage of Amersham Typhoon laser scanning platform to record multi-channel fluorescence images of the gels. Details of the channels used are given in Table S4.

### Protocol for the preparation of fluorescence gel images and their analysis

We used **Fiji** software (*63*) to prepare gel images and analyse the results. Composite images are prepared to optimise visual clarity in each case, and scaling of colours is not uniform between images. Both the identification of bands and the calculation of yields were performed using the separate images of each channel, rather than with these processed composites. The details of the image preparation workflow are provided in the Zenodo upload https://doi.org/10.5281/zenodo. 20822987.

### Inferring the identity of the bands

Wherever possible, the samples were run alongside pre-made controls. The gels were then scanned in up to three fluorescence channels. The migration distances of the bands and the presence or absence of signals from particular channels corresponding to the fluorescence labels of constituent strands (listed in Table S1) were used to propose the band identity. For a representative example, *ABC_s_* should appear only Cy5 channel (fluorescence label of *A*), and not in anything other (figure S27). *ĀDE*, on the other hand, should appear in Cy2 (*Ā*) and Cy3 (*D*) channels, and not in Cy5 (figure S27). Tetrameric product ^4^ *P* migrates more slowly than the trimers, and contains Cy5 (*A*), and Cy2 (*C* and *D_s_*), and no Cy3 (figure S28). Pentameric ^5^ *P* moves even slower and is visible in all 3 channels (figure S30). A comprehensive table list of complexes of interest in their corresponding gels and the expected signals from the fluorescence channels is given in Table S24.

### Estimating the yields of reactions

For the purposes of calculating yields, electrophoreses was performed with aliquots of the reaction mixtures alongside known concentrations of the pre-formed products, which were used as the references. First, the backgrounds were subtracted from 16 bit images in each channel to obtain sharp bands. Then the reference band (for trimers) or the entire positive control lane (for the products that cannot be synthesised by directly annealing the strands, tetramer and pentamer) was selected to calculate the integrated pixel density, which was treated as the reference value for 100% yield. Then the integrated pixel densities were also calculated for the concerned bands from the other lanes, and were compared against the reference values to estimate the yields of the reactions. When appropriate, this procedure was repeated separately for all chanels in which the product was expected to be visible.

### Generating the pixel density plots

Pixel density plots can be a qualitative way to compare the yields and contents of several lanes of a gel image. Images obtained by scanning the gels in different channels were analysed independently to generate the plots The profiles were generated by first subtracting the background, and then plotting the pixel brightness along a straight line path drawn from the top to the bottom of each lane of interest of a gel image. Where complexes had multiple channels, the clearest channel with the fewest peaks was shown in the main text; plots for the other channels are provided in the Zenodo upload https://doi.org/10.5281/zenodo.20822987. The step-by-step methods for generating the plots from raw data are also provided there. Composite images were not used because different scaling of the individual channels can lead to erroneous results.

### Experimental protocol for monitoring binding of monomers to the templates

This protocol was used to gather data reported in Figures 2a, S9 and S10.

#### Step 1 for baseline

20 pL of 20% Tween20 in 1X TAE and 1 M NaCl, and 120-130 pL of experimental buffer was added to each experiment and control well. Then 10 μL of pre-annealed blocked monomer complexes (200 nM stock) was added to the buffer in the experiment wells to get a final concentration of 10 nM (calculated for the total reaction volume of 200 μL, which is attained after Step 2). For the positive controls, 3-10 μL of pre-annealed monomer-template complex (200 nM stock concentration) was added to get final concentration of 3-10 nM. For the blank, nothing was added except Tween20 and experimental buffer. All the wells had equal volume of 150 µL at this point. The fluorescence values were measured over at least 10 data points. Reactions were performed in triplicate simultaneously.

#### Step 2 for reaction kinetics

Different volumes (0-20 μL) of template strands (100 nM stocks) and 50-40 μL experimental buffer were injected into the experimental wells via the injector of the plate reader to make the final volume in each well 200 μL. The total injected volume of template and buffer was 50 µL for each experimental well. Template injection volumes were varied to attain a final *T* concentration of 0-10 nM in the wells. In the positive control and blank well, 50 pL experimental buffer was injected. The initial kinetics of template binding was followed by monitoring the change in fluorescence intensities for several minutes to several hours depending on the reaction rates.

#### Step 3 for unblocking all available monomers (saturation)

When the kinetic traces reached a plateau, an excess of the template (10 pL of 2 pM stocks) was added to the reaction wells and allowed to react for at least 3 hours before measurement. For the control wells, 10 μL of experimental buffer was added to keep the volumes of all wells equal. The fluorescence values were measured over at least 10 data points.

The required volumes for this method are tabulated in Table S5.

### Experimental protocol for trimer formation at standard monomer concentration

This protocol was used to gather data reported in Figures 2b, and S11.

#### Step 1 for baseline

20 pL of 20% Tween20 in 1X TAE and 1 M NaCl, and 100-130 pL of experimental buffer were added to each experiment and control well. Then 10 μL each of pre-annealed, blocked, first (*AL*_1_, *ĀL*_1_) and second (*BL*_2_*, DL*_2_) monomer complexes (200 nM stock) and free final monomer (*C_S_*, *E*, 200 nM stock) were added to the buffer in the experiment wells to obtain a final concentration of 10 nM of each component (calculated for the total reaction volume of 200 μL, which is attained after Step 2). For the positive controls, 3-10 μL of pre-annealed trimer complex (200 nM stock concentration) was added to get final concentration of 3-10 nM. For the blank, nothing was added except Tween20 and experimental buffer. All the wells had equal volume of 150 µL at this point. The fluorescence values of all required channels were simultaneously measured over at least 10 data points of step 1. Reactions were performed in triplicate simultaneously.

#### Step 2 for reaction kinetics

Different volumes (0-20 μL) of template strands (100 nM stocks) and 50-40 μL experimental buffer were injected into the experimental wells via the injector of the plate reader to make the final volume in each well 200 μL. The total injected volume of template and buffer was 50 μL for each experimental well. Template injection volumes were varied to attain a final T concentration of 0-10 nM in the wells. In the positive control and blank well, 50 pL experimental buffer was injected. The initial kinetics of template binding was followed by monitoring the change in fluorescence intensities for several minutes to several hours depending on the reaction rates.

The required volumes for this method are tabulated in Table S6.

### Experimental protocol for trimer formation at higher monomer concentrations

This protocol was used to gather data reported in Figures 3a, and S12.

#### Step 1 for baseline

20 pL of 20% Tween20 in 1X TAE and 1 M NaCl, and 70-130 pL of experimental buffer was added to each experiment and control well. Then 20 μL each of pre-annealed, blocked, first (*AL*_1_, *ĀL*_1_) and second (*BL*_2_*, DL*_2_) monomer complexes (1 pMstock) and free final monomer (*C_S_*, *E*, 1 pMstock) were added to the buffer in the experiment wells to obtain a final concentration of 100 nM of each component (calculated for the total reaction volume of 200 μL, which is attained after Step 2). For the positive controls, 5-20 μL of pre-annealed trimer complex (1 pMstock concentration) was added to get final concentration of 25-100 nM. For the blank, nothing was added except Tween20 and experimental buffer. All the wells had equal volume of 150 µL at this point. The fluorescence values of each required channel were simultaneously measured over at least 10 data points at the end of step 1.

#### Step 2 for reaction kinetics

Different volumes (0-10 μL) of template strands (100 nM stocks) and 50-40 μL experimental buffer were injected into the experimental wells via the injector of the plate reader to make the final volume in each well 200 μL. The total injected volume of template and buffer was 50 μL for each experimental well. Template injection volumes were varied to attain a final T concentration of 0-5 nM in the wells. In the positive control and blank well, 50 pL experimental buffer was injected. The initial kinetics of template binding was followed by monitoring the change in fluorescence intensities for several minutes to several hours depending on the reaction rates.

The required volumes for this method are tabulated in Table S7.

### Experimental protocol for tetramer formation at standard monomer concentrations

This protocol was used to gather data reported in figure S13.

#### Step 1 for baseline

20 pL of 20% Tween20 in 1X TAE and 1 M NaCl, and 90-130 pL of experimental buffer was added to each experiment and control well. Then 10 μL each of pre-annealed blocked monomer complexes *AL*_1_, *BL*_2_*, CL*_1_ (200 nM stock) and free final monomer *D_S_* (200 nM stock) was added to the buffer in the experiment wells to obtain a final concentration of 10 nM of each component (calculated for the total reaction volume of 200 μL, which is attained after Step 2). For the positive controls, 3-10 μL of pre-annealed complexes *AB* and *CD_s_*(200 nM stocks) was added to get final concentration of 3-10 nM tetramer. For the blank, nothing was added except Tween20 and experimental buffer. All the wells had equal volume of 150 µL at this point. The fluorescence values of each required channel were simultaneously measured over at least 10 data points at the end of step 1. Reactions were performed in triplicate simultaneously.

#### Step 2 for reaction kinetics

Different volumes (0-20 μL) of ^4^*T* (100 nM stocks) and 50-40 μL experimental buffer were injected into the experimental wells via the injector of the plate reader to make the final volume in each well 200 μL. The total injected volume of template and buffer was 50 μL for each experimental well. Template injection volumes were varied to attain a final T concentration of 0-10 nM in the wells. In the positive control and blank well, 50 μL experimental buffer was injected. All the wells had equal volume of 150 µL at this point. The initial kinetics of template binding was followed by monitoring the change in fluorescence intensities for several minutes to several hours depending on the reaction rates.

The required volumes for this method are tabulated in Table S9.

### Experimental protocol for tetramer formation at high monomer concentrations

This protocol was used to gather data reported in figure S14.

#### Step 1 for baseline

20 pL of 20% Tween20 in 1X TAE and 1 M NaCl, and 90-130 pL of experimental buffer were added to each experiment and control well. Then 20 μL each of pre-annealed blocked monomer complexes *AL*_1_, *BL*_2_*, CL*_1_ (1 pMstock) and free final monomer *D_S_* (1 pMstock) was added to the buffer in the experiment wells to obtain a final concentration of 100 nM of each component (calculated for the total reaction volume of 200 μL, which is attained after Step 2). For the positive controls, 5-20 μL of pre-annealed complexes *AB* and *CD_s_* (200 nM stocks) was added to get final concentration of 25-100 nM tetramer. For the blank, nothing was added except Tween20 and experimental buffer. All the wells had equal volume of 150 µL at this point. The fluorescence values of each required channel were simultaneously measured over at least 10 data points at the end of step 1.

#### Step 2 for reaction kinetics

Different volumes (0-10 μL) of ^4^*T* (100 nM stocks) and 50-40 μL experimental buffer were injected into the experimental wells via the injector of the plate reader to make the final volume in each well 200 μL. The total injected volume of template and buffer was 50 μL for each experimental well. Template injection volumes were varied to attain a final T concentration of 0-5 nM in the wells. In the positive control and blank well, 50 μL experimental buffer was injected. The initial kinetics of template binding was followed by monitoring the change in fluorescence intensities for several minutes to several hours depending on the reaction rates.

The required volumes for this method are tabulated in Table S10.

### Experimental protocol for tetramer formation at high monomer concentrations with 0–10 nM template

This protocol was used to gather data reported in figure S41.

#### Step 1 for baseline

20 pL of 20% Tween20 in 1X TAE and 1 M NaCl, and 90-130 pL of experimental buffer were added to each experiment and control well. Then 20 µL each of pre-annealed blocked monomer complexes *AL*_1_, *BL*_2_*, CL*_1_ (1 pMstock) and free final monomer *D_S_* (1 pMstock) was added to the buffer in the experiment wells to obtain a final concentration of 100 nM of each component (calculated for the total reaction volume of 200 µL, which is attained after Step 2). For the positive controls, 5-20 µL of pre-annealed complexes *AB* and *CD_s_* (200 nM stocks) was added to get final concentration of 25-100 nM tetramer. For the blank, nothing was added except Tween20 and experimental buffer. All the wells had equal volume of 150 µL at this point. The fluorescence values of each required channel were simultaneously measured over at least 10 data points at the end of step 1.

#### Step 2 for reaction kinetics

Different volumes (0–20 µL) of ^4^*T* (100 nM stocks) and 30–50 µL experimental buffer were injected into the experimental wells via the injector of the plate reader to make the final volume in each well 200 µL. The total injected volume of template and buffer was 50 µL for each experimental well. Template injection volumes were varied to attain a final T concentration of 0-5 nM in the wells. In the positive control and blank well, 50 µL experimental buffer was injected. The initial kinetics of template binding was followed by monitoring the change in fluorescence intensities for several minutes to several hours depending on the reaction rates.

The required volumes for this method are tabulated in Table S11.

### Experimental protocol for pentamer formation with standard monomer concentrations

This protocol was used to gather data reported in Figures 4C, S15.

#### Step 1 for baseline

20 pL of 20% Tween20 in 1X TAE and 1 M NaCl, and 80-130 pL of experimental buffer were added to each experiment and control well. Then 10 μL of pre-annealed blocked monomer complexes *AL*_1_, *BL*_2_*, CL*_1_*, DL*_2_, (200 nM stock) and free final monomer *E* (200 nM stock) were added to the buffer in the experiment wells to obtain a final concentration of 10 nM of each component (calculated for the total reaction volume of 200 μL, which is attained after Step 2). For the positive controls, 3-10 μL of pre-annealed complexes *AB* and *DE* (200 nM stocks) was added to the wells first, and the plate was shaken at 600 rpm on a double orbital shaker for 30 seconds. Then 3-10 μL *C* (200 nM stock) was added to the positive controls to get final concentration of 3-10 nM of each strand in the wells. A fraction of *C* therefore simultaneously binds to both *AB* and *DE* to make the pentamer. This system is then used as a control signal for the fluorescence monitoring, and also as the size marker in PAGE experiments. For the blank, nothing was added except Tween20 and experimental buffer. All the wells had equal volume of 150 µL at this point. The fluorescence values of each required channel were simultaneously measured over at least 10 data points at the end of step 1. Reactions were performed in triplicate simultaneously.

#### Step 2 for reaction kinetics

Different volumes (0-20 μL) of ^5^*T* (100 nM stocks) and 50-40 μL experimental buffer were injected into the experimental wells via the injector of the plate reader to make the final volume in each well 200 μL. The total injected volume of template and buffer was 50 μL for each experimental well. Template injection volumes were varied to attain a final ^5^*T* concentration of 0-10 nM in the wells. In the positive control and blank well, 50 μL experimental buffer was injected. The initial kinetics of template binding were followed by monitoring the change in fluorescence intensities for several minutes to several hours depending on the reaction rates.

The required volumes for this method are tabulated in Table S12.

### Experimental protocol for pentamer formation with high monomer concentrations

This protocol was used to gather data reported in Figures 4B, S16.

#### Step 1 for baseline

20 pL of 20% Tween20 in 1X TAE and 1 M NaCl, and 30-130 pL of experimental buffer were added to each experiment and control well. Then 20 μL each of pre-annealed blocked monomer complexes *AL*_1_, *BL*_2_*, CL*_1_*, DL*_2_ (1 pM stock) and free final monomer *E* (1 pM stock) was added to the buffer in the experiment wells to obtain a final concentration of 100 nM of each component (calculated for the total reaction volume of 200 μL, which is attained after Step 2). For the positive controls, 5-20 μL of pre-annealed complexes *AB* and *DE* (1 pM stocks) was added to the wells first, and the plate was shaken at 600 rpm on a double orbital shaker for 30 seconds. Then 5-20 μL *C* (200 nM stock) was added to the positive controls to get final concentration of 25-100 nM of each strand in the wells. A fraction of *C* therefore simultaneously binds to both *AB* and *DE* to make the pentamer. This system is then used as a control signal for the fluorescence monitoring, and also as the size marker in PAGE experiments. For the blank, nothing was added except Tween20 and experimental buffer. All the wells had equal volume of 150 µL at this point. The fluorescence values of each required channel were simultaneously measured over at least 10 data points at the end of step 1.

#### Step 2 for reaction kinetics

Different volumes (0-10 μL) of ^5^*T* (100 nM stocks) and 50-40 μL experimental buffer were injected into the experimental wells via the injector of the plate reader to make the final volume in each well 200 μL. The total injected volume of template and buffer was 50 μL for each experimental well. Template injection volumes were varied to attain a final ^5^*T* concentration of 0-5 nM in the wells. In the positive control and blank well, 50 μL experimental buffer was injected. The initial kinetics of template binding were followed by monitoring the change in fluorescence intensities for several minutes to several hours depending on the reaction rates.

The required volumes for this method are tabulated in Table S13.

### Experimental protocol for pentamer formation with high monomer concentrations and 0–10 nM template

This protocol was used to gather data reported in figure S42.

#### Step 1 for baseline

20 pL of 20% Tween20 in 1X TAE and 1 M NaCl, and 30-130 pL of experimental buffer were added to each experiment and control well. Then 20 µL each of pre-annealed blocked monomer complexes *AL*_1_, *BL*_2_*, CL*_1_*, DL*_2_ (1 pM stock) and free final monomer *E* (1 µM stock) was added to the buffer in the experiment wells to obtain a final concentration of 100 nM of each component (calculated for the total reaction volume of 200 µL, which is attained after Step 2). For the positive controls, 5-20 µL of pre-annealed complexes *AB* and *DE* (1 µM stocks) was added to the wells first, and the plate was shaken at 600 rpm on a double orbital shaker for 30 seconds. Then 5-20 µL *C* (200 nM stock) was added to the positive controls to get final concentration of 25-100 nM of each strand in the wells. A fraction of *C* therefore simultaneously binds to both *AB* and *DE* to make the pentamer. This system is then used as a control signal for the fluorescence monitoring, and also as the size marker in PAGE experiments. For the blank, nothing was added except Tween20 and experimental buffer. All the wells had equal volume of 150 µL at this point. The fluorescence values of each required channel were simultaneously measured over at least 10 data points at the end of step 1.

#### Step 2 for reaction kinetics

Different volumes (0-20 µL) of ^5^*T* (100 nM stocks) and 30–50 µL experimental buffer were injected into the experimental wells via the injector of the plate reader to make the final volume in each well 200 µL. The total injected volume of template and buffer was 50 µL for each experimental well. Template injection volumes were varied to attain a final ^5^*T* concentration of 0-5 nM in the wells. In the positive control and blank well, 50 µL experimental buffer was injected. The initial kinetics of template binding were followed by monitoring the change in fluorescence intensities for several minutes to several hours depending on the reaction rates.

The required volumes for this method are tabulated in Table S14.

### Experimental protocol for testing product inhibition in trimer formation

This protocol was used to gather data reported in Figure 3C.

#### Step 1 for baseline

20 pL of 20% Tween20 in 1X TAE and 1 M NaCl, and 0-130 pL of experimental buffer were added to each experiment and control well. Then 10 μL each of pre-annealed blocked monomer complexes *ĀL*_1_, *DL*_2_, (200 nM stock) and free final monomer *E* (200 nM stock) were added to the buffer in the experiment wells to obtain a final concentration of 10 nM of each component (calculated for the total reaction volume of 200 μL, which is attained after Step 2). Next, 0 - 100 μL of pre-annealed unlabelled *ĀDE* was added to the wells to vary its concentration between 0 and 100 nM. For the positive controls, 3-10 μL of pre-annealed labelled *ĀDE* (200 nM stock concentration) was added to get final concentration of 3-10 nM. For the blank, nothing was added except Tween20 and experimental buffer. The fluorescence values of all required channels were simultaneously measured over at least 10 data points of step 1.

#### Step 2 for reaction kinetics

8 pL of template strands (100 nM stocks) and 42 pL experimental buffer were injected into the experimental wells via the injector of the plate reader to make the final volume in each well 200 μL. The total injected volume of template and buffer was 50 μL for each experimental well. Template injection volumes were varied to attain a final T concentration of 0-10 nM in the wells. In the positive control and blank well, 50 μL experimental buffer was injected. The initial kinetics of template binding was followed by monitoring the change in fluorescence intensities for several hours depending on the reaction rates.

The required volumes for this method are tabulated in Table S8.

### Experimental protocol for testing the stability of tetrameric products

This protocol was used to gather data reported in figure S17.

#### Step 1 for baseline

20 pL of 20% Tween20 in 1X TAE and 1 M NaCl, and 0-130 pL of experimental buffer were added to each experiment and control well. Then 10 µL of pre-annealed monomer complexes *ĈD_s_* (200 nM stock) was added to the buffer in the experiment wells to obtain a final concentration of 10 nM (calculated for the total reaction volume of 200 µL, which is attained after Step 2). For the positive controls, 3-10 µL of pre-annealed labelled *ĀD_s_* (200 nM stock concentration) was added to get final concentration of 3-10 nM. For the blank, nothing was added except Tween20 and experimental buffer. The fluorescence values of all required channels were simultaneously measured over at least 10 data points of step 1.

#### Step 2 for reaction kinetics

Next, 0 - 10 µL of pre-annealed *AB̄* (200 nM stock), and 40-50 µL of experimental buffer were added to the wells to make the final volume in each well 200 pL. *AB̄* injection volumes were varied to attain a final concentration of 0-10 nM in the wells. In the positive control and blank well, 50 µL experimental buffer was injected. The initial kinetics of template binding was followed by monitoring the change in fluorescence intensities for several hours depending on the reaction rates.

The required volumes for this method are tabulated in Table S16.

### Experimental protocol for testing global specificity of templated catalysis

This protocol was used to gather data reported in Figure 5B.

#### Step 1 for baseline

20 pL of 20% Tween20 in 1X TAE and 1 M NaCl, and 50-130 pL of experimental buffer were added to each experiment and control well. Then 10 μL of pre-annealed blocked monomer complexes *AL*_1_, *ĀL*_1_, *BL*_2_*, CL*_1_*, DL*_2_, (200 nM stock) and free final monomers *C_s_, D_s_, E* (200 nM stock) were added to the buffer in the experiment wells to obtain a final concentration of 10 nM of each component (calculated for the total reaction volume of 200 μL, which is attained after Step 2).

Several positive controls were used in this experiment. For ^5^ *P* 3-10 μL of pre-annealed complexes *AB* and *DE* (200 nM stocks) were added to the wells first, and the plate was shaken at 600 rpm on a double orbital shaker for 30 seconds. Then 3-10 μL *C* (200 nM stock) were added to the positive controls to get final concentration of 3-10 nM of each strand in the wells. A fraction of *C* therefore simultaneously binds to both *AB* and *DE* to make the pentamer. The same strategy was used for *ABCBE* and *ADCDE*. For ^4^ *P*, *AB* and *CD_s_* were mixed. For *AD_s_*, *ABC_s_*, and *ĀDE*, pre-annealed complexes were used directly. In each positive control well, 10 nM of *D_s_* was added to account for the baseline signal that was present in the experimental wells from the beginning due to the presence of free *D_s_* with the FAM label. These systems were then used as control signals for fluorescence monitoring, and also as the size marker in PAGE experiments. For the blank, nothing was added except Tween20 and experimental buffer. All the wells had equal volume of 150 µL at this point. The fluorescence values of each required channel were simultaneously measured over at least 10 data points at the end of step 1. Reactions were performed in triplicate simultaneously.

#### Step 2 for reaction kinetics

In the first reaction well, 50 µL of the experimental buffer was added via the injector of the plate reader. In all other reaction wells, 8 µL of the appropriate templates (100 nM stocks) and 42 µL of experimental buffer were injected to attain a final template concentration of 4 nM. The total injected volume of template and buffer was 50 µL for each experimental well. In the positive control and blank wells, 50 µL experimental buffer was injected. All well had the same final volume of 200 µL. The initial kinetics of template binding were followed by monitoring the change in fluorescence intensities for several minutes to several hours depending on the reaction rates.

The required volumes for this method are tabulated in Table S15.

### Experimental protocol for testing template specificity for binding to the blocked monomers

This protocol was used to gather data reported in figure S18.

#### Step 1 for baseline

20 pL of 2% Tween20 in 1X TAE and 1 M NaCl, and 120-130 pL of experimental buffer were added to each experiment and control well. Then 10 µL of pre-annealed blocked monomer complexes *AL*_1_ / *ĀL*_1_ / *BL*_2_/*DL*_2_ (200 nM stock) were added to the buffer in the experiment wells to obtain a final concentration of 10 nM (calculated for the total reaction volume of 200 pL, which is attained after Step 2).

For the positive controls, 3-10 µL of pre-annealed monomer-template complex (200 nM stock concentration) was added to get final concentration of 3-10 nM. For the blank, nothing was added except Tween20 and experimental buffer. All the wells had equal volume of 150 µL at this point. The fluorescence values were measured over at least 10 data points. Reactions were performed in triplicate simultaneously.

#### Step 2 for reaction kinetics

In the first reaction well, 50 µL of the experimental buffer was added via the injector of the plate reader. In next two reaction wells, 4 and 10 pL of ^3^*T_ABC_* (100 nM stock) were added. Similarly, in the next two wells, 4 and 10 pL of ^3^*T_ĀDE_* (100 nM stock) were added. Volumes of injected buffer were adjusted to make to total injection volume of template and buffer 50 µL for each well. In the positive control and blank wells, 50 μL experimental buffer was injected. All well had the same final volume of 200 µL. The initial kinetics of template binding were followed by monitoring the change in fluorescence intensities for several minutes to several hours depending on the reaction rates.

The required volumes for this method are tabulated in Table S17.

### Experimental protocol for testing template specificity for trimer formation with each set of monomers

This protocol was used to gather data reported in figure S19 and S20.

#### Step 1 for baseline

20 pL of 2% Tween20 in 1X TAE and 1 M NaCl, and 100-130 pL of experimental buffer were added to each experiment and control well. Then 10 µL of monomers *AL*_1_, *BL*_2_, and *C_s_* or *ĀL*_1_, *DL*_2_, and *E* (200 nM stock) were added to the buffer in the experiment wells to obtain a final concentration of 10 nM of each monomer(calculated for the total reaction volume of 200 μL, which is attained after Step 2).

For the positive controls, 5 and 10 µL of pre-annealed trimer *ABC_s_* or *ĀDE* (200 nM stock concentration) was added to get final concentration of 5 and 10 nM. For the blank, nothing was added except Tween20 and experimental buffer. All the wells had equal volume of 150 µL at this point. The fluorescence values were measured over at least 10 data points. Reactions were performed in triplicate simultaneously.

#### Step 2 for reaction kinetics

In the first reaction well, 50 µL of the experimental buffer was added via the injector of the plate reader. In next two reaction wells, 4 and 10 pL of ^3^*T_ABC_* (100 nM stock) were added. Similarly, in the next two wells, 4 and 10 pL of ^3^*T_ĀDE_* (100 nM stock) were added. Volumes of injected buffer were adjusted to make to total injection volume of template and buffer 50 µL for each well. In the positive control and blank wells, 50 µL experimental buffer was injected. All well had the same final volume of 200 µL. The initial kinetics of template binding were followed by monitoring the change in fluorescence intensities for several minutes to several hours depending on the reaction rates.

The required volumes for this method are tabulated in Table S18.

### Experimental protocol for testing template specificity for trimer formation in a mixture of all monomers

This protocol was used to gather data reported in figure 5A.

#### Step 1 for baseline

20 pL of 2% Tween20 in 1X TAE and 1 M NaCl, and 70-130 pL of experimental buffer were added to each experiment and control well. Then 10 µL of all monomers *AL*_1_, *BL*_2_, *C_s_*, *ĀL*_1_, *DL*_2_, and *E* (200 nM stock) were added to the buffer in the experiment wells to obtain a final concentration of 10 nM of each monomer(calculated for the total reaction volume of 200 µL, which is attained after Step 2).

For the positive controls, 5 and 10 µL of pre-annealed trimer *ABC_s_* or *ĀDE* (200 nM stock concentration) was added to get final concentration of 5 and 10 nM. For the blank, nothing was added except Tween20 and experimental buffer. All the wells had equal volume of 150 µL at this point. The fluorescence values were measured over at least 10 data points. Reactions were performed in triplicate simultaneously.

#### Step 2 for reaction kinetics

In the first reaction well, 50 µL of the experimental buffer was added via the injector of the plate reader. In next two reaction wells, 4 and 10 pL of ^3^*T_ABC_* (100 nM stock) were added. Similarly, in the next two wells, 4 and 10 pL of ^3^*T_ĀDE_* (100 nM stock) were added. Volumes of injected buffer were adjusted to make to total injection volume of template and buffer 50 µL for each well. In the positive control and blank wells, 50 µL experimental buffer was injected. All well had the same final volume of 200 µL. The initial kinetics of template binding were followed by monitoring the change in fluorescence intensities for several minutes to several hours depending on the reaction rates.

The required volumes for this method are tabulated in Table S19.

### Experimental protocol for systematic omission of monomers

This protocol was used to gather data reported in figure S24.

#### Step 1 for baseline

20 pL of 2% Tween20 in 1X TAE and 1 M NaCl, and 90-130 pL of experimental buffer was added to each experiment and control well. Then 10 μL each of pre-annealed blocked monomer complexes *AL*_1_, *BL*_2_*, CL*_1_*, DL*_2_, (200 nM stock) and free final monomer *E* or *Ē* (200 nM stock) was added to the buffer in the experiment wells to obtain a final concentration of 10 nM of all but one component (calculated for the total reaction volume of 200 μL, which is attained after Step 2). For the positive controls, 3-10 μL of pre-annealed complexes *AB* and *DE* (200 nM stocks) was added to the wells first, and the plate was shaken at 600 rpm on a double orbital shaker for 30 seconds. Then 3-10 μL *C* (200 nM stock) was added to the positive controls to get final concentration of 3-10 nM of each strand in the wells. A fraction of *C* therefore simultaneously binds to both *AB* and *DE* to make the pentamer. This system is then used as a control signal for the fluorescence monitoring, and also as the size marker in PAGE experiments. For the blank, nothing was added except Tween20 and experimental buffer. The fluorescence values of each required channel were simultaneously measured over at least 10 data points at the end of step 1.

#### Step 2 for reaction kinetics

8 pL of ^5^*T* or ^5^*T_Ē_* (100 nM stocks) and 42 μL experimental buffer were injected into the experimental wells via the injector of the plate reader to make the final volume in each well 200 µL. Template injection volumes were set to attain a final ^5^*T* or ^5^*T_Ē_* concentration of 4 nM in the wells. In the positive control and blank well, 50 µL experimental buffer was injected. The initial kinetics of template binding were followed by monitoring the change in fluorescence intensities for several minutes to several hours depending on the reaction rates.

The required volumes for this method are tabulated in Table S23.

### Experimental protocol for stepwise addition of monomers in pentamer formation at standard monomer concentration

This protocol was used to gather data reported in figure S21.

#### Step 1 for baseline

20 pL of 20% Tween20 in 1X TAE and 1 M NaCl, and 80-130 pL of experimental buffer were added to each experiment and control well. Then 10 µL of pre-annealed blocked monomer complex *AL*_1_ were added to the reaction wells. For the positive controls, 3-10 pL of pre-annealed complexes *AB* and *DE* (200 nM stocks) was added to the wells first, and the plate was shaken at 600 rpm on a double orbital shaker for 30 seconds. Then 3-10 µL *C* (200 nM stock) was added to the positive controls to get final concentration of 3-10 nM of each strand in the wells. A fraction of *C* therefore simultaneously binds to both *AB* and *DE* to make the pentamer. This system was then used as a control signal for the fluorescence monitoring, and also as the size marker in PAGE experiments. For the blank, nothing was added except Tween20 and experimental buffer. The fluorescence values of each required channel were simultaneously measured over at least 10 data points at the end of step 1.

#### Step 2 for reaction kinetics

4-10 pL of ^5^*T* or (100 nM stocks) and 40-46 µL experimental buffer were injected into the experimental wells via the injector of the plate reader. Template injection volumes were set to attain a final ^5^*T* concentration of 2-5 nM in the wells in 200 µL. In the positive control and blank wells, 50 µL experimental buffer were injected. The initial kinetics of template binding were followed by monitoring the change in fluorescence intensities for several minutes to several hours depending on the reaction rates.

#### Step 3 for *BL*_2_ addition

After approximately 2 hours, 10 µL of pre-annealed blocked monomer complex *BL*_2_ (200 nM stocks) were added to the reaction wells. The fluorescence signal was monitored continuously.

#### Step 4 for *CL*_1_ addition

After approximately 5 hours, 10 µL of pre-annealed blocked monomer complex *CL*_1_ (200 nM stocks) were added to the reaction wells. The fluorescence signal was monitored continuously.

#### Step 5 for *DL*_2_ addition

After approximately 15 hours, 10 µL of pre-annealed blocked monomer complex *DL*_2_ (200 nM stocks) were added to the reaction wells. The fluorescence signal was monitored continuously.

#### Step 6 for *E* addition

After approximately 8 hours, 10 µL of *E* (200 nM stocks) were added to the reaction wells. The fluorescence signal was monitored continuously.

The required volumes for this method are tabulated in Table S20.

### Experimental protocol for stepwise addition of monomers in pentamer formation at high monomer concentration

This protocol was used to gather data reported in figure S22.

#### Step 1 for baseline

20 pL of 20% Tween20 in 1X TAE and 1 M NaCl, and 30-130 pL of experimental buffer were added to each experiment and control well. Then 20 µL of pre-annealed blocked monomer complex *AL*_1_ were added to the reaction wells. For the positive controls, 5-20 pL of pre-annealed complexes *AB* and *DE* (1000 nM stocks) was added to the wells first, and the plate was shaken at 600 rpm on a double orbital shaker for 30 seconds. Then 5-20 µL *C* (1000 nM stock) was added to the positive controls to get final concentration of 25-100 nM of each strand in the wells. A fraction of *C* therefore simultaneously binds to both *AB* and *DE* to make the pentamer. This system was then used as a control signal for the fluorescence monitoring, and also as the size marker in PAGE experiments. For the blank, nothing was added except Tween20 and experimental buffer. The fluorescence values of each required channel were simultaneously measured over at least 10 data points at the end of step 1.

#### Step 2 for reaction kinetics

0-10 pL of ^5^*T* or (100 nM stocks) and 40-50 µL experimental buffer were injected into the experimental wells via the injector of the plate reader. Template injection volumes were set to attain a final ^5^*T* concentration of 0-5 nM in the wells in 200 µL. In the positive control and blank wells, 50 µL experimental buffer were injected. The initial kinetics of template binding were followed by monitoring the change in fluorescence intensities for several minutes to several hours depending on the reaction rates.

#### Step 3 for *BL*_2_ addition

After approximately 80 minnutes, 20 pL of pre-annealed blocked monomer complex *BL*_2_ (1000 nM stock) were added to the reaction wells. The fluorescence signal was monitored continuously.

#### Step 4 for *CL*_1_ addition

After approximately 200 minutes, 20 µL of pre-annealed blocked monomer complex *CL*_1_ (1000 nM stock) were added to the reaction wells. The fluorescence signal was monitored continuously.

#### Step 5 for *DL*_2_ addition

After approximately 18 hours, 20 µL of pre-annealed blocked monomer complex *DL*_2_ (1000 nM stock) were added to the reaction wells. The fluorescence signal was monitored continuously.

#### Step 6 for *E* addition

After approximately 150 minutes, 20 µL of *E* (1000 nM stock) were added to the reaction wells. The fluorescence signal was monitored continuously.

The required volumes for this method are tabulated in Table S21.

### Experimental protocol for addition of variable numbers of monomer types in pentamer formation

This protocol was used to gather data reported in figure S23.

#### Step 1 for baseline

20 pL of 20% Tween20 in 1X TAE and 1 M NaCl, and 80-130 pL of experimental buffer were added to each experiment and control well. Then 10 µL of pre-annealed blocked monomer complex *AL*_1_ was added in the first experimental well. 10 µL of *AL*_1_ and *BL*_2_ were added in the second well. 10 µL of *AL*_1_, *BL*_2_, and *CL*_1_ were added to the third well. 10 µL of *AL*_1_, *BL*_2_, *CL*_1_, and *DL*_2_ were added to the fourth well. And 10 µL of *AL*_1_, *BL*_2_, *CL*_1_, *DL*_2_, and *E* were added to the fifth well. All monomers had final concentration of 10 nM for all but one component (calculated for the total reaction volume of 200 µL, which is attained after Step 2). For the positive controls, 3-10 µL of pre-annealed *M*^5^*T* complexes (200 nM stocks) were added to separate wells. For the blank, nothing was added except Tween20 and experimental buffer. All the wells had equal volume of 150 µL at this point. The fluorescence values of each required channel were simultaneously measured over at least 10 data points at the end of step 1.

#### Step 2 for reaction kinetics

8 pL of ^5^*T* or (100 nM stocks) and 42 µL experimental buffer were injected into the experimental wells via the injector of the plate reader to make the final volume in each well 200 μL. The total injected volume of template and buffer was 50 μL for each experimental well. Template injection volumes were set to attain a final ^5^*T* or ^5^*T_Ē_* concentration of 4 nM in the wells. In the positive control and blank well, 50 μL experimental buffer were injected. The initial kinetics of template binding were followed by monitoring the change in fluorescence intensities for several minutes to several hours depending on the reaction rates.

The required volumes for this method are tabulated in Table S22.

### Data processing using CLEARISSA software: working principles

This pipeline follows that in the article ‘‘Fuel-driven catalytic molecular templating’’ (*35*), we lay it out again here for completeness of the article, and convenience of the readers.

This section describes how raw fluorescence signals (AFU) are converted into inferred concentrations or into a normalised, dimensionless readout. We first define a plate-level normalisation based on positive controls; then apply assay-specific signal models and inference steps. The data processing pipeline is carried out using the in-house software Clearissa (v1.71), available at https://github.com/PrinciplesKJ/Clearissa_1.71. Step-by-step instructions for reproducing the processing described in the following sections are provided in the accompanying Zenodo submission https://doi.org/10.5281/zenodo.20822987

Raw fluorescence signals can exhibit time-dependent changes due to instrument drift and handling steps (for example, sealing, injections, or plate ejection and reinsertion). We assume that the measured fluorescence is the sum of contributions from the individual species present in a reaction well. We further assume that each species’ contribution is proportional to its concentration, and the *ratios* of the corresponding proportionality constants do not change under these perturbations.

### General experimental context and common notation

A ‘‘well’’ refers to an individual reaction well on a plate. Each plate contains one of the following:

- a buffer-only blank well with raw fluorescence *f*_blank_(*t*) (AFU),
- an assay-specific negative-control well with raw fluorescence *f*_neg_(*t*) (AFU),
- one or more positive-control wells containing a fluorescent reference species at known concentration(s) [*C*]*_q_*, with raw fluorescence *f*_pos*,q*_ (*t*) (AFU). In standard experiments, typically [*C*]*_q_* ∈ {3, 5, 7, 10} nM, and 25, 50, 75, 100 nM for reactions with high monomer concentrations.
- experimental wells with raw fluorescence *f*_raw_(*t*) (AFU).

Here, *f*_raw_(*t*) denotes the raw fluorescence time trace of an arbitrary experimental well; when multiple experimental wells are indexed explicitly we write *f*_raw*,i*_ (*t*).

Unless stated otherwise, background correction is performed by blank subtraction:

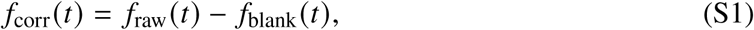

where *f*_corr_(*t*) is the blank-subtracted trace (AFU). The same procedure is applied to positive control and negative control traces.

### Definitions

- Initialisation window: a 10 data point pre-trigger interval

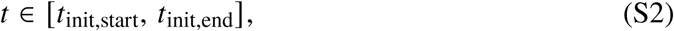

Acquired while wells contain the initial 150 μL mixture (before addition of the remaining 50 µL to reach 200 µL). This window is used to estimate a mini calibration curve.

- Oversaturation window: a 15 min interval after the injection of a saturating level of a strand or complex intended to push one species to react completely

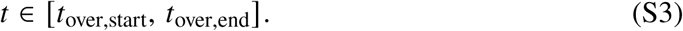

It is necessary to wait for a stable (plateau) signal before this window can be taken.

- Main measurement window: the interval used for kinetic analysis, starting at the trigger injection timepoint *t*_inj_ and ending once the signal has reached a steady state.

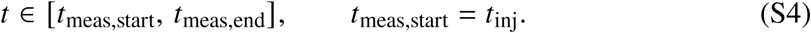

Here, *t*_meas,end_ is defined as the earliest timepoint at which the trace has reached a stable plateau; for slow reactions, *t*_meas,end_ is set to the final recorded timepoint.

- pos*_q,_*_init_: mean blank-subtracted positive-control fluorescence in the initialisation window (AFU). The index q refers to the respective known concentration of the positive control.
- *c*_io_: fluorescence-to-concentration slope (AFU per nM) of the positive control, determined from the initialisation window.
- *C*_ref_: reference concentration used when a concentration-referenced trajectory is required (typically 10 nM).
- *r_q_* (*t*): signal drift ratio for positive control *q* (dimensionless).

### Positive control normalisation

Positive control normalisation defines a time-dependent, background-corrected reference trajectory *p*_corr_(*t*) (AFU) that corrects plate-wide, time-dependent artifacts (gain drift, handling steps, plate ejection/reinsertion).

### Procedure

#### Blank subtraction

We first correct the positive control for the background blank signal.

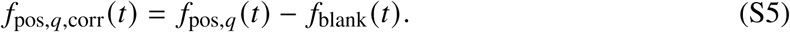

#### Calibration slope from the initialisation window

A mini calibration of fluorescence against concentration for the positive control is formed by linear regression of pos*_q,_*_init_ against [*C*]*_q_*:

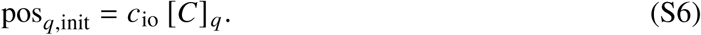

We thus assume a no-intercept model after blank subtraction. For the purposes of our analysis, *c*_io_ is taken as the ground truth with which other concentrations can be converted to fluorescence and vice versa.

#### Plate-level drift profile

Each positive-control trace is converted to a ratio relative to its own initialisation mean:

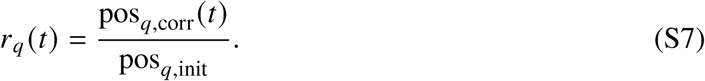

The signal drift profile is the mean across all *Q* positive controls:

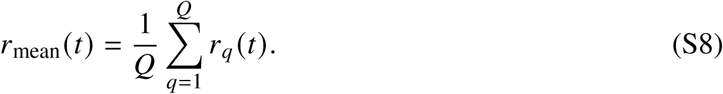

#### Reference trajectory in AFU

The information from all positive controls can be used to define a time-dependent nominal positive control fluorescence signal at any concentration *C*_ref_ according to:

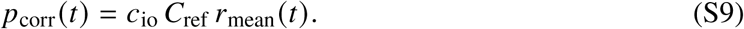

Equivalently,

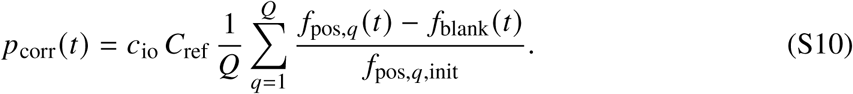

#### Applying the normalisation

For an experimental well with raw fluorescence trace *f*_raw_(*t*) (AFU), we first apply blank subtraction (Eq. S1) and then divide by *p*_corr_(*t*) to give a ratio of the signal to the positive control:

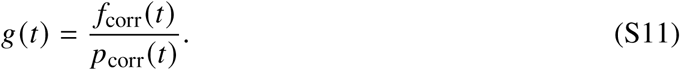

When concentration-referenced units are required, we define *h*(*t*) with units of nM:

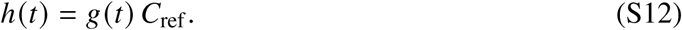

*h*(*t*) gives the concentration of positive control species that would be required to explain the observed fluorescence in the experimental trace.

### Conversion of AFU to concentrations for TMSD reactions

#### Experimental context and signal model

The assay follows a toehold-mediated strand displacement reaction of the form:

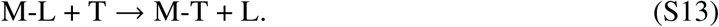

Reaction wells are initially loaded with a fluorophore-quencher duplex M-L. The non-labelled invader strand T is injected to initiate the reaction in which L is displaced and M-T is formed. M-L exhibits residual fluorescence due to incomplete quenching, whereas the displacement product M-T is fully fluorescent.

Positive-control wells contain M-T at known concentrations, which are used to construct *p*_corr_(*t*).

We apply the common preprocessing described above to obtain *h*(*t*) (nM) for each experimental trace (Eq. S12), i.e.

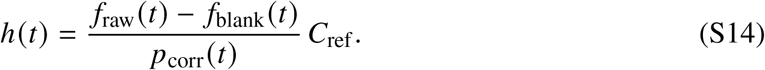

For concentration inference we assume

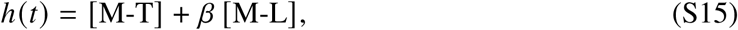

where *β* is the fractional fluorescence of M-L relative to M-T (dimensionless). We further assume mass conservation

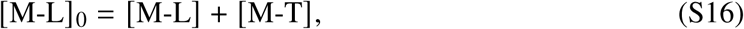

so that

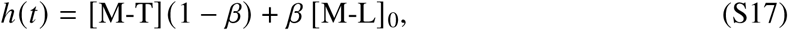

and thus

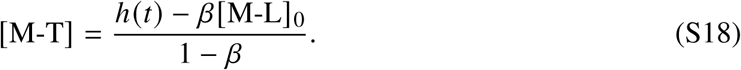

### Concentration inference procedure (per trace)

**1. Determine the initial substrate concentration** [M**-**L]_0_ **by oversaturation.** Under oversaturation by injection of excess T we assume full conversion of M-L to M-T, hence *h*(*t*) = [M-L]_0_ in the oversaturated plateau window. We estimate

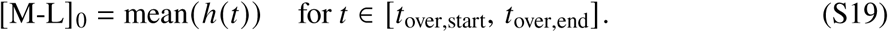

**2. Estimate** *β* **from the pre-trigger window.** In the initialisation window, [M-T] = 0 and *h*(*t*) = *β*[M-L]_0_. We compute

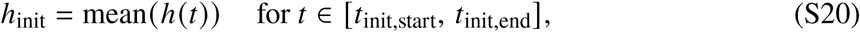

and estimate

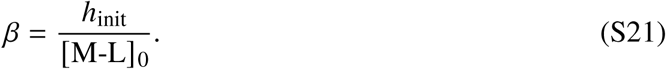

**3. Reconstruct concentrations over time** Having identified the constants [*M*-*L*]_0_ and *β*, equation S18 can be used to infer the concentrations of the product [M-T] over time.

### Calculation of a control-normalised readout in relative units

#### Experimental context

This mode reports a dimensionless normalised readout (relative fluorescence, RF) rather than concentrations. It is used when a fluorescence-to-concentration mapping is not justified (for example, complex stoichiometry, multiple fluorescent species, or unknown quenching fractions). The essential principle is to report the fluorescence on a scale normalised with respect to two controls – typically a negative control assigned the value 0 and a positive control assigned the value 1. Note that the system being reported need not contain only the fluorescent species that define these controls.

For each well, we consider the blank-corrected fluorescence *f*_corr_(*t*) (Eq. S1) and the averaged, blank-corrected positive control *p*_corr_(*t*) (Eq. S9). We also consider a blank-corrected negative control

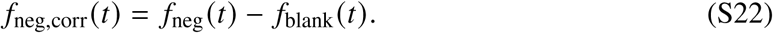

All quantities are defined for the time period corresponding to the initialisation and main measurement interval

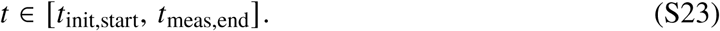

#### Negative-control baseline (AFU)

A baseline window *t* ∈ [*t*_nc,start_*, t*_nc,end_] usually within the initialisation window *t* ∈ [*t*_init,start_*, t*_init,end_], or sometimes in the main measurement interval *t* ∈ [*t*_meas,start_*, t*_meas,end_] is selected after any initial transients in the negative control due to the reaction initiation procedure have subsided. We use this time period to define a fixed negative control baseline

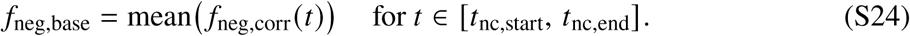

#### Relative fluorescence calculation

We then calculate a relative fluorescence as

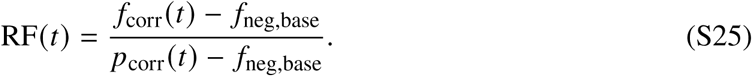

Typically, the negative control value was taken from the fluorescence of the system with 0 nM of added input, at a point immediately prior to injection. The change in fluorescence signal due to buffer injection is negligible. We highlight which wells were used for the negative control in table S6, S7, S8, S9, S10, S12, S13, S15, S17, S18, S19, S20, S21, and S22. In the majority of experiments, these negative controls allow us to correct for any fluorescence from unquenched monomers (such as labelled versions of *E* or *D_s_*) that are present throughout the reaction time course.

### Kinetic model fitting

This section describes how the inferred concentration traces from our TMSD experiments are fitted to kinetic models. The data fitting pipeline is performed using in-house software Clearissa v1.71, available at https://github.com/PrinciplesKJ/Clearissa_1.71.

Fits are performed only over a user-defined kinetic window *t* ∈ [*t*_0_*, t*_meas,end_], where *t*_0_ is the reaction-onset time chosen in the analysis software, typically aligned to the trigger injection and excluding any pre-onset baseline. The model assumes well-mixed mass-action kinetics and constant volume over the fitted window.

All TMSD reactions are modelled as an irreversible bimolecular conversion of the form

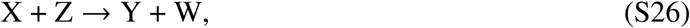

where Y is the product species whose concentration trajectory is to be fitted. W is a placeholder for any by-product(s) that are created within the reaction but are not included in the fitted rate law. The fitted observable is *y* (*t*) = [Y](*t*).

The ordinary differential equation governing the reactions is expected to be of the form

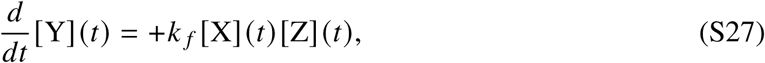

where Y is the product and X and Z are the reactants. The conservation of mass

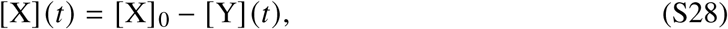

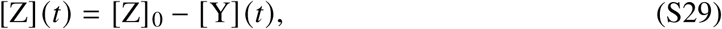

also applies. Substituting these conservation laws yields the reduced one-dimensional form used for fitting,

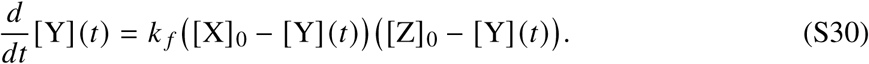

For each experimental trace, we fit a modelled trajectory *y*_sim_(*t*) to the experimentally inferred [Y](*t*), using *k _f_* and [X]_0_ as fitting parameters and using the value of [Z]_0_ calculated when inferring [Y](*t*). For a TMSD reaction,

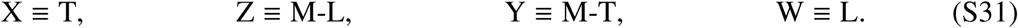

### Fitting procedure

Fits are performed independently for each trace over the fitting window *t* ∈ [*t*_0_*, t*_meas,end_], where *t*_0_ ≥ *t*_meas,start_ = *t*_inj_ is the reaction-onset time chosen in the analysis software (typically aligned to the trigger injection and excluding any pre-onset baseline). For a candidate parameter set *k _f_,* [*X*]_0_ and fixed [*Z*]_0_, the model prediction is evaluated using the analytical solution of the reduced mass-action model. Parameter estimation is performed by nonlinear least squares in Python using scipy.optimize.curve_fit with non-negativity bounds, fitting *y*_sim_(*t*) to the inferred product trace [Y](*t*) over the fitted time points. The default initial guess for the rate constant is *k _f_* = 10^5^ M^−1^s^−1^, and the initial guess for [X]_0_ is 10 nM. Fit quality is quantified using the coefficient of determination *R*^2^, computed from the residual sum of squares between [Y](*t*) and *y*_sim_(*t*), normalised by the total sum of squares of [Y](*t*) about its mean over the fitted time points.

#### Infererence of a representative rate constant from replicate fits

Replicate traces sharing the same nominal initial conditions are grouped together. Each trace in a group is fitted individually, yielding a rate constant *k _f,i_* and fitted concentration *X*_0*,i*_ for trace *i*. The fixed concentration [*Z*]_0*,i*_ (obtained during concentration inference) may also vary slightly across replicates due to experimental variations.

For each group, the representative rate constant is taken as the arithmetic mean of the replicate values *k _f,i_*. Representative concentrations are defined analogously as the arithmetic means of [*X*]_0*,i*_ and [*Z*]_0*,i*_ across replicates.

For visualisation, we plot the group mean experimental trajectory and a shaded envelope spanning the pointwise minimum and maximum across replicate traces. The group fit trajectory is obtained by simulating the kinetic model using the averaged parameters. The average of all fits are presented as the *k _f_* in the main text.

## Supplementary figures

Here we provide supporting figures and processed additional experimental data as described in the previous section.

**Figure S1:**
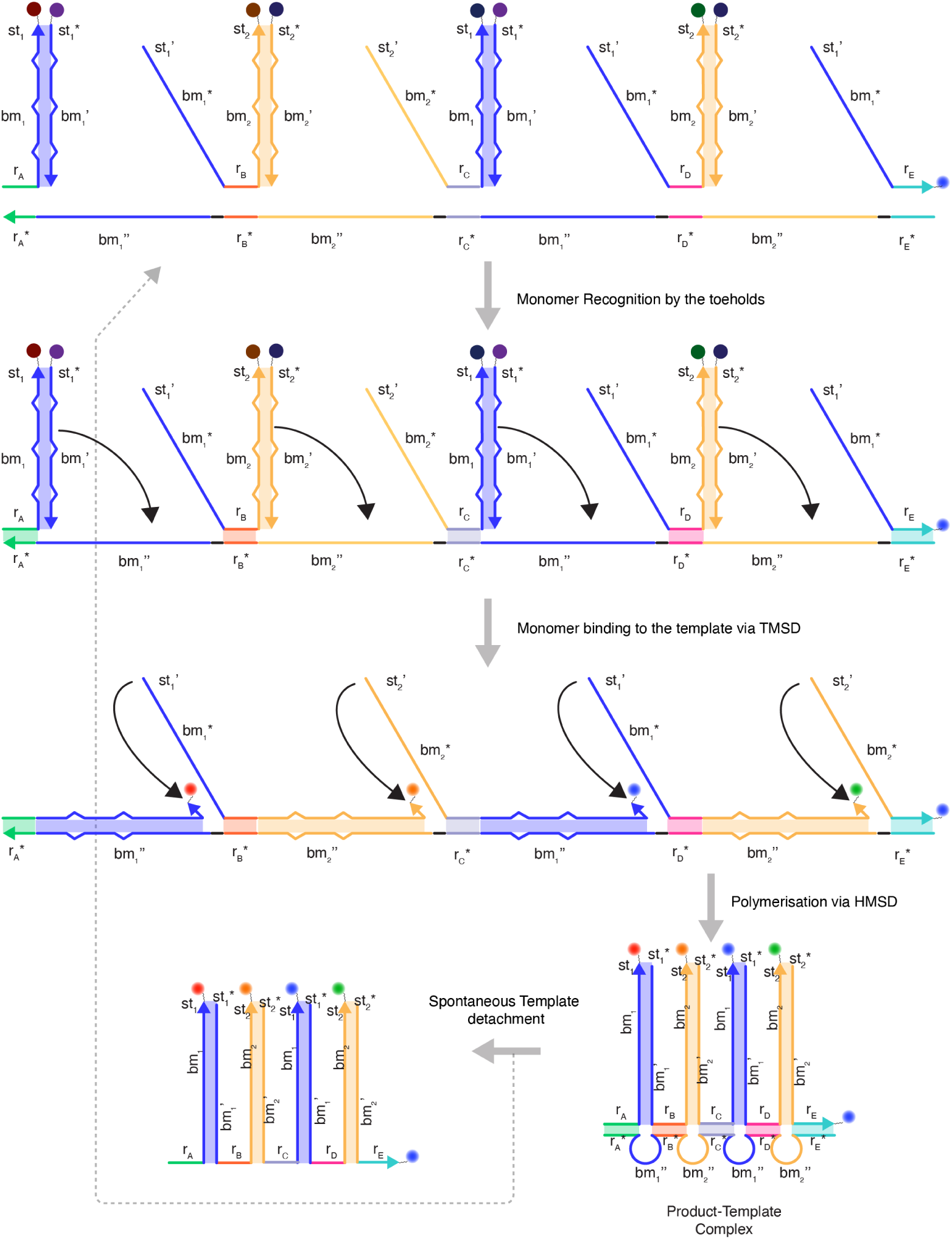
Intended pentamer formation mechanism. Note that it is not necessary for all recognition domains to bind before TMSD occurs; we are including all steps in one for simplicity of representation, and ignoring the possibility of brush formation.

**Figure S2:**
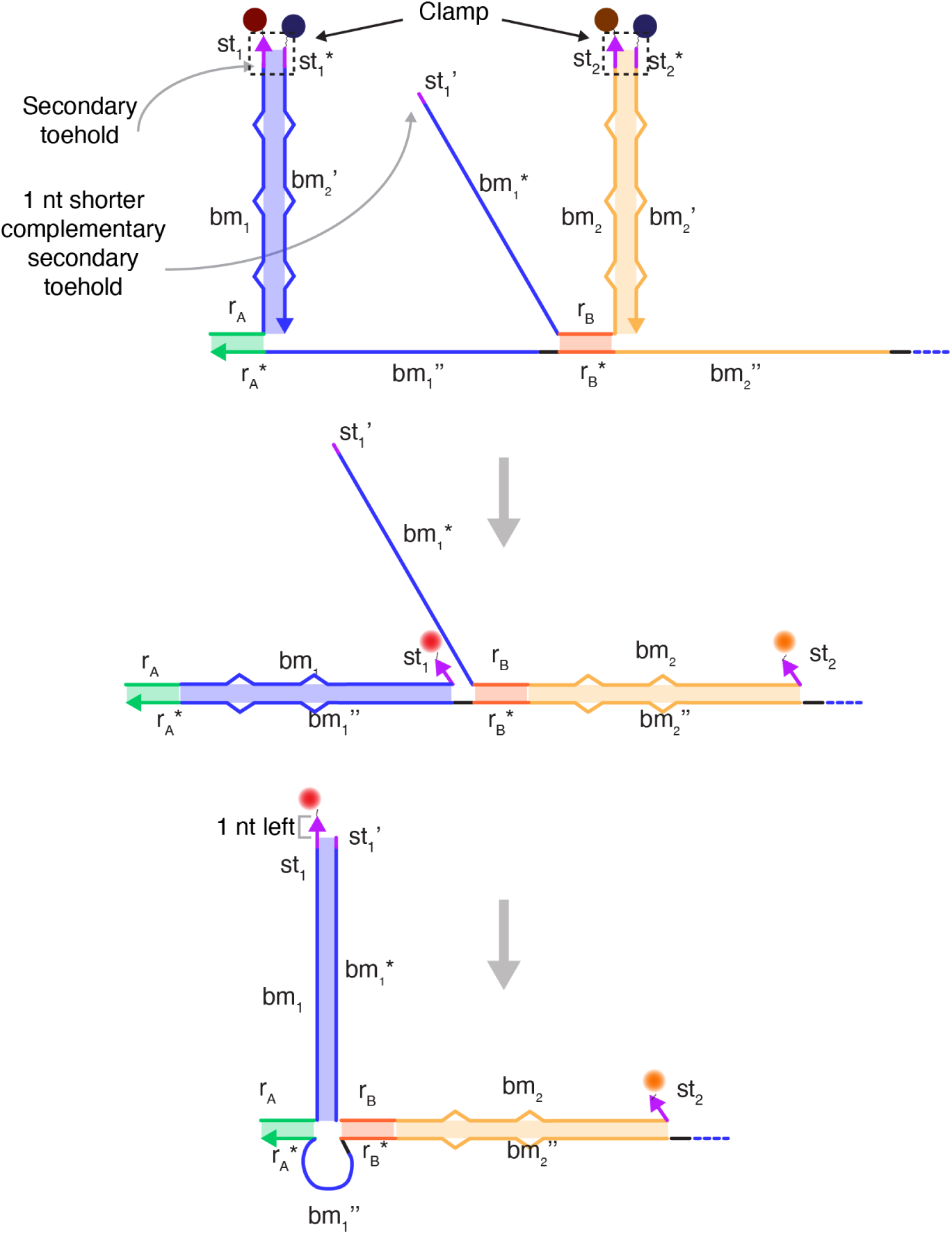
Schematic showing the role of clamps and secondary toeholds. The figure shows two adjacent monomers first binding to the template, upon which they generate the purple output toeholds (*st*_1_ and *st*_2_). During HMSD, *st_1_*^j^ and *st_2_*^j^ domains, which are complementary to *st*_1_ and *st*_2_ and truncated by one nucleotide act as secondary toeholds.

**Figure S3:**
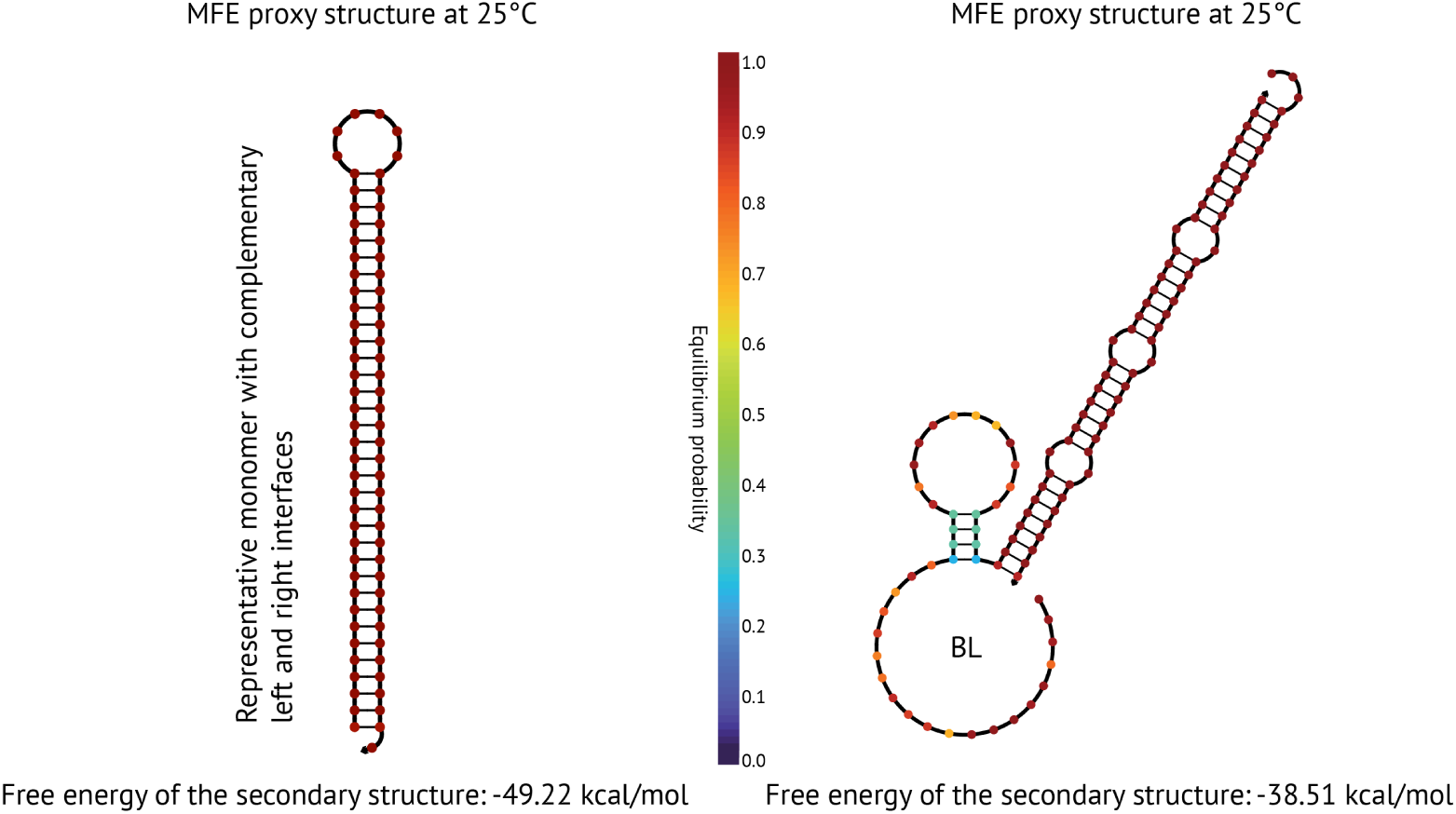
Strong hairpins and stable blocked monomers. NUPACK (*47*)-predicted structures of a dummy monomer with complementary left and right interfaces, and *B* in presence of their corresponding blocker strands shows a strong hairpin structure for the dummy, and monomer-blocker duplex for *B*.

**Figure S4:**
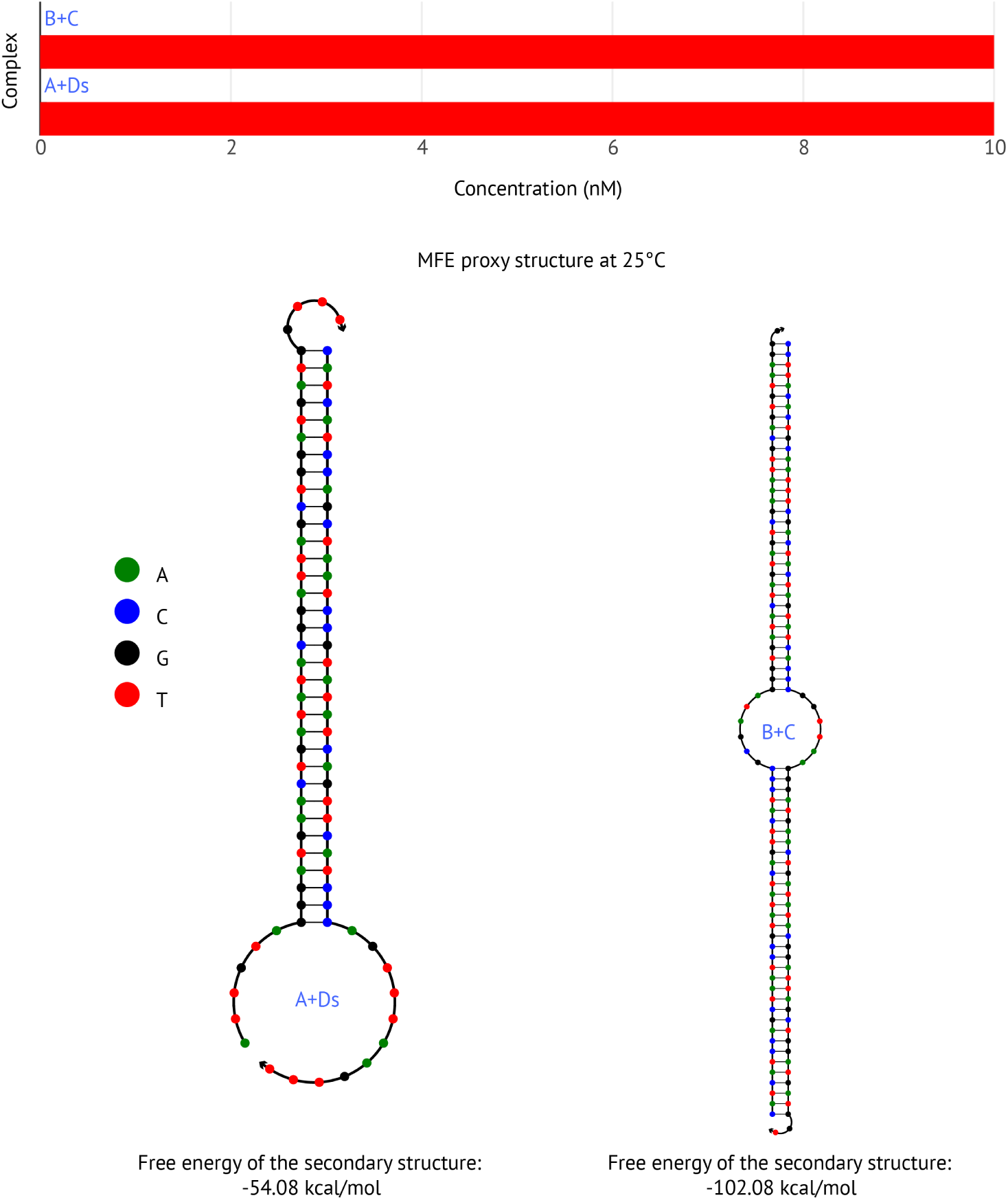
NUPACK analysis of tetramer components. NUPACK analysis of *A*, *B*, *C*, and *D_s_* predicts no tetramer and complete formation of *AD_s_* and *BC* dimers in equilibrium.

**Figure S5:**
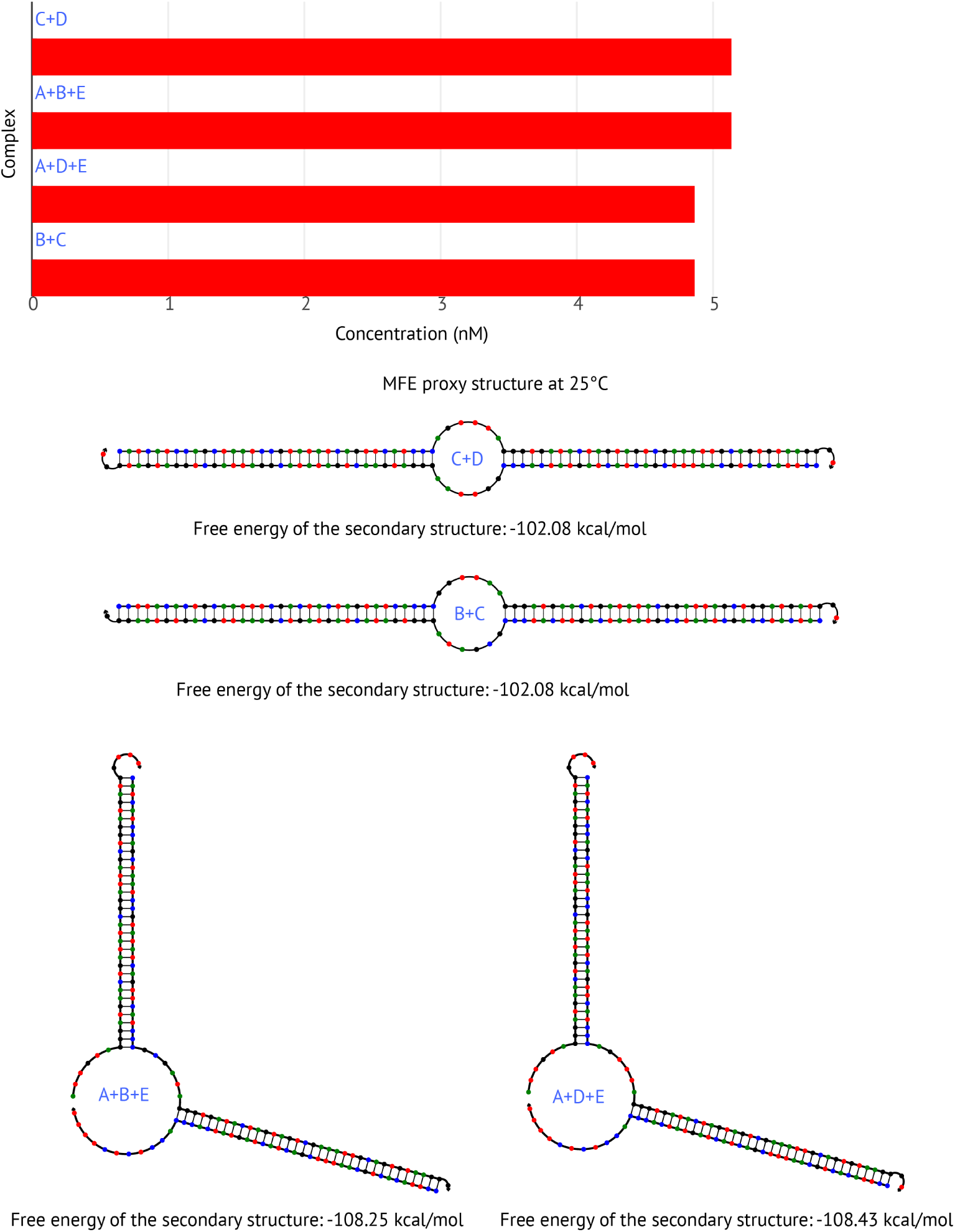
NUPACK analysis of pentamer components. NUPACK analysis of *A*, *B*, *C*, *D* and *E* predicts no pentamer. It predicts formation of *ADE*, *ABE*, *CD*, and *BC* complexes in equilibrium.

**Figure S6:**
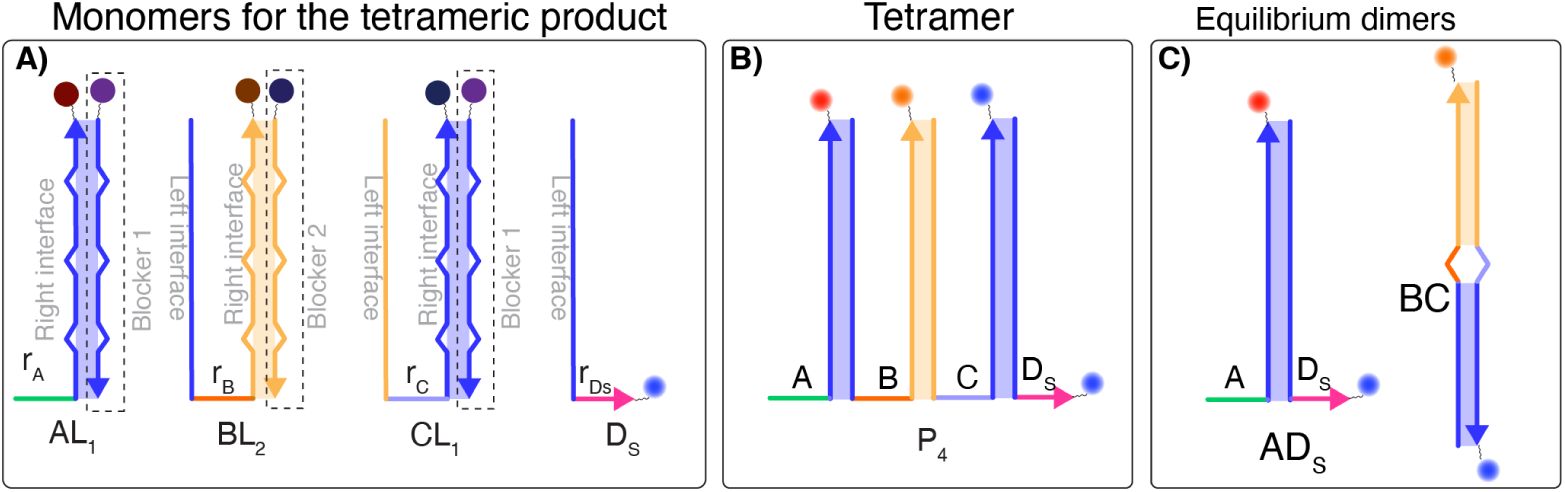
Components of the tetramer forming system. Scheme of **A)** the required monomers *A*, *B*, *C*, and *D_S_*; **B)** the desired non-equilibrium tetramer, and **C)** the more thermodynamically favoured dimers.

**Figure S7:**
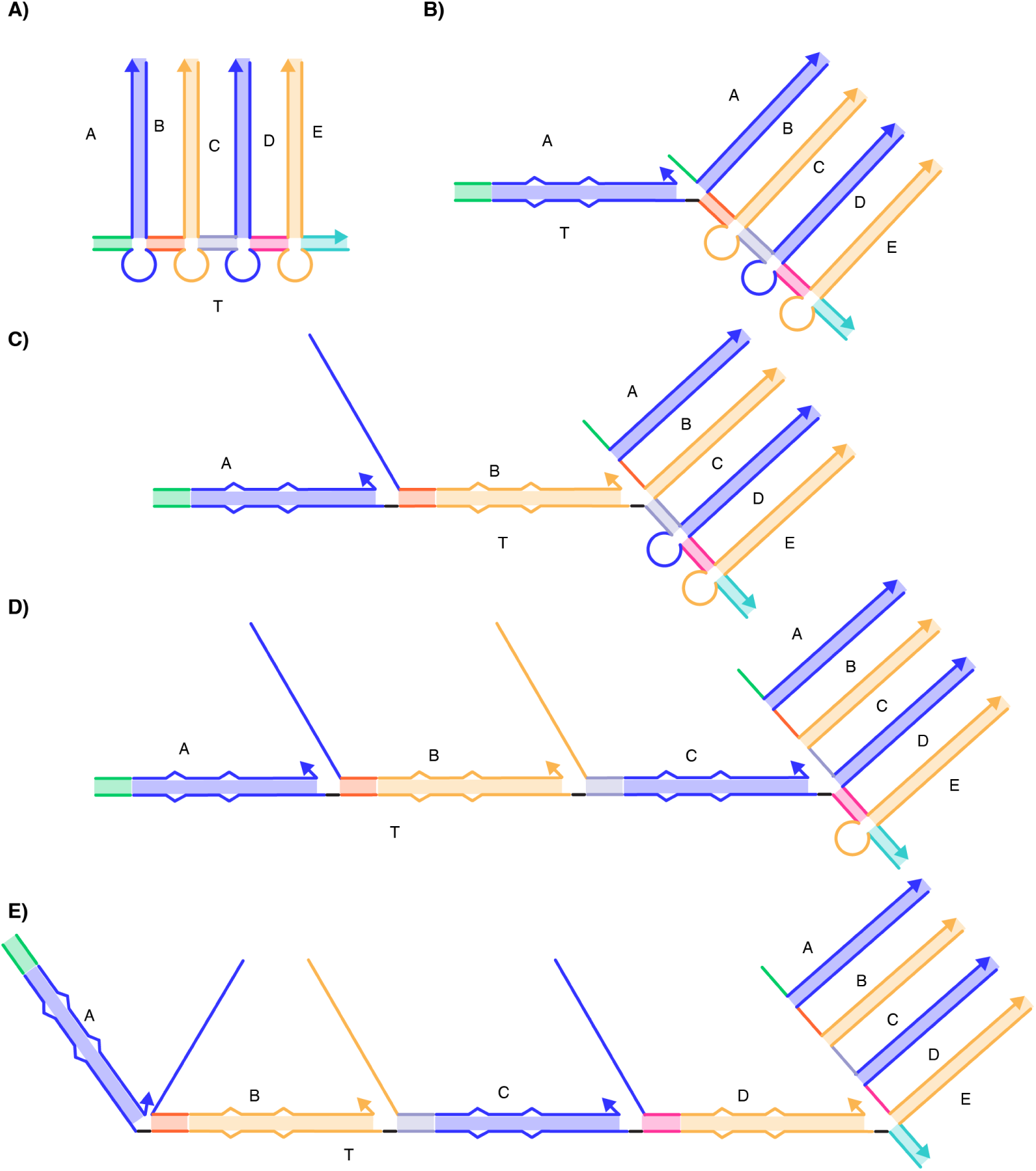
Unsaturated brush conformations. A few possible conformations of unsaturated brushes formed on pentamer template. **A)** *ABCDE* −^5^ *T*, **B)** *A* − *ABCDE* −^5^ *T*, **C)** *A* − *B* − *ABCDE* −^5^ *T*, **D)** *A* − *B* − *C* − *ABCDE* −^5^ *T*, and **E)** *A* − *B* − *C* − *D* − *ABCDE* −^5^ *T*.

**Figure S8:**
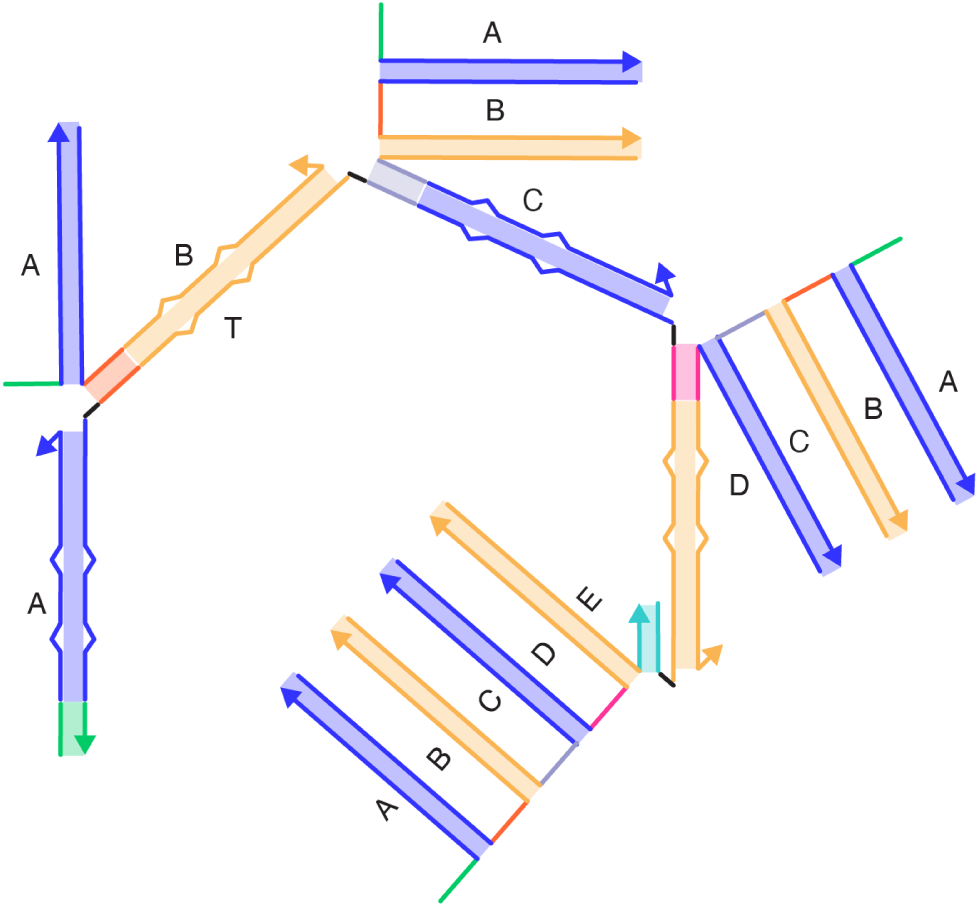
Fully saturated brush conformation. Schematic representation of of a fully saturated brush for formed on a pentamer template. The complex is formed by *A* − *AB* − *ABC* − *ABCD* − *ABCDE* −^5^ *T*.

**Figure S9:**
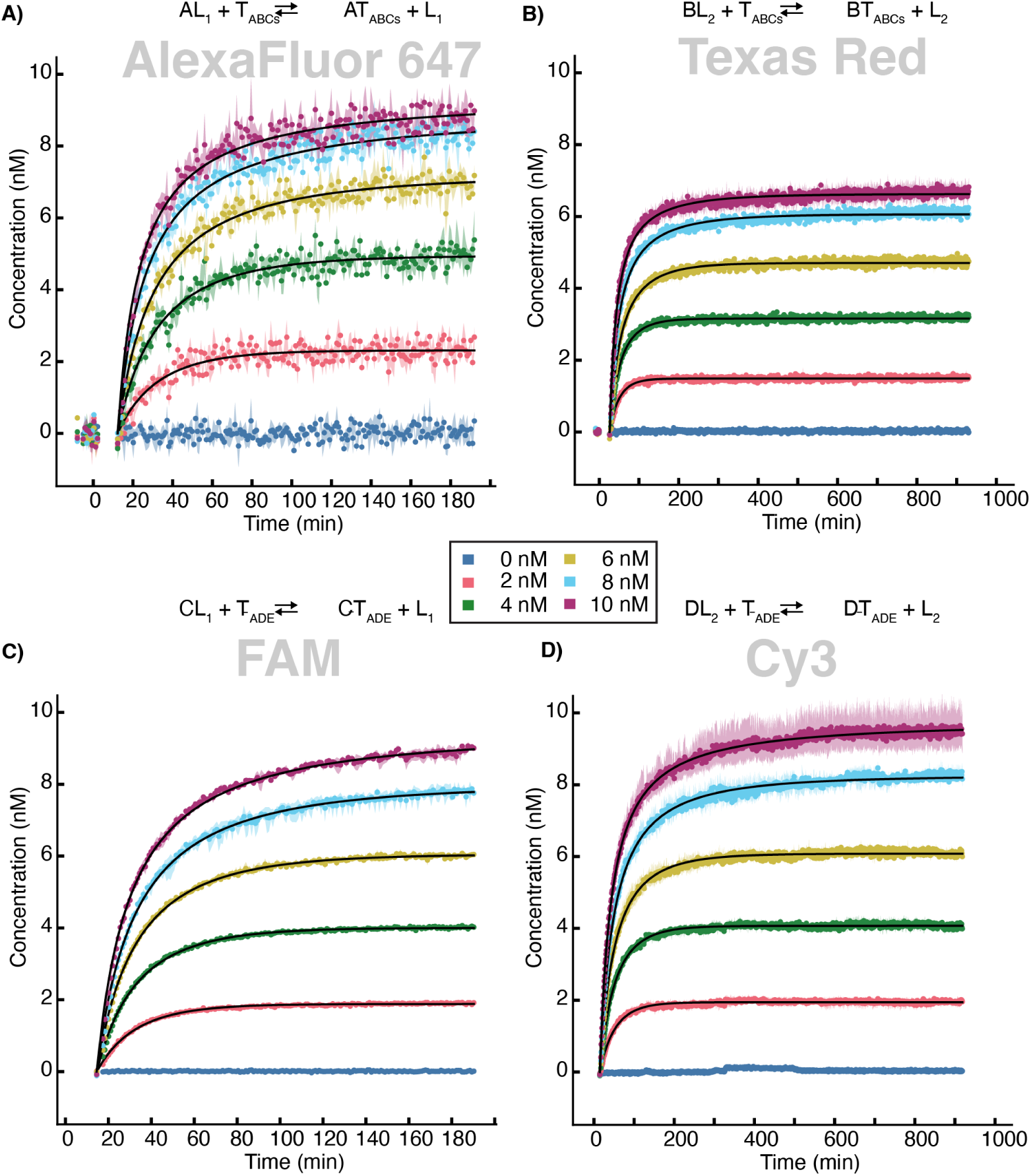
Kinetic traces showing the reaction of monomers with trimer templates. Individual monomer binding to the template ^3^*T_ABCs_* (**A** and **B**) and ^3^*T_Ā_*_*DE*_ (**C** and **D**) *via* TMSD. 2-10 nM of templates was added to 10 nM blocked monomers *AL*_1_ (**A**), *BL*_2_ (**B**) *ĀL*_1_ (**C**), or *DL*_2_ (**D**), with concentrations of reacted monomers inferred as described in the methods. Lines are fits to a second order kinetic model as described in the methods. Concentrations quoted are as intended by the protocol; our data processing pipeline provides updated estimates of these concentrations as outlined in methods.

**Figure S10:**
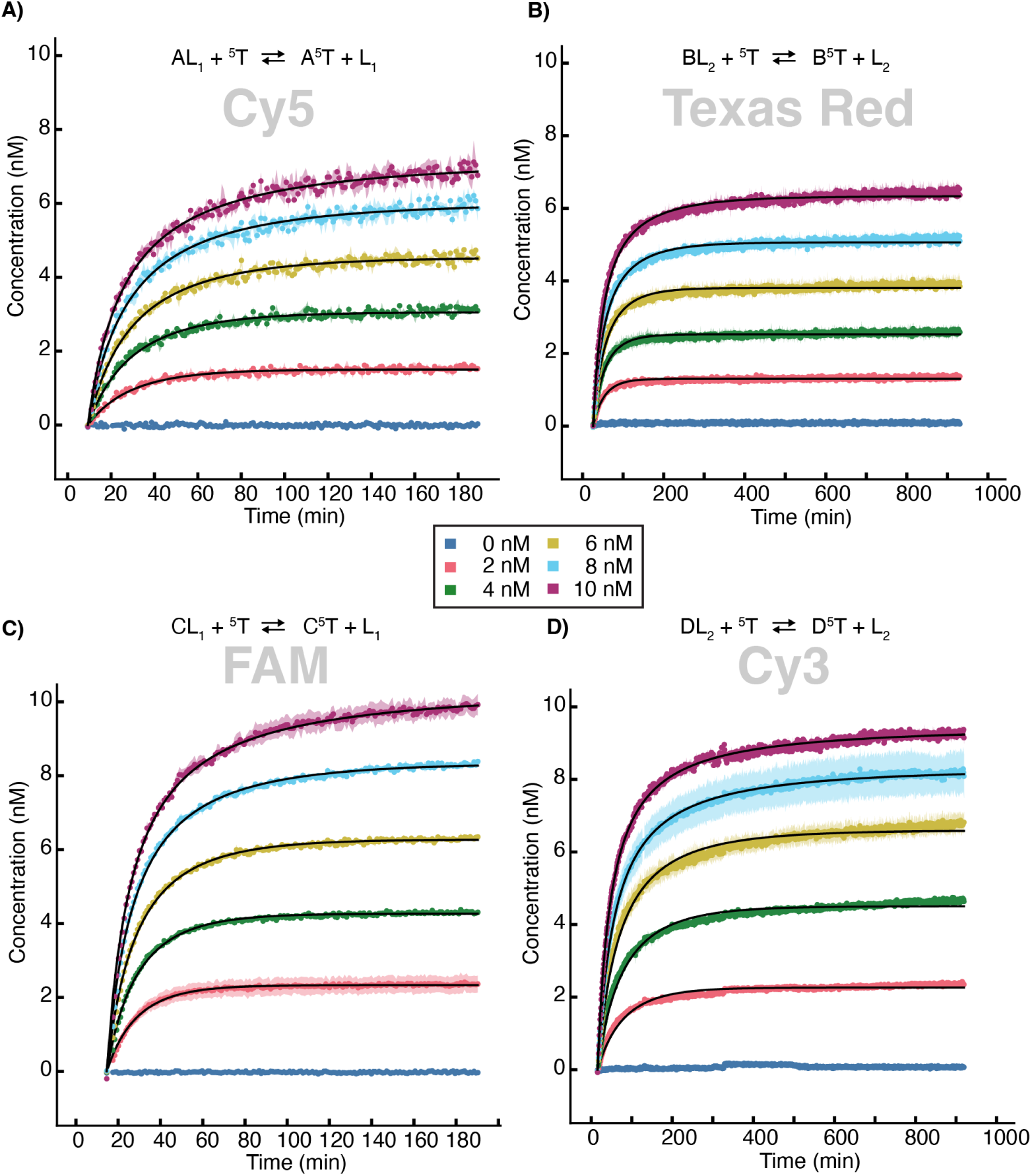
Kinetic traces showing the reaction of monomers with pentamer templates. Individual monomer binding to the template ^5^*T via* TMSD. 2-10 nM of template was added to 10 nM blocked monomers *AL*_1_ (a), *BL*_2_ (b) *CL*_1_ (c), or *DL*_2_ (d), with concentrations of reacted monomers inferred as described in the methods. Lines are fits to a second order kinetic model as decribed in the methods. Concentrations quoted are as intended by the protocol; our data processing pipeline provides updated estimates of these concentrations as outlined in the methods.

**Figure S11:**
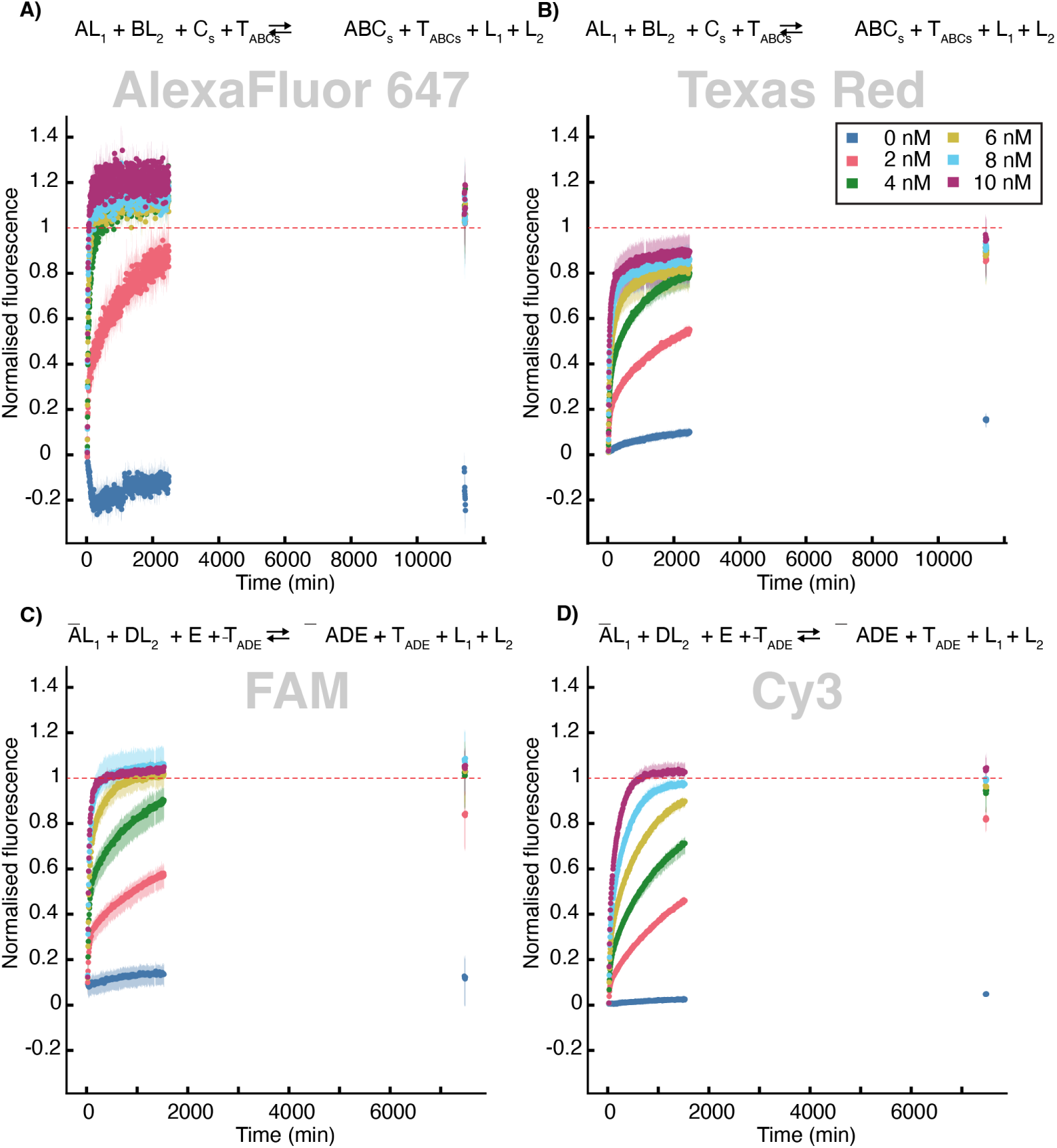
Reaction kinetics of trimer forming reactions. Fluorescence arising from monomer turnover during catalytic trimer formation, relative to a 10 nM control (*ABC_s_* or *ĀDE*) as described in the methods. 2-10 nM of ^3^*T_ABCs_* (**A**, **B**) and ^3^*T_Ā_*_*DE*_ (**C, D**) was added to a mixture of 100 nM blocked monomers *AL*_1_*, BL*_2_, and free monomer *C_s_* (in **A, B**) or *ĀL*_1_, *DL*_2_, and free monomer *E* (in **C, D**), with fluorescence reported in the channels corresponding to *A* (in **A**), *B* (in **B**), *Ā* (in **C**) and *D* (in **D**). The data was recorded in triplicate, with the mean and min-max range shown.

**Figure S12:**
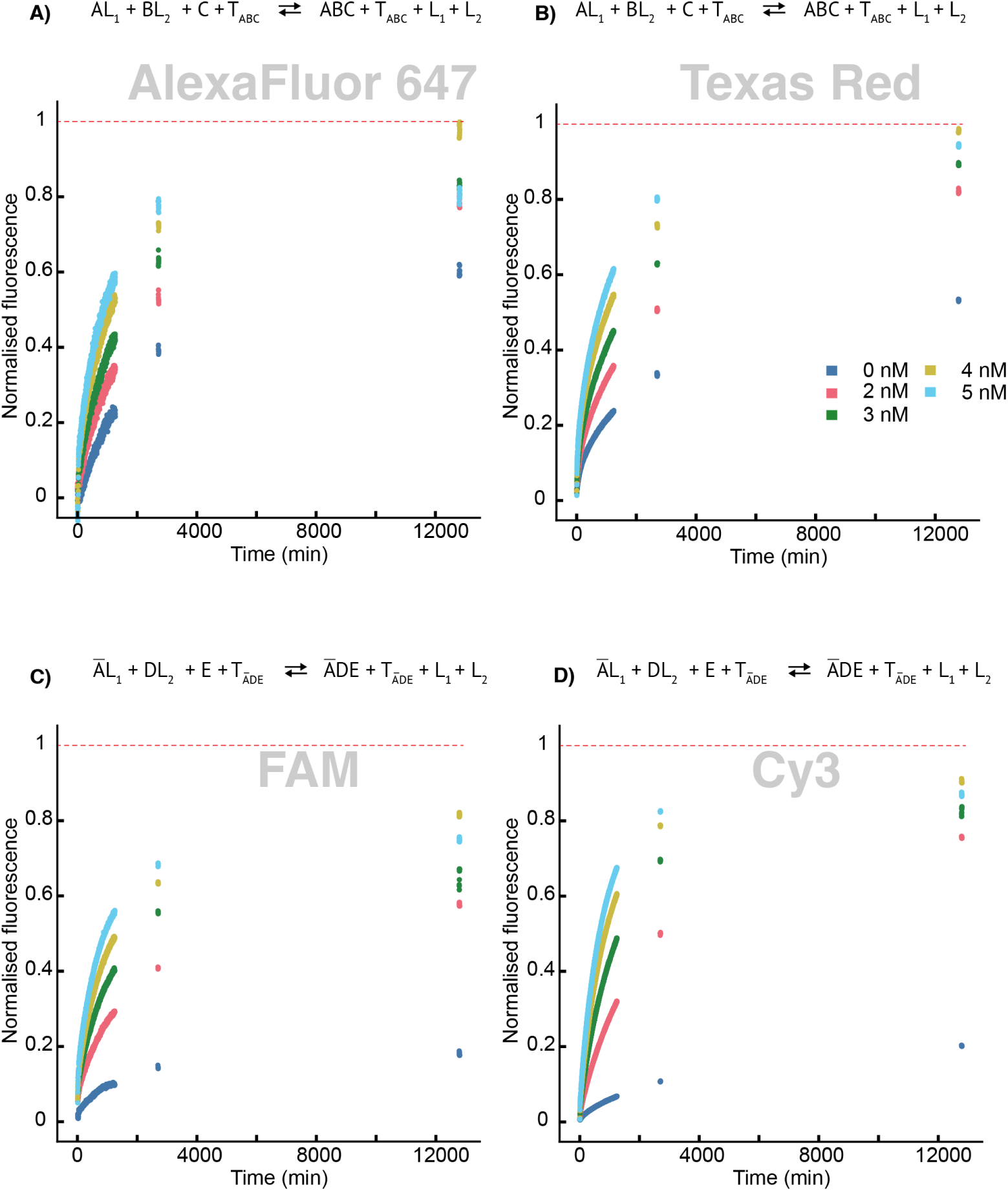
Reaction kinetics of trimer forming reactions with high monomer conformation. Fluorescence arising from monomer turnover during catalytic trimer formation, relative to a 100 nM control (*ABC_s_* or *ĀDE*) as described in the methods. 2-5 nM of ^3^*T_ABC_* (**A**, **B**) and ^3^*T_ĀDE_* (**C, D**) was added to a mixture of 100 nM blocked monomers *AL*_1_*, BL*_2_, and free monomer *C_s_* (in **A, B**) or *ĀL*_1_, *DL*_2_, and free monomer *E* (in **C, D**), with fluorescence reported in the channels corresponding to *A* (in **A**), *B* (in **B**), *Ā* (in **C**) and *D* (in **D**).

**Figure S13:**
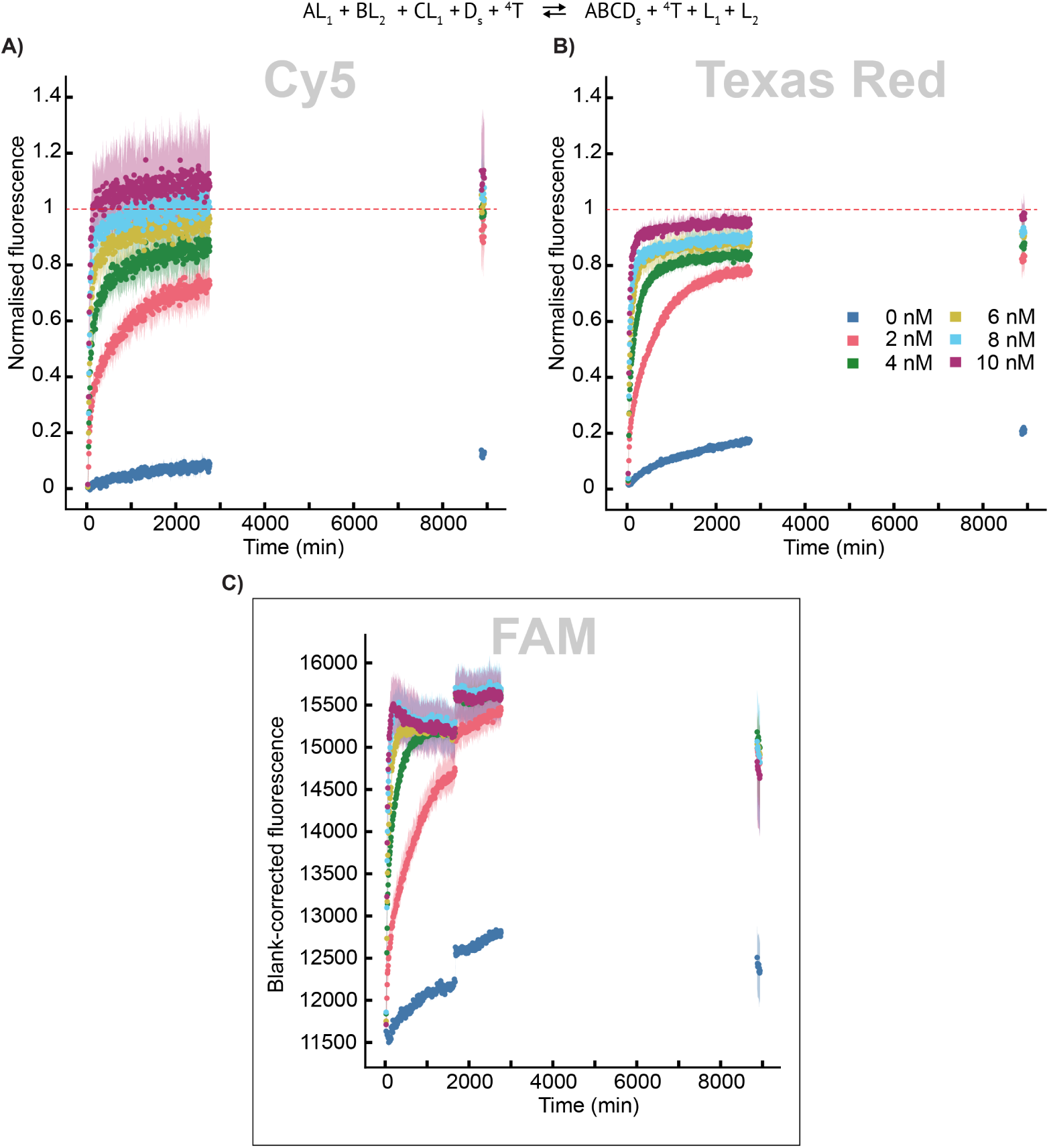
Reaction kinetics of tetramer forming reactions. Fluorescence arising from monomer turnover during catalytic tetramer formation, relative to a 10 nM control (a 1:1 mixture of *AB* and *CD_s_*) as described in the methods. 2-10 nM of ^4^*T* was added to a mixture of 10 nM blocked monomers *AL*_1_*, BL*_2_, *CL*_1_ and free monomer *D_s_* with fluorescence reported in the channels corresponding to *A* (in **A**), *B* (in **B**), and *C* (in **C**). The data was recorded in triplicate, with the mean and min-max range shown.

**Figure S14:**
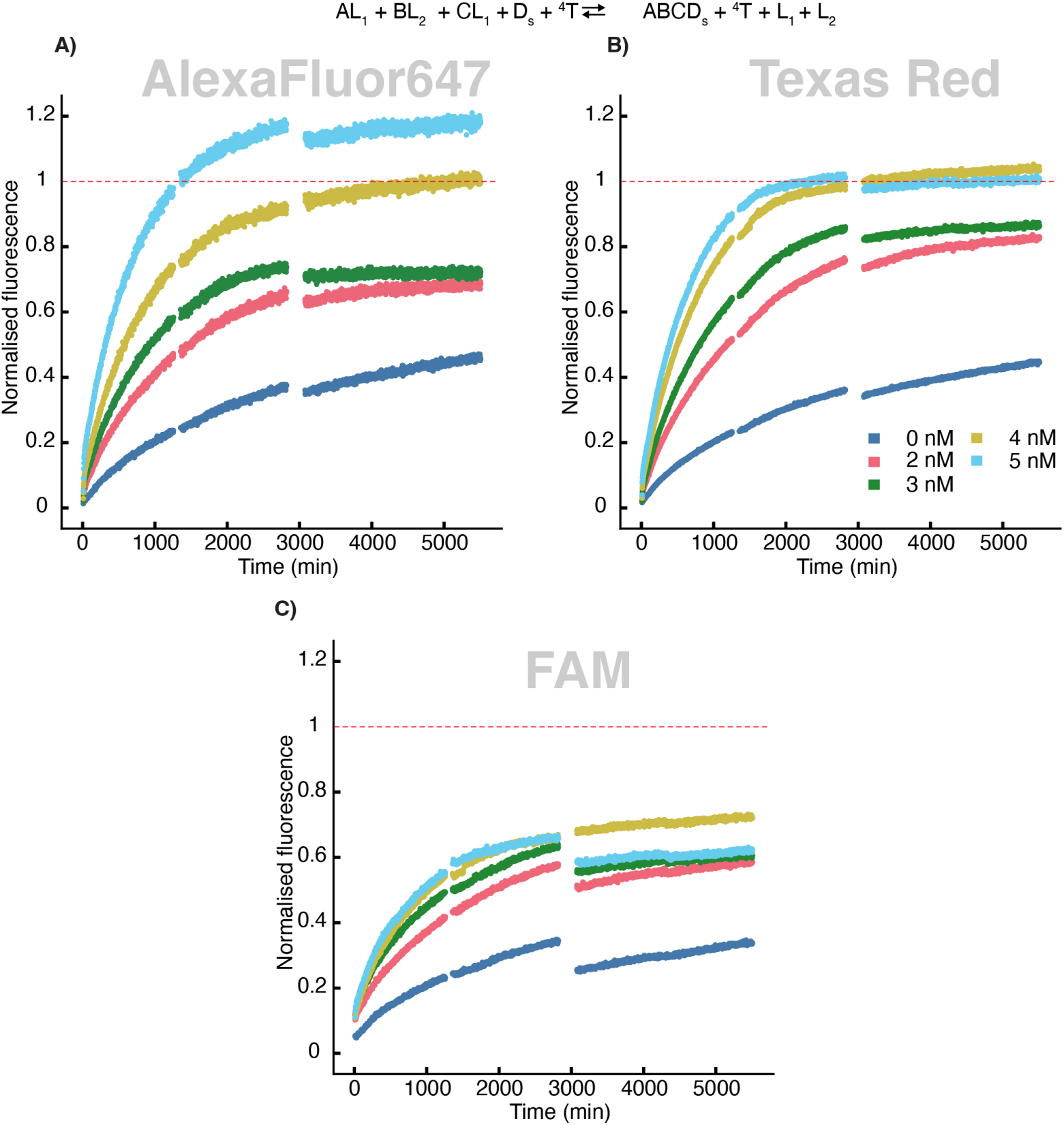
Reaction kinetics of tetramer formation with high monomer concentration. Fluorescence arising from monomer turnover during catalytic tetramer formation, relative to a 100 nM control (a 1:1 mixture of *AB* and *CD_s_*) as described in the methods. 2-5 nM of ^4^*T* was added to a mixture of 10 nM blocked monomers *AL*_1_*, BL*_2_, *CL*_1_ and free monomer *D_s_* with fluorescence reported in the channels corresponding to *A* (in **A**), *B* (in **B**), and *C* (in **C**).

**Figure S15:**
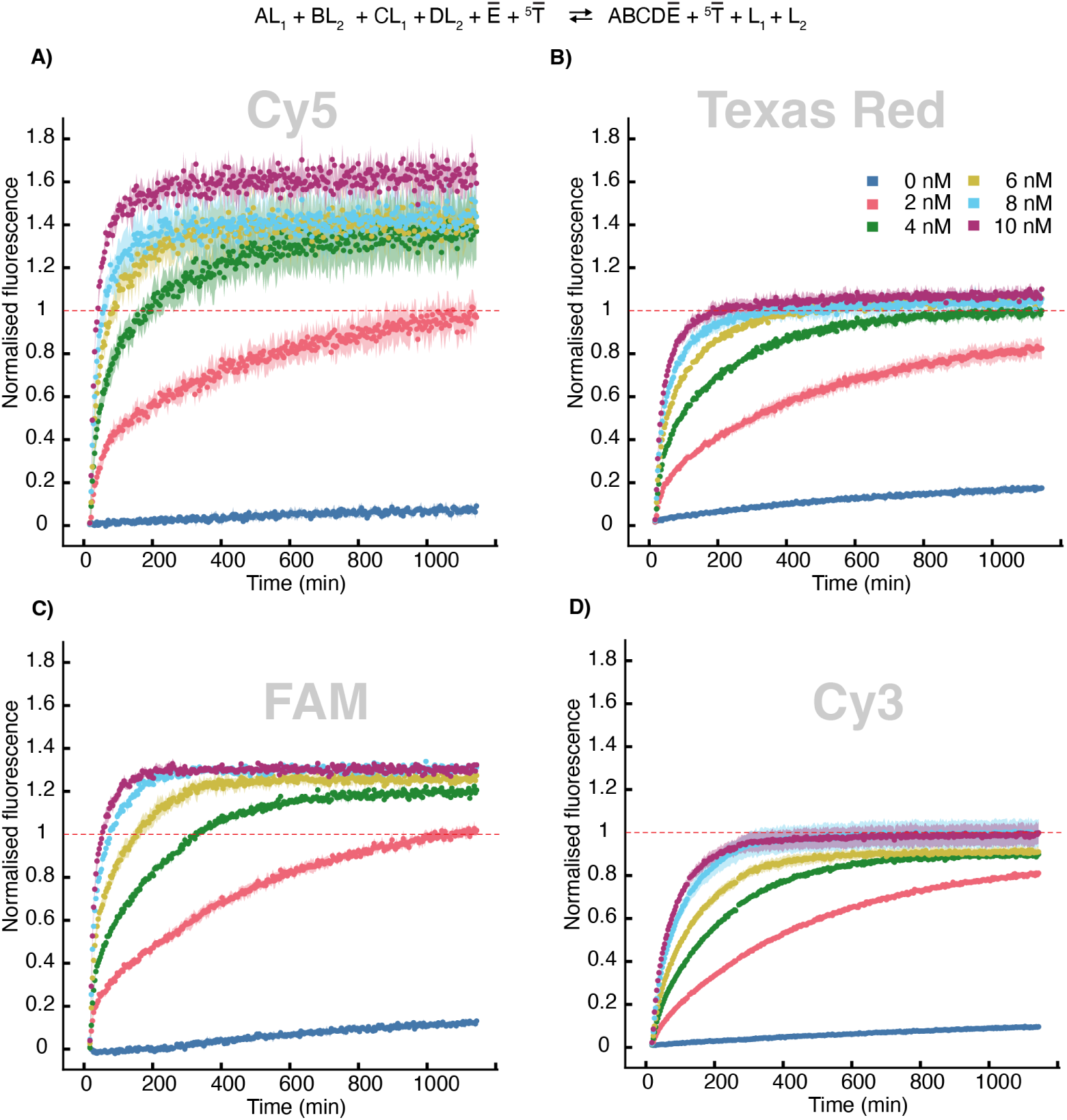
Reaction kinetics of a variant of pentamer forming reactions. Fluorescence arising from monomer turnover during catalytic pentamer formation, relative to a 10 nM control (a 1:1:1 mixture of *AB*, *DĒ*, and *C*) as described in the methods. 2-10 nM of ^5^*T_Ē_* was added to a mixture of 10 nM blocked monomers *AL*_1_, *BL*_2_, *CL*_1_, *DL*_2_, and *Ē* (which is a variant of *E*, differing only in the recognition domain) with fluorescence reported in the channels corresponding to *A* (in **A**), *B* (in **B**), *C* (in **C**), and *D* (in **D**). The data was recorded in triplicate, with the mean and min-max range shown.

**Figure S16:**
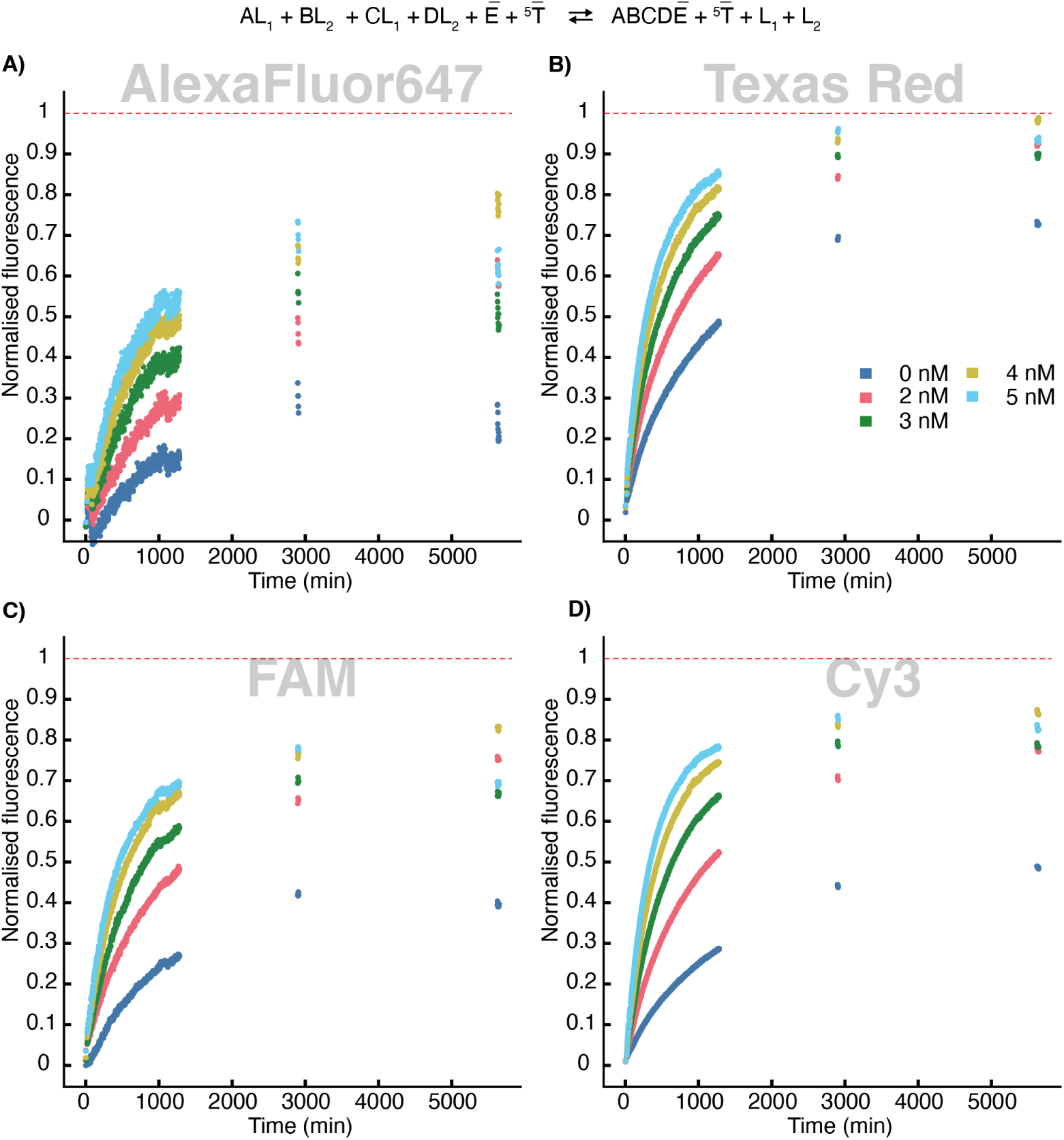
Reaction kinetics of a variant of pentamer formation with high monomer concentration. Fluorescence arising from monomer turnover during catalytic pentamer formation, relative to a 100 nM control (a 1:1:1 mixture of *AB*, *DĒ*, and *C*) as described in the methods. 2-5 nM of ^5^*T_Ē_* was added to a mixture of 100 nM blocked monomers *AL*_1_, *BL*_2_, *CL*_1_, *DL*_2_, and *Ē* (which is a variant of *E*, differing only in the recognition domain) with fluorescence reported in the channels corresponding to *A* (in **A**), *B* (in **B**), *C* (in **C**), and *D* (in **D**).

**Figure S17:**
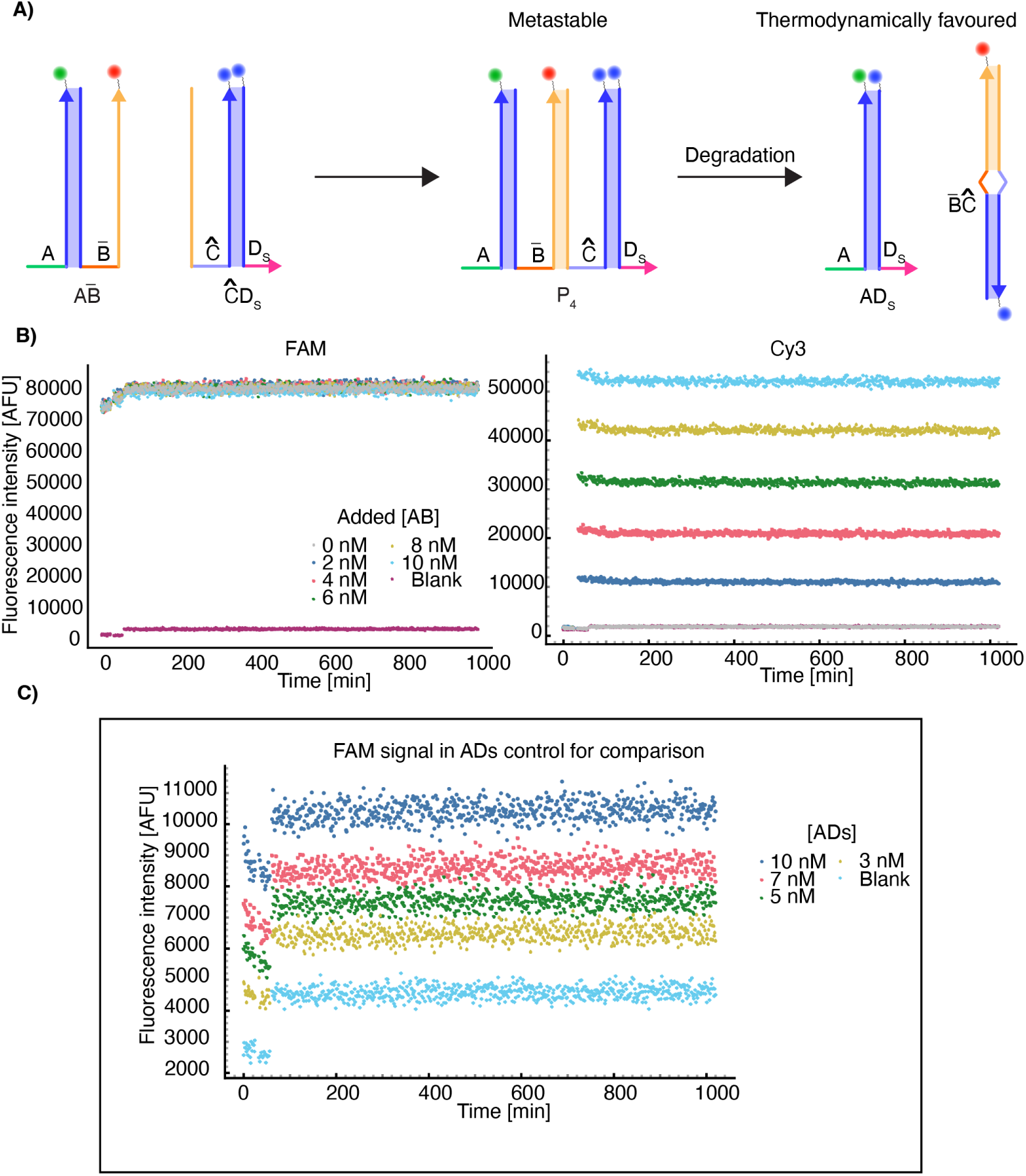
Stability test for the tetramer. **A**) Reaction schematics showing the plausible pathway for tetrameric product degradation into thermodynamically stable dimers as shown in figure S4. A FAM fluorescent label was strategically attached to the 5’ end of the terminal monomer *D_s_*. Upon degradation, when *AD_s_*is formed, the FAM label comes into close proximity with the Cy3 label of *A*, and the fluorescence intensity of FAM decreases. **B**) Signals from FAM and Cy3 channels during the reactions show little change, indicating the stability of the tetramer. **C**) Signal intensities of FAM from pre-annealed *AD_s_* control showing substantially lower fluorescence than in the experiments where *AB̄* is added to *ĈD_s_*, confirming that FAM would be quenched in this complex.

**Figure S18:**
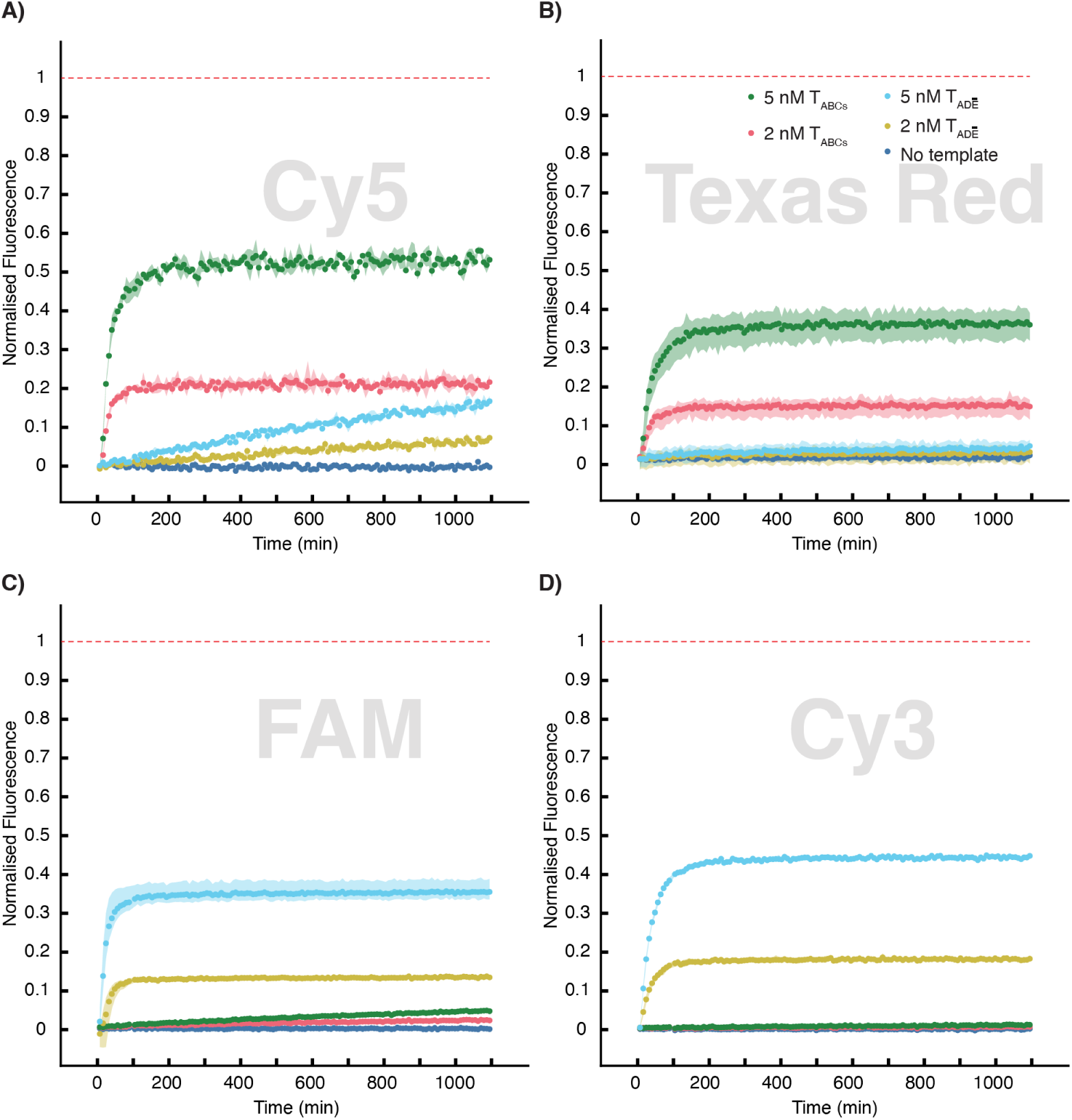
Monomer specificity for template binding. Fluorescence arising from monomers binding to the templates, normalised relative to 10 nM controls (*A*^3^*T_ABCs_* / *B*^3^*T_ABCs_* / *Ā*^3^*T_ĀDE_* / *D*^3^*T_ĀDE_*) as outlined in the methods. 2 and 5 nM of ^3^*T_ABC_* or ^3^*T_ĀDE_* was added separately to each blocked monomers *AL*_1_, *BL*_2_, *ĀL*_1_, and *DL*_2_ along with an untriggered reaction. Fluorescence reported in the channels correspond to *A* (in **A**), *B* (in **B**), *Ā* (in **C**), and *D* (in **D**). The data was recorded in triplicate, with the mean and min-max range shown.

**Figure S19:**
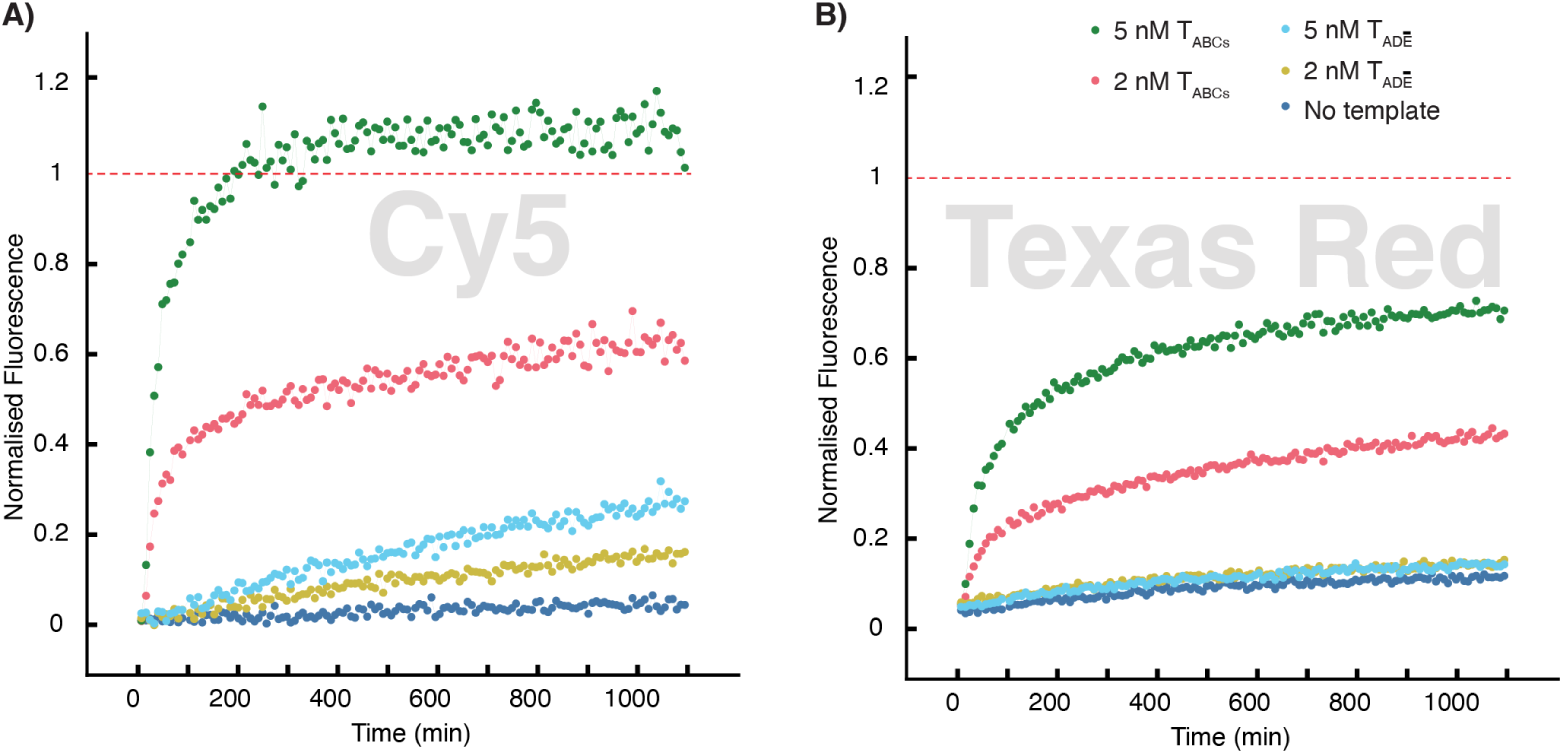
Specificity in *ABC_s_* formation. Fluorescence arising from monomers turnover in a trimer forming reaction, relative to a 10 nM control (*ABC_s_*) as outlined in the methods. 2 nM and 5 nM of ^3^*T_ABCs_* was added to a mixture of blocked monomers *AL*_1_, *BL*_2_, and free monomer *C_s_*, along with an untriggered reaction. Fluorescence reported in the channels corresponds to *A* (in a), and *B* (in b). The data was recorded in triplicate, with the mean and min-max range shown.

**Figure S20:**
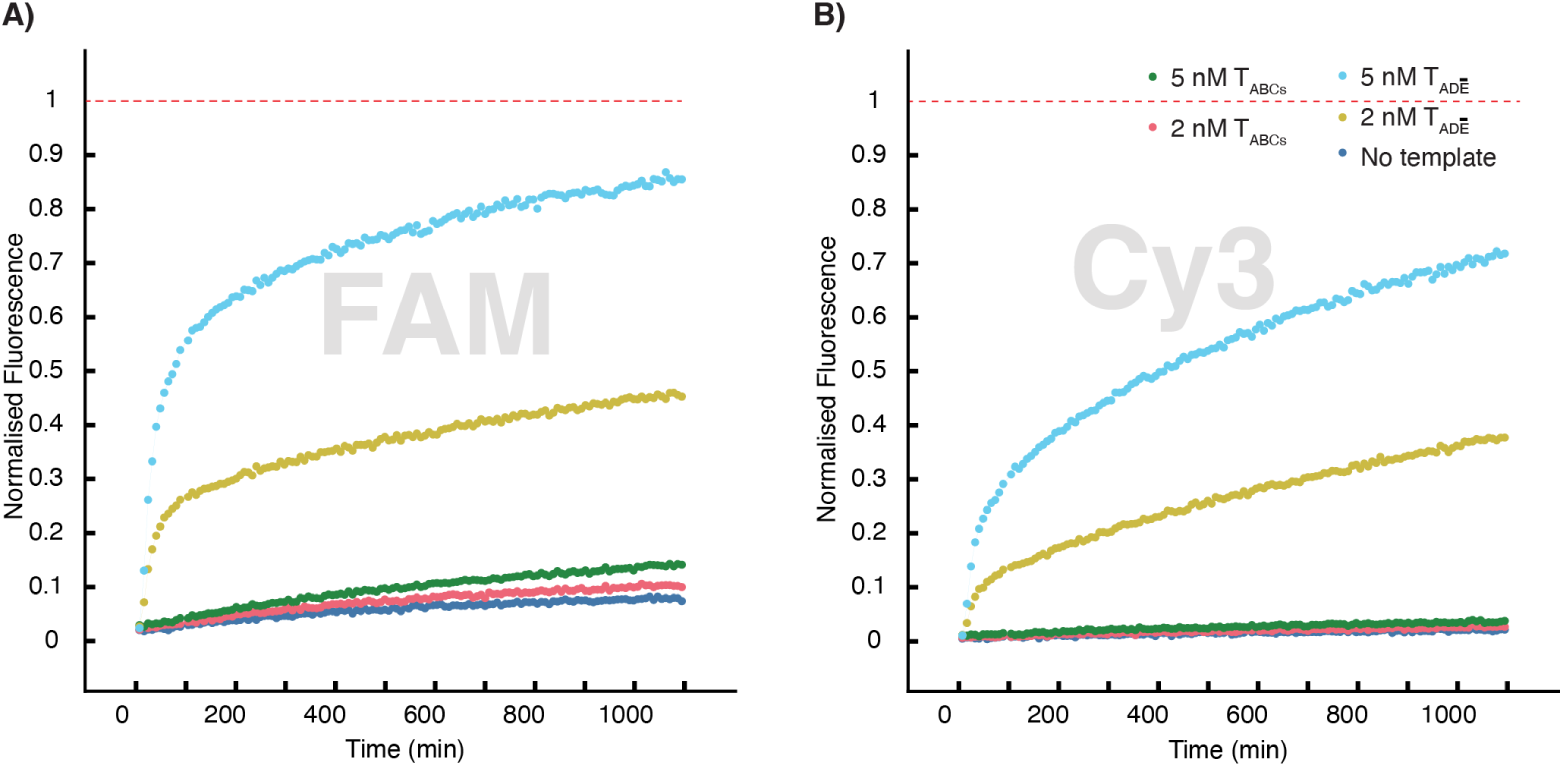
Specificity in *ĀDE* formation. Fluorescence arising from monomers turnover in a trimer forming reaction, relative to a 10 nM control (*ĀDE*) as outlined in the methods. 2 nM and 5 nM of ^3^*T_ĀDE_* was added to a mixture of blocked monomers *ĀL*_1_, *DL*_2_, and free monomer *E*, along with an untriggered reaction. Fluorescence reported in the channels corresponds to *Ā* (in a), and *D* (in b). The data was recorded in triplicate, with the mean and min-max range shown.

**Figure S21:**
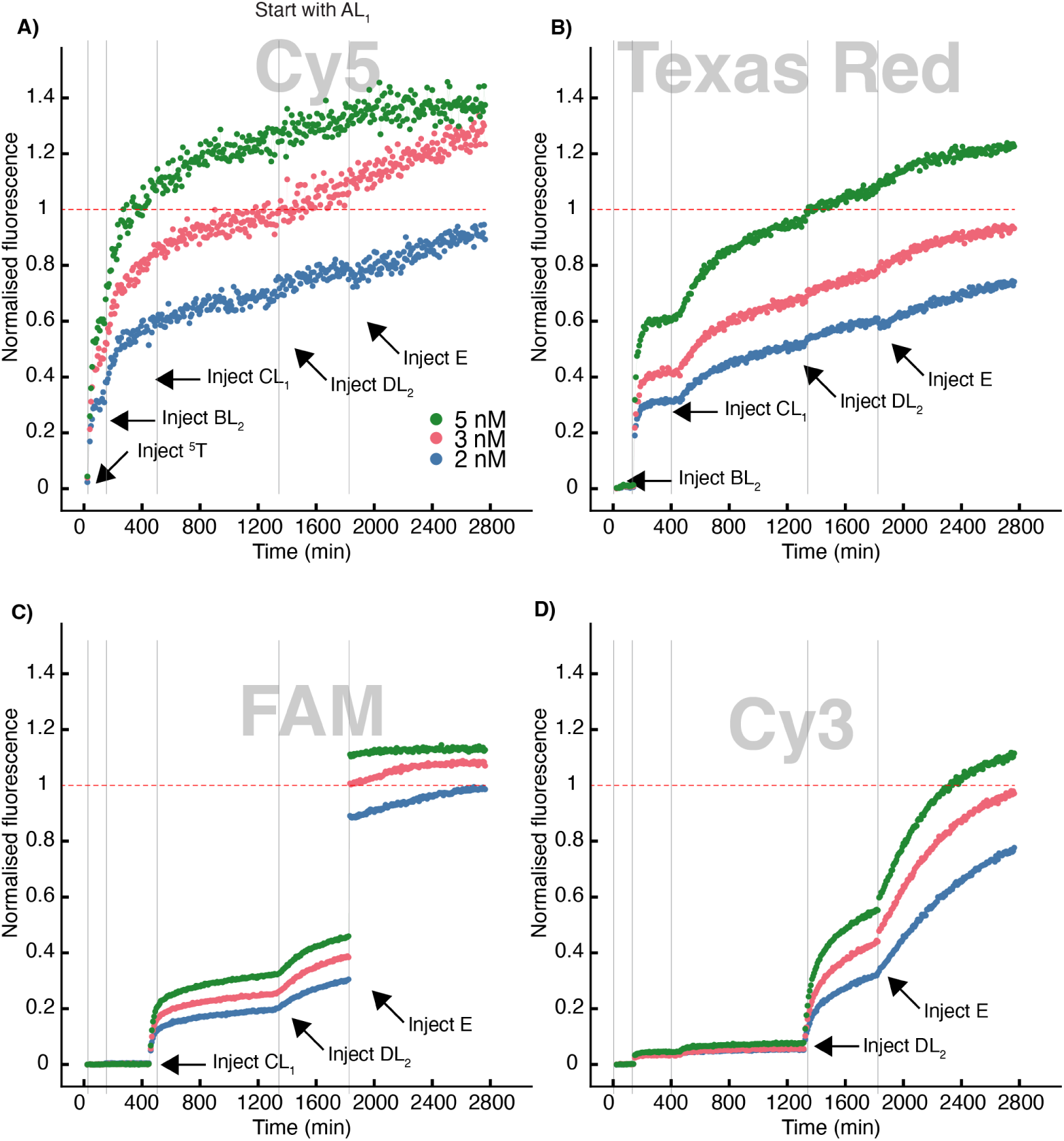
Stepwise monomer addition at standard concentration in pentamer formation reaction. Fluorescence arising from monomer turnover during catalytic pentamer formation, relative to a 100 nM control (a 1:1:1 mixture of *AB*, *DE*, and *C*) as described in the methods. 2, 3 and 5 nM of ^5^*T* was added to 10 nM blocked monomer *AL*_1_. Then 10 nM of *BL*_2_, *CL*_1_, *DL*_2_, and *E* were added in a stepwise fashion. Fluorescence reported in the channels corresponds to *A* (in **A**), *B* (in **B**), *C* & *E* (in **C**), and *D* (in **D**).

**Figure S22:**
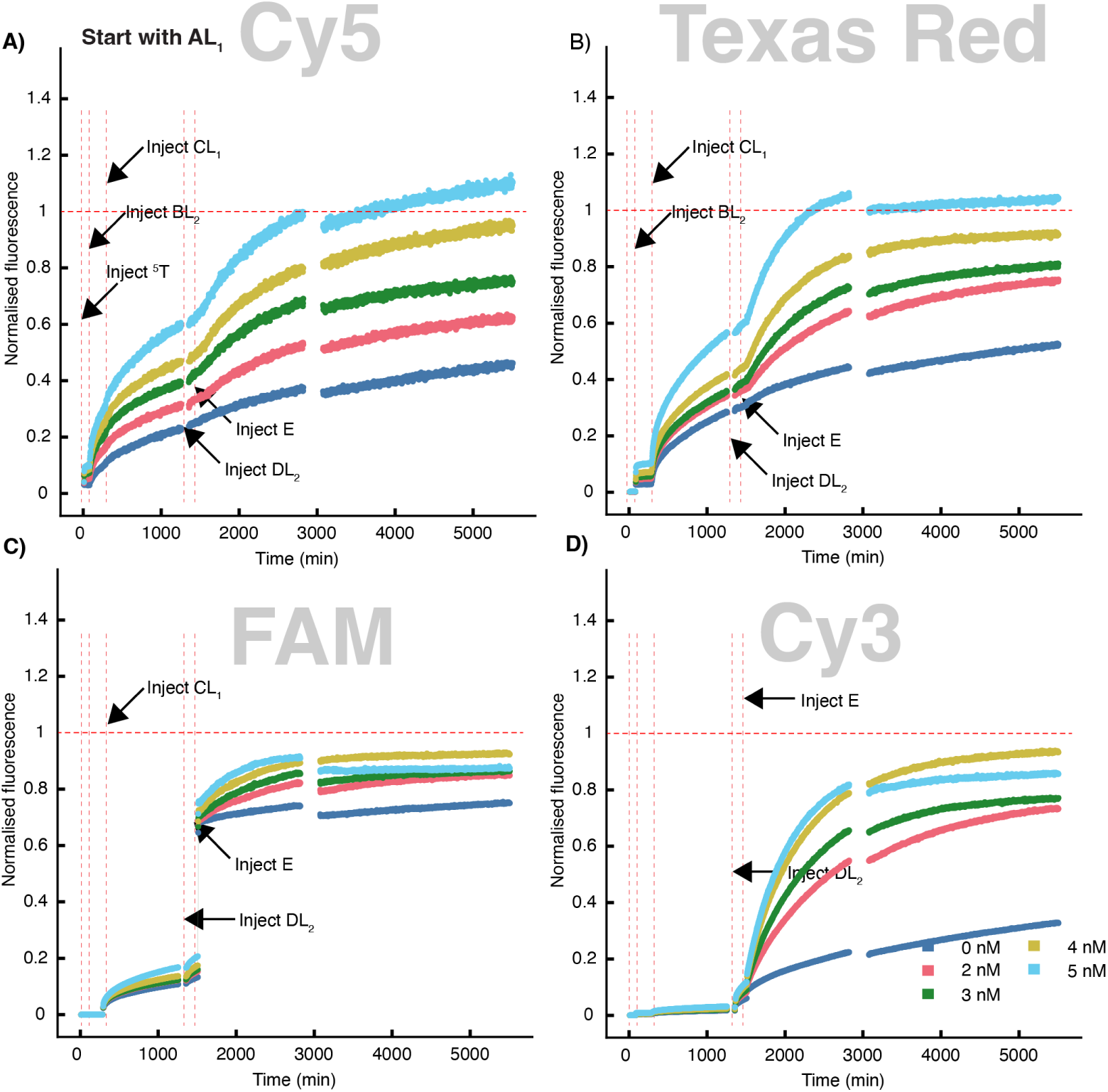
Stepwise monomer addition at high concentration in pentamer formation reaction. Fluorescence arising from monomer turnover during catalytic trimer formation, relative to a 100 nM control (a 1:1:1 mixture of *AB*, *DE*, and *C*) as decribed in the methods. 2-5 nM of ^5^*T* was added to 100 nM blocked monomer *AL*_1_. Then 100 nM of *BL*_2_, *CL*_1_, *DL*_2_, and *E* were added in a stepwise fashion, one monomer at a time. Fluorescence reported in the channels corresponds to *A* (in **A**), *B* (in **B**), *C* & *E* (in **C**), and *D* (in **D**).

**Figure S23:**
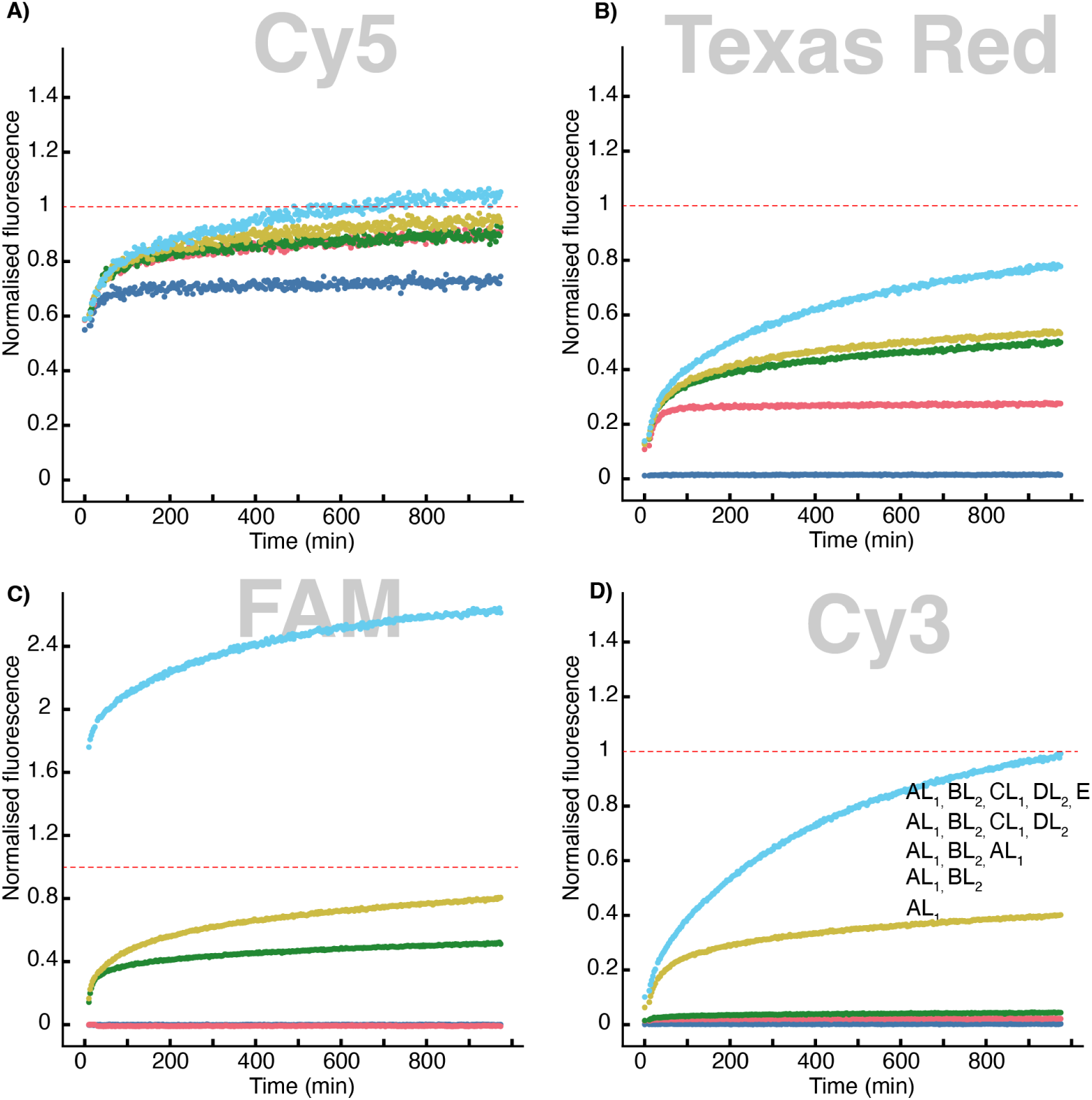
Kinetics of reactions with increasing number of monomers. Fluorescence arising from monomer turnover during catalytic pentamer formation, normalised relative to 10 nM controls (*A*^5^*T* / *B*^5^*T* / *C*^5^*T* / *D*^5^*T*) as outlined in the methods. 4 nM of ^5^*T* was added to the reaction wells containing different numbers of types of 10 nM blocked monomers. First reaction contained 10 nM of *AL*_1_; second reaction contained 10 nM of *AL*_1_ and *BL*_2_; third reaction contained 10 nM of *AL*_1_, *BL*_2_, and *CL*_1_; fourth reaction contained 10 nM of *AL*_1_, *BL*_2_, *CL*_1_, and *DL*_2_; and the last well had 10 nM of *AL*_1_, *BL*_2_, *CL*_1_, *DL*_2_ and *E*. Fluorescence is reported in the channels corresponding to *A* (in **A**), *B* (in **B**), *C* & *E* (in **C**), and *D* (in **D**).

**Figure S24:**
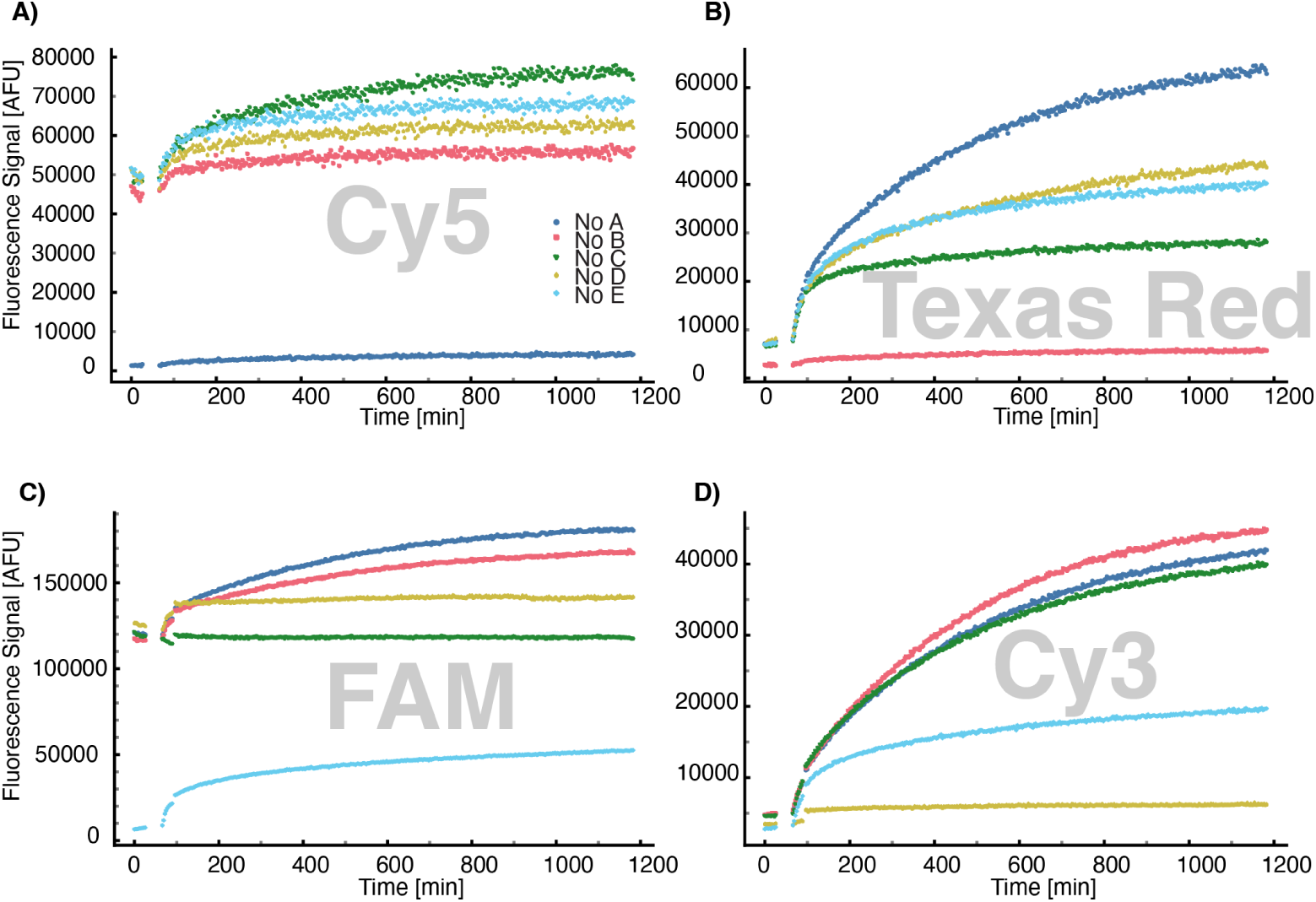
Pentamer templating reactions with systemic omission of monomers. Unprocessed fluorescence arising from monomer unblocking during systematic omission experiment. 4 nM of ^5^*T* was added to reaction wells containing combinations of 10 nM blocked monomers. First reaction contained 10 nM of monomers *BL*_2_, *CL*_1_, *DL*_2_, and *E*, but no *AL*_1_; second reaction contained 10 nM of all monomers except *BL*_2_; third reaction contained 10 nM of all monomers except *CL*_1_; fourth reaction contained 10 nM of all monomers except *DL*_2_; and fifth reaction contained 10 nM of all monomers except *E*. Fluorescence was reported in the channels corresponding to *A* (in **A**), *B* (in **B**), *C* + *E* (in **C**), and *D* (in **D**).

**Figure S25:**
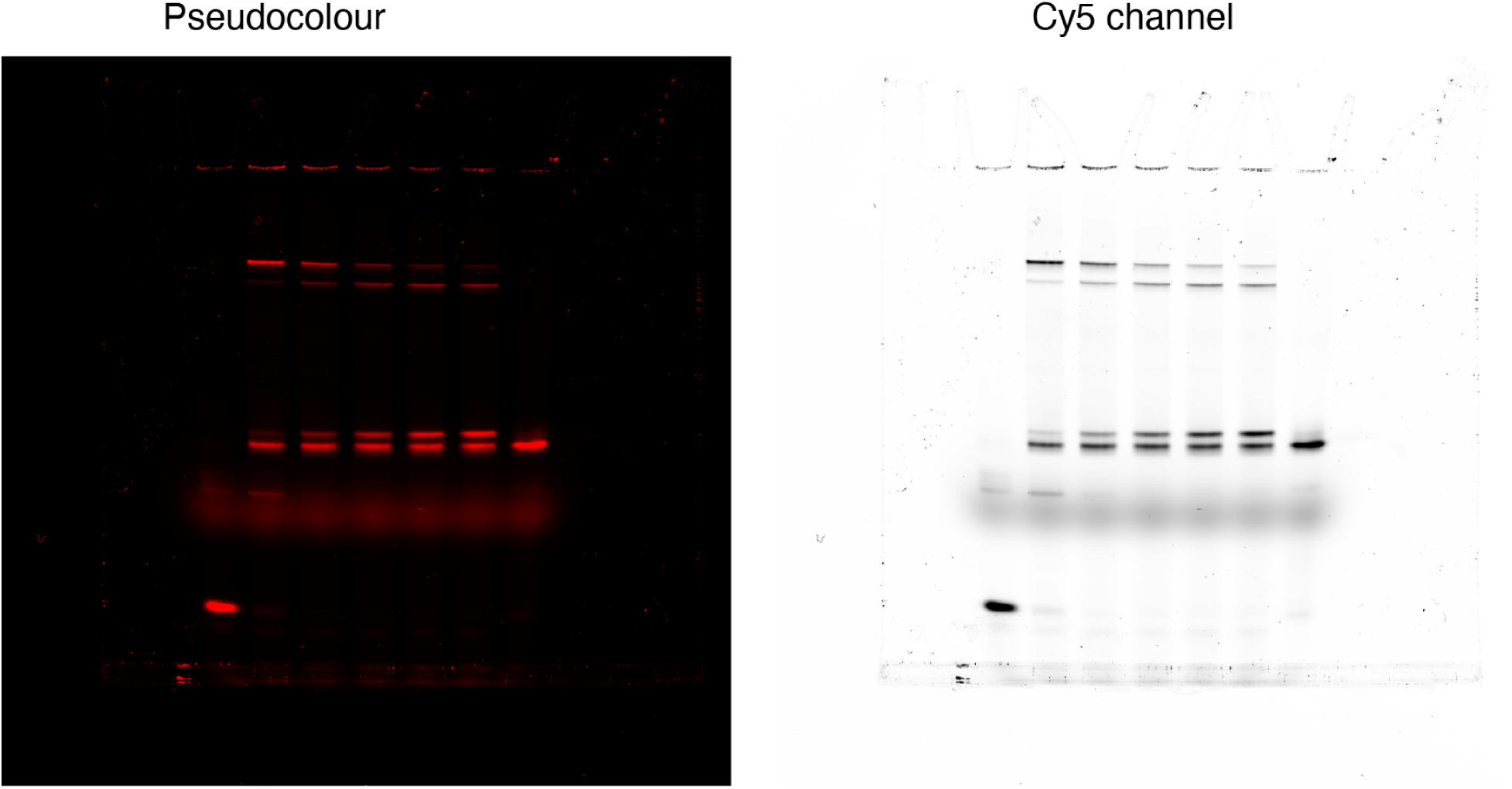
Original fluorescence gel images of *ABC_s_* as shown in 2C. From left to right: pseudocolour image of trimers formation experiments with standard monomer concentrations, and greyscale image scanned in Cy5 channel. In the lanes from left to right, we had monomer mix containing 10 nM of (*AL*_1_, *BL*_2_, and *C_s_*) with 0, 2, 4, 6, 8, and 10 nM *T_ABCs_*. Lane 7 had 10 nM *ABC_s_* as control. The greyscale image has manually adjusted brightness and contrast for better legibility.

**Figure S26:**
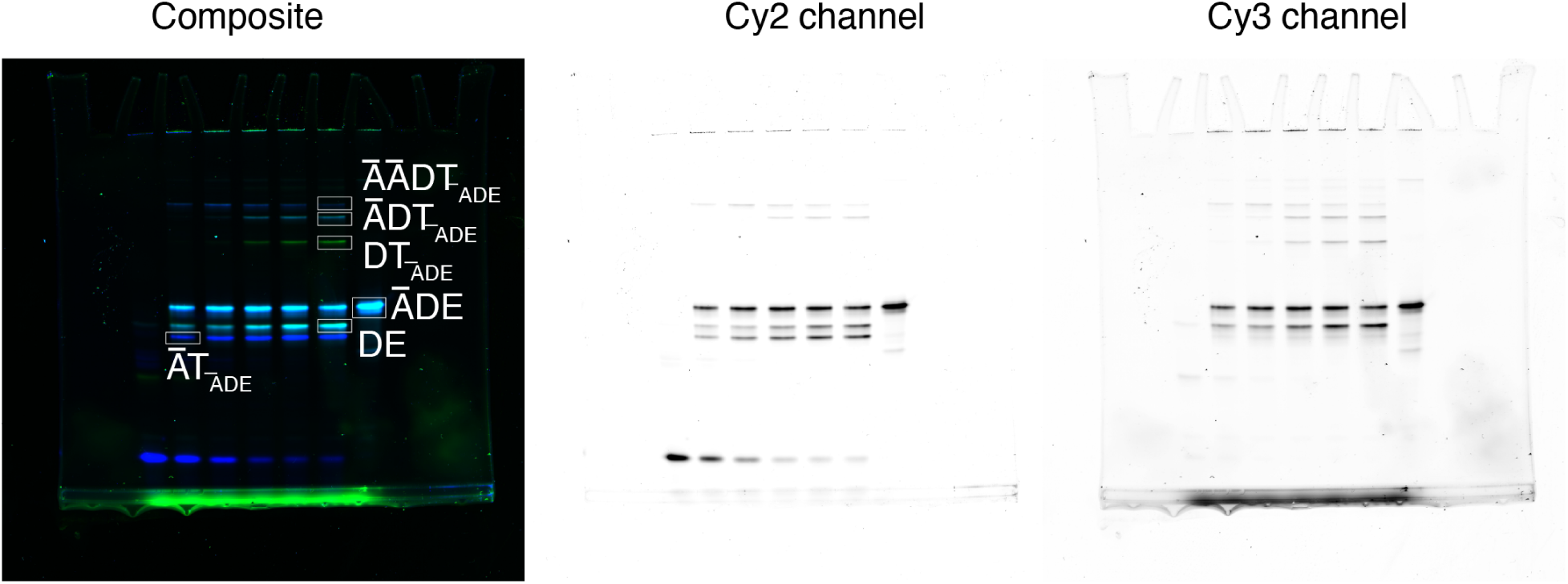
Original fluorescence gel images of *ĀDE* as shown in 2C. From left to right: composite image of trimers-formation experiments with standard monomer concentrations, greyscale image scanned in Cy2 channel, and in Cy3 channel. In the lanes from left to right, we had monomer mix containing 10 nM of (*ĀL*_1_, *DL*_2_, and *E*) with 0, 2, 4, 6, 8, and 10 nM *T_ĀDE_*. Lane 7 had 10 nM *ĀDE* as control. The greyscale images have manually adjusted brightness and contrast for better legibility.

**Figure S27:**
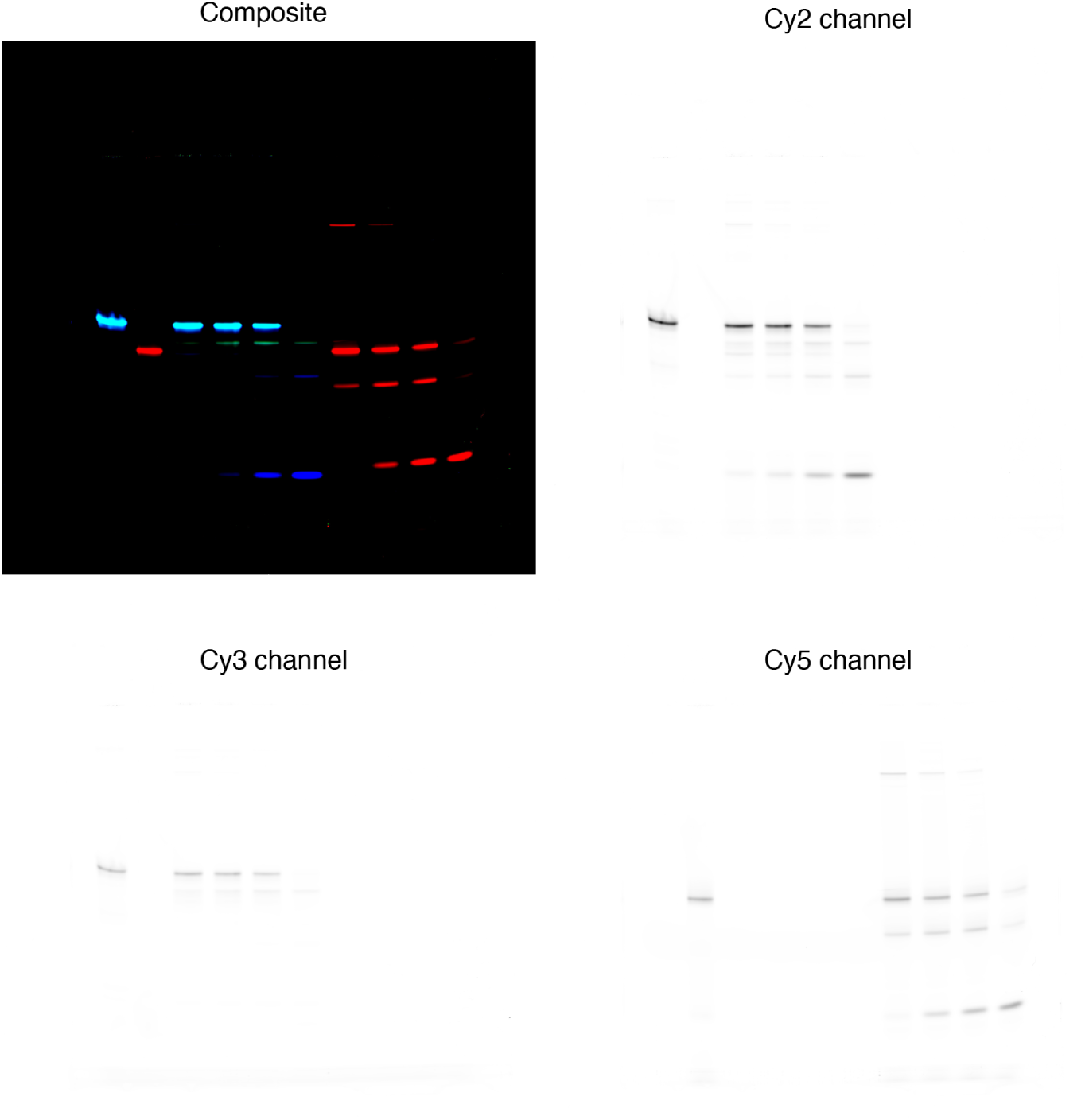
Original fluorescence gel images of both trimers formed from monomers at high concentration as shown in 3B. Clockwise from top left corner: composite image of trimer-formation experiments with high monomer concentrations, greyscale image scanned in Cy2 channel, in Cy5 channel, and in Cy3 channel. Lanes 1 and 2 from left had 100 nM *ĀDE* and *ABC_s_* as controls. In the lanes 3-6, we had monomer mix containing 100 nM of (*ĀL*_1_, *DL*_2_, and *E*) with 5, 3, 2, and 0 nM *T_Ā_*_*DE*_. Lane 7-10 had monomer mix containing 100 nM of (*AL*_1_, *BL*_2_, and *C_s_*) with 5, 3, 2, and 0 nM *T_ABCs_*. The greyscale images have manually adjusted brightness and contrast for better legibility.

**Figure S28:**
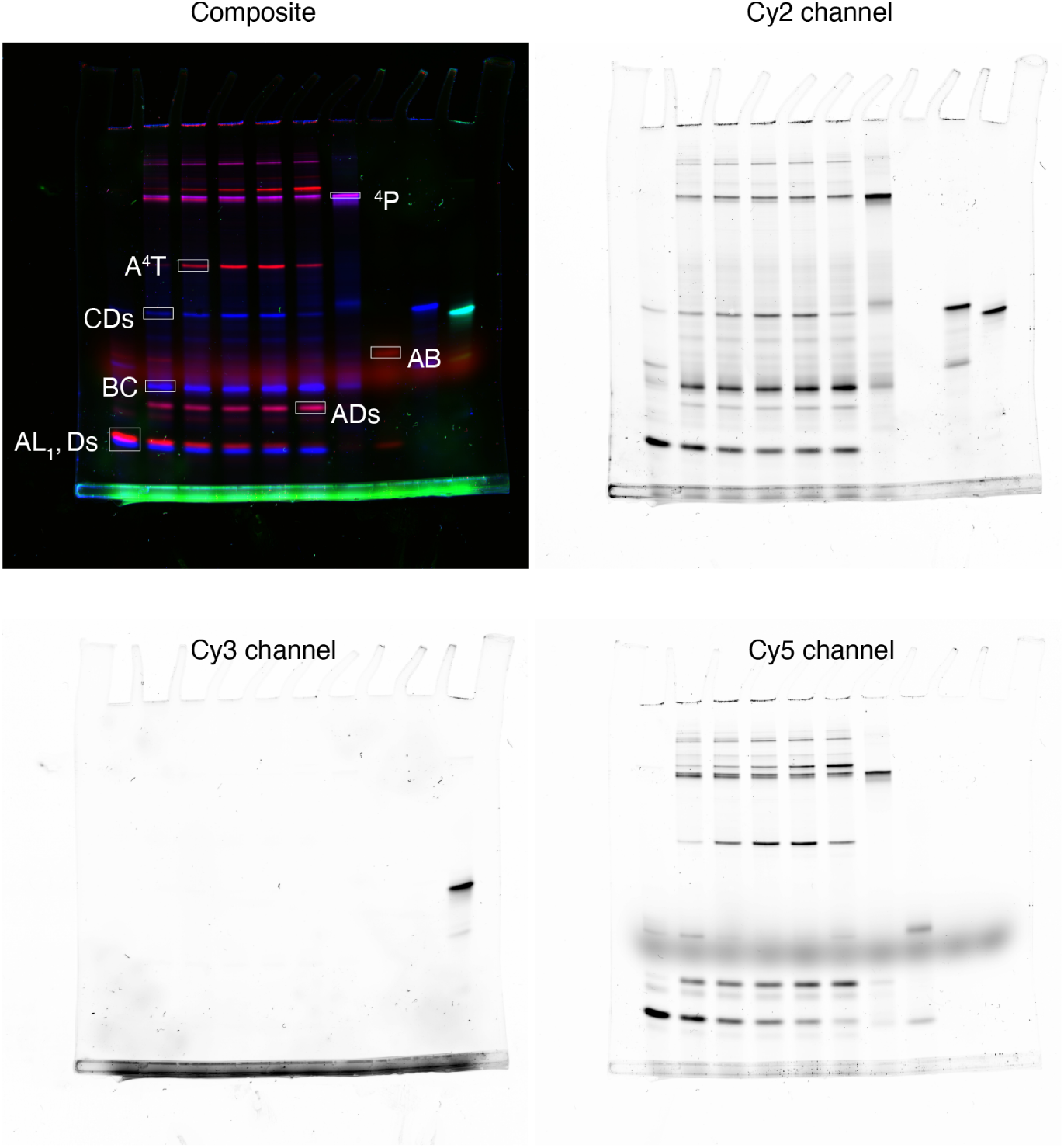
Fluorescence gel images of tetramer-formation experiment. Clockwise from top left corner: composite image of tetramer formation experiments, greyscale image scanned in Cy2 channel, in Cy5 channel, and in Cy3 channel. In the lanes from left to right, we have monomer mix containing 10 nM of (*AL*_1_, *BL*_2_, *CL*_1_, *D_s_*) with 0, 2, 4, 6, 8 and 10 nM ^4^*T*. Lanes 7-10 had 10 nM *AB* and *CD_s_* (for assembling ^4^ *P*), *AB*, *CD_s_*, and *DE* (not relevant for this experiment) controls. The greyscale images have manually adjusted brightness and contrast for better legibility.

**Figure S29:**
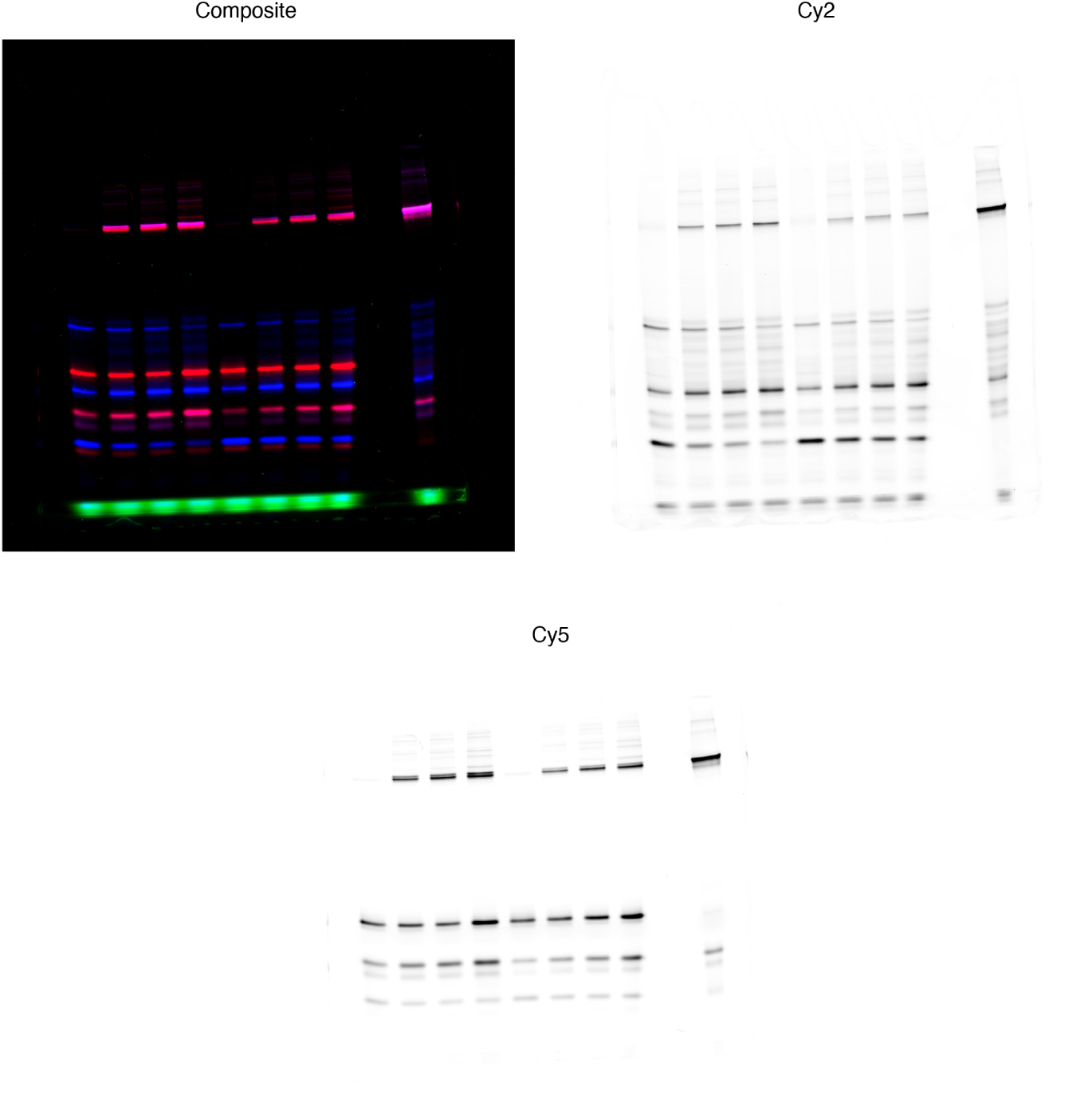
Fluorescence gel images of the tetramer formation reaction at high monomer concentration. Clockwise from top left corner: composite image of tetramer formation at high monomer concentrations, greyscale image scanned in Cy2 channel, and Cy5 channel channel. In the lanes from left to right, we have monomer mix containing 100 nM of (*AL*_1_, *BL*_2_, *CL*_1_, *D_s_*) with 0, 2, 3, and 5 nM ^4^*T*. Lane 10 had 100 nM of each *AB* and *CD_s_*(for assembling ^4^ *P*) control. The greyscale images have manually adjusted brightness and contrast for better legibility.

**Figure S30:**
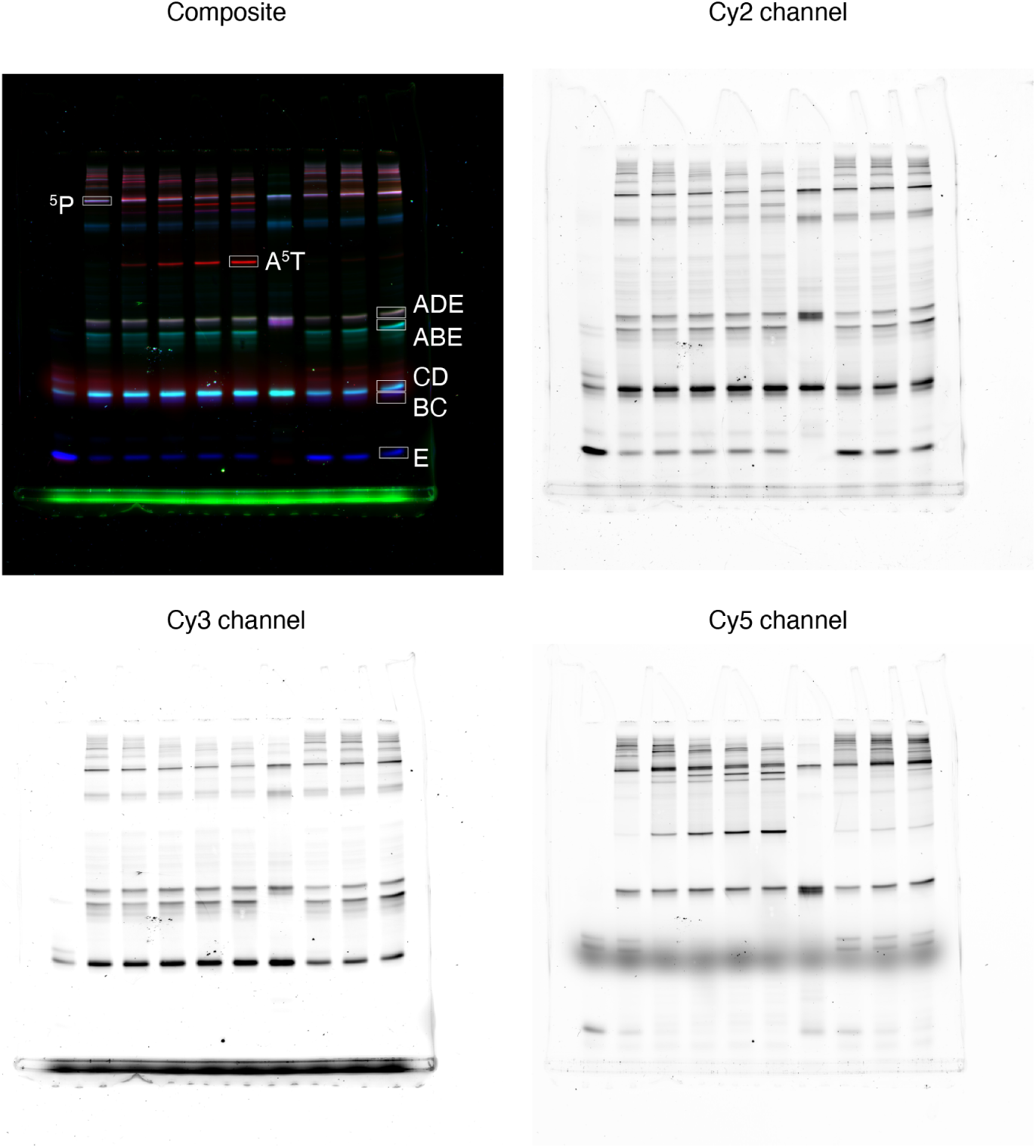
Original fluorescence gel images of pentamer formation reaction as shown in Figure 4E. In the main text, 7 lanes from the left are displayed. Clockwise from top left corner: composite image of pentamer formation experiments, greyscale image scanned in Cy2 channel, in Cy3 channel, and in Cy5 channel. In the lanes from left to right, we have monomer mix containing 10 nM of (*AL*_1_, *BL*_2_, *CL*_1_, *DL*_2_, *E*) with 0, 2, 4, 6, 8 and 10 nM ^5^*T_Ē_*. Lane 7 had 10 nM of each *AB*, *DE*, and *C* (for assembling ^5^ *P*) control. The greyscale images have manually adjusted brightness and contrast for better legibility.

**Figure S31:**
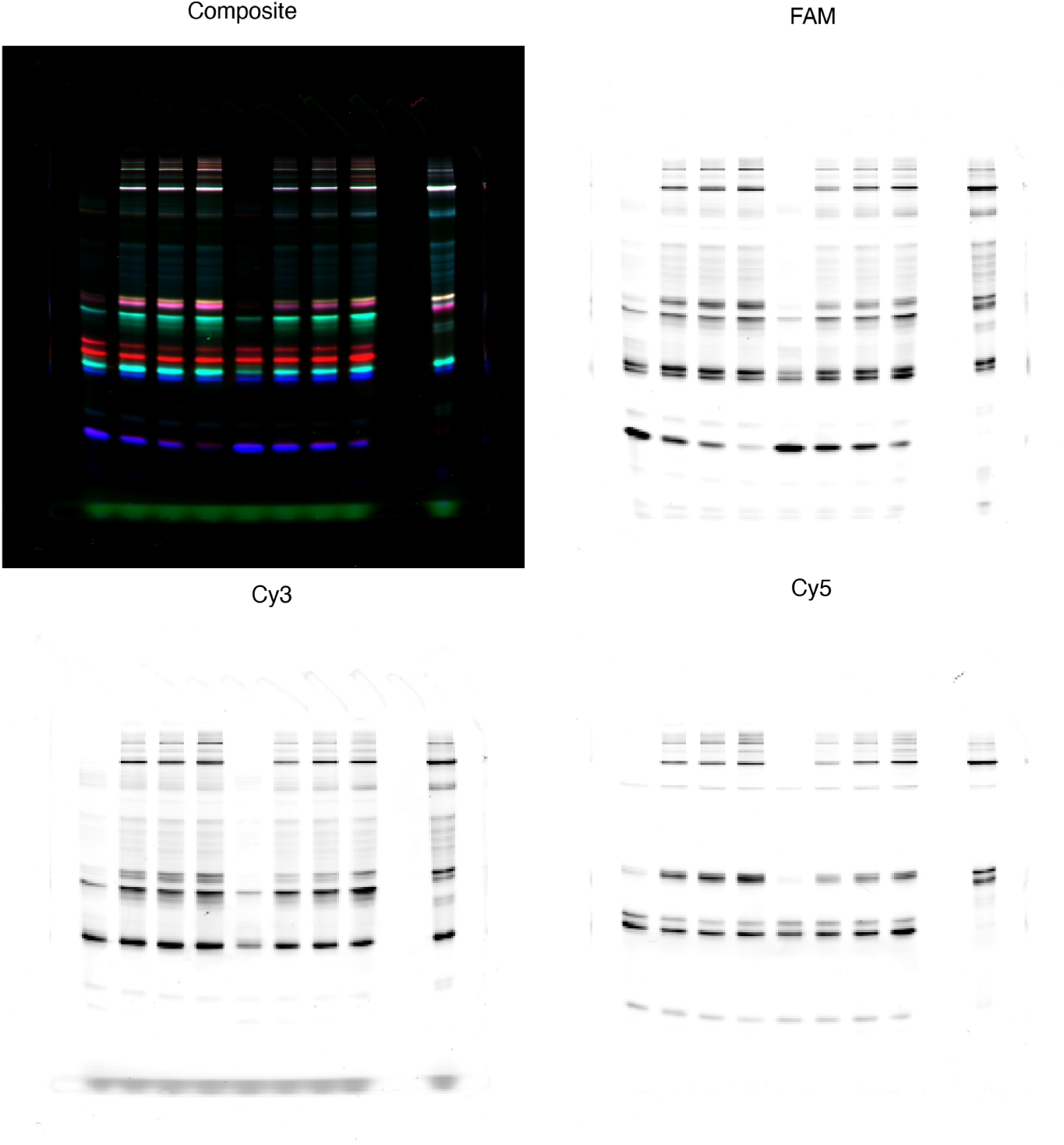
Original fluorescence gel images of pentamer formation reaction at high monomer concentration as shown in Figure 4F. Lanes 1-4 and 10 from the left are displayed in the main text. Clockwise from top left corner: composite image of pentamer formation experiments at high monomer concentration, greyscale image scanned in Cy2 channel, in Cy3 channel, and in Cy5 channel. In the lanes from left to right, we have monomer mix containing 100 nM of (*AL*_1_, *BL*_2_, *CL*_1_, *DL*_2_, *E*) with 0, 2, 3, and 5 nM ^5^*T*. The rightmost lane had 100 nM of each *AB*, *DE*, and *C* (for assembling ^5^ *P*). The greyscale images have manually adjusted brightness and contrast for better legibility.

**Figure S32:**
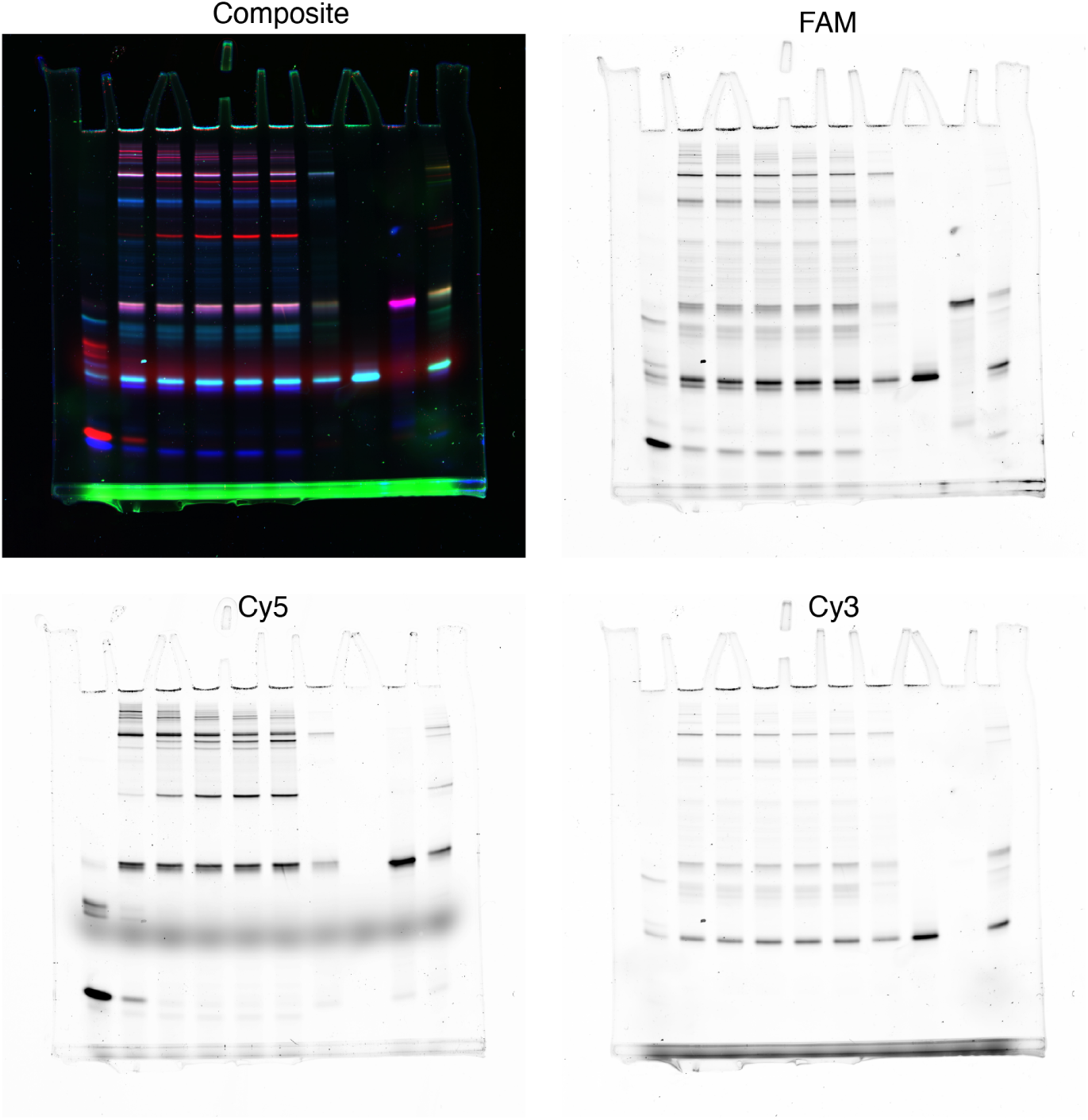
Fluorescence gel images of an alternate pentamer formation reaction. Clockwise from top left corner: composite image of pentamer formation experiments, greyscale image scanned in Cy2 channel, in Cy5 channel, and in Cy3 channel. In the lanes from left to right, we have monomer mix containing 10 nM of (*AL*_1_, *BL*_2_, *CL*_1_, *DL*_2_, *Ē*) with 0, 2, 4, 6, 8 and 10 nM ^5^*T_Ē_*. Lanes 7-10 had 10 nM of each *AB*, *DĒ*, and *C* (for assembling ^5^ *P_Ē_*), *CD*, *ABĒ*, and *ADĒ* controls. The greyscale images have manually adjusted brightness and contrast for better legibility.

**Figure S33:**
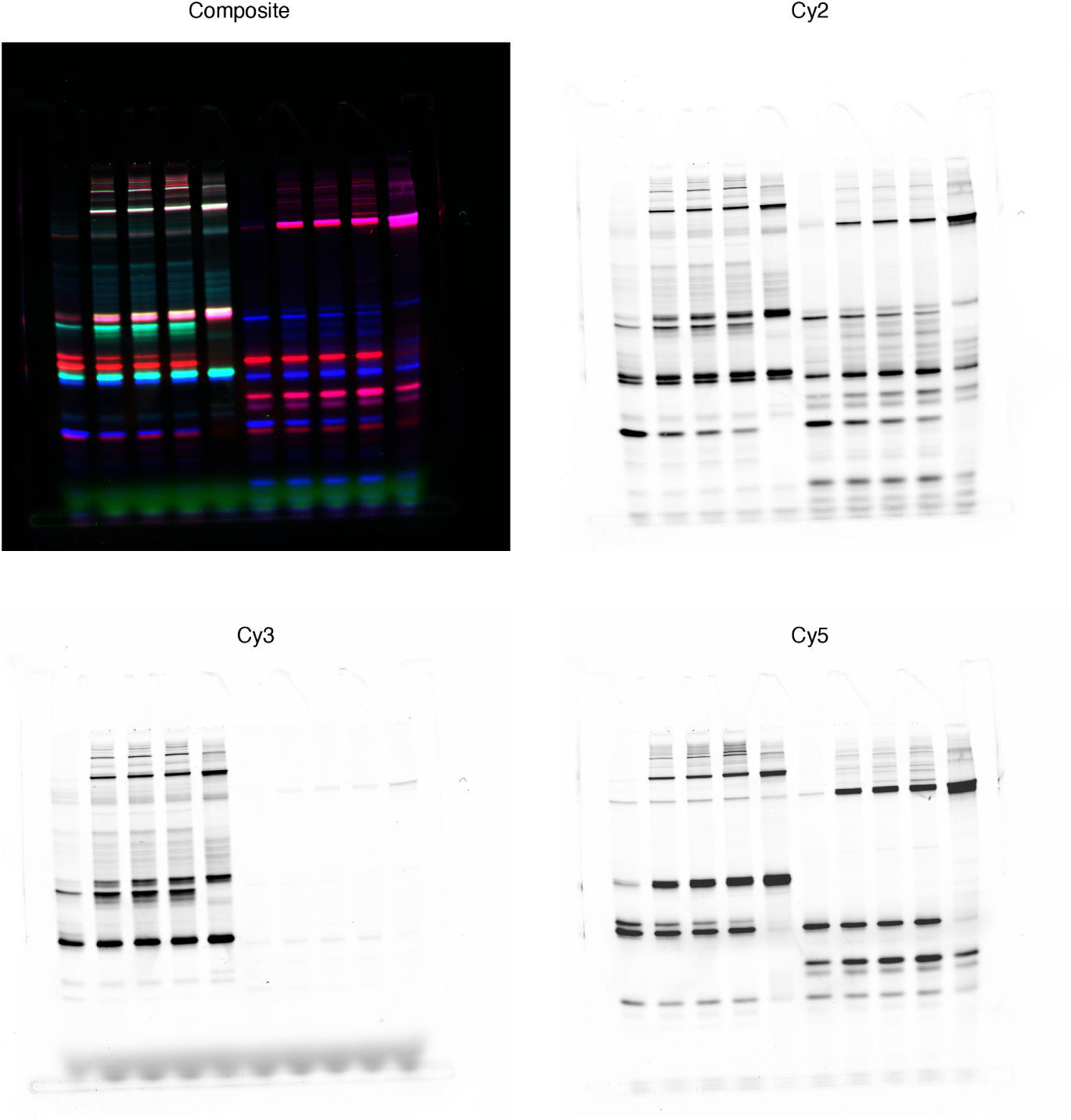
Fluorescence gel images of an alternate pentamer formation reaction at high monomer concentration. Clockwise from top left corner: composite image of pentamer formation experiments at high monomer concentration, greyscale image scanned in Cy2 channel, in Cy5 channel, and in Cy3 channel. In the lanes from left to right, we have monomer mix containing 100 nM of (*AL*_1_, *BL*_2_, *CL*_1_, *DL*_2_, *Ē*) with 0, 2, 3, and 5 nM ^5^*T_Ē_*. Lane 5 has 100 nM of each *AB*, *DĒ*, and *C* (for assembling ^5^ *P_Ē_*). The greyscale images have manually adjusted brightness and contrast for better legibility.

**Figure S34:**
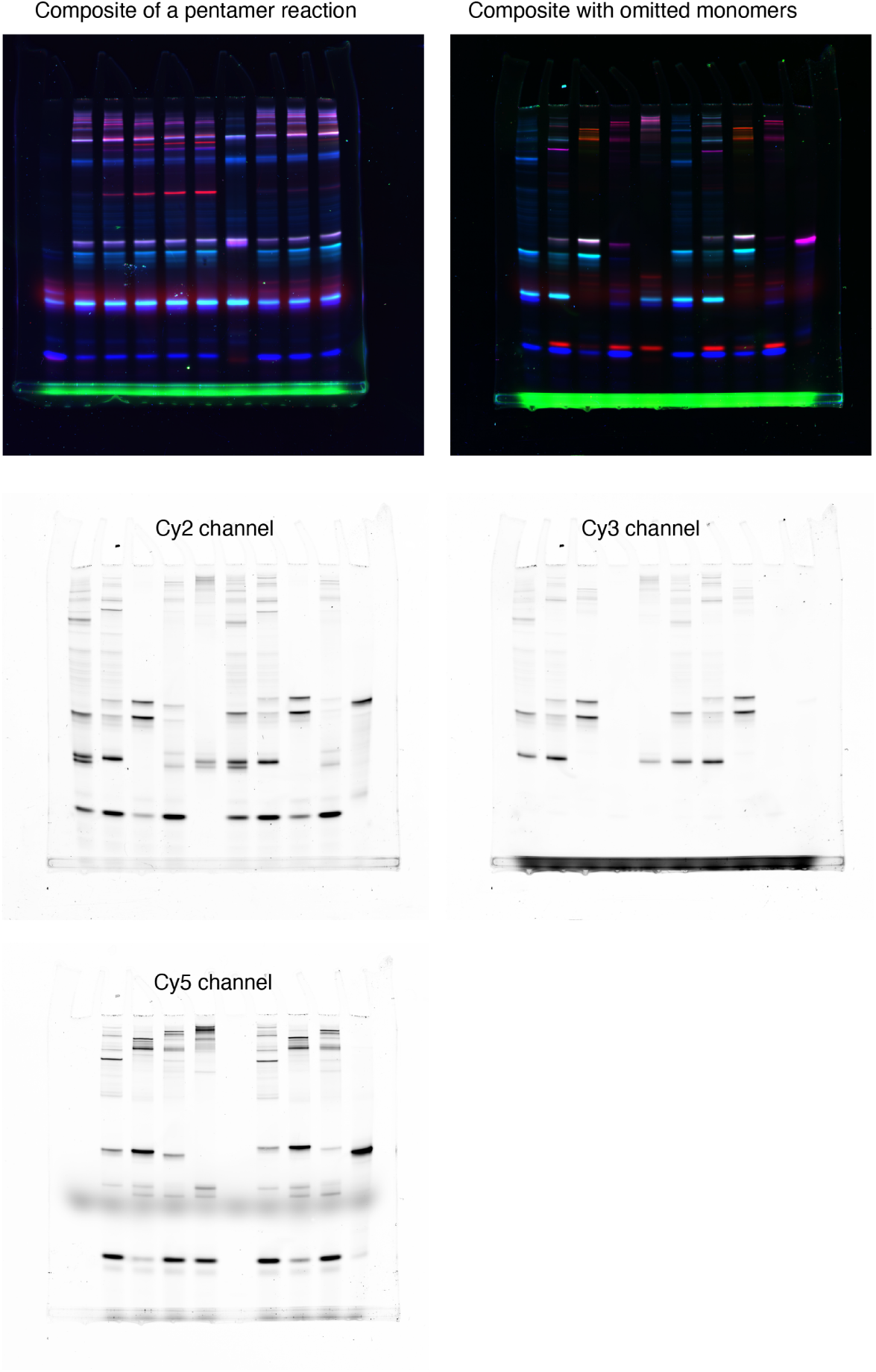
Fluorescence gel images of reaction where each monomer is systematically missing. Clockwise from top left corner: pseudocolour composite image of a normal pentamer formation as shown in figure S30 for comparison, composite image of the systematic omission experiments with ^5^*T* and ^5^*T_Ē_*, greyscale image scanned in Cy3 channel, in Cy5 channel, and in Cy2 channel. In the first five lanes of the top right gel from left to right, we have monomer mix containing 10 nM of (*BL*_2_, *CL*_1_, *DL*_2_, *E*), (*AL*_1_, *CL*_1_, *DL*_2_, *E*), (*AL*_1_, *BL*_2_, *DL*_2_, *E*), (*AL*_1_, *BL*_2_, *CL*_1_, *E*), and (*AL*_1_, *BL*_2_, *CL*_1_, *DL*_2_). All reactions were triggered with 4 nM ^5^*T*. In the lanes 6-9, from left to right, we have monomer mix containing 10 nM of (*BL*_2_, *CL*_1_, *DL*_2_, *Ē*), (*AL*_1_, *CL*_1_, *DL*_2_, *Ē*), (*AL*_1_, *BL*_2_, *DL*_2_, *Ē*), and (*AL*_1_, *BL*_2_, *CL*_1_, *Ē*). All reactions were triggered with 4 nM ^5^*T_Ē_*. The rightmost lane had annealed mixture of all monomers and blockers at 10 nM. The greyscale images have manually adjusted brightness and contrast for better legibit1y.

**Figure S35:**
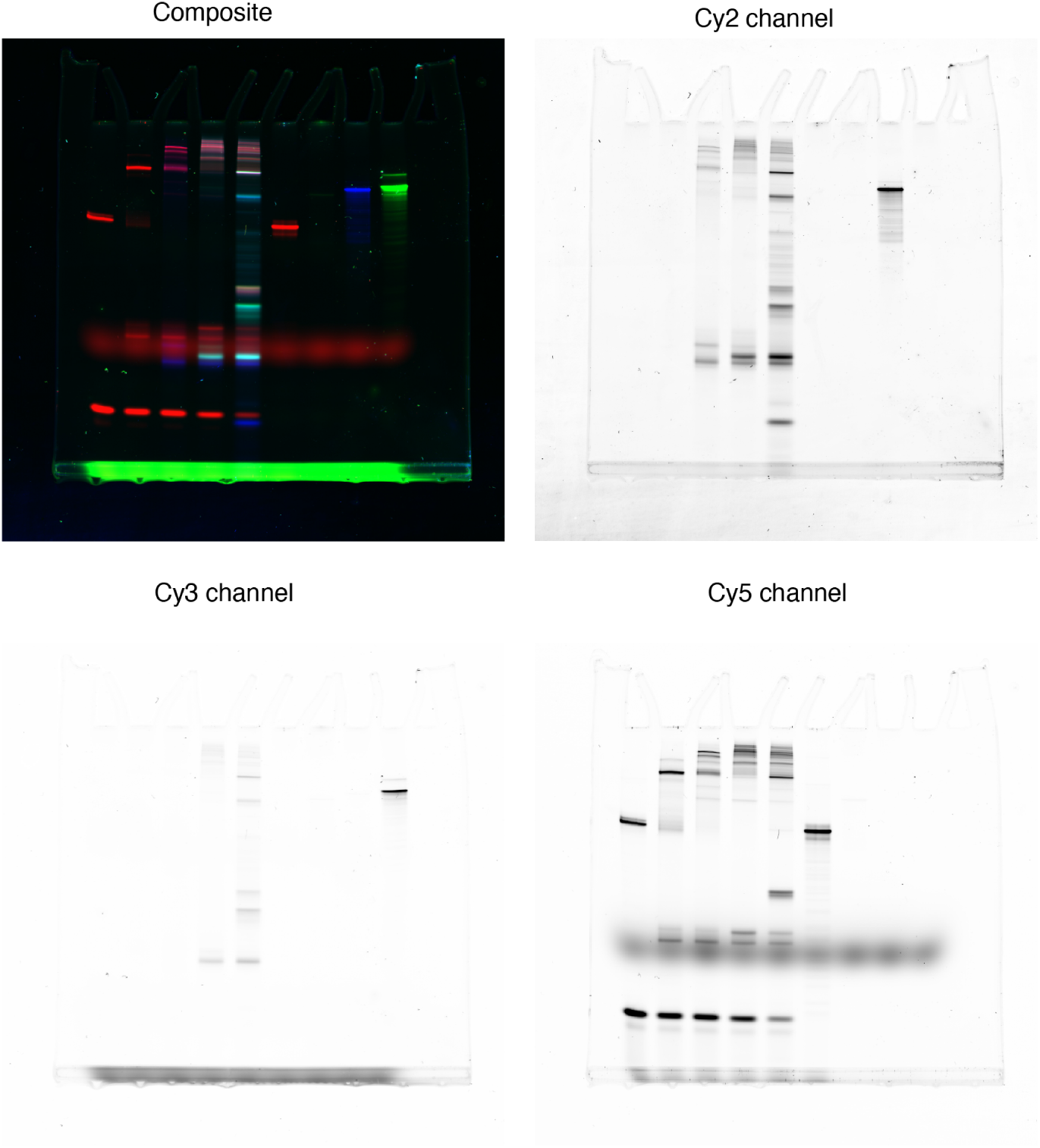
Fluorescence gel images of reaction where one more monomer type is added each time starting from *AL*_1_ only. Clockwise from top left corner: pseudocolour composite image, greyscale image scanned in Cy2 channel, in Cy5 channel, and in Cy3 channel. In the first five lanes from left to right, we have monomer mix containing 10 nM of *AL*_1_; *AL*_1_ + *BL*_2_; *AL*_1_ + *BL*_2_ + *CL*_1_; *AL*_1_ + *BL*_2_ + *CL*_1_ + *DL*_2_; *AL*_1_ + *BL*_2_ + *CL*_1_ + *DL*_2_ + *E*). All reactions were triggered with 4 nM ^5^*T*. Last four lanes had 10 nM of *A*^5^*T*, *B*^5^*T*, *C*^5^*T*, and *D*^5^*T*. The greyscale images have manually adjusted brightness and contrast for better legibility.

**Figure S36:**
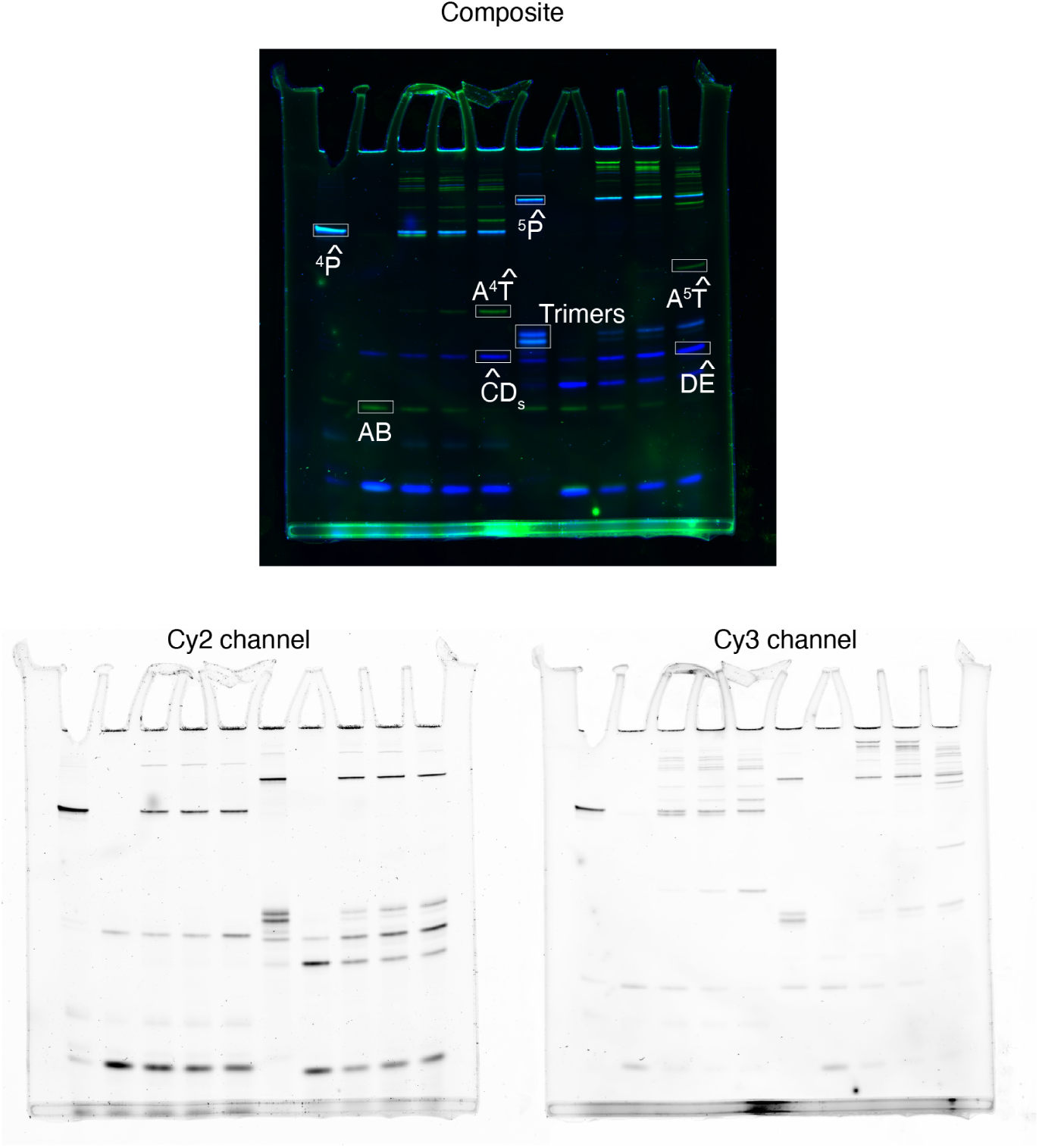
Fluorescence gel images of tetramer and pentamer-formation reactions in which only the terminal monomers are labelled. Clockwise from top: pseudocolour composite image, greyscale image scanned in Cy3 channel, and in Cy2 channel. In the lanes from left to right, we have ^4^ *P̂* control; monomer mix (containing 10 nM of *AL*_1_-Cy3, *BL*_2_-vanilla, *ĈL*_1_-vanilla, and *D_s_*-FAM) with 0 nM, 2 nM, 3 nM, and 5 nM of ^4^*T̂*; ^5^ *P̂* control, monomer mix (containing 10 nM of *AL*_1_-Cy3, *BL*_2_-vanilla, *ĈL*_1_-vanilla, *DL*_2_-vanilla, and *Ê*) with 0 nM, 2 nM, 3 nM, and 5 nM of ^5^*T̂*. The greyscale images have manually adjusted brightness and contrast for better legibility.

**Figure S37:**
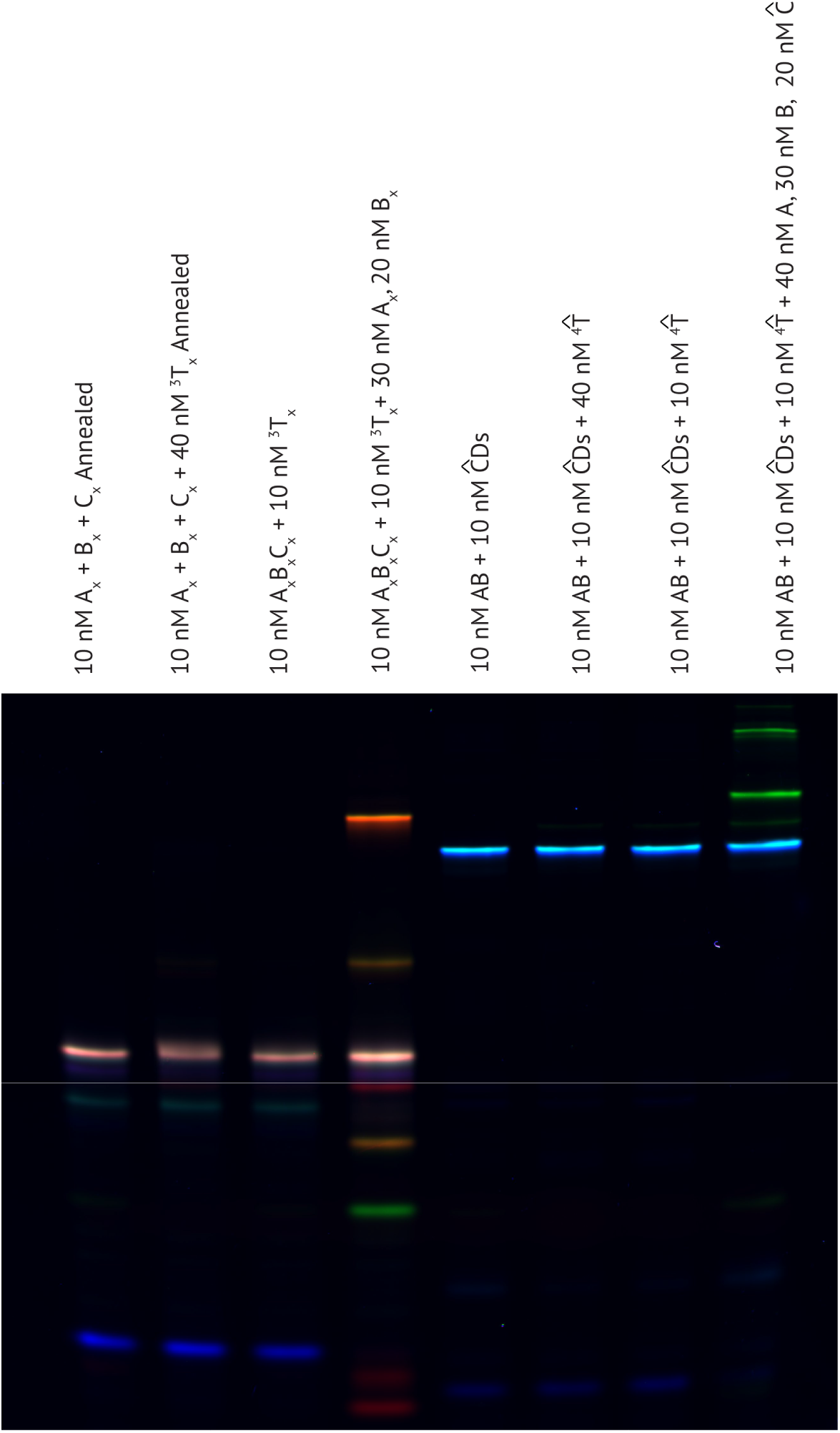
Fluorescence gel images of reactions testing the binding propensity of the templates to products. Different concentrations of monomers and templates were annealed or incubated to test if the templates form a sufficiently stable complex with the fully formed trimeric or tetrameric products. We do not observe such behaviour in either case, even in presence of large excess of the templates evidenced by the absence of any new bands in lanes 2-3 compared to 1, and 6-7 compared to 5 from left. However, with higher monomer concentrations compared to the templates, higher molecular weight ‘‘brushes’’ were formed.

**Figure S38:**
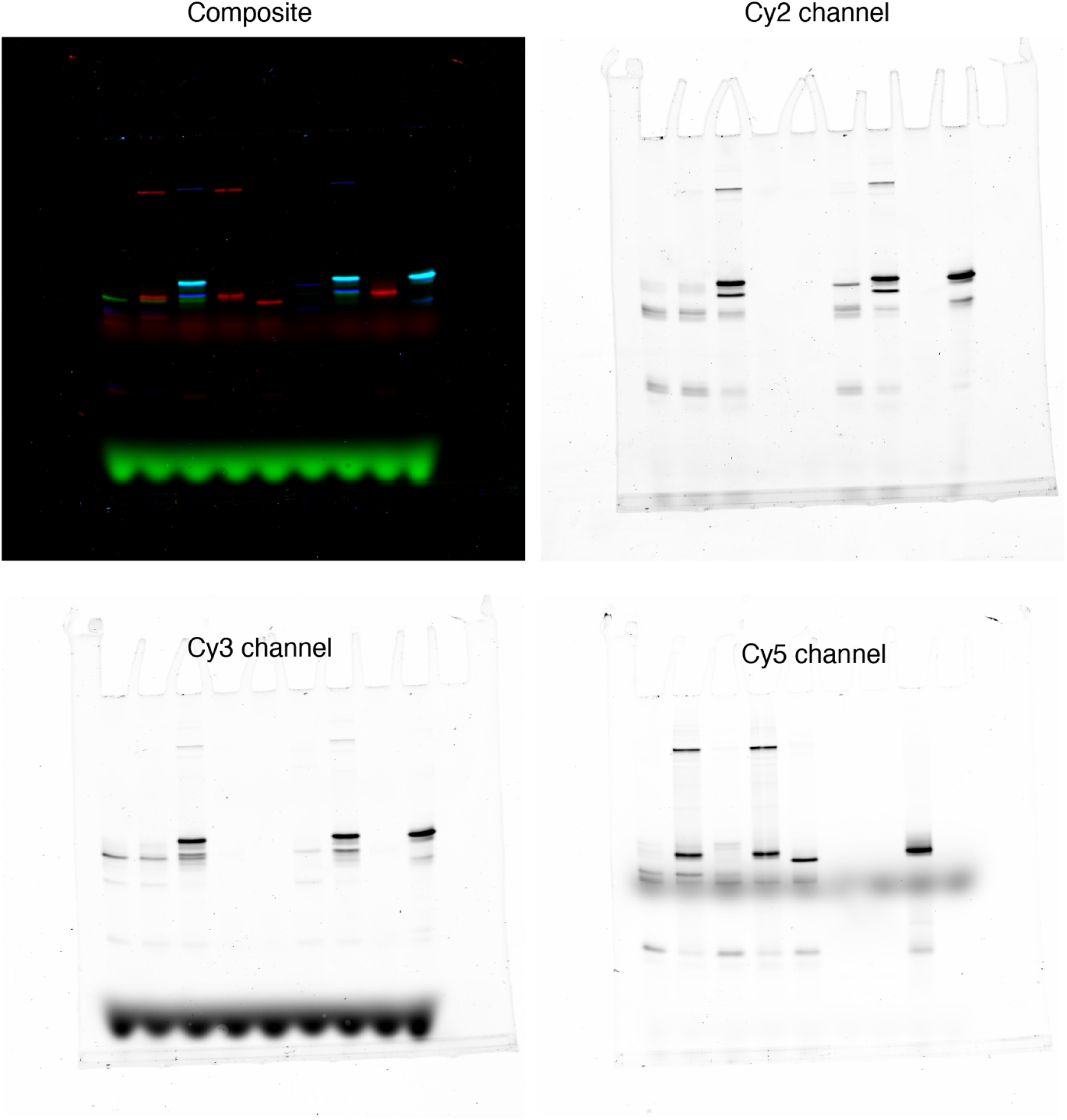
Fluorescence gel images of reactions to test the specificity of trimer templates as shown in Figure 5B. Clockwise from top left corner: pseudocolour composite image, greyscale image scanned in Cy2 channel, in Cy5 channel, and in Cy3 channel. From left to right, in lanes 1-3, we have monomer mix (containing 10 nM of *AL*_1_, *ĀL*_1_, *BL*_2_, *DL*_2_, *C_s_*, and *E*–vanilla) with no template, 2 nM T_ABCs_, 2 nM *T_Ā_*_*DE*_. Lanes 4-5 have monomer mix (containing 10 nM of *AL*_1_, *BL*_2_, and *C_s_*) with 2 nM *T_ABCs_*, and 2 nM *T_Ā_*_*DE*_. Lanes 6-7 have monomer mix (containing 10 nM of *ĀL*_1_, *DL*_2_, and *E*) with 2 nM *T_ABCs_*, and 2 nM *T_ĀDE_*. Lanes 8-9 have 10 nM *ABC_s_* and *ĀDE*. The greyscale images have manually adjusted brightness and contrast for better legibility.

**Figure S39:**
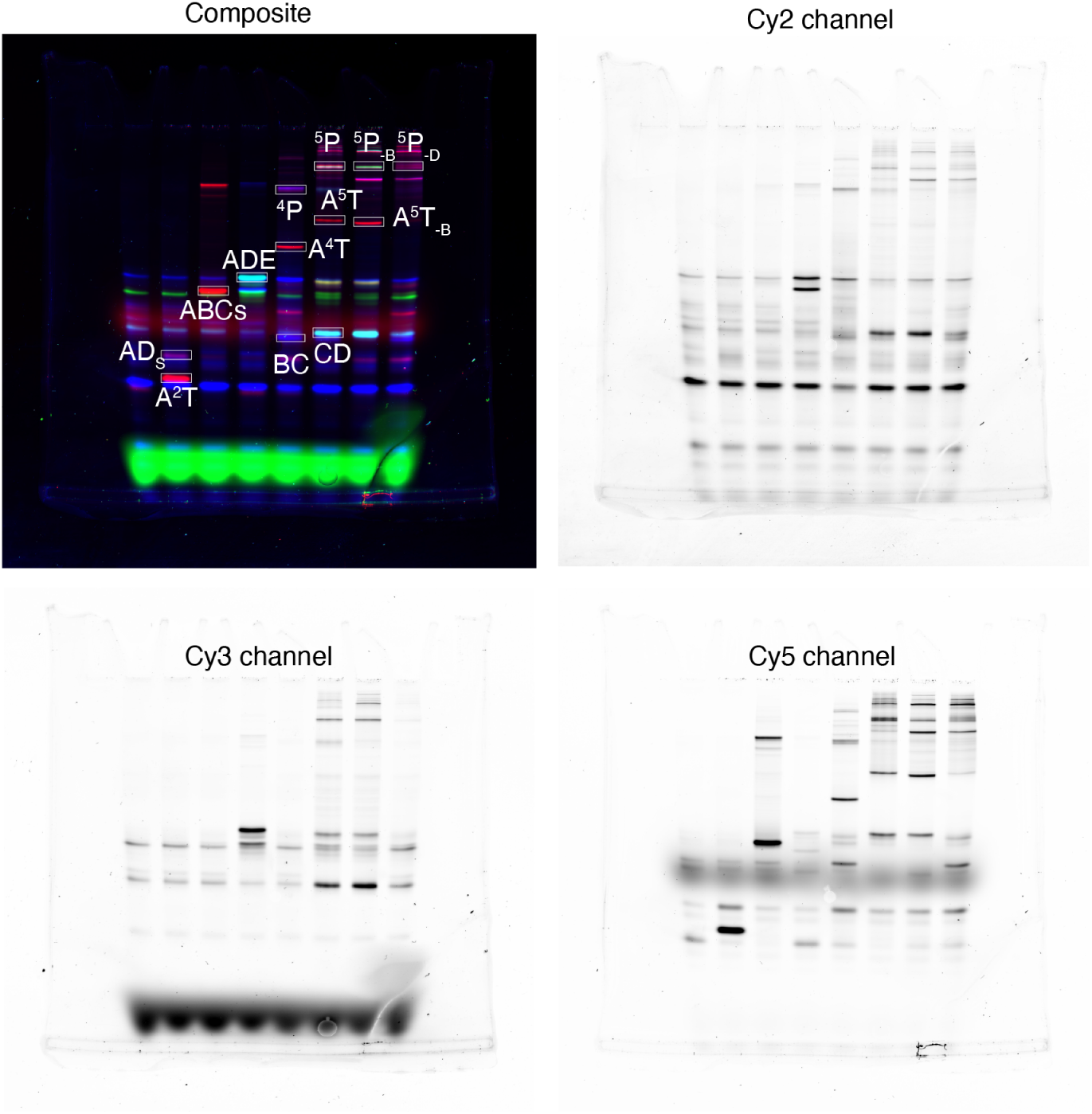
Fluorescence gel images of reactions to test the global specificity of catalytic templates. Clockwise from top left corner: pseudocolour composite image, greyscale image scanned in Cy2 channel, in Cy5 channel, and in Cy3 channel. In the lanes from left to right, we have monomer mix (containing 10 nM of *AL*_1_, *ĀL*_1_, *BL*_2_, *CL*_1_, *DL*_2_, *C_s_*, *D_s_*, and *E*) with the following triggers: no template, 2 nM *T_AD_*, 2 nM *T_ABCs_*, 2 nM *T_Ā_*_*DE*_, 2 nM ^4^*T*, 2 nM ^5^*T*, 2 nM ^5^*T*_−*B*_, and 2 nM ^5^*T*_−*D*_. The greyscale images have manually adjusted brightness and contrast for better legibility.

**Figure S40:**
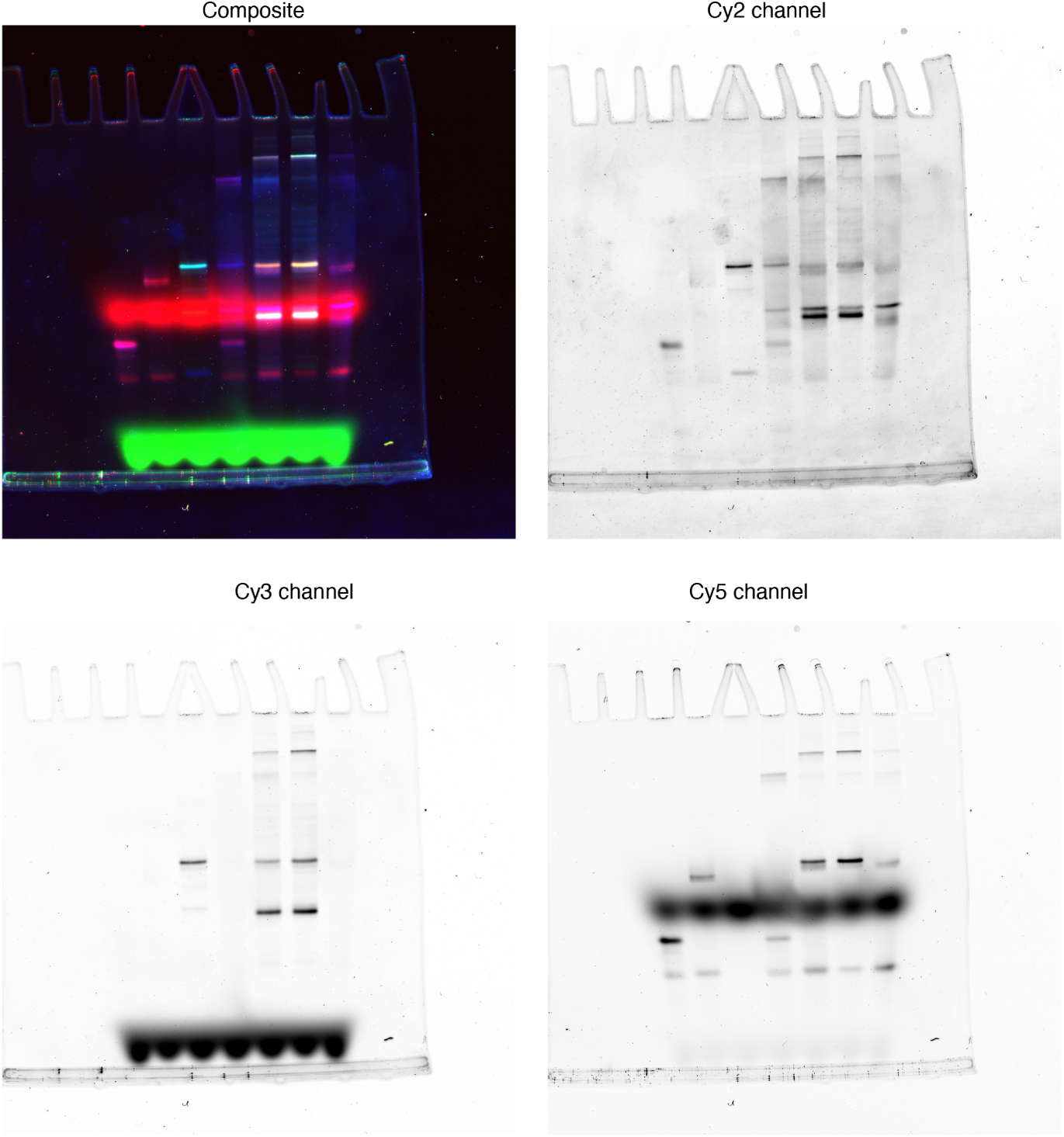
Fluorescence gel images of pre-made controls for the global specificity experiments. Clockwise from top left corner: pseudocolour composite image, greyscale image scanned in Cy2 channel, in Cy5 channel, and in Cy3 channel. In the lanes from left to right, we have 10 nM *AD_s_*; *ABC_s_*; *ĀDE*; *AB* and *CD_s_* (to assemble ^4^ *P*); *AB*, *DE*, and *C* (to assemble ^5^ *P*); *AD*, *DE*, and *C* (to assemble ^5^ *P*_−*B*_); and *AB*, *BE*, and *C* (to assemble ^5^ *P*_−*D*_). The greyscale images have manually adjusted brightness and contrast for better legibility.

**Figure S41:**
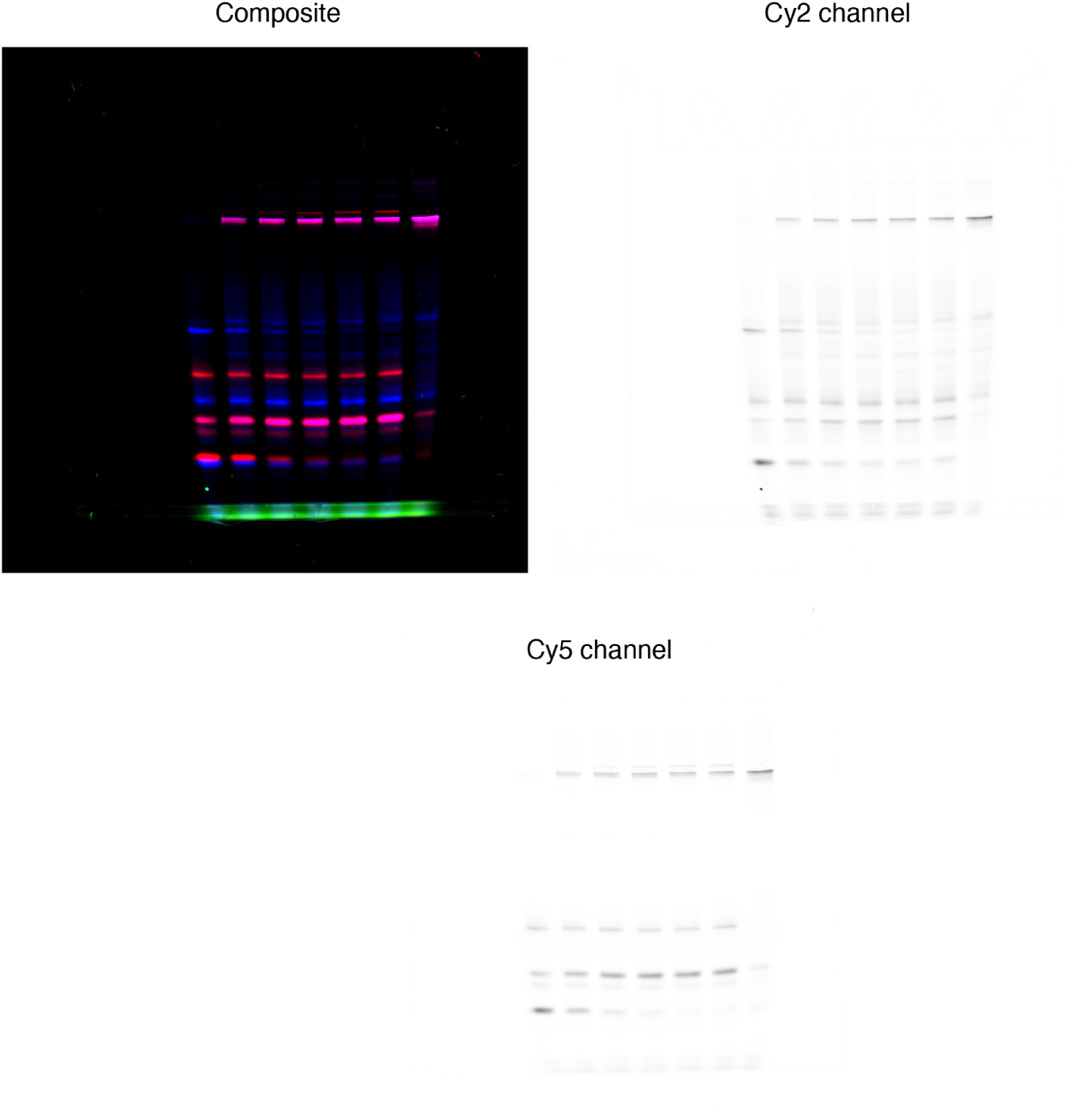
Fluorescence gel images of tetramer formation reaction at high monomer concentration with 0–10 nM template. Clockwise from top left corner: composite image of pentamer formation experiments at high monomer concentration, greyscale image scanned in Cy2 channel, in Cy5 channel, and in Cy3 channel. In the lanes from left to right, we have monomer mix containing 100 nM of (*AL*_1_, *BL*_2_, *CL*_1_, and *D_s_*) with 0, 2, 4, 6, 8, and 10 nM ^4^*T*. The rightmost lane had 100 nM of each *AB*, and *CD_s_* (for assembling ^4^ *P*). The greyscale images have manually adjusted brightness and contrast for better legibility.

**Figure S42:**
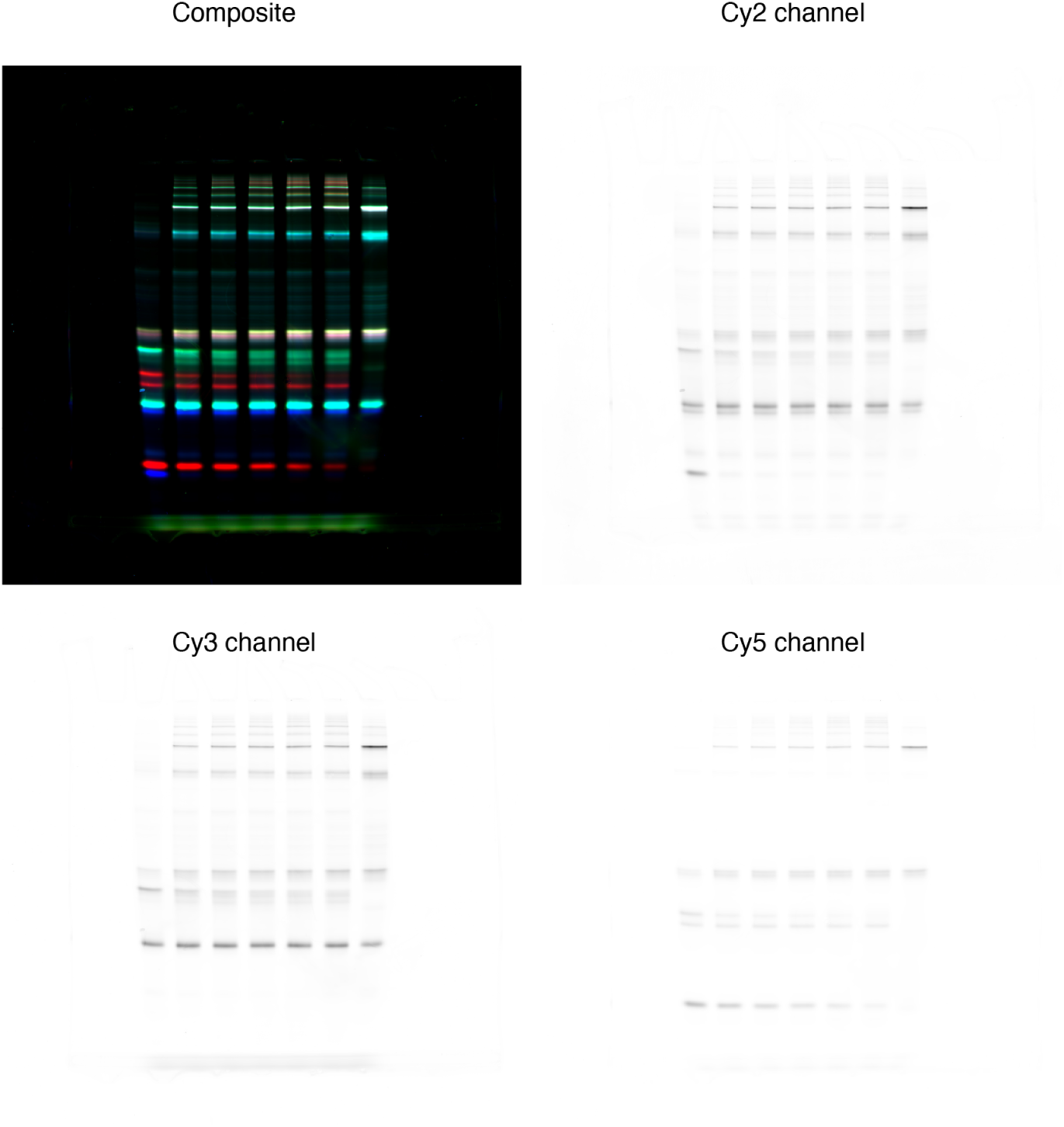
Fluorescence gel images of pentamer formation reaction at high monomer concentration with 0–10 nM template. Clockwise from top left corner: composite image of pentamer formation experiments at high monomer concentration, greyscale image scanned in Cy2 channel, in Cy5 channel, and in Cy3 channel. In the lanes from left to right, we have monomer mix containing 100 nM of (*AL*_1_, *BL*_2_, *CL*_1_, *DL*_2_, *E*) with 0, 2, 4, 6, 8, and 10 nM ^5^*T*. The rightmost lane had 100 nM of each *AB*, *DE*, and *C* (for assembling ^5^ *P*). The greyscale images have manually adjusted brightness and contrast for better legibility.

## Supplementary tables

**Table S1:**
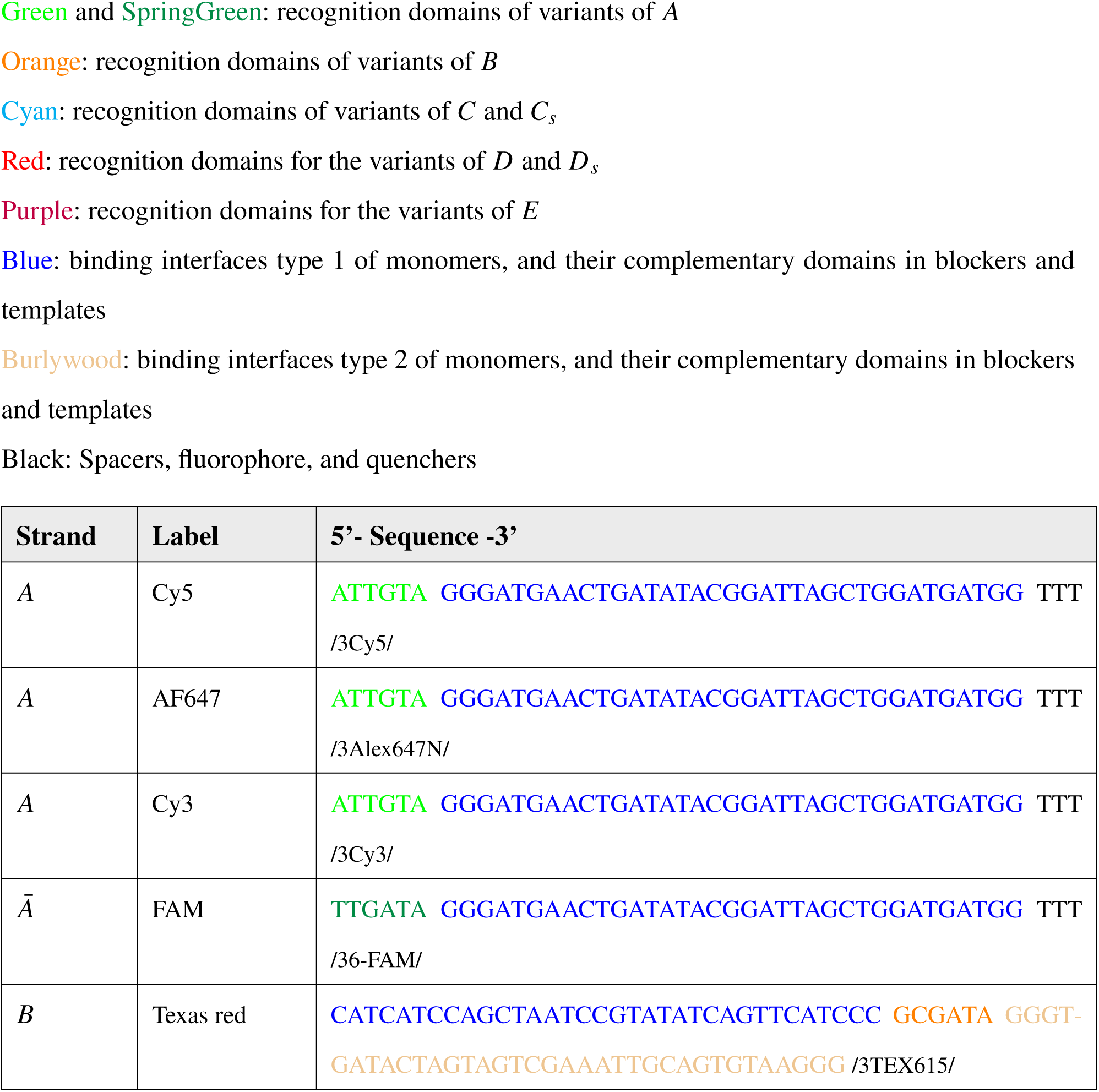
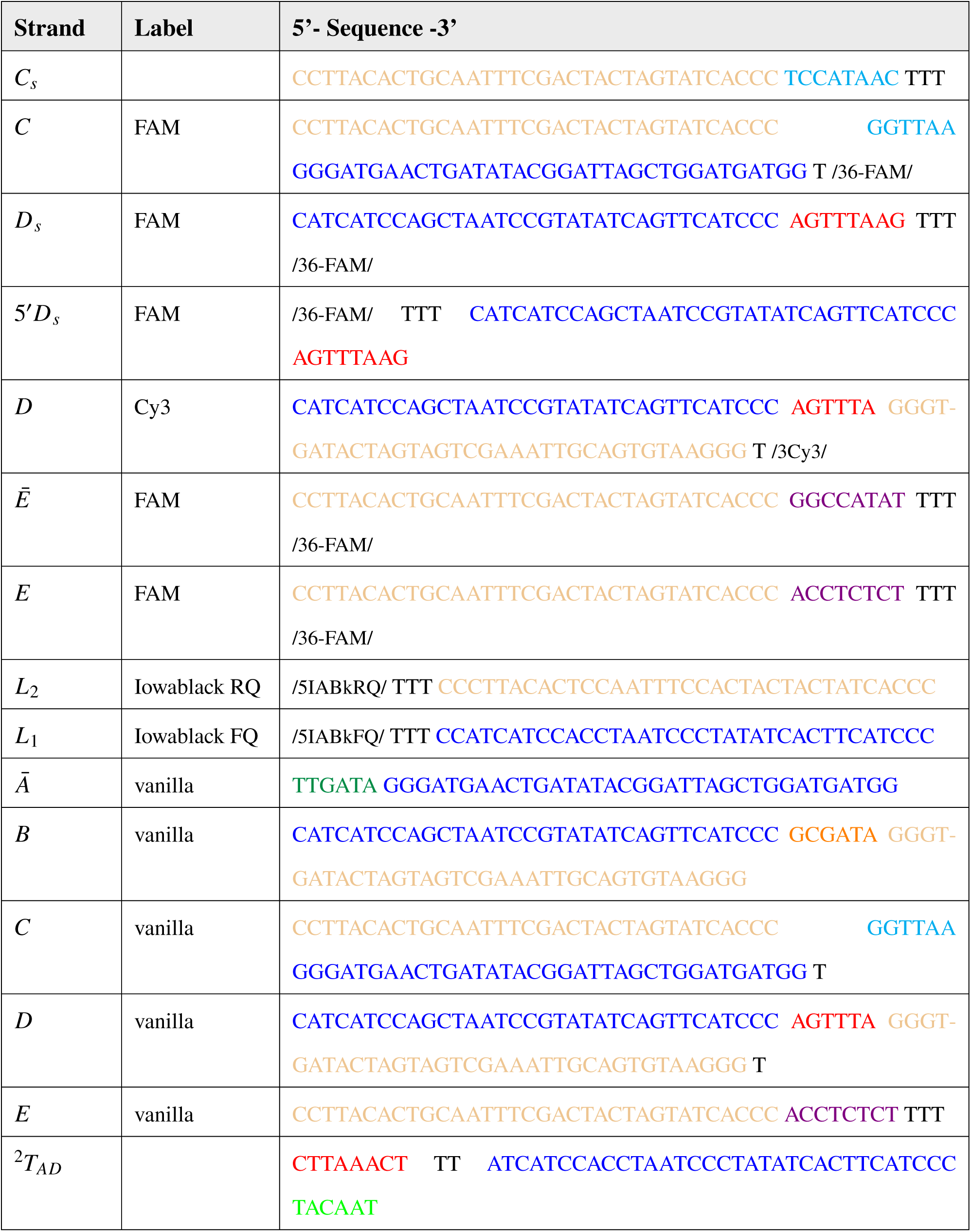
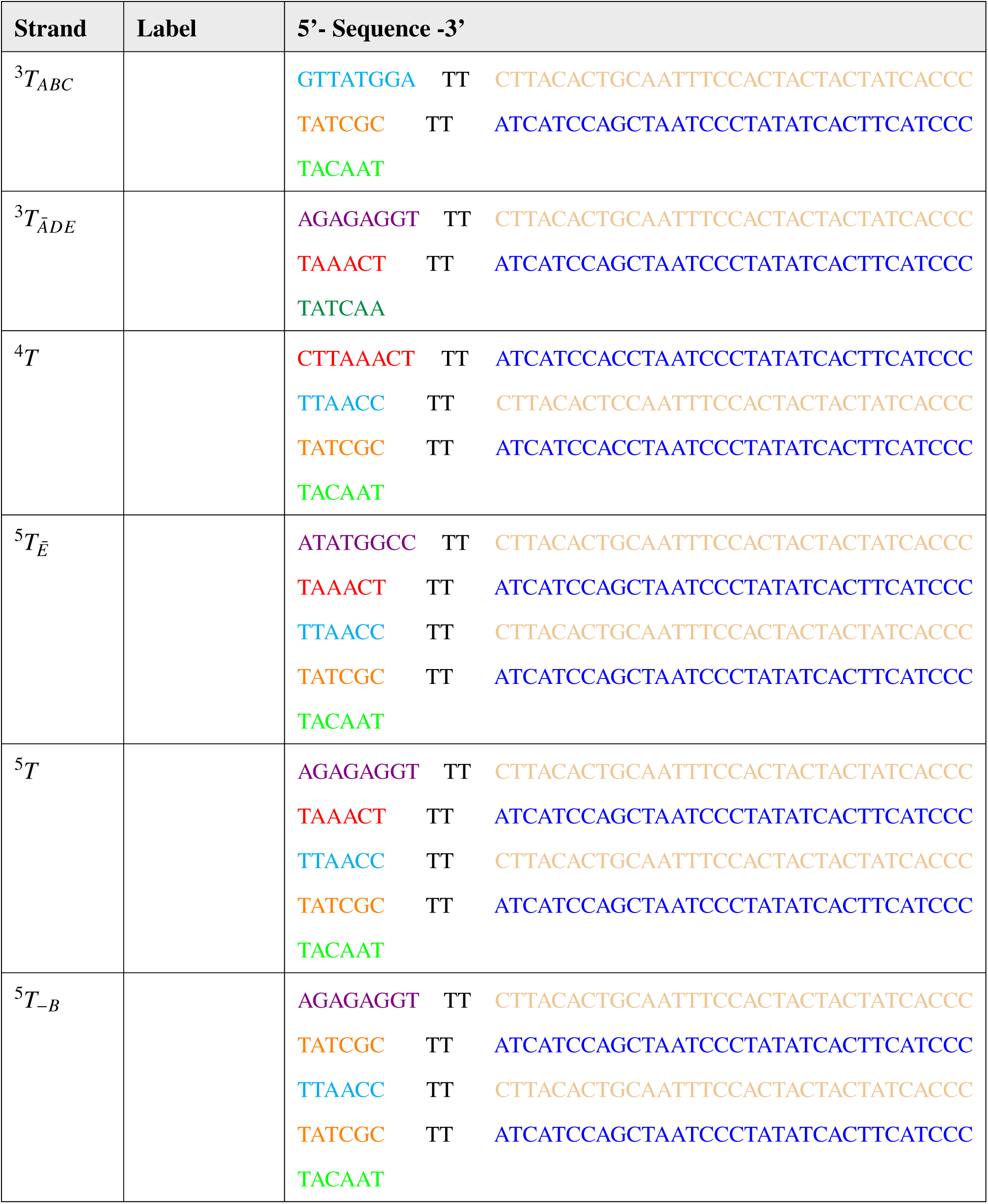
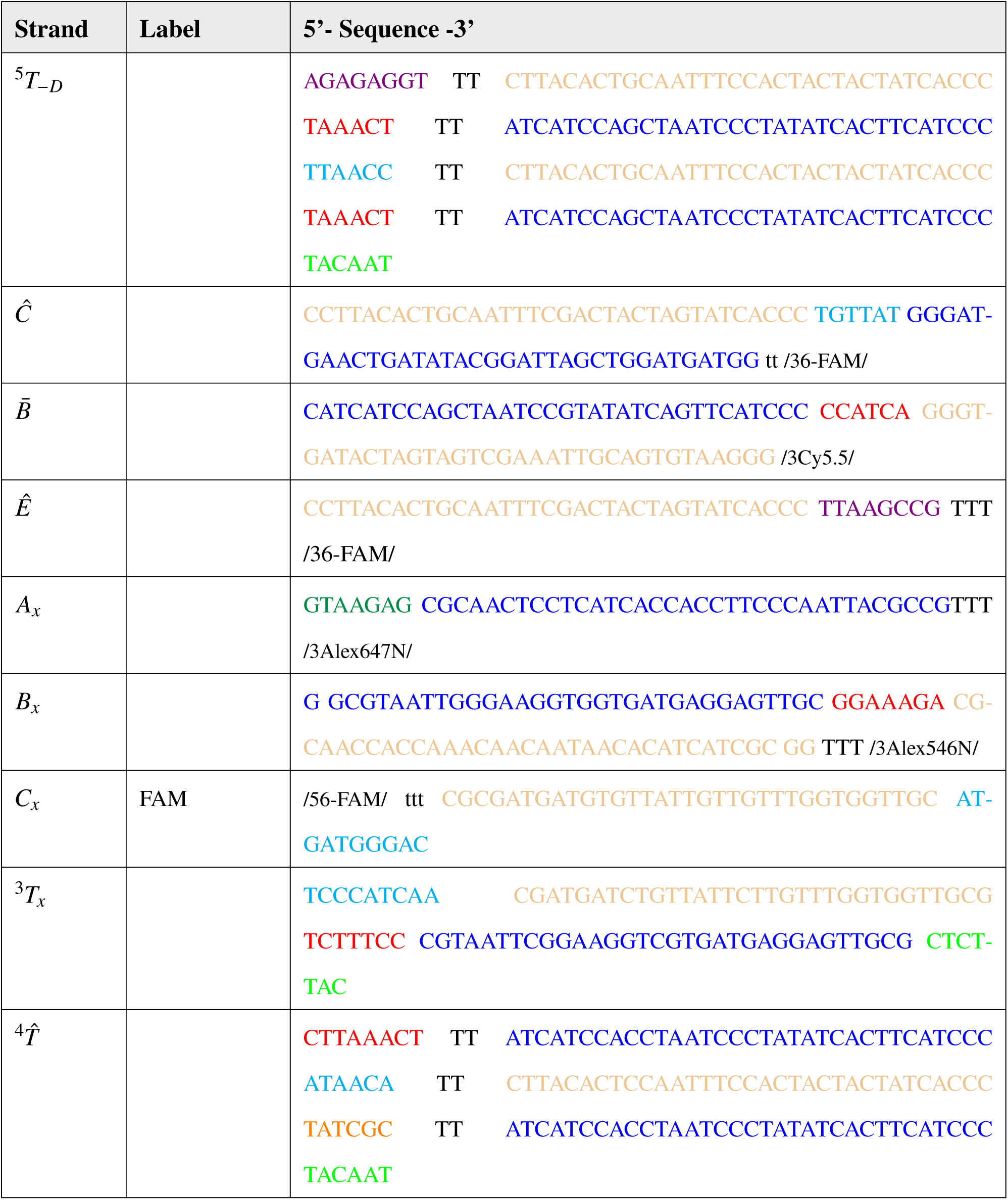
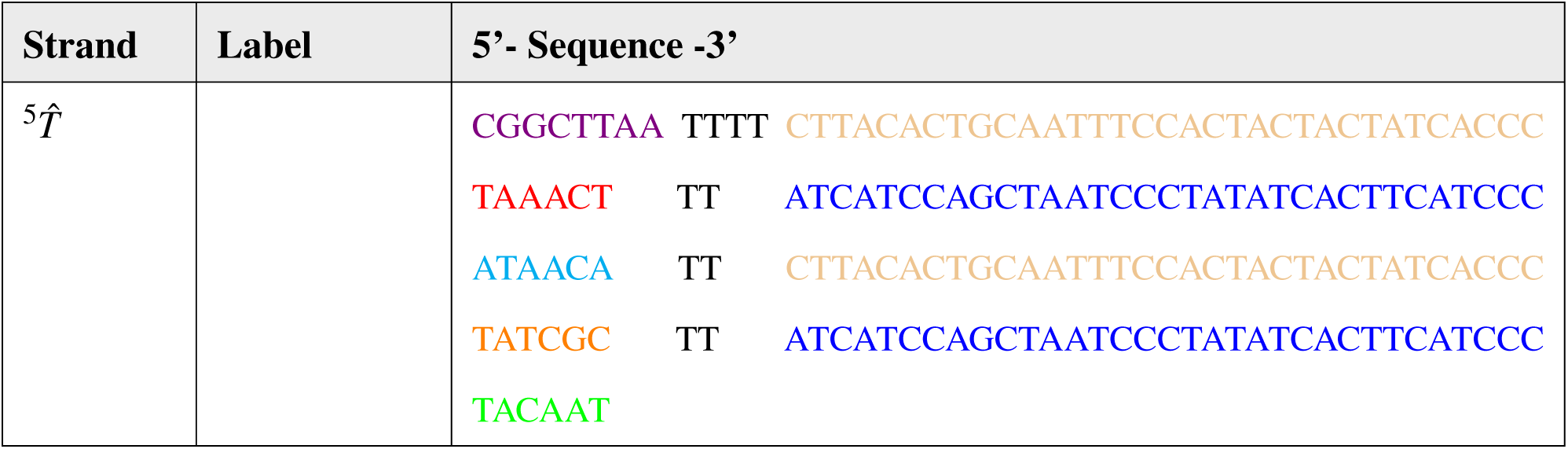
Sequences of all strands used in the experiments. The sequences are written from 5’ to 3’. The ‘‘Label’’ column lists the fluorescence tags of the corresponding strands if appropriate. ‘‘Vanilla’’ denotes an unlabelled variant of a labelled strand. Actual codes used by Integrated DNA Technologies for the fluorescence tags are written within //. Below is an overview of the colour codes:

**Table S2:**
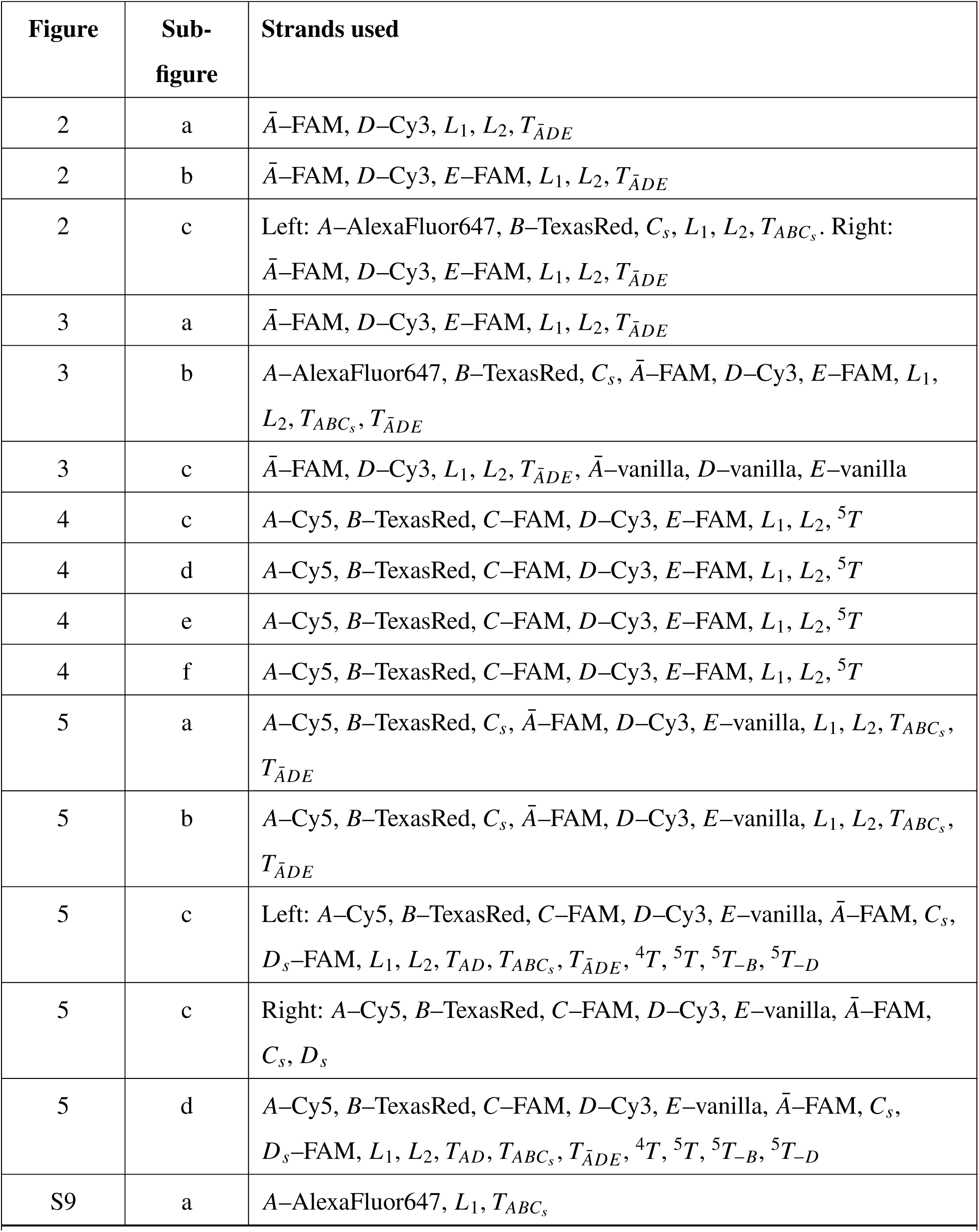
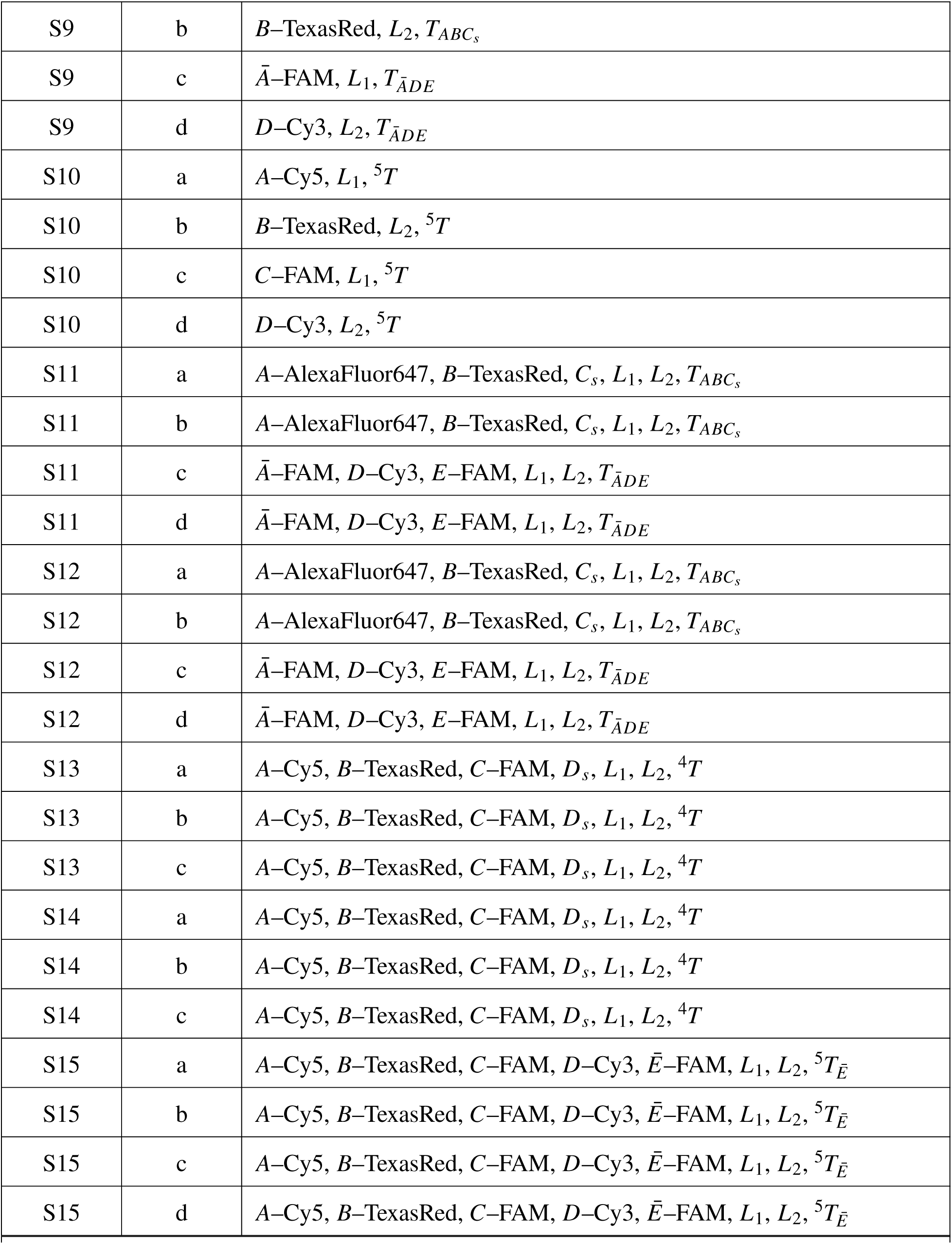
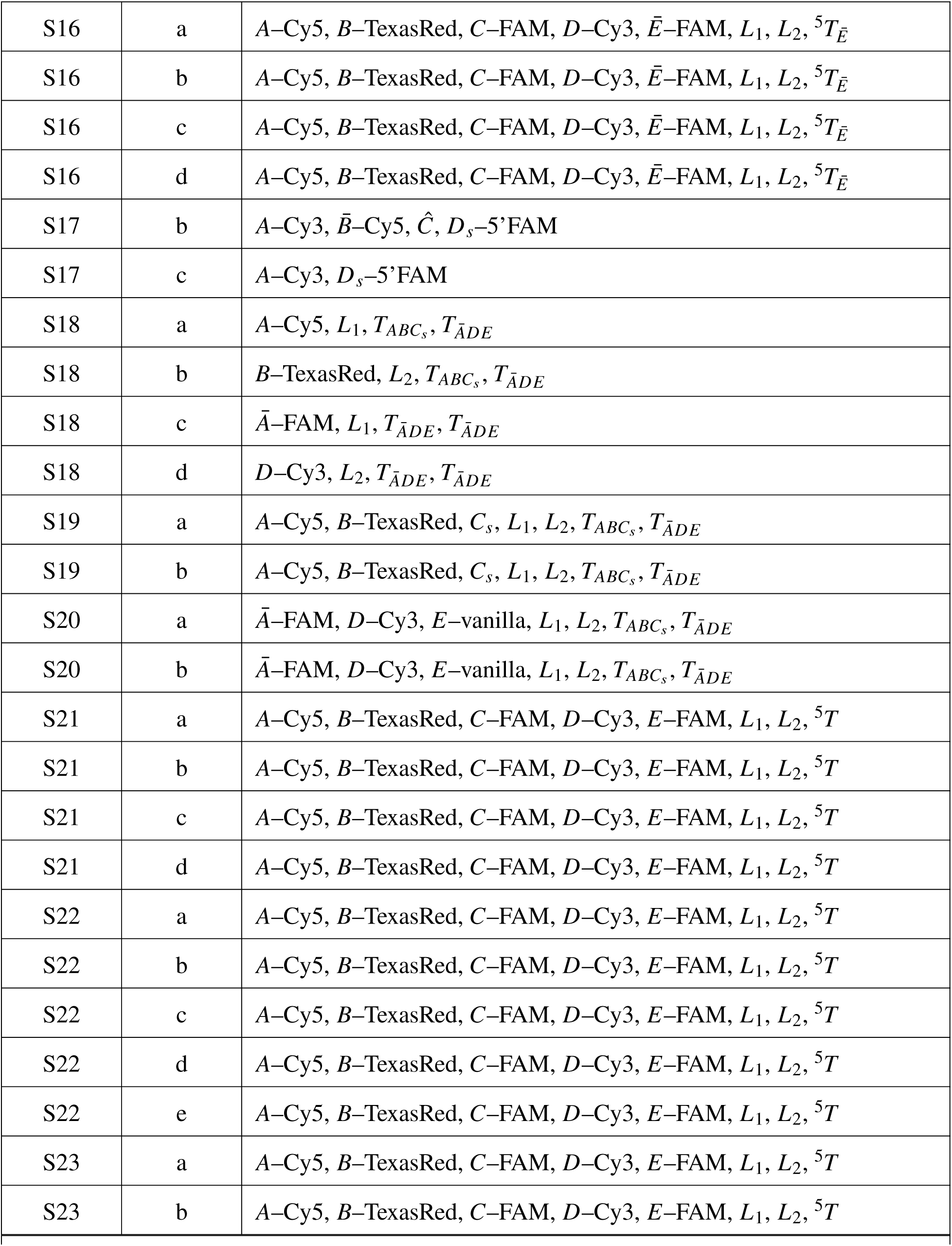
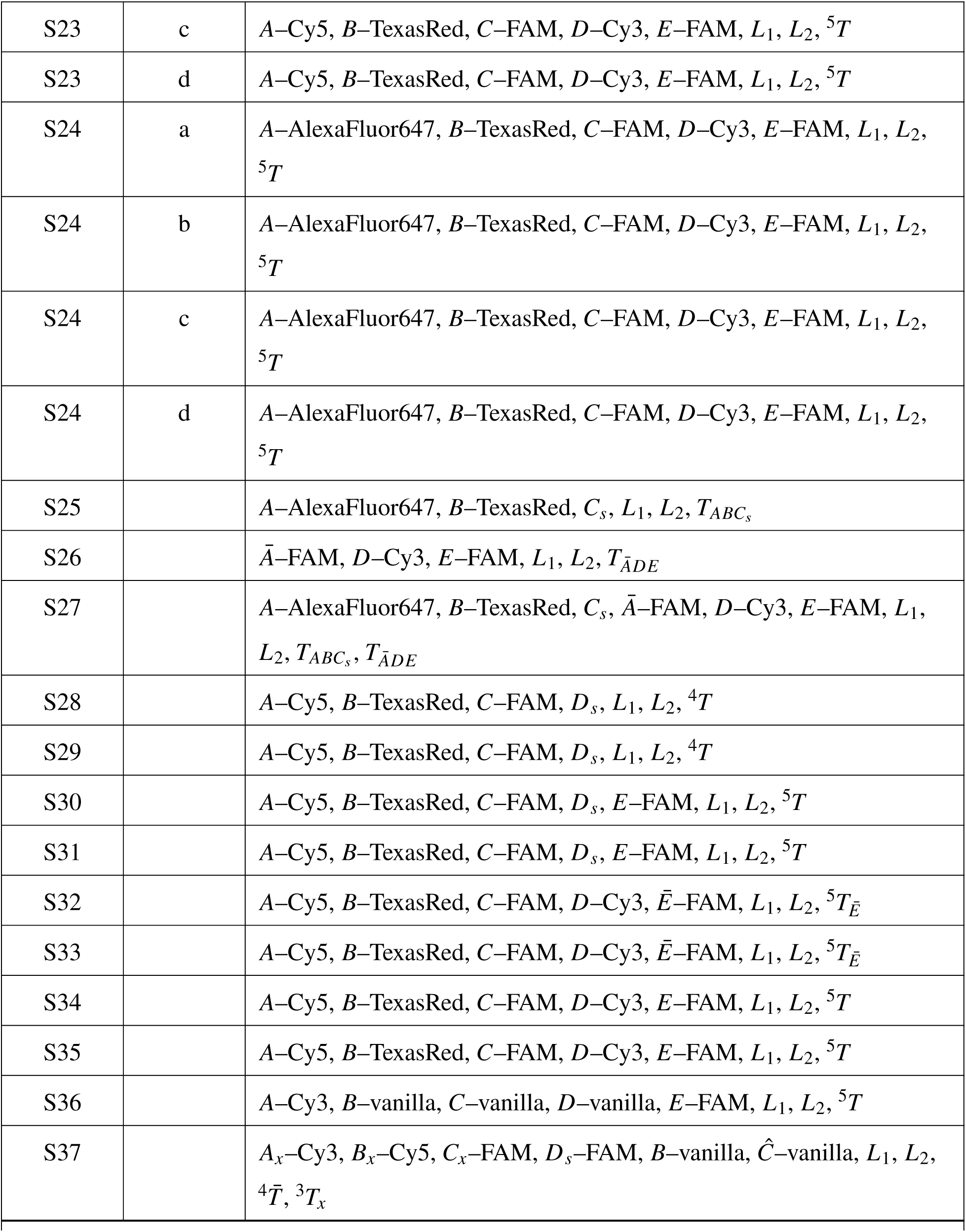
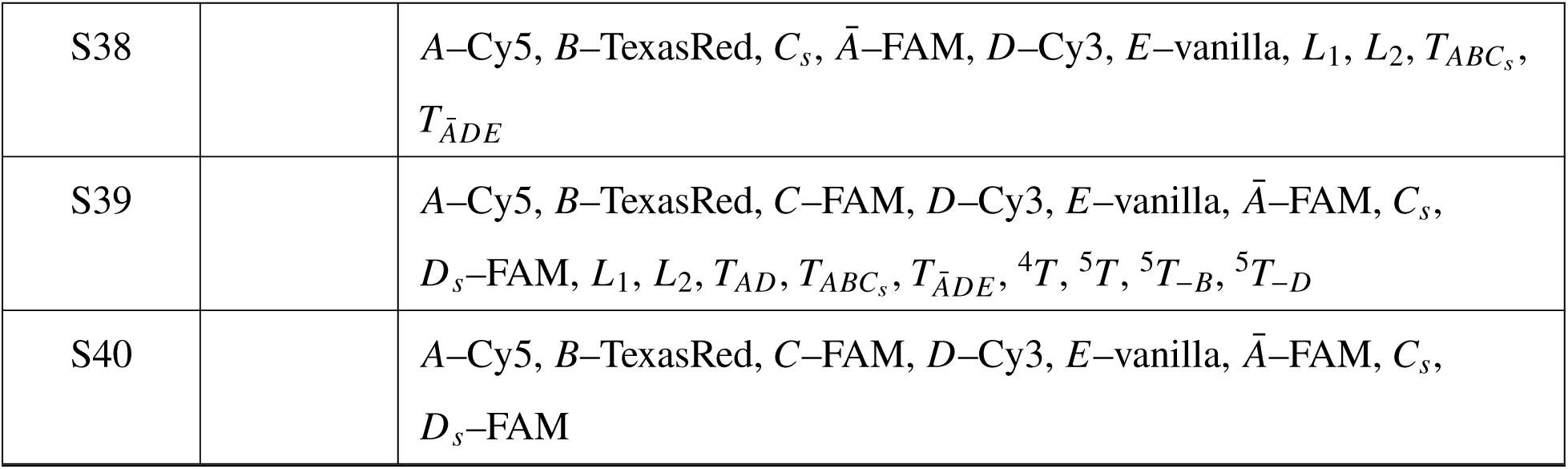
Figure and subfigure contents with strands used.

| Figure | Sub-figure | Strands used |
| --- | --- | --- |
| 2 | a | $\bar{A}$-FAM, $D$-Cy3, $L_1$, $L_2$, $T_{\bar{A}DE}$ |
| 2 | b | $\bar{A}$-FAM, $D$-Cy3, $E$-FAM, $L_1$, $L_2$, $T_{\bar{A}DE}$ |
| 2 | c | Left: $A$-AlexaFluor647, $B$-TexasRed, $C_s$, $L_1$, $L_2$, $T_{ABC_s}$. Right: $\bar{A}$-FAM, $D$-Cy3, $E$-FAM, $L_1$, $L_2$, $T_{\bar{A}DE}$ |
| 3 | a | $\bar{A}$-FAM, $D$-Cy3, $E$-FAM, $L_1$, $L_2$, $T_{\bar{A}DE}$ |
| 3 | b | $A$-AlexaFluor647, $B$-TexasRed, $C_s$, $\bar{A}$-FAM, $D$-Cy3, $E$-FAM, $L_1$, $L_2$, $T_{ABC_s}$, $T_{\bar{A}DE}$ |
| 3 | c | $\bar{A}$-FAM, $D$-Cy3, $L_1$, $L_2$, $T_{\bar{A}DE}$, $\bar{A}$-vanilla, $D$-vanilla, $E$-vanilla |
| 4 | c | $A$-Cy5, $B$-TexasRed, $C$-FAM, $D$-Cy3, $E$-FAM, $L_1$, $L_2$, ${}^5T$ |
| 4 | d | $A$-Cy5, $B$-TexasRed, $C$-FAM, $D$-Cy3, $E$-FAM, $L_1$, $L_2$, ${}^5T$ |
| 4 | e | $A$-Cy5, $B$-TexasRed, $C$-FAM, $D$-Cy3, $E$-FAM, $L_1$, $L_2$, ${}^5T$ |
| 4 | f | $A$-Cy5, $B$-TexasRed, $C$-FAM, $D$-Cy3, $E$-FAM, $L_1$, $L_2$, ${}^5T$ |
| 5 | a | $A$-Cy5, $B$-TexasRed, $C_s$, $\bar{A}$-FAM, $D$-Cy3, $E$-vanilla, $L_1$, $L_2$, $T_{ABC_s}$, $T_{\bar{A}DE}$ |
| 5 | b | $A$-Cy5, $B$-TexasRed, $C_s$, $\bar{A}$-FAM, $D$-Cy3, $E$-vanilla, $L_1$, $L_2$, $T_{ABC_s}$, $T_{\bar{A}DE}$ |
| 5 | c | Left: $A$-Cy5, $B$-TexasRed, $C$-FAM, $D$-Cy3, $E$-vanilla, $\bar{A}$-FAM, $C_s$, $D_s$-FAM, $L_1$, $L_2$, $T_{AD}$, $T_{ABC_s}$, $T_{\bar{A}DE}$, ${}^4T$, ${}^5T$, ${}^5T_{-B}$, ${}^5T_{-D}$ |
| 5 | c | Right: $A$-Cy5, $B$-TexasRed, $C$-FAM, $D$-Cy3, $E$-vanilla, $\bar{A}$-FAM, $C_s$, $D_s$ |
| 5 | d | $A$-Cy5, $B$-TexasRed, $C$-FAM, $D$-Cy3, $E$-vanilla, $\bar{A}$-FAM, $C_s$, $D_s$-FAM, $L_1$, $L_2$, $T_{AD}$, $T_{ABC_s}$, $T_{\bar{A}DE}$, ${}^4T$, ${}^5T$, ${}^5T_{-B}$, ${}^5T_{-D}$ |
| S9 | a | $A$-AlexaFluor647, $L_1$, $T_{ABC_s}$ |
| Continued on next page |  |  |
| S9 | b | $B\text{-TexasRed}, L_2, T_{ABC_s}$ |
| S9 | c | $\bar{A}\text{-FAM}, L_1, T_{\bar{A}DE}$ |
| S9 | d | $D\text{-Cy3}, L_2, T_{\bar{A}DE}$ |
| S10 | a | $A\text{-Cy5}, L_1, {}^5T$ |
| S10 | b | $B\text{-TexasRed}, L_2, {}^5T$ |
| S10 | c | $C\text{-FAM}, L_1, {}^5T$ |
| S10 | d | $D\text{-Cy3}, L_2, {}^5T$ |
| S11 | a | $A\text{-AlexaFluor647}, B\text{-TexasRed}, C_s, L_1, L_2, T_{ABC_s}$ |
| S11 | b | $A\text{-AlexaFluor647}, B\text{-TexasRed}, C_s, L_1, L_2, T_{ABC_s}$ |
| S11 | c | $\bar{A}\text{-FAM}, D\text{-Cy3}, E\text{-FAM}, L_1, L_2, T_{\bar{A}DE}$ |
| S11 | d | $\bar{A}\text{-FAM}, D\text{-Cy3}, E\text{-FAM}, L_1, L_2, T_{\bar{A}DE}$ |
| S12 | a | $A\text{-AlexaFluor647}, B\text{-TexasRed}, C_s, L_1, L_2, T_{ABC_s}$ |
| S12 | b | $A\text{-AlexaFluor647}, B\text{-TexasRed}, C_s, L_1, L_2, T_{ABC_s}$ |
| S12 | c | $\bar{A}\text{-FAM}, D\text{-Cy3}, E\text{-FAM}, L_1, L_2, T_{\bar{A}DE}$ |
| S12 | d | $\bar{A}\text{-FAM}, D\text{-Cy3}, E\text{-FAM}, L_1, L_2, T_{\bar{A}DE}$ |
| S13 | a | $A\text{-Cy5}, B\text{-TexasRed}, C\text{-FAM}, D_s, L_1, L_2, {}^4T$ |
| S13 | b | $A\text{-Cy5}, B\text{-TexasRed}, C\text{-FAM}, D_s, L_1, L_2, {}^4T$ |
| S13 | c | $A\text{-Cy5}, B\text{-TexasRed}, C\text{-FAM}, D_s, L_1, L_2, {}^4T$ |
| S14 | a | $A\text{-Cy5}, B\text{-TexasRed}, C\text{-FAM}, D_s, L_1, L_2, {}^4T$ |
| S14 | b | $A\text{-Cy5}, B\text{-TexasRed}, C\text{-FAM}, D_s, L_1, L_2, {}^4T$ |
| S14 | c | $A\text{-Cy5}, B\text{-TexasRed}, C\text{-FAM}, D_s, L_1, L_2, {}^4T$ |
| S15 | a | $A\text{-Cy5}, B\text{-TexasRed}, C\text{-FAM}, D\text{-Cy3}, \bar{E}\text{-FAM}, L_1, L_2, {}^5T_{\bar{E}}$ |
| S15 | b | $A\text{-Cy5}, B\text{-TexasRed}, C\text{-FAM}, D\text{-Cy3}, \bar{E}\text{-FAM}, L_1, L_2, {}^5T_{\bar{E}}$ |
| S15 | c | $A\text{-Cy5}, B\text{-TexasRed}, C\text{-FAM}, D\text{-Cy3}, \bar{E}\text{-FAM}, L_1, L_2, {}^5T_{\bar{E}}$ |
| S15 | d | $A\text{-Cy5}, B\text{-TexasRed}, C\text{-FAM}, D\text{-Cy3}, \bar{E}\text{-FAM}, L_1, L_2, {}^5T_{\bar{E}}$ |
| Continued on next page |  |  |
| S16 | a | $A\text{-Cy5}, B\text{-TexasRed}, C\text{-FAM}, D\text{-Cy3}, \bar{E}\text{-FAM}, L_1, L_2, {}^5T_{\bar{E}}$ |
| S16 | b | $A\text{-Cy5}, B\text{-TexasRed}, C\text{-FAM}, D\text{-Cy3}, \bar{E}\text{-FAM}, L_1, L_2, {}^5T_{\bar{E}}$ |
| S16 | c | $A\text{-Cy5}, B\text{-TexasRed}, C\text{-FAM}, D\text{-Cy3}, \bar{E}\text{-FAM}, L_1, L_2, {}^5T_{\bar{E}}$ |
| S16 | d | $A\text{-Cy5}, B\text{-TexasRed}, C\text{-FAM}, D\text{-Cy3}, \bar{E}\text{-FAM}, L_1, L_2, {}^5T_{\bar{E}}$ |
| S17 | b | $A\text{-Cy3}, \bar{B}\text{-Cy5}, \hat{C}, D_s\text{-5'FAM}$ |
| S17 | c | $A\text{-Cy3}, D_s\text{-5'FAM}$ |
| S18 | a | $A\text{-Cy5}, L_1, T_{ABC_s}, T_{\bar{A}DE}$ |
| S18 | b | $B\text{-TexasRed}, L_2, T_{ABC_s}, T_{\bar{A}DE}$ |
| S18 | c | $\bar{A}\text{-FAM}, L_1, T_{\bar{A}DE}, T_{\bar{A}DE}$ |
| S18 | d | $D\text{-Cy3}, L_2, T_{\bar{A}DE}, T_{\bar{A}DE}$ |
| S19 | a | $A\text{-Cy5}, B\text{-TexasRed}, C_s, L_1, L_2, T_{ABC_s}, T_{\bar{A}DE}$ |
| S19 | b | $A\text{-Cy5}, B\text{-TexasRed}, C_s, L_1, L_2, T_{ABC_s}, T_{\bar{A}DE}$ |
| S20 | a | $\bar{A}\text{-FAM}, D\text{-Cy3}, E\text{-vanilla}, L_1, L_2, T_{ABC_s}, T_{\bar{A}DE}$ |
| S20 | b | $\bar{A}\text{-FAM}, D\text{-Cy3}, E\text{-vanilla}, L_1, L_2, T_{ABC_s}, T_{\bar{A}DE}$ |
| S21 | a | $A\text{-Cy5}, B\text{-TexasRed}, C\text{-FAM}, D\text{-Cy3}, E\text{-FAM}, L_1, L_2, {}^5T$ |
| S21 | b | $A\text{-Cy5}, B\text{-TexasRed}, C\text{-FAM}, D\text{-Cy3}, E\text{-FAM}, L_1, L_2, {}^5T$ |
| S21 | c | $A\text{-Cy5}, B\text{-TexasRed}, C\text{-FAM}, D\text{-Cy3}, E\text{-FAM}, L_1, L_2, {}^5T$ |
| S21 | d | $A\text{-Cy5}, B\text{-TexasRed}, C\text{-FAM}, D\text{-Cy3}, E\text{-FAM}, L_1, L_2, {}^5T$ |
| S22 | a | $A\text{-Cy5}, B\text{-TexasRed}, C\text{-FAM}, D\text{-Cy3}, E\text{-FAM}, L_1, L_2, {}^5T$ |
| S22 | b | $A\text{-Cy5}, B\text{-TexasRed}, C\text{-FAM}, D\text{-Cy3}, E\text{-FAM}, L_1, L_2, {}^5T$ |
| S22 | c | $A\text{-Cy5}, B\text{-TexasRed}, C\text{-FAM}, D\text{-Cy3}, E\text{-FAM}, L_1, L_2, {}^5T$ |
| S22 | d | $A\text{-Cy5}, B\text{-TexasRed}, C\text{-FAM}, D\text{-Cy3}, E\text{-FAM}, L_1, L_2, {}^5T$ |
| S22 | e | $A\text{-Cy5}, B\text{-TexasRed}, C\text{-FAM}, D\text{-Cy3}, E\text{-FAM}, L_1, L_2, {}^5T$ |
| S23 | a | $A\text{-Cy5}, B\text{-TexasRed}, C\text{-FAM}, D\text{-Cy3}, E\text{-FAM}, L_1, L_2, {}^5T$ |
| S23 | b | $A\text{-Cy5}, B\text{-TexasRed}, C\text{-FAM}, D\text{-Cy3}, E\text{-FAM}, L_1, L_2, {}^5T$ |
| S23 | c | $A\text{-Cy5}, B\text{-TexasRed}, C\text{-FAM}, D\text{-Cy3}, E\text{-FAM}, L_1, L_2, {}^5T$ |
| S23 | d | $A\text{-Cy5}, B\text{-TexasRed}, C\text{-FAM}, D\text{-Cy3}, E\text{-FAM}, L_1, L_2, {}^5T$ |
| S24 | a | $A\text{-AlexaFluor647}, B\text{-TexasRed}, C\text{-FAM}, D\text{-Cy3}, E\text{-FAM}, L_1, L_2, {}^5T$ |
| S24 | b | $A\text{-AlexaFluor647}, B\text{-TexasRed}, C\text{-FAM}, D\text{-Cy3}, E\text{-FAM}, L_1, L_2, {}^5T$ |
| S24 | c | $A\text{-AlexaFluor647}, B\text{-TexasRed}, C\text{-FAM}, D\text{-Cy3}, E\text{-FAM}, L_1, L_2, {}^5T$ |
| S24 | d | $A\text{-AlexaFluor647}, B\text{-TexasRed}, C\text{-FAM}, D\text{-Cy3}, E\text{-FAM}, L_1, L_2, {}^5T$ |
| S25 | | $A\text{-AlexaFluor647}, B\text{-TexasRed}, C_s, L_1, L_2, T_{ABC_s}$ |
| S26 | | $\bar{A}\text{-FAM}, D\text{-Cy3}, E\text{-FAM}, L_1, L_2, T_{\bar{A}DE}$ |
| S27 | | $A\text{-AlexaFluor647}, B\text{-TexasRed}, C_s, \bar{A}\text{-FAM}, D\text{-Cy3}, E\text{-FAM}, L_1, L_2, T_{ABC_s}, T_{\bar{A}DE}$ |
| S28 | | $A\text{-Cy5}, B\text{-TexasRed}, C\text{-FAM}, D_s, L_1, L_2, {}^4T$ |
| S29 | | $A\text{-Cy5}, B\text{-TexasRed}, C\text{-FAM}, D_s, L_1, L_2, {}^4T$ |
| S30 | | $A\text{-Cy5}, B\text{-TexasRed}, C\text{-FAM}, D_s, E\text{-FAM}, L_1, L_2, {}^5T$ |
| S31 | | $A\text{-Cy5}, B\text{-TexasRed}, C\text{-FAM}, D_s, E\text{-FAM}, L_1, L_2, {}^5T$ |
| S32 | | $A\text{-Cy5}, B\text{-TexasRed}, C\text{-FAM}, D\text{-Cy3}, \bar{E}\text{-FAM}, L_1, L_2, {}^5T_{\bar{E}}$ |
| S33 | | $A\text{-Cy5}, B\text{-TexasRed}, C\text{-FAM}, D\text{-Cy3}, \bar{E}\text{-FAM}, L_1, L_2, {}^5T_{\bar{E}}$ |
| S34 | | $A\text{-Cy5}, B\text{-TexasRed}, C\text{-FAM}, D\text{-Cy3}, E\text{-FAM}, L_1, L_2, {}^5T$ |
| S35 | | $A\text{-Cy5}, B\text{-TexasRed}, C\text{-FAM}, D\text{-Cy3}, E\text{-FAM}, L_1, L_2, {}^5T$ |
| S36 | | $A\text{-Cy3}, B\text{-vanilla}, C\text{-vanilla}, D\text{-vanilla}, E\text{-FAM}, L_1, L_2, {}^5T$ |
| S37 | | $A_x\text{-Cy3}, B_x\text{-Cy5}, C_x\text{-FAM}, D_s\text{-FAM}, B\text{-vanilla}, \hat{C}\text{-vanilla}, L_1, L_2, {}^4\bar{T}, {}^3T_x$ |
| Continued on next page |  |  |
| S38 | | $A\text{-Cy5}, B\text{-TexasRed}, C_s, \bar{A}\text{-FAM}, D\text{-Cy3}, E\text{-vanilla}, L_1, L_2, T_{ABC_s}, T_{\bar{A}DE}$ |
| S39 | | $A\text{-Cy5}, B\text{-TexasRed}, C\text{-FAM}, D\text{-Cy3}, E\text{-vanilla}, \bar{A}\text{-FAM}, C_s, D_s\text{-FAM}, L_1, L_2, T_{AD}, T_{ABC_s}, T_{\bar{A}DE}, {}^4T, {}^5T, {}^5T_{-B}, {}^5T_{-D}$ |
| S40 | | $A\text{-Cy5}, B\text{-TexasRed}, C\text{-FAM}, D\text{-Cy3}, E\text{-vanilla}, \bar{A}\text{-FAM}, C_s, D_s\text{-FAM}$ |

**Table S3:** Plate reader settings. The table contains the excitation and emission wavelength, and gain settings used for different fluorophores to follow the reaction kinetics at different concentrations.

| Channel | Excitation wavelength (nm) | Dichroic filter | Emission wavelength (nm) | Gain for 10 nM reactions | Gain for 100 nM reactions |
| --- | --- | --- | --- | --- | --- |
| Cy5, AlexaFluor647 | 625-30 | 652.5 | 680-30 | 2900 | 2400 |
| Texas Red | 580-20 | 602.5 | 630-30 | 2500 | 1900 |
| Cy3, AlexaFluor546 | 530-20 | 552.5 | 535-30 | 2000 | 1500 |
| FAM, AlexaFluor488 | 488-14 | 507.5 | 580-30 | 2000 | 1400 |

**Table S4:** Photomultiplier tube (PMT) voltage settings for different excitation lasers and emission filters as used for gel scanning. A set of lower PMT voltage settings was used in the experiments with high monomer concentrations to ensure that the signals were not saturated.

| Name | Compatible fluorophores | Optical Settings |  | PMT Voltage (V) |  |
| --- | --- | --- | --- | --- | --- |
|  |  | Excitation laser (nm) | Emission filter (nm) | Standard | High |
| Cy2 | FAM, AlexaFluor 488 | 488 | 525BP20 | 542 | 444 |
| Cy3 | Cy3, AlexaFluor 546 | 532 | 570BP20 | 688 | 569 |
| Cy5 | Cy5, AlexaFluor 647 | 635 | 670BP30 | 596 | 451 |

**Table S5:** Protocols used for the reactions where templates bind to blocked monomers *via* TMSD. The table shows the concentrations of the stock solutions used, and the required amounts for the TMSD reactions. Reac*^n^* denotes reaction wells, PC denotes positive controls. Target concentrations are shown in [brackets]

| Components with stock concentrations | Required volume (μL) [Target Concentration (nM) in 200 μL volume] |  |  |  |  |  |  |  |  |  |  |
| --- | --- | --- | --- | --- | --- | --- | --- | --- | --- | --- | --- |
|  | Reac<sup>n</sup> 1 | Reac<sup>n</sup> 2 | Reac<sup>n</sup> 3 | Reac<sup>n</sup> 4 | Reac<sup>n</sup> 5 | Reac<sup>n</sup> 6 | PC1 | PC2 | PC3 | PC4 | Blank |
| **Step 1: Baseline** |  |  |  |  |  |  |  |  |  |  |  |
| 2% Tween20 in 1X TAE, 1 M NaCl | 20 | 20 | 20 | 20 | 20 | 20 | 20 | 20 | 20 | 20 | 20 |
| 1X TAE, 1 M NaCl | 120 | 120 | 120 | 120 | 120 | 120 | 120 | 123 | 125 | 127 | 130 |
| ML [200 nM] | 10 [10] | 10 [10] | 10 [10] | 10 [10] | 10 [10] | 10 [10] | - | - | - | - | - |
| MT [200 nM] | - | - | - | - | - | - | 10 [10] | 7 [7] | 5 [5] | 3 [3] | 0 [0] |
| **Step 2: Reaction kinetics** |  |  |  |  |  |  |  |  |  |  |  |
| Template [100 nM] | 0 [0] | 4 [2] | 8 [4] | 12 [6] | 16 [8] | 20 [10] | - | - | - | - | - |
| 1X TAE, 1 M NaCl | 50 | 46 | 42 | 38 | 34 | 30 | 50 | 50 | 50 | 50 | 50 |
| **Step 3: Saturation** |  |  |  |  |  |  |  |  |  |  |  |
| Template [2 μM] | 10 [100] | 10 [100] | 10 [100] | 10 [100] | 10 [100] | 10 [100] | - | - | - | - | - |
| 1X TAE, 1 M NaCl | - | - | - | - | - | - | 10 | 10 | 10 | 10 | 10 |

**Table S6:** Protocols used for trimer formation with standard monomer concentrations. The table shows the concentrations of the stock solutions used, and the required amounts for the TMSD reactions. Reac*^n^* denotes reaction wells, PC denotes positive controls. Target concentrations are shown in [brackets]. The first 10 data points in the initialisation phase of Reac*^n^*1, where the extent of undesired leak reactions is negligible, are used as the negative control baseline for the fluorescence data analysis as described in the methods.

| Components with stock concentrations | Required volume (μL) [Target Concentration (nM) in 200 μL volume] |  |  |  |  |  |  |  |  |  |  |
| --- | --- | --- | --- | --- | --- | --- | --- | --- | --- | --- | --- |
|  | Reac<sup>n</sup> 1 | Reac<sup>n</sup> 2 | Reac<sup>n</sup> 3 | Reac<sup>n</sup> 4 | Reac<sup>n</sup> 5 | Reac<sup>n</sup> 6 | PC1 | PC2 | PC3 | PC4 | Blank |
| Step1:Baseline |  |  |  |  |  |  |  |  |  |  |  |
| 2% Tween20 in 1X TAE, 1M NaCl | 20 | 20 | 20 | 20 | 20 | 20 | 20 | 20 | 20 | 20 | 20 |
| 1X TAE, 1M NaCl | 100 | 100 | 100 | 100 | 100 | 100 | 120 | 123 | 125 | 127 | 130 |
| $A_1L_1$ or $\bar{A}_1L_1$ [200 nM] | 10 [10] | 10 [10] | 10 [10] | 10 [10] | 10 [10] | 10 [10] | - | - | - | - | - |
| $BL_2$ or $DL_2$ [200 nM] | 10 [10] | 10 [10] | 10 [10] | 10 [10] | 10 [10] | 10 [10] | - | - | - | - | - |
| $C_s$ or $E$ [200 nM] | 10 [10] | 10 [10] | 10 [10] | 10 [10] | 10 [10] | 10 [10] | - | - | - | - | - |
| $ABC_s$ or $\bar{A}DE$ [200 nM] | - | - | - | - | - | - | 10 [10] | 7 [7] | 5 [5] | 3 [3] | - |
| Step 2: Reaction kinetics |  |  |  |  |  |  |  |  |  |  |  |
| Template [100 nM] | 0 [0] | 4 [2] | 8 [4] | 12 [6] | 16 [8] | 20 [10] | - | - | - | - | - |
| 1X TAE, 1M NaCl | 50 | 46 | 42 | 38 | 34 | 30 | 50 | 50 | 50 | 50 | 50 |

**Table S7:** Protocols used for trimer formation with high monomer concentrations. The table lists stock concentrations and required volumes for each reaction well (Reac*^n^*), positive control (PC), and blank. Target concentrations are shown in [brackets]. The first 10 data points in the initialisation phase of Reac*^n^*1, where the extent of undesired leak reactions is negligible, are used as the negative control baseline for the fluorescence data analysis as described in the methods.

| Components with stock concentrations | Required volume (μL) [Target Concentration (nM) in 200 μL volume] |  |  |  |  |  |  |  |  |  |
| --- | --- | --- | --- | --- | --- | --- | --- | --- | --- | --- |
|  | Reac<sup>n</sup> 1 | Reac<sup>n</sup> 2 | Reac<sup>n</sup> 3 | Reac<sup>n</sup> 4 | Reac<sup>n</sup> 5 | PC1 | PC2 | PC3 | PC4 | Blank |
| **Step 1: Baseline** |  |  |  |  |  |  |  |  |  |  |
| 2% Tween20 in 1X TAE, 1M NaCl | 20 | 20 | 20 | 20 | 20 | 20 | 20 | 20 | 20 | 20 |
| 1X TAE, 1M NaCl | 70 | 70 | 70 | 70 | 70 | 120 | 123 | 125 | 127 | 130 |
| AL<sub>1</sub> or $\bar{A}L_1$ [1000 nM] | 20 [100] | 20 [100] | 20 [100] | 20 [100] | 20 [100] | - | - | - | - | - |
| BL<sub>2</sub> or DL<sub>2</sub> [1000 nM] | 20 [100] | 20 [100] | 20 [100] | 20 [100] | 20 [100] | - | - | - | - | - |
| C<sub>s</sub> or E [1000 nM] | 20 [100] | 20 [100] | 20 [100] | 20 [100] | 20 [100] | - | - | - | - | - |
| ABC<sub>s</sub> or $\bar{A}DE$ [1000 nM] | - | - | - | - | - | 20 [100] | 15 [75] | 10 [50] | 5 [25] | - |
| **Step 2: Reaction kinetics** |  |  |  |  |  |  |  |  |  |  |
| Template [100 nM] | 0 [0] | 4 [2] | 6 [3] | 8 [4] | 10 [5] | - | - | - | - | - |
| 1X TAE, 1M NaCl | 50 | 46 | 44 | 42 | 40 | 50 | 50 | 50 | 50 | 50 |

**Table S8:** Protocol used for testing product inhibition in *ĀDE* trimer formation. The table lists stock concentrations and required volumes for each reaction well (Reac*^n^*), positive control (PC), and blank. Target concentrations are shown in [brackets]. The first 10 data points in the initialisation phase of Reac*^n^*1, where the extent of undesired leak reactions is negligible, are used as the negative control baseline for the fluorescence data analysis as described in the methods.

| Components with stock concentrations | Reac<sup>n</sup>1 | Reac<sup>n</sup>2 | Reac<sup>n</sup>3 | Reac<sup>n</sup>4 | Reac<sup>n</sup>5 | Reac<sup>n</sup>6 | Reac<sup>n</sup>7 | Reac<sup>n</sup>8 | PC1 | PC2 | PC3 | PC4 | Blank |
| --- | --- | --- | --- | --- | --- | --- | --- | --- | --- | --- | --- | --- | --- |
| **Step1:Baseline** |  |  |  |  |  |  |  |  |  |  |  |  |  |
| 2% Tween20 in 1X TAE, 1M NaCl | 20 | 20 | 20 | 20 | 20 | 20 | 20 | 20 | 20 | 20 | 20 | 20 | 20 |
| 1X TAE, 1M NaCl | 100 | 90 | 80 | 70 | 60 | 50 | 25 | 0 | 120 | 123 | 125 | 127 | 130 |
| $\bar{A}L_1$ [200 nM] | 10 [10] | 10 [10] | 10 [10] | 10 [10] | 10 [10] | 10 [10] | 10 [10] | 10 [10] | - | - | - | - | - |
| $DL_2$ [200 nM] | 10 [10] | 10 [10] | 10 [10] | 10 [10] | 10 [10] | 10 [10] | 10 [10] | 10 [10] | - | - | - | - | - |
| $E - vanilla$ [200 nM] | 10 [10] | 10 [10] | 10 [10] | 10 [10] | 10 [10] | 10 [10] | 10 [10] | 10 [10] | - | - | - | - | - |
| Unlabelled $\bar{A}DE$ [200 nM] | - | 10 [10] | 20 [20] | 30 [30] | 40 [40] | 50 [50] | 75 [50] | 100 [100] | - | - | - | - | - |
| $\bar{A}DE$ [200 nM] | - | - | - | - | - | - | - | - | 10 [10] | 7 [7] | 5 [5] | 3 [3] | - |
| **Step 2: Reaction kinetics** |  |  |  |  |  |  |  |  |  |  |  |  |  |
| Template [100 nM] | 8 [4] | 8 [4] | 8 [4] | 8 [4] | 8 [4] | 8 [4] | 8 [4] | 8 [4] | - | - | - | - | - |
| 1X TAE, 1M NaCl | 42 | 42 | 42 | 42 | 42 | 42 | 42 | 42 | 50 | 50 | 50 | 50 | 50 |

**Table S9:** Protocol used for tetramer formation with standard monomer concentrations. The table lists stock concentrations and required volumes for each reaction well (Reac*^n^*), positive control (PC), and blank. Target concentrations are shown in [brackets]. The first 10 data points in the initialisation phase of Reac*^n^*1, where the extent of undesired leak reactions is negligible, are used as the negative control baseline for the fluorescence data analysis as described in the methods.

| Components with stock concentrations | Required volume (μL) [Target Concentration (nM) in 200 μL volume] |  |  |  |  |  |  |  |  |  |  |
| --- | --- | --- | --- | --- | --- | --- | --- | --- | --- | --- | --- |
|  | Reac<sup>n</sup> 1 | Reac<sup>n</sup> 2 | Reac<sup>n</sup> 3 | Reac<sup>n</sup> 4 | Reac<sup>n</sup> 5 | Reac<sup>n</sup> 6 | PC1 | PC2 | PC3 | PC4 | Blank |
| **Step1:Baseline** |  |  |  |  |  |  |  |  |  |  |  |
| 2% Tween20 in 1X TAE, 1 M NaCl | 20 | 20 | 20 | 20 | 20 | 20 | 20 | 20 | 20 | 20 | 20 |
| 1X TAE, 1 M NaCl | 100 | 100 | 100 | 100 | 100 | 100 | 110 | 116 | 120 | 124 | 130 |
| AL<sub>1</sub> [200 nM] | 10 [10] | 10 [10] | 10 [10] | 10 [10] | 10 [10] | 10 [10] | - | - | - | - | - |
| BL<sub>2</sub> [200 nM] | 10 [10] | 10 [10] | 10 [10] | 10 [10] | 10 [10] | 10 [10] | - | - | - | - | - |
| CL<sub>1</sub> [200 nM] | 10 [10] | 10 [10] | 10 [10] | 10 [10] | 10 [10] | 10 [10] | - | - | - | - | - |
| D<sub>s</sub> [200 nM] | 10 [10] | 10 [10] | 10 [10] | 10 [10] | 10 [10] | 10 [10] | - | - | - | - | - |
| Pre-annealed AB [200 nM] | - | - | - | - | - | - | 10 [10] | 7 [7] | 5 [5] | 3 [3] | - |
| Pre-annealed CD<sub>s</sub> [200 nM] | - | - | - | - | - | - | 10 [10] | 7 [7] | 5 [5] | 3 [3] | - |
| **Step 2: Reaction kinetics** |  |  |  |  |  |  |  |  |  |  |  |
| <sup>4</sup>T [100 nM] | 0 [0] | 4 [2] | 8 [4] | 12 [6] | 16 [8] | 20 [10] | - | - | - | - | - |
| 1X TAE, 1 M NaCl | 50 | 46 | 42 | 38 | 34 | 30 | 50 | 50 | 50 | 50 | 50 |

**Table S10:**
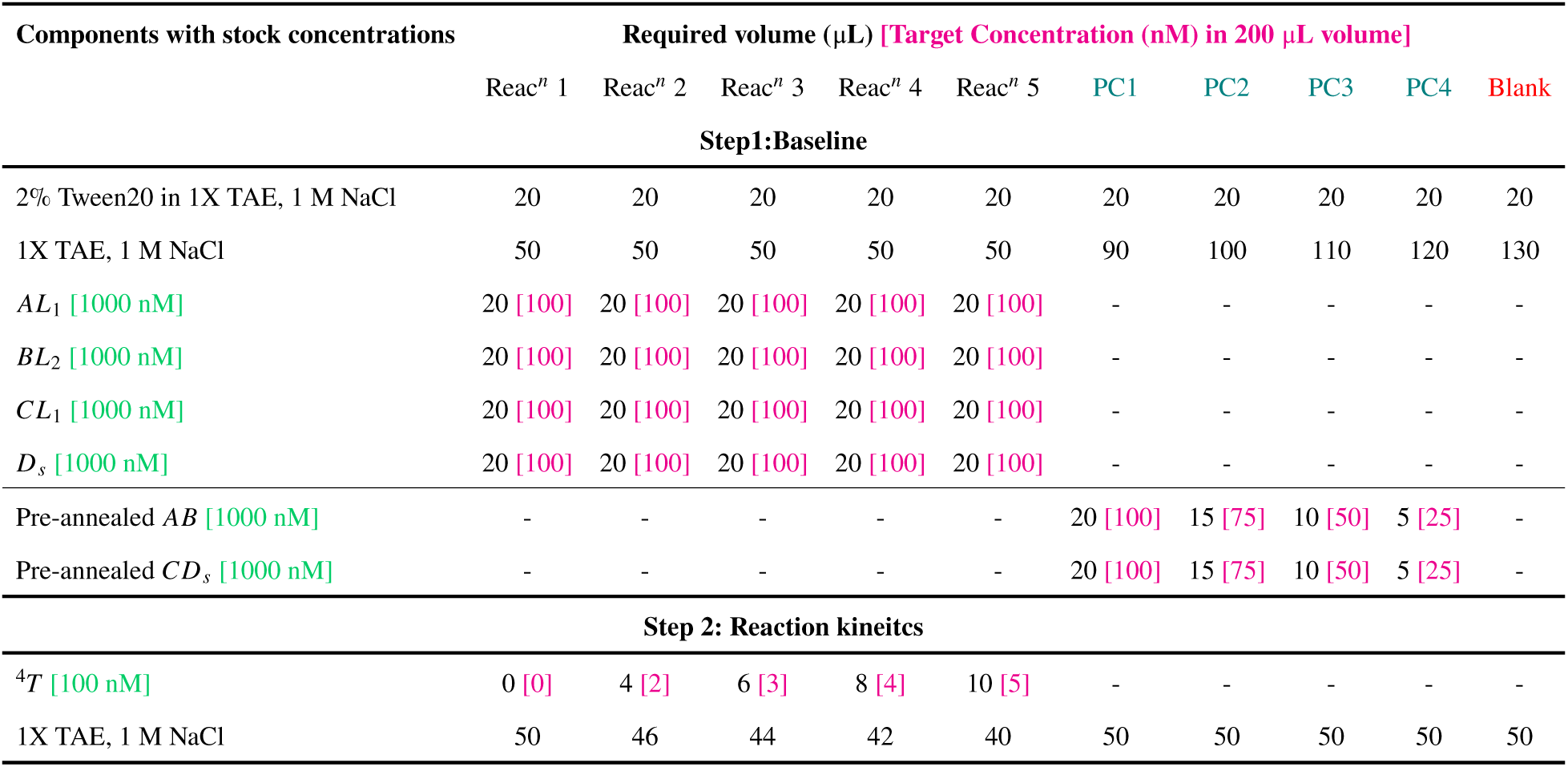
Protocol used for tetramer formation with high monomer concentrations. The table lists stock concentrations and required volumes for each reaction well (Reac*^n^*), positive control (PC), and blank. Target concentrations are shown in [brackets]. The first 10 data points in the initialisation phase of Reac*^n^*1, where the extent of undesired leak reactions is negligible, are used as the negative control baseline for the fluorescence data analysis as described in the methods.

**Table S11:**
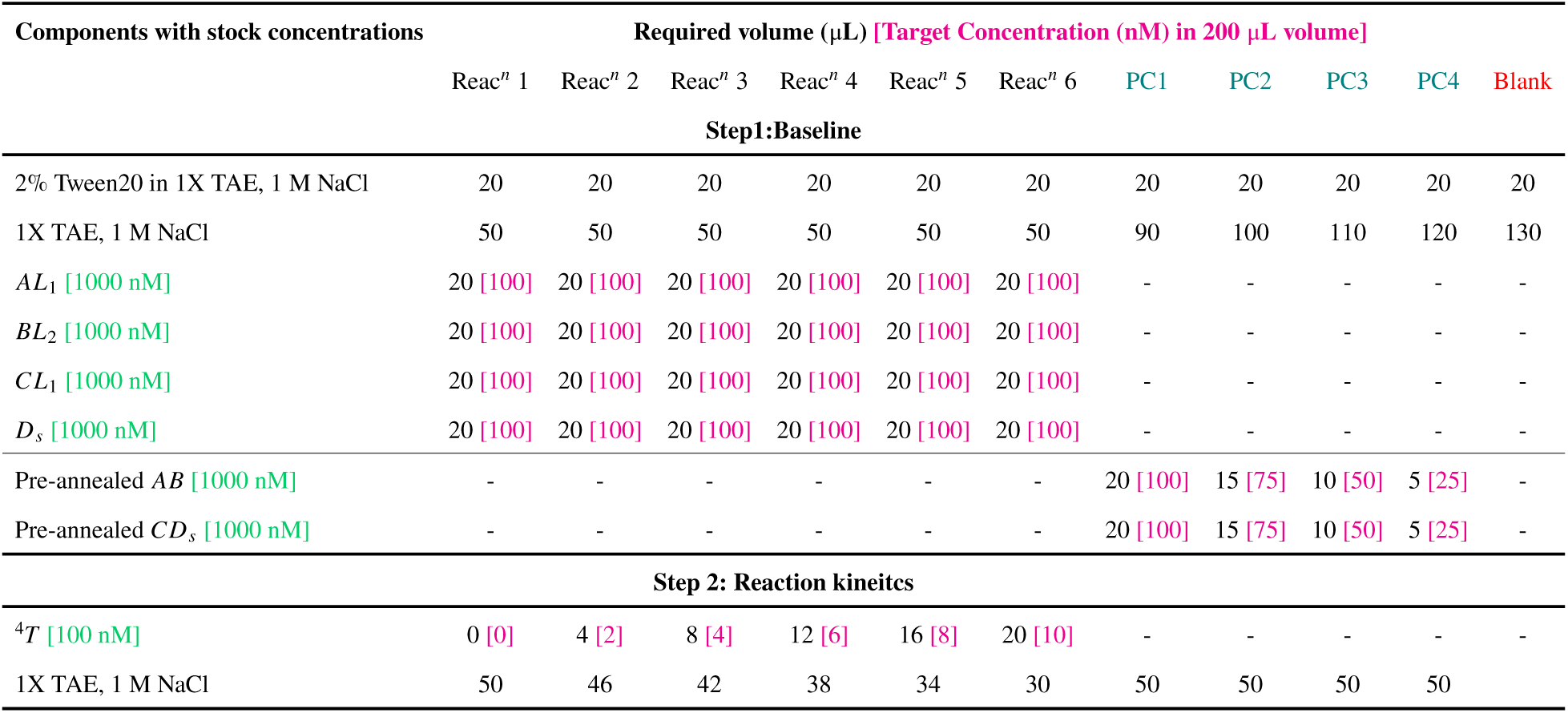
Protocol used for tetramer formation with high monomer concentrations and 0–10 nM template. The table lists stock concentrations and required volumes for each reaction well (Reac*^n^*), positive control (PC), and blank. Target concentrations are shown in [brackets].

| Components with stock concentrations | Required volume (μL) [Target Concentration (nM) in 200 μL volume] |  |  |  |  |  |  |  |  |  |  |
| --- | --- | --- | --- | --- | --- | --- | --- | --- | --- | --- | --- |
|  | Reac<sup>n</sup> 1 | Reac<sup>n</sup> 2 | Reac<sup>n</sup> 3 | Reac<sup>n</sup> 4 | Reac<sup>n</sup> 5 | Reac<sup>n</sup> 6 | PC1 | PC2 | PC3 | PC4 | Blank |
| Step1:Baseline |  |  |  |  |  |  |  |  |  |  |  |
| 2% Tween20 in 1X TAE, 1 M NaCl | 20 | 20 | 20 | 20 | 20 | 20 | 20 | 20 | 20 | 20 | 20 |
| 1X TAE, 1 M NaCl | 50 | 50 | 50 | 50 | 50 | 50 | 90 | 100 | 110 | 120 | 130 |
| *AL*<sub>1</sub> [1000 nM] | 20 [100] | 20 [100] | 20 [100] | 20 [100] | 20 [100] | 20 [100] | - | - | - | - | - |
| *BL*<sub>2</sub> [1000 nM] | 20 [100] | 20 [100] | 20 [100] | 20 [100] | 20 [100] | 20 [100] | - | - | - | - | - |
| *CL*<sub>1</sub> [1000 nM] | 20 [100] | 20 [100] | 20 [100] | 20 [100] | 20 [100] | 20 [100] | - | - | - | - | - |
| *D*<sub>s</sub> [1000 nM] | 20 [100] | 20 [100] | 20 [100] | 20 [100] | 20 [100] | 20 [100] | - | - | - | - | - |
| Pre-annealed *AB* [1000 nM] | - | - | - | - | - | - | 20 [100] | 15 [75] | 10 [50] | 5 [25] | - |
| Pre-annealed *CD*<sub>s</sub> [1000 nM] | - | - | - | - | - | - | 20 [100] | 15 [75] | 10 [50] | 5 [25] | - |
| Step 2: Reaction kineites |  |  |  |  |  |  |  |  |  |  |  |
| <sup>4</sup>*T* [100 nM] | 0 [0] | 4 [2] | 8 [4] | 12 [6] | 16 [8] | 20 [10] | - | - | - | - | - |
| 1X TAE, 1 M NaCl | 50 | 46 | 42 | 38 | 34 | 30 | 50 | 50 | 50 | 50 |  |

**Table S12:** Protocol used for pentamer formation with standard monomer concentrations. The table lists stock concentrations and required volumes for each reaction well (Reac*^n^*), positive control (PC), and blank. Target concentrations are shown in [brackets]. The first 10 data points in the initialisation phase of Reac*^n^*1, where the extent of undesired leak reactions is negligible, are used as the negative control baseline for the fluorescence data analysis as described in the methods.

| Components with stock concentrations | Required volume (μL) [Target Concentration (nM) in 200 μL volume] |  |  |  |  |  |  |  |  |  |  |
| --- | --- | --- | --- | --- | --- | --- | --- | --- | --- | --- | --- |
|  | Reac<sup>n</sup> 1 | Reac<sup>n</sup> 2 | Reac<sup>n</sup> 3 | Reac<sup>n</sup> 4 | Reac<sup>n</sup> 5 | Reac<sup>n</sup> 6 | PC1 | PC2 | PC3 | PC4 | Blank |
| **Step1:Baseline** |  |  |  |  |  |  |  |  |  |  |  |
| 2% Tween20 in 1X TAE, 1M NaCl | 20 | 20 | 20 | 20 | 20 | 20 | 20 | 20 | 20 | 20 | 20 |
| 1X TAE, 1M NaCl | 80 | 80 | 80 | 80 | 80 | 80 | 100 | 109 | 115 | 121 | 130 |
| *AL*<sub>1</sub> [200 nM] | 10 [10] | 10 [10] | 10 [10] | 10 [10] | 10 [10] | 10 [10] | - | - | - | - | - |
| *BL*<sub>2</sub> [200 nM] | 10 [10] | 10 [10] | 10 [10] | 10 [10] | 10 [10] | 10 [10] | - | - | - | - | - |
| *CL*<sub>1</sub> [200 nM] | 10 [10] | 10 [10] | 10 [10] | 10 [10] | 10 [10] | 10 [10] | - | - | - | - | - |
| *DL*<sub>2</sub> [200 nM] | 10 [10] | 10 [10] | 10 [10] | 10 [10] | 10 [10] | 10 [10] | - | - | - | - | - |
| *E* [200 nM] | 10 [10] | 10 [10] | 10 [10] | 10 [10] | 10 [10] | 10 [10] | - | - | - | - | - |
| *AB* [200 nM] | - | - | - | - | - | - | 10 [10] | 7 [7] | 5 [5] | 3 [3] | - |
| *DE* [200 nM] | - | - | - | - | - | - | 10 [10] | 7 [7] | 5 [5] | 3 [3] | - |
| *C* [200 nM] | - | - | - | - | - | - | 10 [10] | 7 [7] | 5 [5] | 3 [3] | - |
| **Step 2: Reaction kinetics** |  |  |  |  |  |  |  |  |  |  |  |
| Template [100 nM] | 0 [0] | 4 [2] | 8 [4] | 12 [6] | 16 [8] | 20 [10] | - | - | - | - | - |
| 1X TAE, 1M NaCl | 50 | 46 | 42 | 38 | 34 | 30 | 50 | 50 | 50 | 50 | 50 |

**Table S13:** Protocol used for pentamer formation with high monomer concentrations. The table lists stock concentrations and required volumes for each reaction well (Reac*^n^*), positive control (PC), and blank. Target concentrations are shown in [brackets]. The first 10 data points in the initialisation phase of Reac*^n^*1, where the extent of undesired leak reactions is negligible, are used as the negative control baseline for the fluorescence data analysis as described in the methods.

| Components with stock concentrations | Required volume (μL) [Target Concentration (nM) in 200 μL volume] |  |  |  |  |  |  |  |  |  |
| --- | --- | --- | --- | --- | --- | --- | --- | --- | --- | --- |
|  | Reac<sup>n</sup> 1 | Reac<sup>n</sup> 2 | Reac<sup>n</sup> 3 | Reac<sup>n</sup> 4 | Reac<sup>n</sup> 5 | PC1 | PC2 | PC3 | PC4 | Blank |
| **Step1:Baseline** |  |  |  |  |  |  |  |  |  |  |
| 2% Tween20 in 1X TAE, 1 M NaCl | 20 | 20 | 20 | 20 | 20 | 20 | 20 | 20 | 20 | 20 |
| 1X TAE, 1 M NaCl | 30 | 30 | 30 | 30 | 30 | 70 | 85 | 100 | 115 | 130 |
| AL<sub>1</sub> [1000 nM] | 20 [100] | 20 [100] | 20 [100] | 20 [100] | 20 [100] | - | - | - | - | - |
| BL<sub>2</sub> [1000 nM] | 20 [100] | 20 [100] | 20 [100] | 20 [100] | 20 [100] | - | - | - | - | - |
| CL<sub>1</sub> [1000 nM] | 20 [100] | 20 [100] | 20 [100] | 20 [100] | 20 [100] | - | - | - | - | - |
| DL<sub>2</sub> [1000 nM] | 20 [100] | 20 [100] | 20 [100] | 20 [100] | 20 [100] | - | - | - | - | - |
| E [1000 nM] | 20 [100] | 20 [100] | 20 [100] | 20 [100] | 20 [100] | - | - | - | - | - |
| AB [1000 nM] | - | - | - | - | - | 20 [100] | 15 [75] | 10 [50] | 5 [25] | - |
| DE [1000 nM] | - | - | - | - | - | 20 [100] | 15 [75] | 10 [50] | 5 [25] | - |
| C [1000 nM] | - | - | - | - | - | 20 [100] | 15 [75] | 10 [50] | 5 [25] | - |
| **Step 2: Reaction kinetics** |  |  |  |  |  |  |  |  |  |  |
| Template [100 nM] | 0 [0] | 4 [2] | 6 [3] | 8 [4] | 10 [5] | - | - | - | - | - |
| 1X TAE, 1 M NaCl | 50 | 46 | 44 | 42 | 40 | 50 | 50 | 50 | 50 | 50 |

**Table S14:**
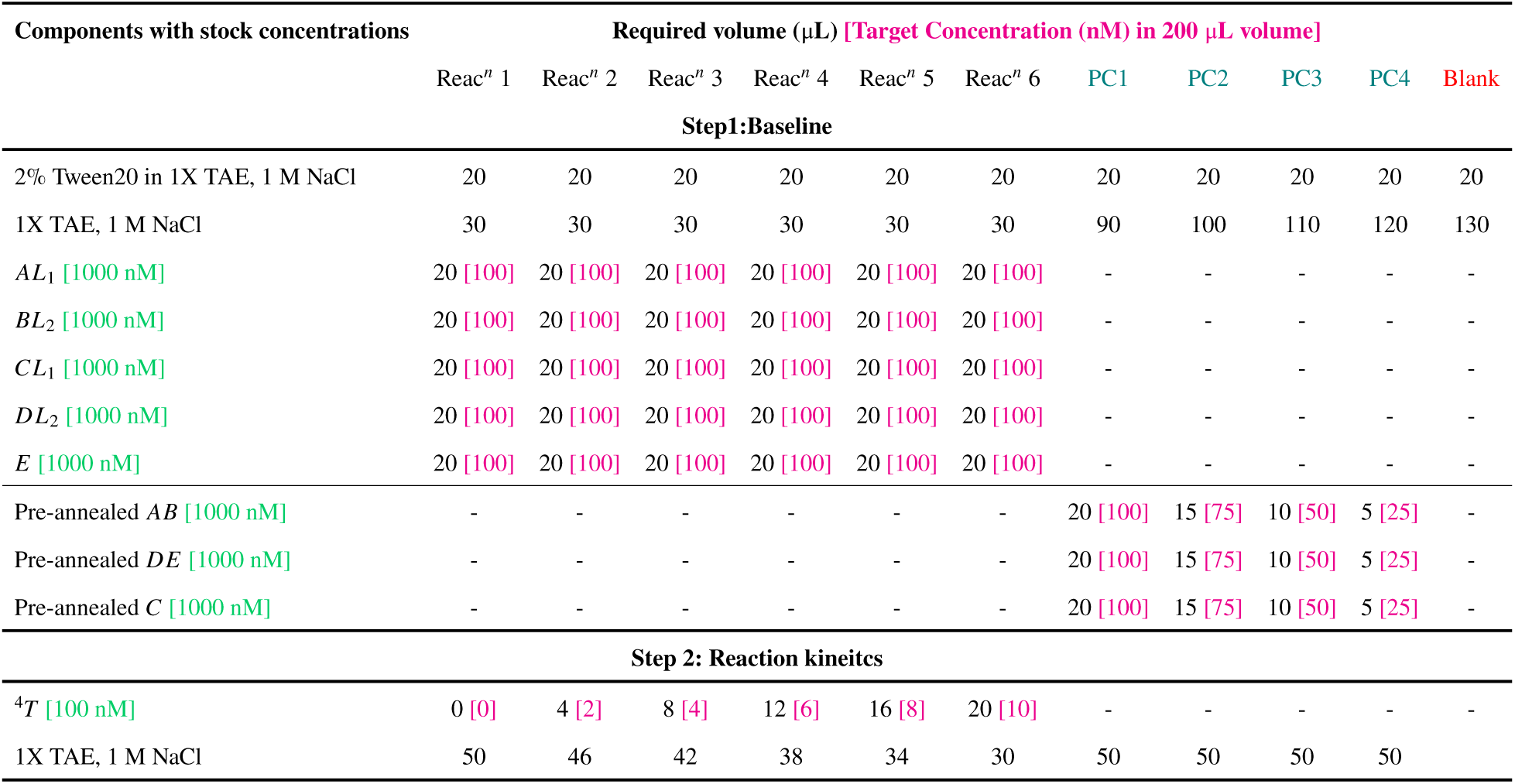
Protocol used for pentamer formation with high monomer concentrations and 0–10 nM template. The table lists stock concentrations and required volumes for each reaction well (Reac*^n^*), positive control (PC), and blank. Target concentrations are shown in [brackets].

**Table S15:** Protocol used for testing global specificity of the templating process. The table lists stock concentrations and required volumes for each reaction well and blank conditions. Target concentrations are shown in [brackets]. The necessary controls are listed in the next page. The first 10 data points in the initialisation phase of Reac*^n^*1, where the extent of undesired leak reactions is negligible, are used as the negative control baseline for the fluorescence data analysis as described in the methods.

| Components with stock concentrations | Required volume (μL) [Target Concentration (nM) in 200 μL volume] |  |  |  |  |  |  |  |  |
| --- | --- | --- | --- | --- | --- | --- | --- | --- | --- |
|  | Reac<sup>n</sup> 1 | Reac<sup>n</sup> 2 | Reac<sup>n</sup> 3 | Reac<sup>n</sup> 4 | Reac<sup>n</sup> 5 | Reac<sup>n</sup> 6 | Reac<sup>n</sup> 7 | Reac<sup>n</sup> 8 | Blank |
| Step1:Baseline |  |  |  |  |  |  |  |  |  |
| 2% Tween20 in 1X TAE, 1M NaCl | 20 | 20 | 20 | 20 | 20 | 20 | 20 | 20 | 20 |
| 1X TAE, 1M NaCl | 50 | 50 | 50 | 50 | 50 | 50 | 50 | 50 | 130 |
| AL<sub>1</sub> [200 nM] | 10 [10] | 10 [10] | 10 [10] | 10 [10] | 10 [10] | 10 [10] | 10 [10] | 10 [10] | 0 [0] |
| BL<sub>2</sub> [200 nM] | 10 [10] | 10 [10] | 10 [10] | 10 [10] | 10 [10] | 10 [10] | 10 [10] | 10 [10] | 0 [0] |
| CL<sub>1</sub> [200 nM] | 10 [10] | 10 [10] | 10 [10] | 10 [10] | 10 [10] | 10 [10] | 10 [10] | 10 [10] | 0 [0] |
| DL<sub>1</sub> [200 nM] | 10 [10] | 10 [10] | 10 [10] | 10 [10] | 10 [10] | 10 [10] | 10 [10] | 10 [10] | 0 [0] |
| E – vanilla [200 nM] | 10 [10] | 10 [10] | 10 [10] | 10 [10] | 10 [10] | 10 [10] | 10 [10] | 10 [10] | 0 [0] |
| AL<sub>1</sub> [200 nM] | 10 [10] | 10 [10] | 10 [10] | 10 [10] | 10 [10] | 10 [10] | 10 [10] | 10 [10] | 0 [0] |
| Cs [200 nM] | 10 [10] | 10 [10] | 10 [10] | 10 [10] | 10 [10] | 10 [10] | 10 [10] | 10 [10] | 0 [0] |
| Ds [200 nM] | 10 [10] | 10 [10] | 10 [10] | 10 [10] | 10 [10] | 10 [10] | 10 [10] | 10 [10] | 0 [0] |
| Step 2: Reaction kinetics, triggered by templates | - | $T_{AD_s}$ | ${}^3T_{ABC_s}$ | ${}^3T_{\bar{A}DE}$ | ${}^4T$ | ${}^5T$ | ${}^5T_{-B}$ | ${}^5T_{-D}$ | - |
| Templates [100 nM] | 0 [0] | 4 [2] | 4 [2] | 4 [2] | 4 [2] | 4 [2] | 4 [2] | 4 [2] | 0 [0] |
| 1X TAE, 1M NaCl | 50 | 46 | 46 | 46 | 46 | 46 | 46 | 46 | 50 |

| Components with stock concentrations | Required volume (μL) [Target Concentration (nM) in 200 μL volume] |  |  |  |
| --- | --- | --- | --- | --- |
|  | PC1 | PC 2 | PC 3 | PC 4 |
|  | ABCDE / ABCBE / ADCDE |  |  |  |
| 2% Tween20 in 1X TAE, 1M NaCl | 20 | 20 | 20 | 20 |
| 1X TAE, 1M NaCl | 90 | 99 | 105 | 111 |
| Pre-annealed AB/AD [200 nM] | 10 [10] | 7 [7] | 5 [5] | 3 [3] |
| Pre-annealed DE-vanilla/BE-vanilla [200 nM] | 10 [10] | 7 [7] | 5 [5] | 3 [3] |
| C [200 nM] | 10 [10] | 7 [7] | 5 [5] | 3 [3] |
| D<sub>s</sub> [200 nM] | 10 [10] | 10 [10] | 10 [10] | 10 [10] |
|  | ABCDs |  |  |  |
| 2% Tween20 in 1X TAE, 1M NaCl | 20 | 20 | 20 | 20 |
| 1X TAE, 1M NaCl | 100 | 106 | 110 | 114 |
| Pre-annealed AB [200 nM] | 10 [10] | 7 [7] | 5 [5] | 3 [3] |
| Pre-annealed CDs [200 nM] | 10 [10] | 7 [7] | 5 [5] | 3 [3] |
| D<sub>s</sub> [200 nM] | 10 [10] | 10 [10] | 10 [10] | 10 [10] |
|  | ADs / ABCs / ADE |  |  |  |
| 2% Tween20 in 1X TAE, 1M NaCl | 20 | 20 | 20 | 20 |
| 1X TAE, 1M NaCl | 110 | 113 | 115 | 117 |
| Pre-annealed ADs/ABCs/ADE-vanilla [200 nM] | 10 [10] | 7 [7] | 5 [5] | 3 [3] |
| D<sub>s</sub> [200 nM] | 10 [10] | 10 [10] | 10 [10] | 10 [10] |

**Table S16:**
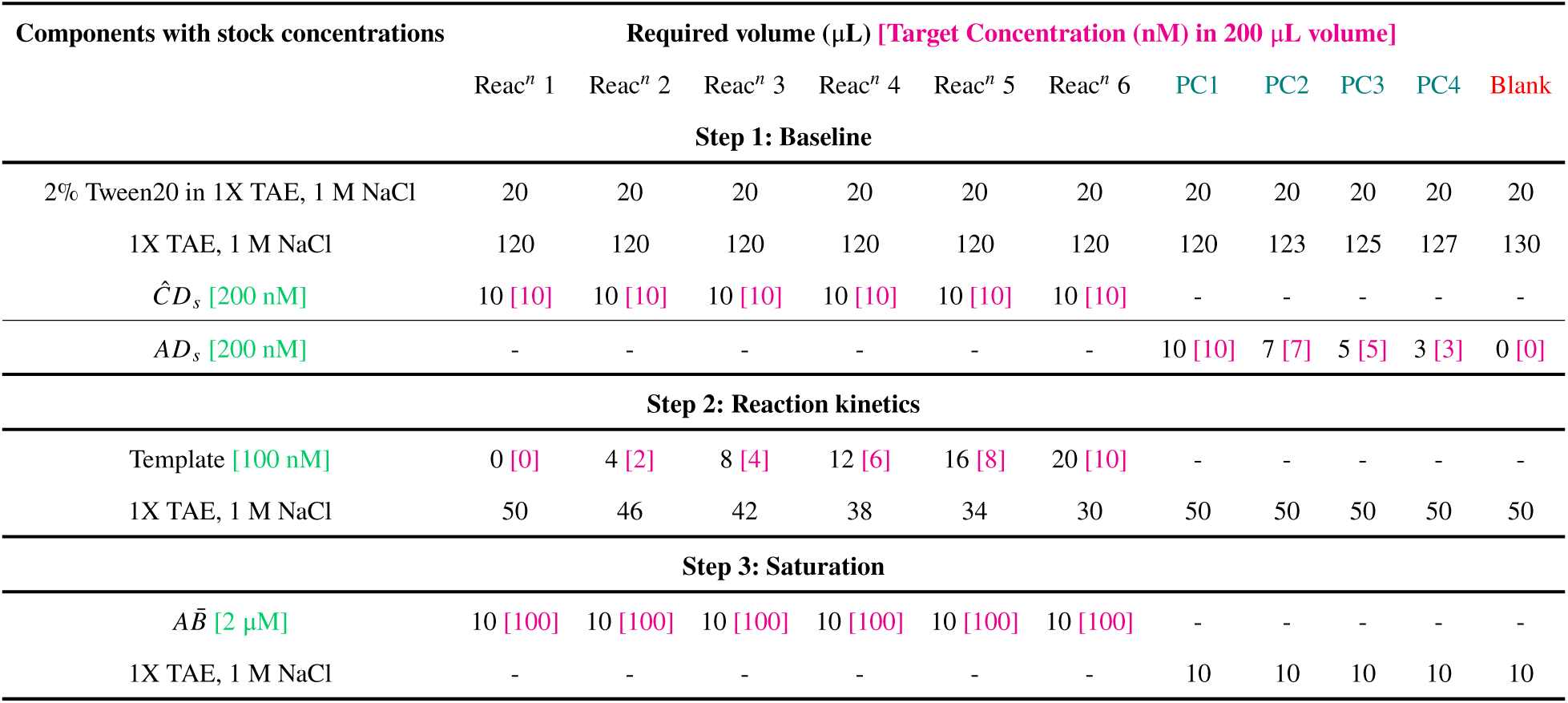
Protocols used for the experiments to test the stability of the tetrameric products. The table shows the concentrations of the stock solutions used, and the required amounts for the reactions. Reac*^n^* denotes reaction wells, PC denotes positive controls.

| Components with stock concentrations | Required volume ($\mu\text{L}$) [Target Concentration (nM) in 200 $\mu\text{L}$ volume] | | | | | | | | | | |
| --- | --- | --- | --- | --- | --- | --- | --- | --- | --- | --- | --- |
| | $\text{Reac}^n$ 1 | $\text{Reac}^n$ 2 | $\text{Reac}^n$ 3 | $\text{Reac}^n$ 4 | $\text{Reac}^n$ 5 | $\text{Reac}^n$ 6 | PC1 | PC2 | PC3 | PC4 | Blank |
| Step 1: Baseline |  |  |  |  |  |  |  |  |  |  |  |
| 2% Tween20 in 1X TAE, 1 M NaCl | 20 | 20 | 20 | 20 | 20 | 20 | 20 | 20 | 20 | 20 | 20 |
| 1X TAE, 1 M NaCl | 120 | 120 | 120 | 120 | 120 | 120 | 120 | 123 | 125 | 127 | 130 |
| $\hat{C}D_s$ [200 nM] | 10 [10] | 10 [10] | 10 [10] | 10 [10] | 10 [10] | 10 [10] | - | - | - | - | - |
| $AD_s$ [200 nM] | - | - | - | - | - | - | 10 [10] | 7 [7] | 5 [5] | 3 [3] | 0 [0] |
| Step 2: Reaction kinetics |  |  |  |  |  |  |  |  |  |  |  |
| Template [100 nM] | 0 [0] | 4 [2] | 8 [4] | 12 [6] | 16 [8] | 20 [10] | - | - | - | - | - |
| 1X TAE, 1 M NaCl | 50 | 46 | 42 | 38 | 34 | 30 | 50 | 50 | 50 | 50 | 50 |
| Step 3: Saturation |  |  |  |  |  |  |  |  |  |  |  |
| $A\bar{B}$ [2 $\mu\text{M}$] | 10 [100] | 10 [100] | 10 [100] | 10 [100] | 10 [100] | 10 [100] | - | - | - | - | - |
| 1X TAE, 1 M NaCl | - | - | - | - | - | - | 10 | 10 | 10 | 10 | 10 |

**Table S17:**
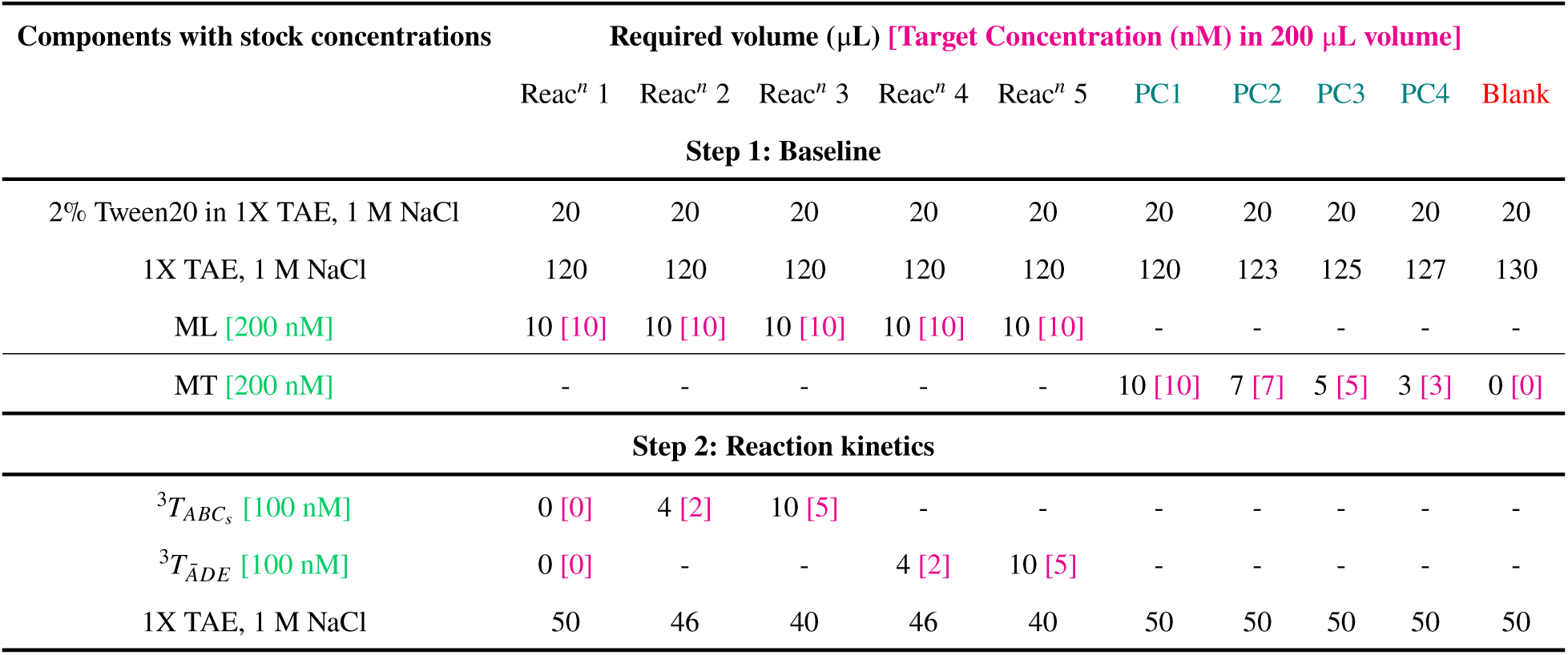
Protocols used for testing template specificity for binding to blocked monomers *via* TMSD. The table shows the concentrations of the stock solutions used, and the required amounts for the TMSD reactions. Reac*^n^* denotes reaction wells, PC denotes positive controls. Target concentrations are shown in [brackets]. The first 10 data points in the initialisation phase of Reac*^n^*1, where the extent of undesired leak reactions is negligible, are used as the negative control baseline for the fluorescence data analysis as described in the methods.

**Table S18:** Protocols used for testing template specificity of trimer formation from a mixture of monomers of either set. The table shows the concentrations of the stock solutions used, and the required amounts for the TMSD reactions. Reac*^n^* denotes reaction wells, PC denotes positive controls. Target concentrations are shown in [brackets]. The first 10 data points in the initialisation phase of Reac*^n^*1, where the extent of undesired leak reactions is negligible, are used as the negative control baseline for the fluorescence data analysis as described in the methods.

| Components with stock concentrations | Required volume ($\mu\text{L}$) [Target Concentration (nM) in 200 $\mu\text{L}$ volume] | | | | | | | |
| --- | --- | --- | --- | --- | --- | --- | --- | --- |
| | $\text{Reac}^n 1$ | $\text{Reac}^n 2$ | $\text{Reac}^n 3$ | $\text{Reac}^n 4$ | $\text{Reac}^n 5$ | PC1 | PC2 | Blank |
| **Step 1: Baseline** |  |  |  |  |  |  |  |  |
| 2% Tween20 in 1X TAE, 1 M NaCl | 20 | 20 | 20 | 20 | 20 | 20 | 20 | 20 |
| 1X TAE, 1 M NaCl | 100 | 100 | 100 | 100 | 100 | 120 | 125 | 130 |
| $AL_1 / \bar{A}L_1$ [200 nM] | 10 [10] | 10 [10] | 10 [10] | 10 [10] | 10 [10] | - | - | - |
| $BL_2 / DL_2$ [200 nM] | 10 [10] | 10 [10] | 10 [10] | 10 [10] | 10 [10] | - | - | - |
| $C_s / E$ [200 nM] | 10 [10] | 10 [10] | 10 [10] | 10 [10] | 10 [10] | - | - | - |
| $ABC_S / \bar{A}DE$ [200 nM] | - | - | - | - | - | 10 [10] | 5 [5] | 0 [0] |
| **Step 2: Reaction kinetics** |  |  |  |  |  |  |  |  |
| ${}^3T_{ABC_S}$ [100 nM] | 0 [0] | 4 [2] | 10 [5] | - | - | - | - | - |
| ${}^3T_{\bar{A}DE}$ [100 nM] | 0 [0] | - | - | 4 [2] | 10 [5] | - | - | - |
| 1X TAE, 1 M NaCl | 50 | 46 | 40 | 46 | 40 | 50 | 50 | 50 |

**Table S19:**
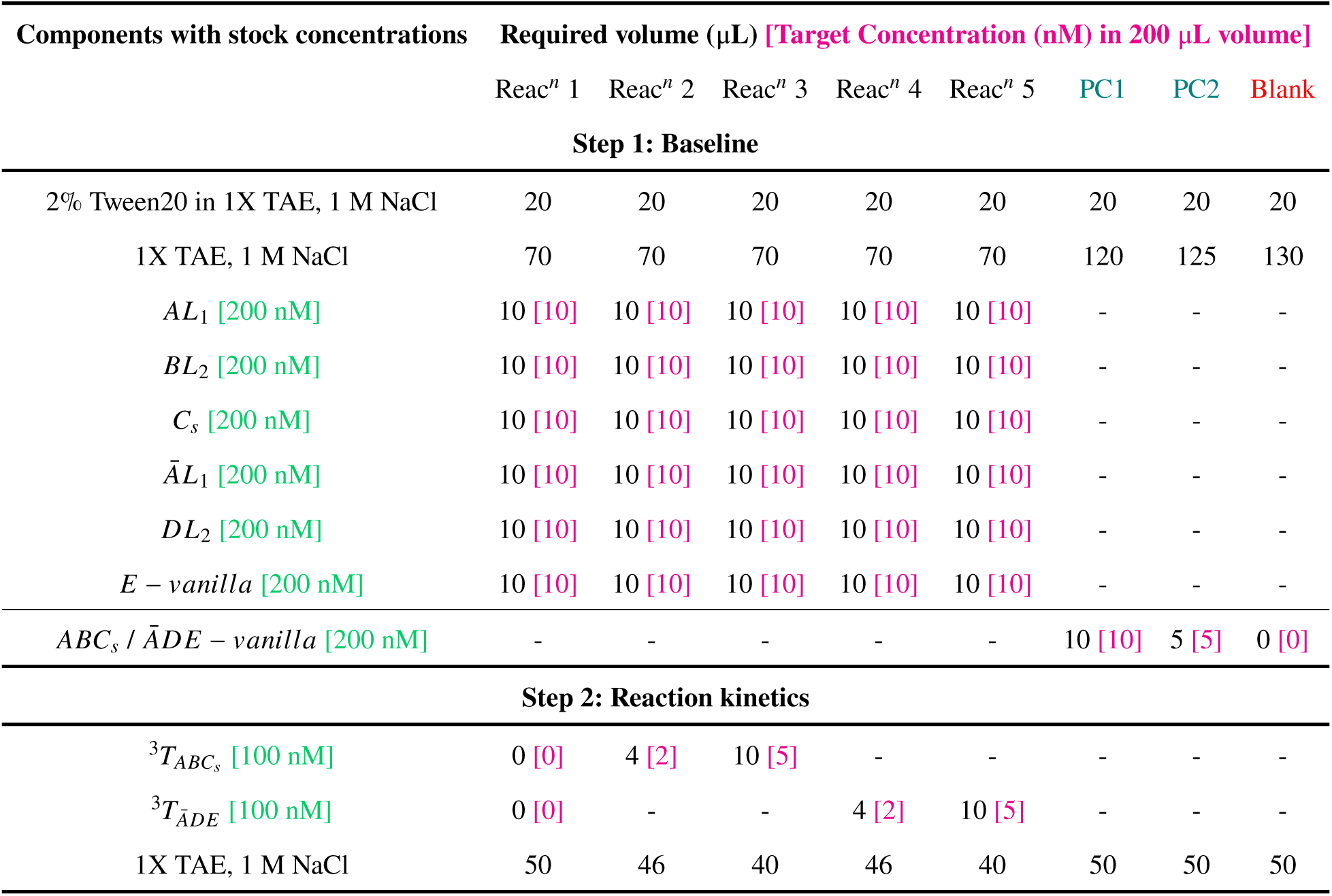
Protocols used for testing template specificity of trimer formation from a mixture of all monomers. The table shows the concentrations of the stock solutions used, and the required amounts for the TMSD reactions. Reac*^n^* denotes reaction wells, PC denotes positive controls. Target concentrations are shown in [brackets]. The first 10 data points in the initialisation phase of Reac*^n^*1, where the extent of undesired leak reactions is negligible, are used as the negative control baseline for the fluorescence data analysis as described in the methods.

**Table S20:** Protocol used for stepwise addition of monomers at standard concentrations for pentamer formation. The table lists stock concentrations and required volumes for each reaction well (Reac*^n^*), positive control (PC), and blank. Target concentrations are shown in [brackets]. Since different components were added at different time points, the experiment could not have a global negative control. We therefore used the first 10 data points of the initialisation phase of the blank well as the negative control in this instance.

| Components with stock concentrations | Reac<sup>n</sup> 2 | Reac<sup>n</sup> 3 | Reac<sup>n</sup> 4 | PC1 | PC2 | PC3 | PC4 | Blank |
| --- | --- | --- | --- | --- | --- | --- | --- | --- |
| **Step1:Baseline** |  |  |  |  |  |  |  |  |
| 2% Tween20 in 1X TAE, 1M NaCl | 20 | 20 | 20 | 20 | 20 | 20 | 20 | 20 |
| 1X TAE, 1M NaCl | 80 | 80 | 80 | 100 | 109 | 115 | 121 | 130 |
| *AL*<sub>1</sub> [200 nM] | 10 [10] | 10 [10] | 10 [10] | - | - | - | - | - |
| Pre-annealed *AB* [200 nM] | - | - | - | 10 [10] | 7 [7] | 5 [5] | 3 [3] | - |
| Pre-annealed *DE* [200 nM] | - | - | - | 10 [10] | 7 [7] | 5 [5] | 3 [3] | - |
| *C* | - | - | - | 10 [10] | 7 [7] | 5 [5] | 3 [3] | - |
| **Step 2: Reaction kinetics** |  |  |  |  |  |  |  |  |
| Template [100 nM] | 4 [2] | 6 [3] | 10 [5] | - | - | - | - | - |
| 1X TAE, 1M NaCl | 46 | 44 | 40 | 50 | 50 | 50 | 50 | 50 |
| **Step 3: Reaction kinetics** |  |  |  |  |  |  |  |  |
| *BL*<sub>2</sub> [200 nM] | 10 [10] | 10 [10] | 10 [10] | - | - | - | - | - |
| **Step 4: Reaction kinetics** |  |  |  |  |  |  |  |  |
| *CL*<sub>1</sub> [200 nM] | 10 [10] | 10 [10] | 10 [10] | - | - | - | - | - |
| **Step 5: Reaction kinetics** |  |  |  |  |  |  |  |  |
| *DL*<sub>2</sub> [200 nM] | 10 [10] | 10 [10] | 10 [10] | - | - | - | - | - |
| **Step 6: Reaction kinetics** |  |  |  |  |  |  |  |  |
| *E* [200 nM] | 10 [10] | 10 [10] | 10 [10] | - | - | - | - | - |

**Table S21:** Protocol used for stepwise addition of monomer at high concentrations for pentamer formation. The table lists stock concentrations and required volumes for each reaction well (Reac*^n^*), positive control (PC), and blank. Target concentrations are shown in [brackets]. Since different components were added at different time points, the experiment could not have a global negative control. We therefore used the first 10 data points of the initialisation phase of the blank well as the negative ocntrol in this instance.

| Components with stock concentrations | Reac<sup>n</sup> 1 | Reac<sup>n</sup> 2 | Reac<sup>n</sup> 3 | Reac<sup>n</sup> 4 | Reac<sup>n</sup> 5 | PC1 | PC2 | PC3 | PC4 | Blank |
| --- | --- | --- | --- | --- | --- | --- | --- | --- | --- | --- |
| **Step 1: Baseline** |  |  |  |  |  |  |  |  |  |  |
| 2% Tween20 in 1X TAE, 1M NaCl | 20 | 20 | 20 | 20 | 20 | 20 | 20 | 20 | 20 | 20 |
| 1X TAE, 1M NaCl | 30 | 30 | 30 | 30 | 30 | 70 | 85 | 100 | 115 | 130 |
| AL<sub>1</sub> [200 nM] | 20 [100] | 20 [100] | 20 [100] | 20 [100] | 20 [100] | - | - | - | - | - |
| Pre-annealed AB [1000 nM] | - | - | - | - | - | 20 [100] | 15 [75] | 10 [50] | 5 [25] | - |
| Pre-annealed DE [1000 nM] | - | - | - | - | - | 20 [100] | 15 [75] | 10 [50] | 5 [25] | - |
| C | - | - | - | - | - | 20 [100] | 15 [75] | 10 [50] | 5 [25] | - |
| **Step 2: Reaction kinetics** |  |  |  |  |  |  |  |  |  |  |
| Template [100 nM] | - | 4 [2] | 6 [3] | 8 [4] | 10 [5] | - | - | - | - | - |
| 1X TAE, 1M NaCl | 50 | 46 | 44 | 42 | 40 | 50 | 50 | 50 | 50 | 50 |
| **Step 3: Reaction kinetics** |  |  |  |  |  |  |  |  |  |  |
| BL<sub>2</sub> [1000 nM] | 20 [100] | 20 [100] | 20 [100] | 20 [100] | 20 [100] | - | - | - | - | - |
| **Step 4: Reaction kinetics** |  |  |  |  |  |  |  |  |  |  |
| CL<sub>1</sub> [1000 nM] | 20 [100] | 20 [100] | 20 [100] | 20 [100] | 20 [100] | - | - | - | - | - |
| **Step 5: Reaction kinetics** |  |  |  |  |  |  |  |  |  |  |
| DL<sub>2</sub> [1000 nM] | 20 [100] | 20 [100] | 20 [100] | 20 [100] | 20 [100] | - | - | - | - | - |
| **Step 6: Reaction kinetics** |  |  |  |  |  |  |  |  |  |  |
| E [1000 nM] | 20 [100] | 20 [100] | 20 [100] | 20 [100] | 20 [100] | - | - | - | - | - |

**Table S22:** Protocol used for a set of reactions with increasing numbers of monomers present. The table lists stock concentrations and required volumes for each reaction well (Reac*^n^*), positive control (PC), and blank. Target concentrations are shown in [brackets]. Since different reactions had different components, the experiment could not have a global negative control. So we used the The first 10 data points of the initialisation phase of the blank well as the negative ocntrol in this instance.

| Components with stock concentrations | Required volume (μL) [Target Concentration (nM) in 200 μL volume] |  |  |  |  |  |  |  |  |  |
| --- | --- | --- | --- | --- | --- | --- | --- | --- | --- | --- |
|  | Reac<sup>n</sup> 1 | Reac<sup>n</sup> 2 | Reac<sup>n</sup> 3 | Reac<sup>n</sup> 4 | Reac<sup>n</sup> 5 | PC1 | PC2 | PC3 | PC4 | Blank |
| **Step1:Baseline** |  |  |  |  |  |  |  |  |  |  |
| 2% Tween20 in 1X TAE, 1M NaCl | 20 | 20 | 20 | 20 | 20 | 20 | 20 | 20 | 20 | 20 |
| 1X TAE, 1M NaCl | 120 | 110 | 100 | 90 | 80 | 100 | 109 | 115 | 121 | 130 |
| AL<sub>1</sub> [200 nM] | 10 [10] | 10 [10] | 10 [10] | 10 [10] | 10 [10] | - | - | - | - | - |
| BL<sub>2</sub> [200 nM] | - | 10 [10] | 10 [10] | 10 [10] | 10 [10] | - | - | - | - | - |
| CL<sub>1</sub> [200 nM] | - | - | 10 [10] | 10 [10] | 10 [10] | - | - | - | - | - |
| DL<sub>2</sub> [200 nM] | - | - | - | 10 [10] | 10 [10] | - | - | - | - | - |
| E / $\bar{E}$ [200 nM] | - | - | - | - | 10 [10] | - | - | - | - | - |
| **Positive Controls: A<sup>5</sup>T/ B<sup>5</sup>T/ C<sup>5</sup>T/ D<sup>5</sup>T** |  |  |  |  |  |  |  |  |  |  |
| M<sup>5</sup>T [200 nM] | - | - | - | - | - | 10 [10] | 7 [7] | 5 [5] | 3 [3] | - |
| **Step 2: Reaction kinetics** |  |  |  |  |  |  |  |  |  |  |
| Template [100 nM] | 8 [4] | 8 [4] | 8 [4] | 8 [4] | 8 [4] | - | - | - | - | - |
| 1X TAE, 1M NaCl | 42 | 42 | 42 | 42 | 42 | 50 | 50 | 50 | 50 | 50 |

**Table S23:** Protocol used for testing the effect of systematic omission of each monomer in pentamer formation with standard monomer concentrations. The table lists stock concentrations and required volumes for each reaction well (Reac*^n^*), positive control (PC), and blank. Target concentrations are shown in [brackets].

| Components with stock concentrations | Required volume (μL) [Target Concentration (nM) in 200 μL volume] |  |  |  |  |  |  |  |  |  |
| --- | --- | --- | --- | --- | --- | --- | --- | --- | --- | --- |
|  | Reac<sup>n</sup> 1 | Reac<sup>n</sup> 2 | Reac<sup>n</sup> 3 | Reac<sup>n</sup> 4 | Reac<sup>n</sup> 5 | PC1 | PC2 | PC3 | PC4 | Blank |
| Step1:Baseline |  |  |  |  |  |  |  |  |  |  |
| 2% Tween20 in 1X TAE, 1M NaCl | 20 | 20 | 20 | 20 | 20 | 20 | 20 | 20 | 20 | 20 |
| 1X TAE, 1M NaCl | 90 | 90 | 90 | 90 | 90 | 100 | 109 | 115 | 121 | 130 |
| AL<sub>1</sub> [200 nM] | - | 10 [10] | 10 [10] | 10 [10] | 10 [10] | - | - | - | - | - |
| BL<sub>2</sub> [200 nM] | 10 [10] | - | 10 [10] | 10 [10] | 10 [10] | - | - | - | - | - |
| CL<sub>1</sub> [200 nM] | 10 [10] | 10 [10] | - | 10 [10] | 10 [10] | - | - | - | - | - |
| DL<sub>2</sub> [200 nM] | 10 [10] | 10 [10] | 10 [10] | - | 10 [10] | - | - | - | - | - |
| E / $\bar{E}$ [200 nM] | 10 [10] | 10 [10] | 10 [10] | 10 [10] | - | - | - | - | - | - |
| AB [200 nM] | - | - | - | - | - | 10 [10] | 7 [7] | 5 [5] | 3 [3] | - |
| DE [200 nM] | - | - | - | - | - | 10 [10] | 7 [7] | 5 [5] | 3 [3] | - |
| C [200 nM] | - | - | - | - | - | 10 [10] | 7 [7] | 5 [5] | 3 [3] | - |
| Step 2: Reaction kinetics |  |  |  |  |  |  |  |  |  |  |
| Template [100 nM] | 8 [4] | 8 [4] | 8 [4] | 8 [4] | 8 [4] | - | - | - | - | - |
| 1X TAE, 1M NaCl | 42 | 42 | 42 | 42 | 42 | 50 | 50 | 50 | 50 | 50 |

**Table S24:** Fluorescence checklist for band identity validation. The table indicates the theoretical presence or absence of different fluorescence signals in the complexes. This table can be consulted for cross-checking the identities of the bands of concern in the fluorescent gel images by scrutinising the individual scans. The colours of the channels are used for making the pseudocolour images.

| Figure | Complex | Cy2 | Cy3 | Cy5 |
| --- | --- | --- | --- | --- |
| 2C / 3B / 5B / S25 / S27 | $ABC_s$ | – | – | Y |
| 2C / 3B / 5B / S25 / S27 | $ABT_{ABC_s}$ | – | – | Y |
| 2C / 3B / 5B / S25 / S27 | $AABT_{ABC_s}$ | – | – | Y |
| 2C / 3B / 5B / S25 / S27 | $AT_{ABC_s}$ | – | – | Y |
| 2C / 3B / 5B / S26 / S27 | $\bar{A}\bar{A}DT_{\bar{A}DE}$ | Y | Y | – |
| 2C / 3B / 5B / S26 / S27 | $\bar{A}DT_{\bar{A}DE}$ | Y | Y | – |
| 2C / 3B / 5B / S26 / S27 | $DT_{\bar{A}DE}$ | – | Y | – |
| 2C / 3B / 5B / S26 / S27 | $\bar{A}DE$ | Y | Y | – |
| 2C / 3B / 5B / S26 / S27 | $\bar{A}T_{\bar{A}DE}$ | Y | – | – |
| 2C / 3B / 5B / S26 / S27 | $DE$ | – | Y | – |
| S28 | ${}^4P$ | Y | – | Y |
| S28 | $A^4T$ | – | – | Y |
| S28 | $CD_s$ | Y | – | – |
| S28 | $AB$ | – | – | Y |
| S28 | $BC$ | Y | – | – |
| S28 | $AD_s$ | Y | – | Y |
| S28 | Monomers | Y | – | Y |
| 4E, F / S30 / S31 / S32 / S33 | ${}^5P$ | Y | Y | Y |
| 4E, F / S30 / S31 / S32 / S33 | $A^5T$ | – | – | Y |
| 4E, F / S30 / S31 / S32 / S33 | $ADE$ | Y | Y | Y |
| 4E, F / S30 / S31 / S32 / S33 | $ABE$ | Y | – | Y |
| 4E, F / S30 / S31 / S32 / S33 | $CD$ | Y | Y | – |
| 4E, F / S30 / S31 / S32 / S33 | $BC$ | Y | — | — |
| 4E, F / S30 / S31 / S32 / S33 | $E$ | Y | — | — |
| S36 | ${}^5\hat{P}$ | Y | Y | — |
| S36 | ${}^4\hat{P}$ | Y | Y | — |
| S36 | $A^5\hat{T}$ | — | Y | — |
| S36 | $A^4\hat{T}$ | — | Y | Y |
| S36 | Trimers | Y | Y | — |
| S36 | $D\hat{E}$ | Y | — | — |
| S36 | $\hat{C}D_s$ | Y | — | — |
| S36 | $AB$ | — | Y | — |
| 5C / S39 | $AD_s$ | Y | — | Y |
| 5C / S39 | $A^2T$ | — | — | Y |
| 5C / S39 | $ABC_s$ | — | — | Y |
| 5C / S39 | $\bar{A}DE$ | Y | Y | — |
| 5C / S39 | $A^4T$ | — | — | Y |
| 5C / S39 | $BC$ | Y | — | — |
| 5C / S39 | $CD$ | Y | Y | — |
| 5C / S39 | ${}^4P$ | Y | — | Y |
| 5C / S39 | ${}^5P$ | Y | Y | Y |
| 5C / S39 | ${}^5P_{-B}$ | Y | Y | Y |
| 5C / S39 | ${}^5P_{-D}$ | Y | — | Y |
| 5C / S39 | $A^5T$ | — | — | Y |
| 5C / S39 | $A^5T_{-D}$ | — | — | Y |

**Table S25:**
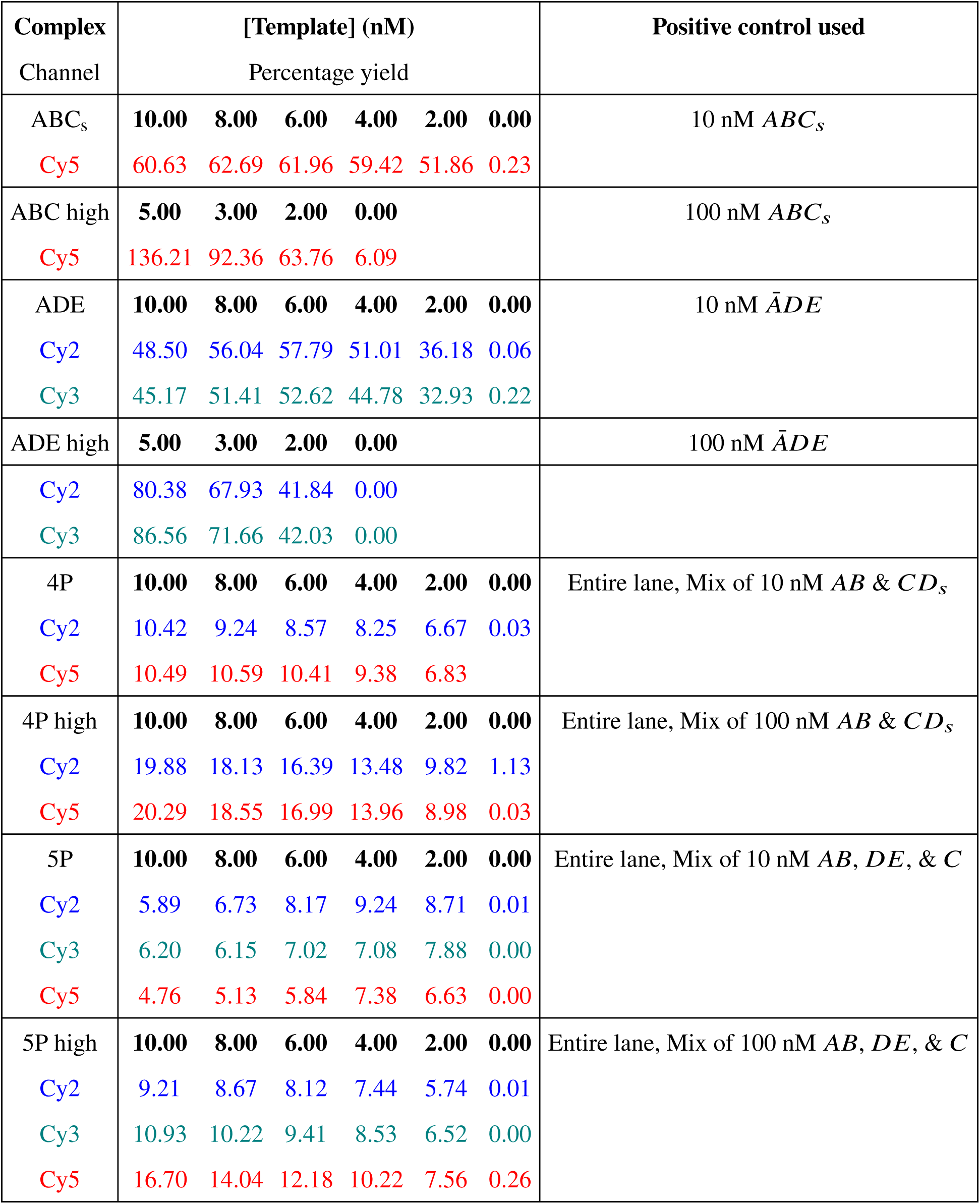
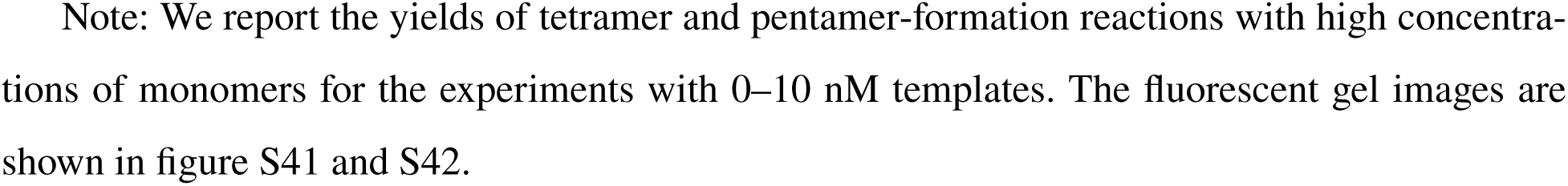
Percentage yield relative to corresponding positive controls. Where the concerned complexes had more than one fluorophore channels, the yields were separately calculated for each channel, and indicated by the names on the concerned channels.

**Table S26:**
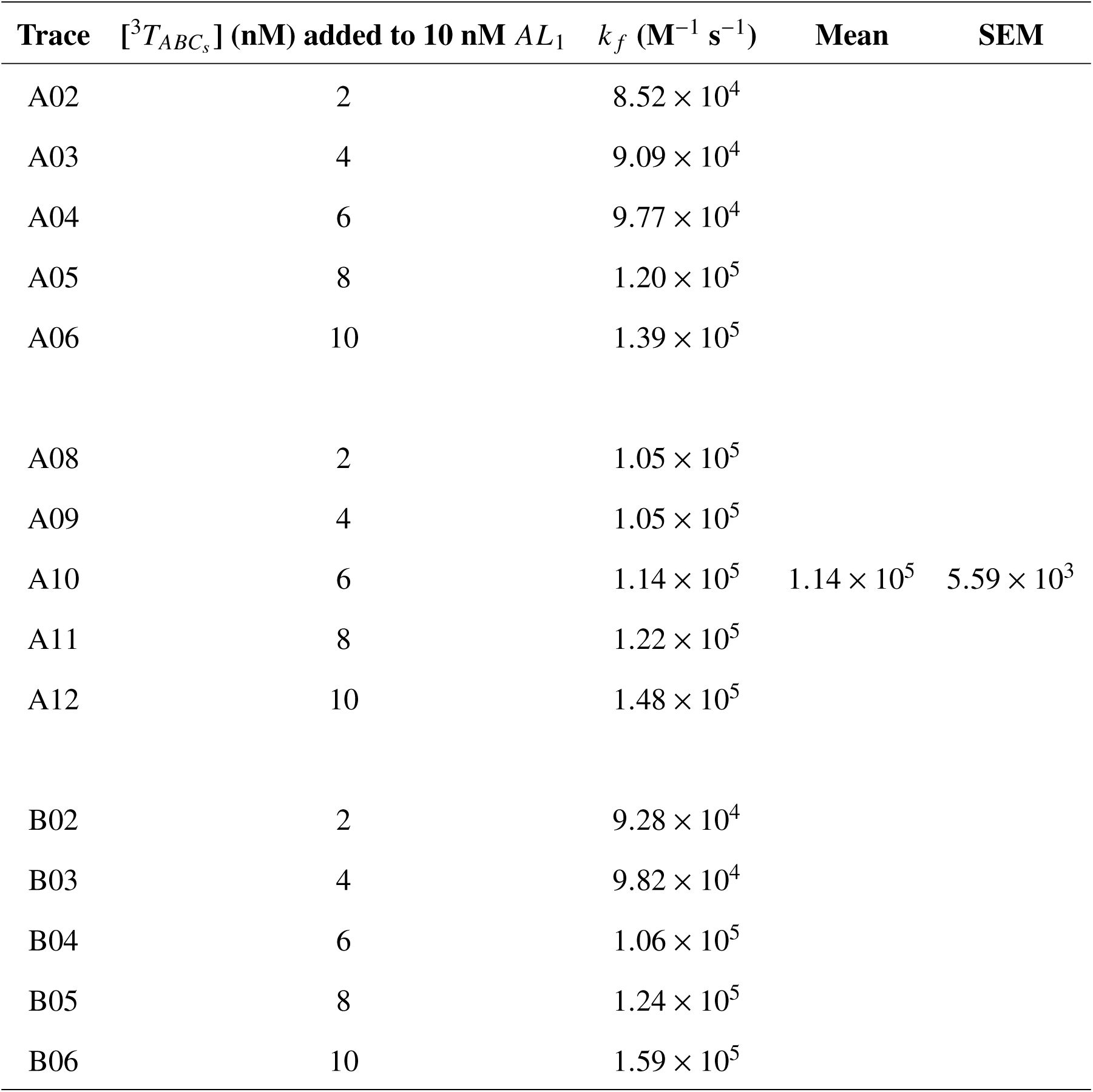
Estimated *k _f_* values for the reaction between *AL*_1_ and ^3^*T_ABCs_*. 15 reaction traces were fitted independently to an ODE model of a second order reaction. Mean and Standard Error of the Mean are reported.

**Table S27:**
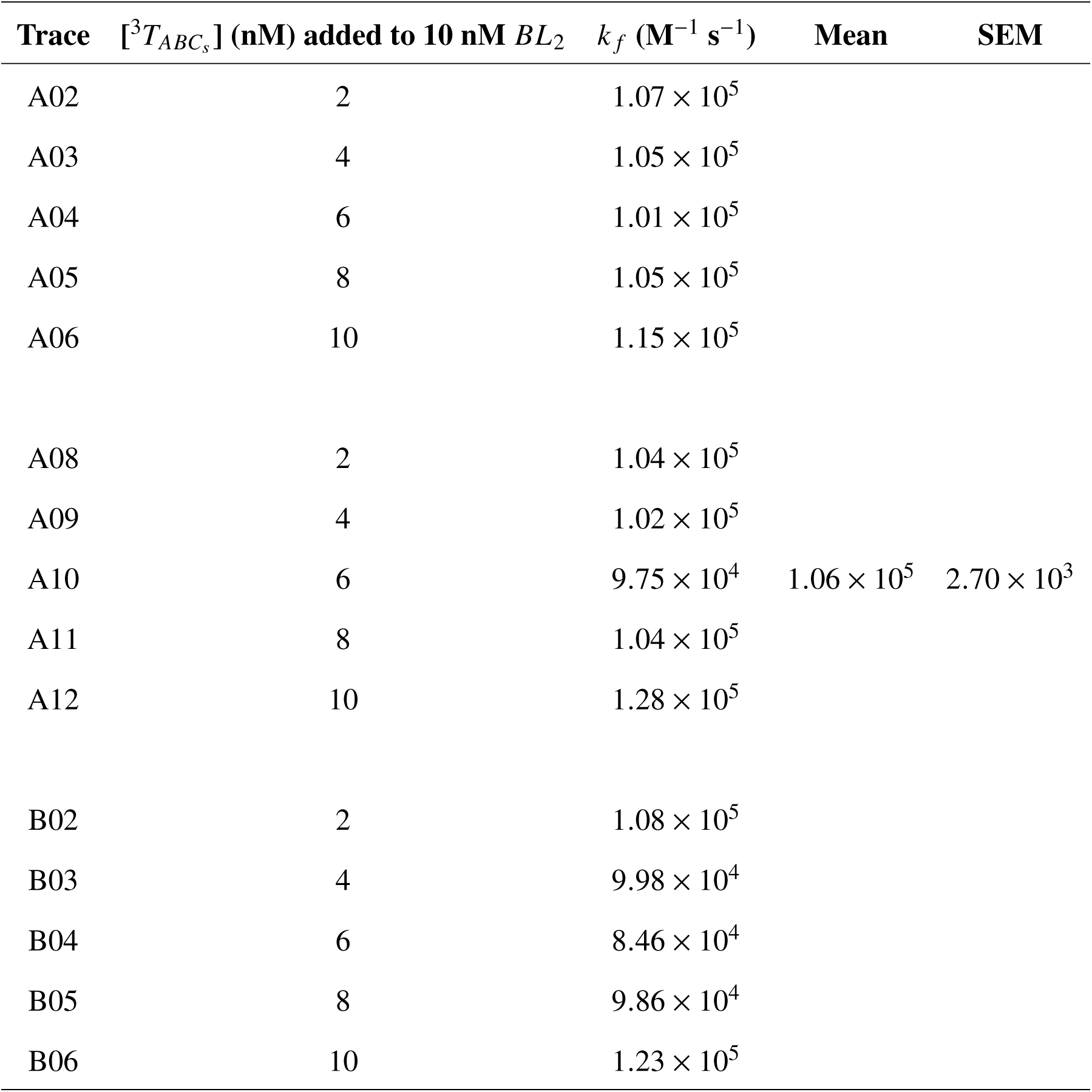
Estimated *k _f_* values for the reaction between *BL*_2_ and *T_ABCs_*. 15 reaction traces were fitted independently to an ODE model of a second order reaction. Mean and Standard Error of the Mean are reported.

| Trace | $[^3T_{ABC_s}]$ (nM) added to 10 nM $BL_2$ | $k_f$ ($M^{-1} s^{-1}$) | Mean | SEM |
| --- | --- | --- | --- | --- |
| A02 | 2 | $1.07 \times 10^5$ | | |
| A03 | 4 | $1.05 \times 10^5$ | | |
| A04 | 6 | $1.01 \times 10^5$ | | |
| A05 | 8 | $1.05 \times 10^5$ | | |
| A06 | 10 | $1.15 \times 10^5$ | | |
| A08 | 2 | $1.04 \times 10^5$ | | |
| A09 | 4 | $1.02 \times 10^5$ | | |
| A10 | 6 | $9.75 \times 10^4$ | $1.06 \times 10^5$ | $2.70 \times 10^3$ |
| A11 | 8 | $1.04 \times 10^5$ | | |
| A12 | 10 | $1.28 \times 10^5$ | | |
| B02 | 2 | $1.08 \times 10^5$ | | |
| B03 | 4 | $9.98 \times 10^4$ | | |
| B04 | 6 | $8.46 \times 10^4$ | | |
| B05 | 8 | $9.86 \times 10^4$ | | |
| B06 | 10 | $1.23 \times 10^5$ | | |

**Table S28:** Estimated *k _f_* values for the reaction between *ĀL*_1_ and *T_ĀDE_*. 15 reaction traces were fitted independently to an ODE model of a second order reaction. Mean and Standard Error of the Mean are reported.

| Trace | $[^3T_{\bar{A}DE}]$ (nM) added to 10 nM $\bar{A}L_1$ | $k_f$ ($\text{M}^{-1} \text{s}^{-1}$) | Mean | SEM |
| --- | --- | --- | --- | --- |
| E02 | 2 | $9.75 \times 10^4$ | | |
| E03 | 4 | $9.54 \times 10^4$ | | |
| E04 | 6 | $9.24 \times 10^4$ | | |
| E05 | 8 | $9.60 \times 10^4$ | | |
| E06 | 10 | $1.03 \times 10^5$ | | |
| E08 | 2 | $9.31 \times 10^4$ | | |
| E09 | 4 | $9.60 \times 10^4$ | | |
| E10 | 6 | $9.61 \times 10^4$ | $9.63 \times 10^4$ | $9.62 \times 10^2$ |
| E11 | 8 | $9.25 \times 10^4$ | | |
| E12 | 10 | $9.89 \times 10^4$ | | |
| F02 | 2 | $9.76 \times 10^4$ | | |
| F03 | 4 | $9.62 \times 10^4$ | | |
| F04 | 6 | $9.53 \times 10^4$ | | |
| F05 | 8 | $9.01 \times 10^4$ | | |
| F06 | 10 | $1.04 \times 10^5$ | | |

**Table S29:** Estimated *k _f_* values for the reaction between *DL*_2_ and *T_Ā_*_*DE*_. 15 reaction traces were fitted independently to an ODE model of a second order reaction. Mean and Standard Error of the Mean are reported.

| Trace | $[^3T_{\bar{A}DE}]$ (nM) added to 10 nM $DL_2$ | $k_f$ ($M^{-1} s^{-1}$) | Mean | SEM |
| --- | --- | --- | --- | --- |
| E02 | 2 | $5.34 \times 10^4$ | | |
| E03 | 4 | $5.62 \times 10^4$ | | |
| E04 | 6 | $5.70 \times 10^4$ | | |
| E05 | 8 | $5.40 \times 10^4$ | | |
| E06 | 10 | $5.92 \times 10^4$ | | |
| E08 | 2 | $5.24 \times 10^4$ | | |
| E09 | 4 | $5.38 \times 10^4$ | | |
| E10 | 6 | $5.53 \times 10^4$ | $5.49 \times 10^4$ | $5.57 \times 10^2$ |
| E11 | 8 | $5.55 \times 10^4$ | | |
| E12 | 10 | $5.38 \times 10^4$ | | |
| F02 | 2 | $5.59 \times 10^4$ | | |
| F03 | 4 | $5.40 \times 10^4$ | | |
| F04 | 6 | $5.22 \times 10^4$ | | |
| F05 | 8 | $5.11 \times 10^4$ | | |
| F06 | 10 | $6.03 \times 10^4$ | | |

**Table S30:** Estimated *k _f_* values for the reaction between *AL*_1_ and ^5^*T*. 15 reaction traces were fitted independently to an ODE model of a second order reaction. Mean and Standard Error of the Mean are reported.

| Trace | $[^5T]$ (nM) added to 10 nM $AL_1$ | $k_f$ ($M^{-1} s^{-1}$) | Mean | SEM |
| --- | --- | --- | --- | --- |
| C02 | 2 | $1.06 \times 10^5$ | | |
| C03 | 4 | $1.13 \times 10^5$ | | |
| C04 | 6 | $1.13 \times 10^5$ | | |
| C05 | 8 | $1.19 \times 10^5$ | | |
| C06 | 10 | $1.22 \times 10^5$ | | |
| C08 | 2 | $1.04 \times 10^5$ | | |
| C09 | 4 | $1.05 \times 10^5$ | | |
| C10 | 6 | $1.08 \times 10^5$ | $1.14 \times 10^5$ | $5.60 \times 10^3$ |
| C11 | 8 | $1.13 \times 10^5$ | | |
| C12 | 10 | $1.19 \times 10^5$ | | |
| D02 | 2 | $1.14 \times 10^5$ | | |
| D03 | 4 | $1.11 \times 10^5$ | | |
| D04 | 6 | $1.13 \times 10^5$ | | |
| D05 | 8 | $1.16 \times 10^5$ | | |
| D06 | 10 | $1.24 \times 10^5$ | | |

**Table S31:** Estimated *k _f_* values for the reaction between *BL*_2_ and ^5^*T*. 15 reaction traces were fitted independently to an ODE model of a second order reaction. Mean and Standard Error of the Mean are reported.

| Well | $[^5T]$ (nM) added to 10 nM $BL_2$ | $k_f$ ($M^{-1} s^{-1}$) | Mean | SEM |
| --- | --- | --- | --- | --- |
| C02 | 2 | $1.24 \times 10^5$ | | |
| C03 | 4 | $1.05 \times 10^5$ | | |
| C04 | 6 | $9.54 \times 10^4$ | | |
| C05 | 8 | $9.31 \times 10^4$ | | |
| C06 | 10 | $9.10 \times 10^4$ | | |
| C08 | 2 | $1.21 \times 10^5$ | | |
| C09 | 4 | $1.07 \times 10^5$ | | |
| C10 | 6 | $9.93 \times 10^4$ | $1.04 \times 10^5$ | $2.87 \times 10^3$ |
| C11 | 8 | $9.70 \times 10^4$ | | |
| C12 | 10 | $9.54 \times 10^4$ | | |
| D02 | 2 | $1.22 \times 10^5$ | | |
| D03 | 4 | $1.14 \times 10^5$ | | |
| D04 | 6 | $1.04 \times 10^5$ | | |
| D05 | 8 | $1.01 \times 10^5$ | | |
| D06 | 10 | $9.42 \times 10^4$ | | |

**Table S32:** Estimated *k _f_* values for the reaction between *CL*_1_ and ^5^*T*. 5 reaction traces were fitted independently to an ODE model of a second order reaction. Mean and Standard Error of the Mean are reported.

| Trace | $[^5T]$ (nM) added to 10 nM $CL_1$ | $k_f$ ($M^{-1} s^{-1}$) | Mean | SEM |
| --- | --- | --- | --- | --- |
| G02 | 2 | $1.16 \times 10^5$ | | |
| G03 | 4 | $1.18 \times 10^5$ | | |
| G04 | 6 | $1.21 \times 10^5$ | | |
| G05 | 8 | $1.20 \times 10^5$ | | |
| G06 | 10 | $1.13 \times 10^5$ | | |
| G08 | 2 | $1.19 \times 10^5$ | | |
| G09 | 4 | $1.20 \times 10^5$ | | |
| G10 | 6 | $1.21 \times 10^5$ | $1.21 \times 10^5$ | $8.1 \times 10^2$ |
| G11 | 8 | $1.24 \times 10^5$ | | |
| G12 | 10 | $1.27 \times 10^5$ | | |
| H02 | 2 | $1.21 \times 10^5$ | | |
| H03 | 4 | $1.21 \times 10^5$ | | |
| H04 | 6 | $1.22 \times 10^5$ | | |
| H05 | 8 | $1.23 \times 10^5$ | | |
| H06 | 10 | $1.25 \times 10^5$ | | |

**Table S33:**
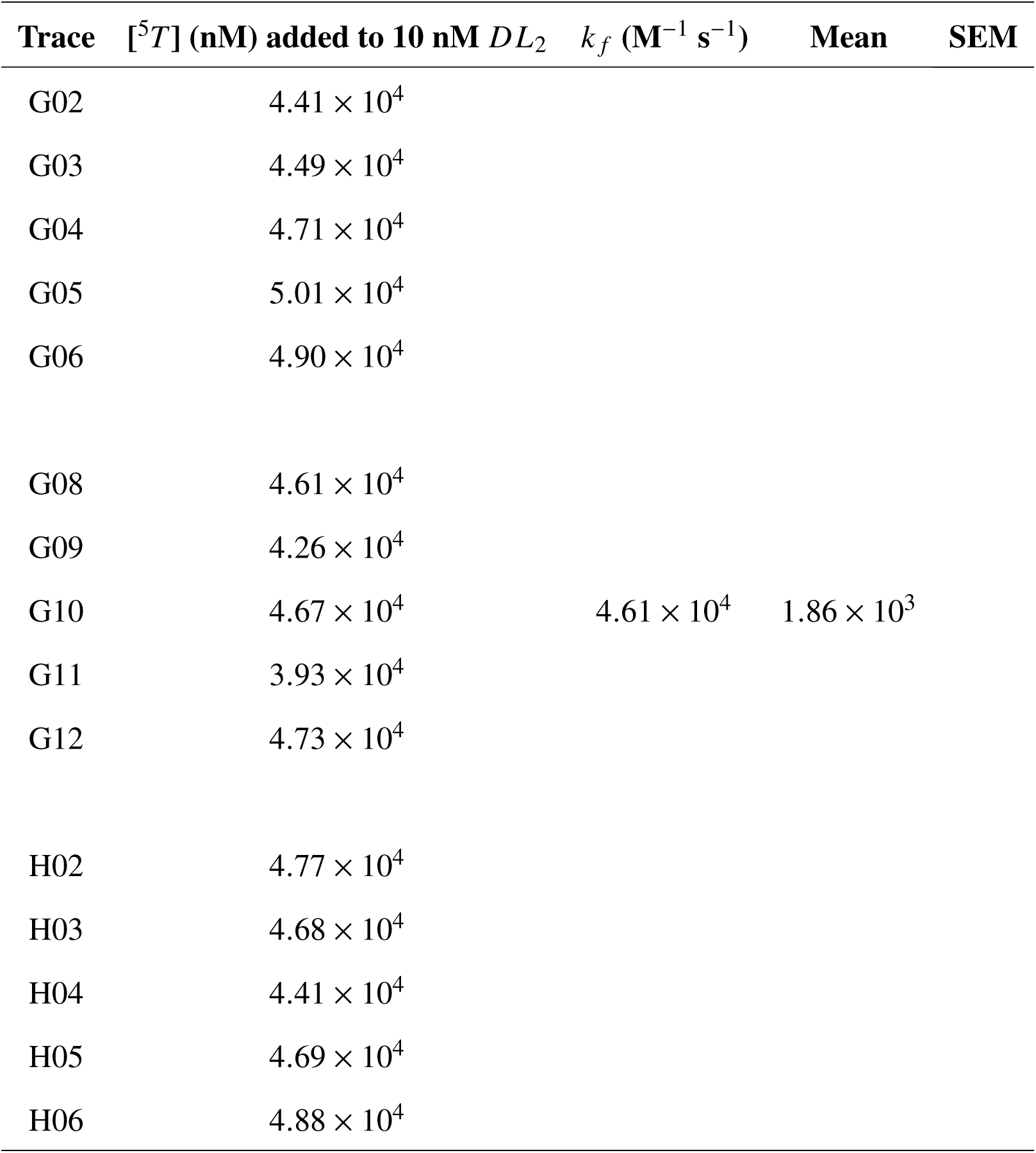
Estimated *k _f_* values for the reaction between *DL*_2_ and ^5^*T*. 15 reaction traces were fitted independently to an ODE model of a second order reaction. Mean and Standard Error of the Mean are reported.

